# Ot2Rec: A Semi-Automatic, Extensible, Multi-Software Tomographic Reconstruction Workflow

**DOI:** 10.1101/2022.12.15.520632

**Authors:** Neville B.-y. Yee, Elaine M. L. Ho, Win Tun, Jake L. R. Smith, Maud Dumoux, Michael Grange, Michele C. Darrow, Mark Basham

**Affiliations:** Artificial Intelligence & Informatics, Rosalind Franklin Institute, Harwell Campus, Didcot, OX11 0FA, UK; Faculty of Medical Sciences, Newcastle University, Newcastle upon Tyne, NE1 7RU, UK; Diamond Light Source Ltd., Diamond House, Harwell Campus, Didcot, OX11 0DE, UK; Structural Biology, Rosalind Franklin Institute, Harwell Campus, Didcot, OX11 0FA, UK; Division of Structural Biology, Wellcome Centre for Human Genetics, University of Oxford, Oxford, OX3 7BN, UK; SPT Labtech, Melbourn Science Park, Cambridge Road, Melbourn, SG8 6HB, UK

**Keywords:** cryo-electron tomography, image processing, cryo-electron microscopy

## Abstract

Electron cryo-tomography (cryo-ET) is an imaging technique for probing 3D structures with at the nanometre scale. This technique has been used extensively in the biomedical field to study the complex structures of proteins and other macromolecules. With the advancement in technology, microscopes are currently capable of producing images amounting to terabytes of data per day, posing great challenges for scientists as the speed of processing of the images cannot keep up with the ever-higher throughput of the microscopes. Therefore, automation is an essential and natural pathway on which image processing – from individual micrographs to full tomograms – is developing. In this paper, we present Ot2Rec, an open-source pipelining tool which aims to enable scientists to build their own processing workflows in a flexible and automatic manner. The basic building blocks of Ot2Rec are plugins which follow a unified API structure, making it simple for scientists to contribute to Ot2Rec by adding features which are not already available. In this paper, we also present three case studies of image processing using Ot2Rec, through which we demonstrate the speedup of using a semi-automatic workflow over a manual one, the possibility of writing and using custom (prototype) plugins, and the flexibility of Ot2Rec which enables the mix-and-match of plugins. We also demonstrate, in the supplementary information, a built-in reporting feature in Ot2Rec which aggregates the metadata from all process being run, and output them in the Jupyter Notebook and/or HTML formats for quick review of image processing quality. Ot2Rec can be found at https://github.com/rosalindfranklininstitute/ot2rec.

**Impact Statement:** The field of cryo electron tomography has grown substantially in recent years, bringing about new advances in hardware and software which enable visualisation of cell and tissue architecture and proteins found in their native context. These same advances have, in some ways, stratified the field into those with access and those without. On the software side, this has emphasised the need for open-source options that do not require high levels of computational literacy to access. Additionally, it has highlighted the need for ways to both mix-and-match software for easy prototyping and comparisons between parameters and methods. Ot2Rec addresses these needs through a simple, unified plugin structure allowing the addition of existing software or the development of new and does so in a way which democratises access.

## 1. Introduction

### 1.1. Cryo-ET & data acquisition

Electron cryo-tomography (cryo-ET), is an imaging technique that produces three-dimensional outputs in the nanometre resolution range^(1,2)^. In the biomedical field, it has been used to study heterogeneous purified proteins using sub-tomogram averaging^(3,4)^, structures within thin or smaller cells^(5)^, the edges of larger cells^(6)^, and more recently in conjunction with cryo focused ion beam milling (cryoFIB), sub-cellular structures and proteins^(7–9)^.

The sample must first be vitrified using either traditional plunge freezing methods ^(10)^ or high pressure freezing for larger samples^(11)^. In some cases, the frozen sample is subsequently thinned by cryosectioning^(12)^ or cryoFIB milling ^(13–15)^). These cryogenic specimen preparation techniques alleviate the need for chemical fixatives and stains which are commonly used to preserve biological structures and bolster contrast in room temperature volume electron microscopy (vEM) techniques^(16,17)^.

During data collection images are taken at a series of angles by tilting the specimen, typically in the range of approximately −60° to +60° with a 2° to 5°step between tilts, though many tilt acquisition schemes exist to ration exposure of the specimen^(18)^. In all cases, a portion of information is not collected due to mechanical constraints of the microscope stage and increasing specimen thickness during tilting. This leads to a missing wedge or cone of information in Fourier space and an elongation of the resultant reconstructed data in the direction of the electron beam in real space^(19)^.

### 1.2. Current image processing workflow for cryo-ET

Several stages of image processing tasks are required to produce tomograms from raw micrograph movies (Figure 1). Many software tools exist for each stage, which have been reviewed recently by Pyle and Zanetti^(20)^. Raw micrograph movies at each tilt angle are first motion-corrected to compensate for specimen movement during acquisition. Individual frames of the movies recorded at each angle are aligned and averaged to produce a single image for each tilt angle. These motion-corrected images are sorted in order of their tilt angles to produce a tilt series. Tilt series projections are then aligned to ensure the rotation axis is consistent between projections. This can be performed with or without fiducial markers. At this stage, the contrast transfer function (CTF) can be optionally estimated from the aligned tilt series, and deconvolved with the tomogram at later stages to improve the image quality. Aligned projections are then reconstructed into tomograms, which provide the 3D representation of the specimen. Post-processing strategies can then be applied to the tomograms, which will be dependent on the specific research question. This often involves sub-tomogram averaging (STA) to obtain high-resolution structures of repeating structures in the tomogram, or segmentation to determine spatial relationships between structures.

**Figure 1.**
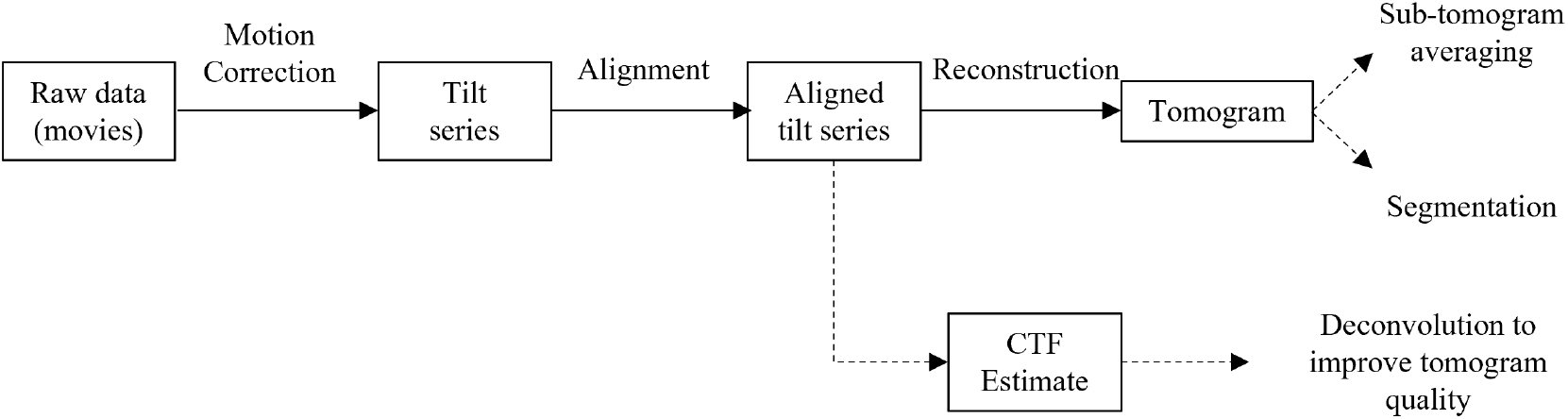
Typical image processing tasks for reconstructing tomograms from raw micrograph movies. Key: boxes - datasets, solid arrows - processes necessary for tomogram reconstruction, dotted arrows - optional processes.

### 1.3. Existing cryo-ET pipelines

The image processing tasks required to reconstruct tomograms from the movies are generally consistent for different experiments, though the post-processing stages afterwards often require more customisation. However, many software tools exist for each stage of the reconstruction process, each with their own disparate data structures, standards, and user interfaces. Users must learn how to use each individual tool and adapt their data structures to fit the specific software package, increasing the barrier to trying alternative tools which may have better performance or functionality. Therefore, pipelines which manage the data flow and provide a unified interface for several software tools would enable easier combination and trialling of the different software tools to optimise reconstruction outcomes.

A few pipelines already exist to automate the reconstruction process within a single framework, to varying degrees of customisation (Table 1). Generally, these pipelines were developed to automate reconstruction for datasets collected within a single research group, and thus, do not enable substitution of different tools developed elsewhere. These pipelines include TomoBear^(21)^, TomoRobot^(22)^, EMAN2^(23)^, and Warp^(25)^. Scipion 3.0 gives users a wide range of software tools to choose in their framework and offers a user-friendly workflow builder in their graphical user interface (GUI). However, accessing the Scipion tools programmatically is not straightforward, so automation of the process is more challenging.

**Table 1.**
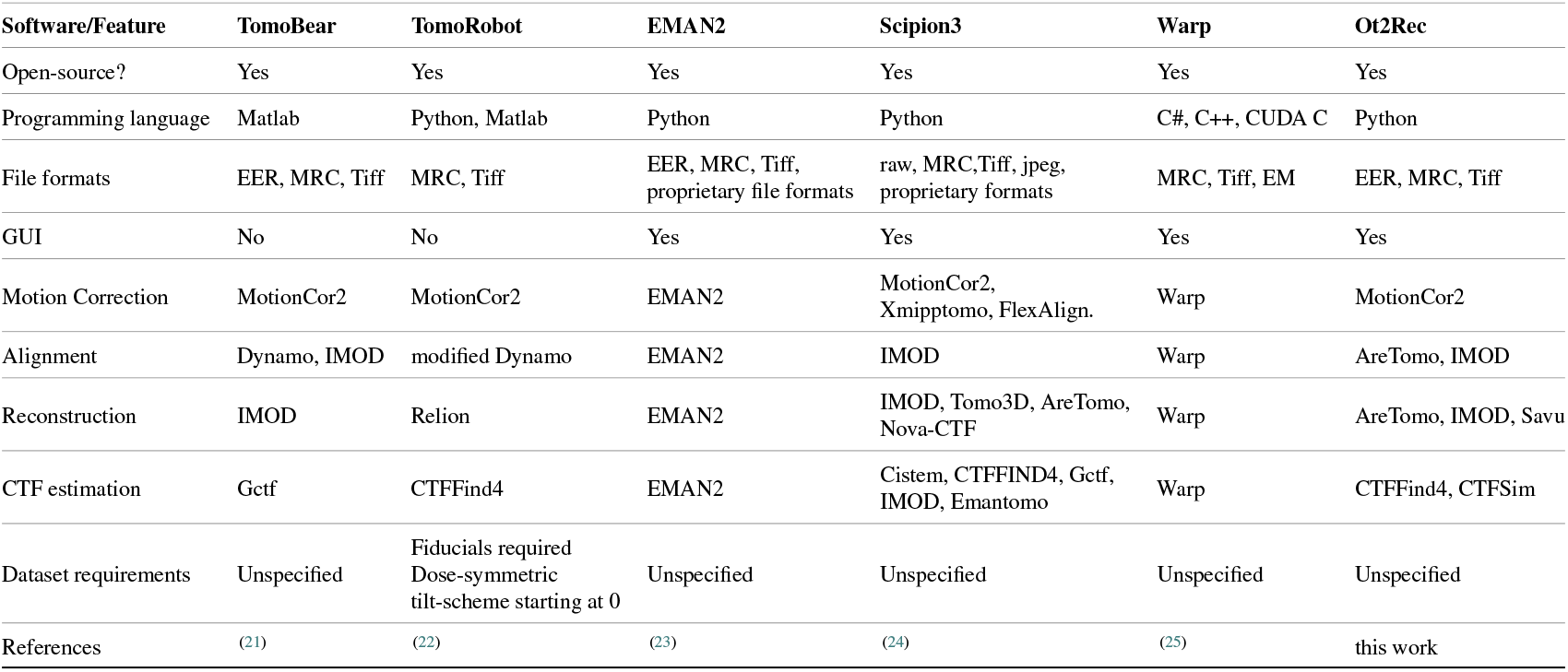
Comparison of cryo-ET reconstruction pipelines.

### 1.4. Motivation

Here, we present Ot2Rec, a pipeline for reconstruction of cryo-ET tilt series which allows users to combine different software tools within a single framework. Ot2Rec can be used via a GUI but can also be automated programatically if required. Ot2Rec is easily extensible through its plugin architecture, and published open-source under the Apache v2.0 license. Ot2Rec is developed as an open-source project with a strong focus on user involvement. We welcome contributions from users and developers alike, our contributors guide is available [on our Github Wiki](https://github.com/rosalindfranklininstitute/Ot2Rec/wiki/Contributing-Guide). A written tutorial guiding users through reproducing one of our case studies is available in the Supplementary Information, in addition to a guide to writing Ot2Rec plugins for developers. These documents are also online on our wiki at https://github.com/rosalindfranklininstitute/ot2rec/wiki.

## 2. Development philosophy & features

Ot2Rec was developed to achieve three key characteristics of a tomography reconstruction pipeline, with the ultimate aim of fully automating tomography reconstruction to obtain high quality tomograms. Firstly, the pipeline had to offer different routes to combine tools for each stage of tomogram reconstruction. The pipeline had to be user-friendly and accessible to users without programming experience. Finally, a mechanism to evaluate all tilt series at a glance was also required. These functions together provide the framework for which a completely automated tomogram reconstruction pipeline can be built, where the optimum combination of software tools and parameters can be automatically selected and applied to several tilt series at once, with minimal user intervention. The following sections describe the design choices that were made towards achieving this functionality in Ot2Rec.

### 2.1. Different routes to tomogram reconstruction

#### 2.1.1. Plugin architecture

In Ot2Rec, the basic program infrastructure is designed such that each feature is a standalone plugin, which has minimal interaction with other plugins. Such design is beneficial for developers as a fault in one plugin does not directly cause other plugins to fail, making further development and maintenance easier. This plugin architecture also facilitates extension of Ot2Rec to cover other image processing tools, whilst maintaining easy integration with existing workflows.

Each plugin performs one task within the reconstruction process, e.g., motion correction or alignment. The plugin has two main subroutines, one to capture user input and configure the parameters for the task to be performed, and the other which generates the commands to run the task on all tilt series. Each plugin follows a simple yet unified structure for its application programming interface (API): a Python class which encapsulates all the essential plugin-specific methods, followed by subroutines that enable the plugin to communicate with the Ot2Rec main API. These subroutines are called and executed directly as entry-points by users. This well-defined, simple structure is helpful for developers and users alike as they become accustomed to the patterns of using Ot2Rec.

Plugins were chosen based on the software tools already used by users in our institute. The nine plugins currently available in Ot2Rec 0.2 are:

- Motion Correction
  – MotionCor2^(26)^
- CTF Estimation and Deconvolution
  – CTFFind4^(27,28)^
  – CTFSim (based on^(28)^; described in Section 5.2)
  – RLDeconv (described in Section 5.2)
- Tilt series alignment
  – IMOD^(29,30)^
  – AreTomo^(31)^
- Reconstruction
  – IMOD^(29,30)^
  – AreTomo^(31)^
  – Savu^(32)^

#### 2.1.2. Metadata handling

Metadata are small files used to record the locations and other useful information about actual data being processed. Metadata often plays a key role in a multi-step computational workflow as it defines how the data are linked to each other. Metadata generated from various software and processes are often incompatible (for instance, having different headers or different metadata file formats) creating obstructions in data flow. This is a currently known problem in the electron tomography community – there are many workflows and packages which use different and incompatible metadata formats, making interoperability between separate software packages extremely challenging.

In view of this, Ot2Rec has been designed with a central philosophy of unified metadata structure. To use Ot2Rec for image processing, the user first runs the command `o2r.new` with which Ot2Rec aggregates the master metadata that includes the path to the raw images (micrographs) and the tilt series indices of the individual micrographs.

In subsequent processing steps (i.e. plugins), Ot2Rec stores the metadata files from the individual programs and outputs an independent metadata file. This metadata file is a human-readable easily parsed YAML format that carries essential information for the next steps.

The unified internal metadata system allows workflows to be built flexibly by interchanging the different plugins for each task. Whilst several default workflows are implemented in the package (cf. Fig. 2), for instance, MotionCor2 → IMOD (alignment + reconstruction), or MotionCor2 → IMOD (alignment) → Savu (reconstruction), the “network” of possible routes can continue to grow as more plugins are implemented into Ot2Rec. This flexibility and customisation allows the user to maximise both result quality and computational performance.

**Figure 2.**
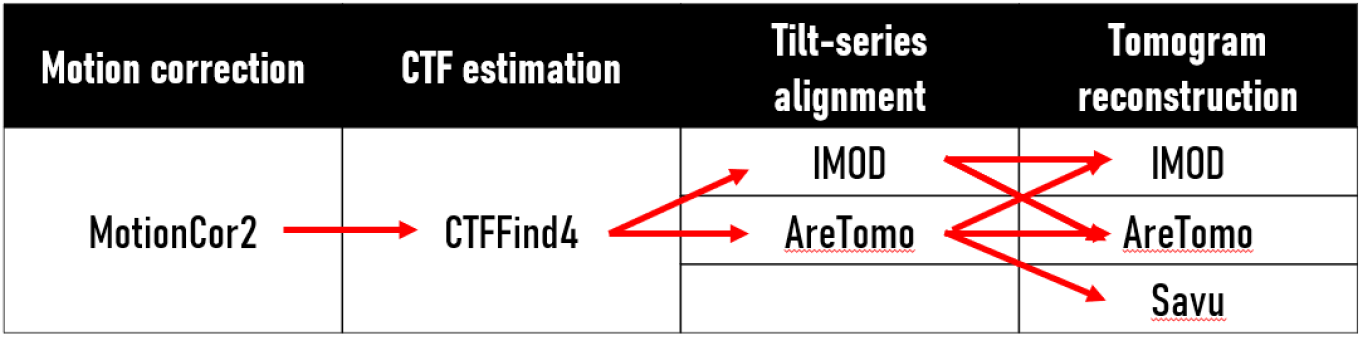
Standard workflows implemented in Ot2Rec. Red arrows denote possible branches of the workflow, which currently covers all permutations of existing implemented plugins (though not necessarily all available options within a software e.g., IMOD).

As image processing programs each have different parameters, the quality of the output and computational performance can vary vastly depending on the chosen parameters. With a flexible workflow, users can choose the tools that best suit their purposes and test different parameters provided by the same tool. They can also compare the results using the reporting feature of Ot2Rec. Once the user has completed a comparative study, preferably using a small, representative subset of their data, they can continue to process the rest of their data using the optimal workflow and parameters they have identified.

### 2.2. User-friendliness

Graphical user interfaces (GUI) are a means for a program to communicate with the user. Its usage can range from the collection of essential parameters to the interactive display of results.

In Ot2Rec, a Linux-based program, since the building blocks are the individual plugins which have a wide range of parameters as inputs, a GUI is necessary to remove the use of command-line flags. For this reason, we have implemented MagicGUI^(33)^ as Ot2Rec’s GUI for gathering parameters from users. One of the biggest benefits of using a simple GUI over the command-line is that parameters can be presented in a more human-readable and descriptive way rather than using internal variable names, which in many cases need to be brief. Another benefit of using a GUI as the first point of communication between the user and the program is that since the parameters are more descriptively presented, the chance of incorrect inputs can be reduced, as additional information can be displayed alongside the parameter inputs for explanation.

### 2.3. Report generation

Comparison of the outputs of different tomogram reconstruction workflows is facilitated in Ot2Rec by automatically generated reports, which contain a workflow diagram describing the plugins that have been applied, selected input parameters, and performance metrics for each plugin. This report is generated for the entire project, allowing the user to evaluate the reconstruction process for all tilt series in the project at a glance. These reports can be generated in a document, slideshow or Jupyter notebook format^(34)^, allowing users to customise how they would like to interact with the report. Reports for all the case studies in this work are included in the Supplementary Information as examples.

## 3. Case Studies

### 3.1. Case Study 1: Semi-automatic processing of large datasets

Tomographic image processing is often performed manually, with users processing one tilt-series at a time. Scripts can be written to automate some of the procedures especially for those being carried out using command-line programs, such as MotionCor2, though this can be difficult and frustrating for many users without a programming background. Ot2Rec enables users to automatically set up and apply a reconstruction workflow to entire datasets of several tilt series at once, without any scripting or individual processing of datasets. A report can be automatically generated at the end for the user to evaluate the performance of the reconstruction process on all tilt series at a glance.

Here, we demonstrate the use of Ot2Rec to reconstruct tomograms from the EMPIAR-10364 dataset^(35)^. This dataset consisted of 17 tilt series of *E. coli* minicells acquired on a Titan Krios at the electron Bio-Imaging Centre, Diamond Light Source. Each tilt series had 60 projections from −60° to +60° in 2° increments. The movie taken at each tilt angle contained 5 frames. Images were binned by a factor of 2 at the alignment step for faster processing, and no binning was applied in reconstruction for a final bin factor of 2x between the raw data and tomogram. Motion correction was performed with MotionCor2, and alignment and weighted back-projection (WBP) reconstruction were performed with IMOD. See Section 5 for more details.

Ot2Rec substantially reduced the amount of user intervention required to process tomograms as configured workflows are automatically applied to all tilt series in the dataset. Typically, the user would have to take each individual tilt series through several software packages, which is feasible for experiments with a few tilt series, but quickly becomes a bottleneck when scaling to dozens or hundreds of tilt series. This case study shows that typical reconstruction workflows with popular software tools can be deployed with Ot2Rec for large datasets, with the potential for automated parameter tuning to determine the optimum reconstruction workflow in the future.

The report generated summarised the performance of each step in the reconstruction process. First, a diagram showing the processes run on the dataset is shown (Figure 3a). The shifts between frames reported by MotionCor2 were within 1–2 pixels and fairly uniform across all tilt series (Figure 3b). The average shifts between patches from the alignment patch-tracking step were between 10 and 38 Angstroms, and all tilt series had relatively similar shifts (Figure 3c). The report also contains representative thumbnails of the central x-y, x-z, and y-z slices of every tomogram, allowing users to visually assess all tomograms at once. The full report for this case study is available in the Supplementary Information.

**Figure 3.**
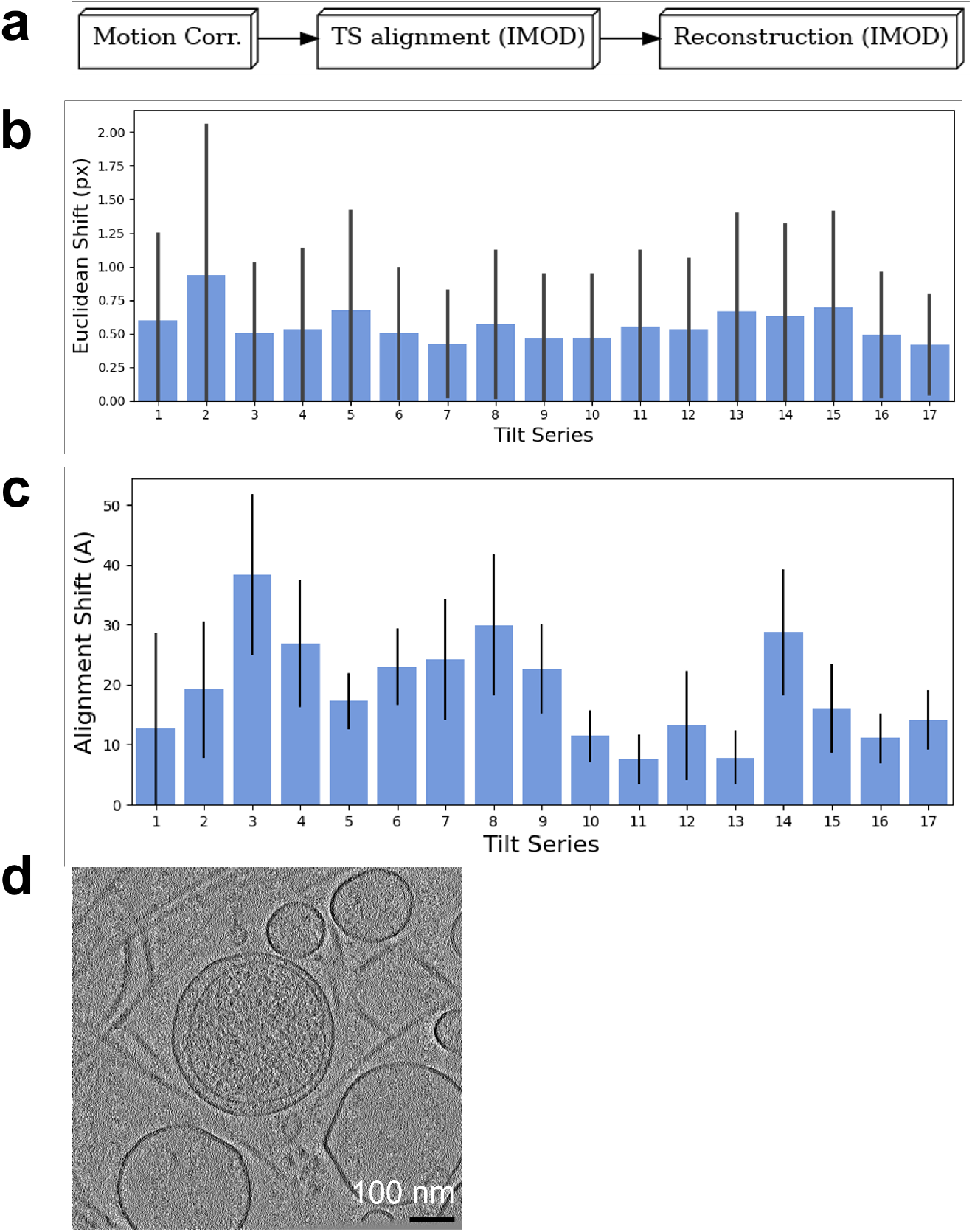
All tilt series in EMPIAR-10364 could be reconstructed at once in Ot2Rec and evaluated at a glance with the automatically generated report, some sections of which are highlighted below. (a) Workflow diagram for processes performed in Case Study 1. (b) Means and standard deviations of motion correction shifts between movie frames for all tilt series. (c) Means and standard deviations of alignment shifts between patches (L_2_-norm) for all tilt series. (d) Central x-y slice of tomogram from tilt series 18 in EMPIAR-10364, with a Gaussian blur filter applied (σ = 2.0 px ≈ 8.972Å). The full report is included in the Supplementary Information.

### 3.2. Case Study 2: Demonstration and testing of prototype features

Implementing custom plugins in Ot2Rec enables rapid development and deployment of bespoke algorithms and tools which can easily be integrated into existing tomogram reconstruction workflows. This case study demonstrates the results of a custom plugin called CTFSim, and how this bespoke tool was used in conjunction with other existing Ot2Rec plugins to enhance the quality of the reconstructed tomograms.

This case study investigated the use of deconvolution to improve contrast with the CTF estimated from CTFSim on tomograms of human choriocarcinoma cells (Figure 4). Images were acquired on a Titan Krios at the electron Bio-Imaging Centre, Diamond Light Source. Each tilt series had 41 projections taken at −53° to +27° degrees in increments of 2°. Ot2Rec was used for motion correction, alignment, and reconstruction of the tomograms for all tilt series in the dataset. The tomograms were then deconvolved using the CTF estimated with the custom Ot2Rec plugin CTFSim (based on^(28,47)^; see Methods for details).

**Figure 4.**
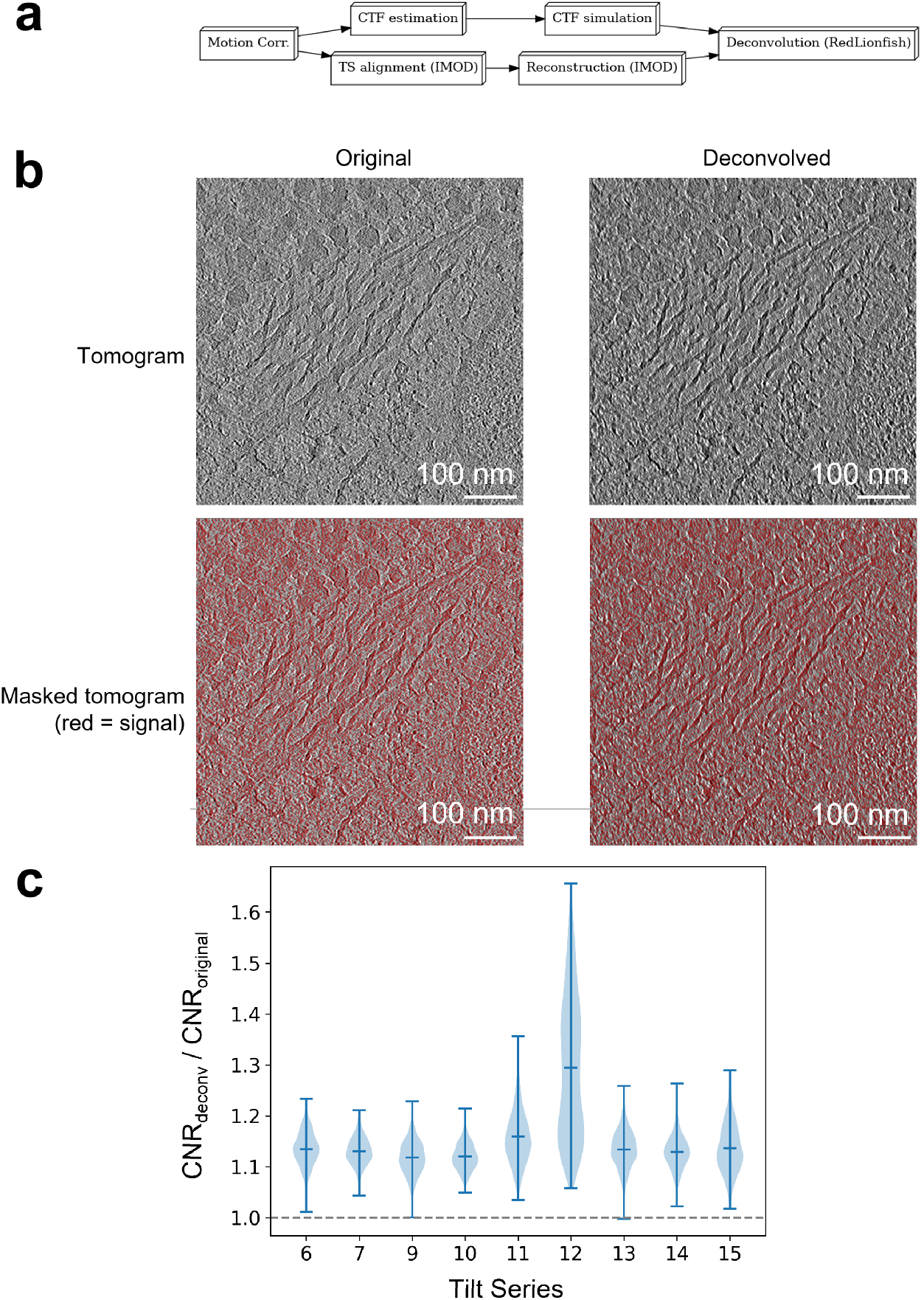
PSF simulation and deconvolution implemented as custom plugins in Ot2Rec can be easily integrated into existing image processing workflows. Deconvolution improved image quality of all tomograms in this case study of human choriocarcinoma cells.(a) Workflow diagram for processes performed in Case Study 2. (b) Thumbnails showing an example original and deconvolved tomogram (tilt series 9), unmasked (top row) and masked (bottom row). The masked regions were used to calculate the CNR. (c) Violin plot showing a general boost (> 1) in contrast-to-noise ratios after deconvolution.

The improvement in image quality was measured using the contrast-to-noise ratio (CNR)^(36)^. Briefly, the signal was segmented from the background by thresholding the grey values with the automatically determined Otsu’s threshold^(37)^. The grey value distributions in the segmented signal and background regions were used to calculate the CNR.

The deconvolved tomograms showed improved contrast compared to the original, as shown visually and as measured by improved contrast-to-noise ratios (CNR) in the deconvolved images (Figure 4). CNR was higher in the deconvolved tilt series compared to non-deconvolved, with the largest increase in CNR seen in tilt series 12 with deconvolution increasing CNR by 65.6% over the original tomogram.

### 3.3. Case Study 3: Optimisation of tomogram reconstruction workflow

Reconstruction of tomograms is a multi-stage process, where the final tomogram quality is affected by the results of earlier stages. Therefore, optimisation of the entire workflow at each stage is necessary to obtain the best results. Ot2Rec provides the framework for this optimisation by enabling users to mix-and-match different software tools on the same tilt series, and compare these results in a single report.

In this case study, different combinations of alignment and reconstruction tools were applied to the same dataset, and Ot2Rec reports were generated to compare the performance of each (Figure 5). This case study consisted of 23 tilt series composed of 41 projection images each ranging from −60°to +60°degrees.

**Figure 5.**
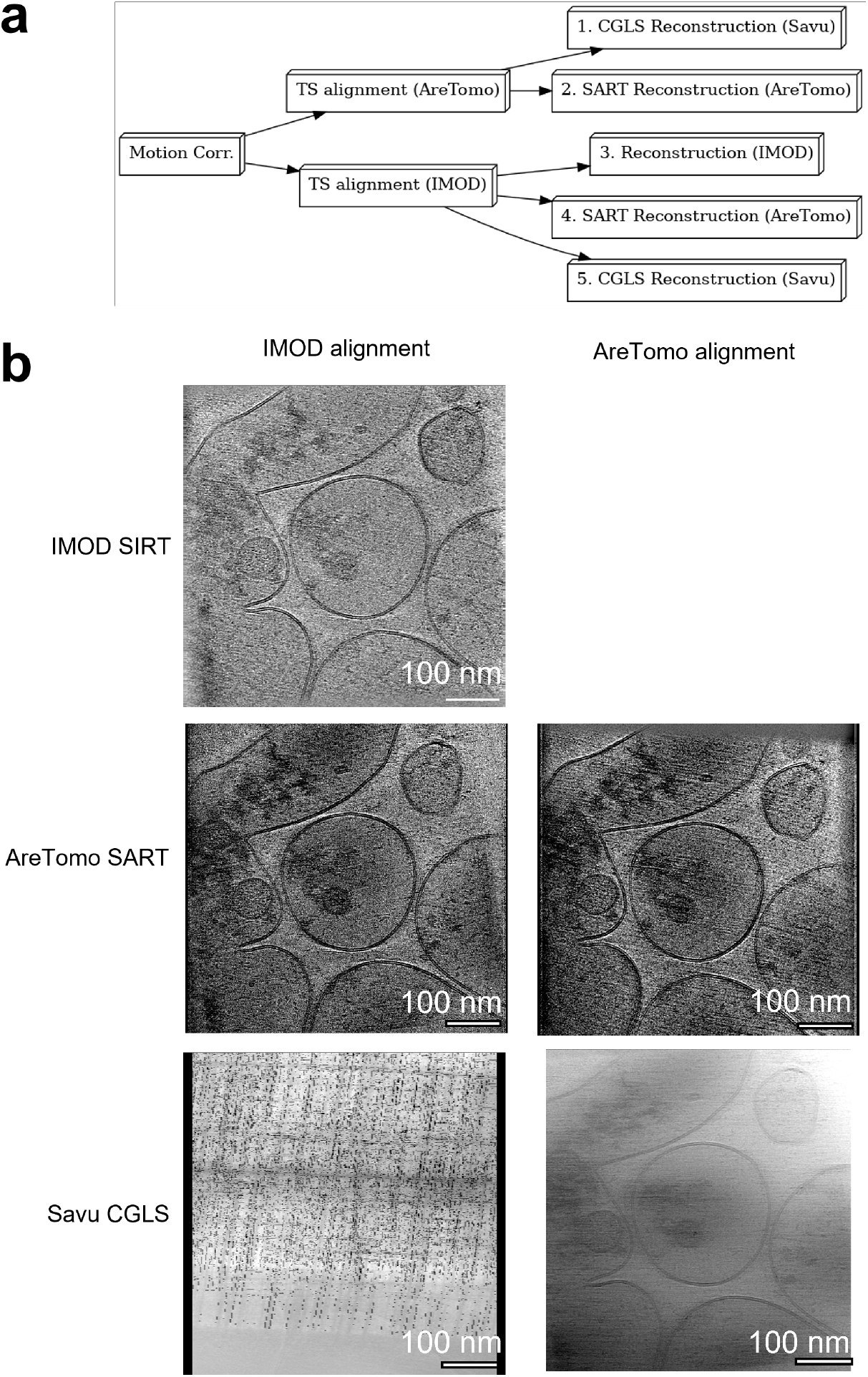
Different combinations of IMOD, AreTomo, and Savu for alignment and reconstruction yielded varying results. Five combinations (1 - 5) were tested, as shown in the workflow diagram a. NB. Iterative reconstruction with CGLS in Savu (Route 5) failed due to poor alignment results from IMOD alignment. However, the same reconstruction method produced a good quality reconstruction when the AreTomo alignment method was used instead (Route 1) (a) Workflow diagram used in Case Study 3, (b) Central x-y slice of the tomogram reconstructed with IMOD, AreTomo, and Savu on IMOD and AreTomo aligned data. Note that IMOD reconstruction with AreTomo aligned data is not available on the current version of Ot2Rec, but will be supported in later versions. A movie showing this figure in 3D is available in the Supplementary Information (Movie S1).

After motion-correction, fiducial-less alignment of the tilt series was performed with either IMOD or AreTomo, followed by reconstruction with either IMOD simultaneous iterative reconstruction technique (SIRT), AreTomo simultaneous algebraic reconstruction technique (SART), or Savu conjugate gradient least squares (CGLS) reconstruction. Different combinations of alignment and reconstruction methods were expected to produce different results, which will be investigated in this section. The full reports for this case study are available in the Supplementary Information.

The shifts reported at the motion correction stage were found to be much larger for tilt series 13 onwards (Figure 6a), indicating a potential issue with tilt series acquisition from this and future datasets in the experiment. Shifts between patches in alignment were also larger overall for tilt series 13 onwards compared to the earlier datasets (Figure 6b). Inspection of the tilt series data showed obstructed views at high tilt angles for those affected by large motion correction shifts. In the future, Ot2Rec could include metrics to determine low quality tilt angles to be excluded.

**Figure 6.**
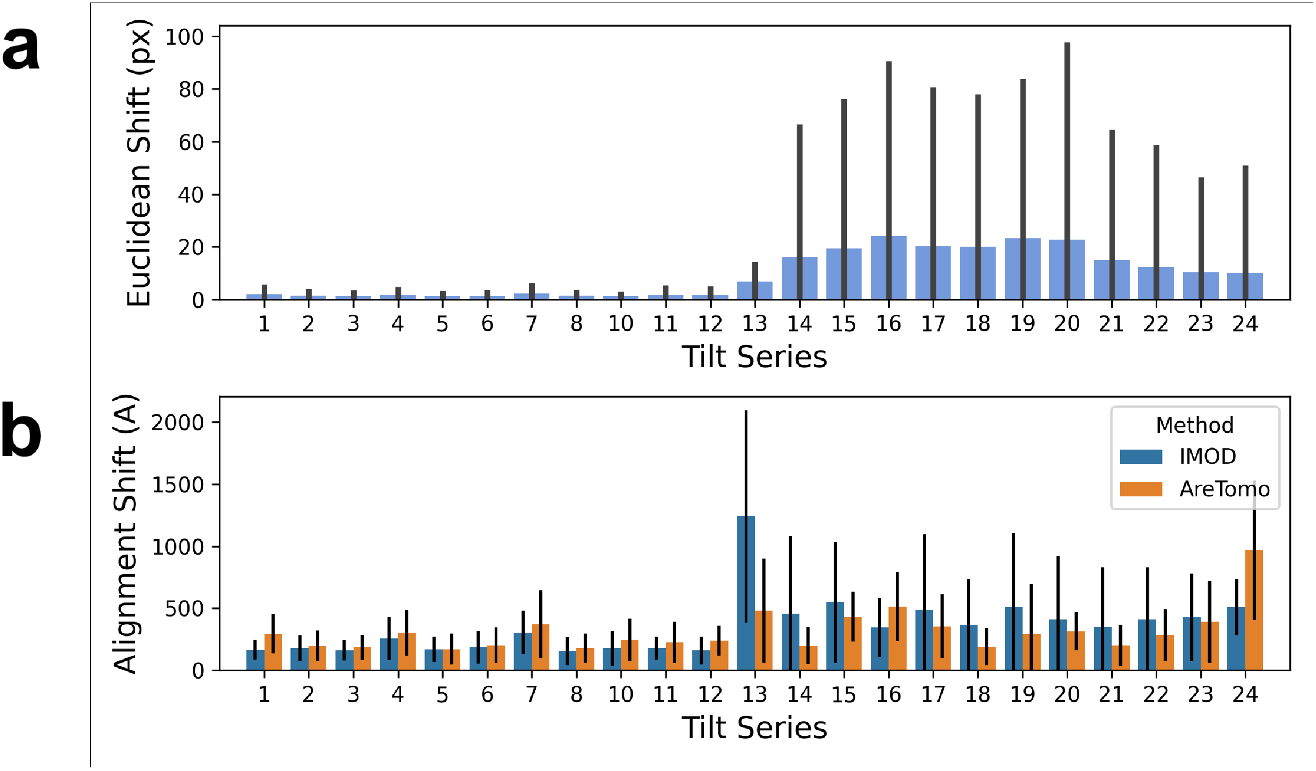
Evaluation of all tilt series shifts from motion correction and alignment at once show substantially larger shifts from tilt series 13 onwards. (a) Euclidean shifts reported by motioncor2 for all tilt series in Case Study 3 (Sect. 3.3) show a large increase from tilt series 13 onwards. (b) Alignment shifts reported by IMOD and AreTomo alignment processes. The alignment shift here is the Euclidean distance between patches which are recorded as metadata from IMOD and AreTomo directly. In both (a) and (b), The bar plots represent the mean and the error bars are the standard deviations in shifts.

An example tomogram (tilt series 18) affected by the large shifts observed in motion correction was chosen to demonstrate differences in outcomes associated with each workflow (Figure 5b). Savu CGLS reconstruction failed with the IMOD aligned data, but not with the AreTomo alignment, and the resultant tomogram had reasonable image quality. However, the CGLS reconstruction had lower contrast than the SIRT or SART reconstruction methods of IMOD or AreTomo. Reconstruction with AreTomo’s SART was successful for both alignment methods (IMOD and AreTomo) with very few visual differences observed between the two outcomes. The IMOD SIRT reconstruction using IMOD alignments also produces a reasonable output which is visually similar to the AreTomo SART reconstructions created from either IMOD or AreTomo alignments. IMOD reconstruction using AreTomo alignments are not currently supported in Ot2Rec, but will be in the future.

## 4. Discussion

Ot2Rec enables reconstruction of tomograms from raw micrograph movies using a range of software tools, all within a single framework. This pipeline is distinct from available alternatives as users can choose different combinations of software tools to suit their specific experiment and automatically apply their workflow to several tilt series at once. Metadata is recorded at each stage and is easily retrievable from text files, and the performance of the overall reconstruction workflow is summarised in a human-readable, automatically generated report.

### 4.1. Future work

The case studies in this work have demonstrated that there are differences in the resultant tomograms from different reconstruction workflows. The reports generated by Ot2Rec also incorporate performance statistics from the individual plugins, e.g., shifts between patches in fiducial-less IMOD tilt series alignment. However, independently measured quality metrics for each stage of the workflow would enable a more equitable comparison of different software tools, so the user (and eventually an algorithm) can quantitatively choose the best combination of software tools and the optimum input parameters for each. The next steps for improving the Ot2Rec reporting system will include development and implementation of such metrics. For example, Fourier Ring Correlation between reconstructions of even and odd projections is a commonly adopted, though computationally intensive, measure of resolution in final tomograms, which could be used to compare reconstruction tools if they are applied on the same aligned tilt series^(38)^. In the future, tomograms could be reconstructed with a pre-defined range of software tools and selected input parameters, and the best overall workflow could be determined quantitatively and then applied to other tilt series in the experiment.

In the case study from Section 3.3, the difference in motion correction shifts between the first 12 and the remaining tilt series was easily observed from the plot of shifts for all tilt series in the project. Observing performance of the tomogram reconstruction process over time can be used as an indication of the quality of the upstream processes (sample preparation, data collection, microscope health) - enabling facility managers to capture valuable data to correct issues at these stages efficiently. Performance in specific metrics could help troubleshoot potential issues, for example, consistent large deviations between calculated and input tilt angles from the alignment process could be a sign of issues with the microscope stage. Issues with poor quality images at specific tilt angles could also be detected and these images excluded from subsequent analysis.

Case Study 2 has demonstrated that custom image processing tools developed within the Ot2Rec framework can be integrated easily into existing image processing workflows, which is beneficial to both developers and users. For developers, maintenance and deployment efforts are significantly reduced as Ot2Rec already provides implementations for file handling of all intermediate steps in the tomography process. For users, customised image processing tools can be accessed quickly and tested alongside their existing workflows, all within a familiar framework.

Ot2Rec will also be extended to perform other cryo-ET processing tasks, e.g., denoising tomograms, 3D particle picking or sub-tomogram averaging. Some of these tasks will be supported in Ot2Rec by developing new plugins for existing software tools, or if no suitable tools are available, bespoke plugins can be developed which would then be available in conjunction with all the standard plugins in Ot2Rec. More work is needed to adapt the Ot2Rec architecture to handle iterative tasks which combine steps, e.g., simultaneous CTF correction and reconstruction as implemented in NovaCTF^(39)^ by Obr and colleagues^(40)^, or iterative alignment and reconstruction^(41)^. And further work is needed to enable a more automated approach that combines multiple plugins into a workflow that can be launched with a single command.

### 4.2. Conclusion

Ot2Rec currently includes the canonical initial steps of tilt series reconstruction, necessary for either sub-tomogram averaging or segmentation. Alongside of this, it also contains prototype plugins allowing for deconvolution of tomograms to enhance contrast. Ot2Rec acts both as an open-source wrapper with a simple unified plugin structure and a developer and user friendly prototyping tool for new plugins. The reporting functionality provides easy-to-access information, enabling comparisons and optimisation of parameters and software packages. The further development of this pipeline opens the door to options such as automated data processing parameter tuning based on the purpose of data collection or automatic monitoring of upstream steps such as microscope health. Continued open-source cross-software development of the cryoET pipeline with a focus on ease-of-interaction for both the user and developer is the future of cryoET data processing.

## 5. Methods

### 5.1. Sample preparation and image acquisition

#### 5.1.1. Case Study 1

Details of the sample preparation and image acquisition protocols are available in the original publication for the EMPIAR dataset^(35)^.

For this study, motion correction of the micrographs was performed using MotionCor2, followed by a fiducial-less (patch tracking based) tilt series alignment using IMOD. The patch-tracking algorithm was configured such that there were 24 ×24 patches, each patch with the dimensions of 210 ×203 pixels, allowing a 15% overlap between patches. The aligned stacks were binned by a factor of 2 before tomogram reconstruction, which was also performed using the batchruntomo tool in the IMOD suite using the default weighted back-projection (WBP) algorithm. The unbinned thickness of the volume was set to 1920 px (≈861Å), such that after the overall factor-2 binning, the dimensions of the output tomograms are 1920 × 1856 × 1920 pixels, with the pixel spacing of 4.486Å in all dimensions. Specific parameters used in Ot2Rec for this case study are in Table S1.

In all case studies presented in this paper, processing was performed on a virtual machine with 12 Intel Xeon Gold 5218 2.30GHz CPUs and 1 Tesla V100 GP.

#### 5.1.2. Case Study 2

JEG-3 human choriocarcinoma cell line was purchased from Merck Life Science (UK) and cultured in Dulbecco’s modified Eagle’s media (DMEM/F-12, HEPES, Gibco, 11330057) supplemented with 10% fetal bovine serum (Gibco, 10500) and 1% Penicillin-streptomycin (10,000U/ml, Gibco, 11548876). Cells were cultured in T75 flasks and incubated at 37°C with 5% CO_2_.

200 mesh gold holey carbon grids (either R2/2 Quantifoil or R3.5/1 Quantifoil, Agar scientific) were sterilised by dipping into 100% ethanol for 2-5min and rinsing with phosphate buffered saline (PBS) (Gibco, 10010). The grids were coated with 30μl of 1:20 fibronectin (Merck, F0895) in PBS, and kept in the incubator for 2 hours. The grids were then rinsed with PBS two times and put in a petri dish containing JEG-3 media, and incubated at 37°C and 5% CO_2_ for overnight.

Prior to cell seeding, the grids were transferred into a 6-well plate (one grid per well) containing fresh JEG-3 media. A total number of trypsinised JEG-3 (3×10^4^ cells) were seeded directly onto each grid. The cells were then allowed to settle in the incubator for 2-3 days until they were well spread on the grids.

For vitrification, the GP2 (Leica) was used. The humidity-controlled chamber was set to reach >80% humidity. After the grid containing the cells was placed into the GP2, a 2.5 *μ*l JEG-3 medium droplet was applied onto the EM grid, and the grid was blotted from the reverse side for 6-10sec. The grid was plunged into liquid ethane, and the frozen grids were stored until focused ion beam (FIB) milling was performed.

Prior to the FIB milling, the frozen EM grids were clipped into clip rings (ThermoFisher). Scios scanning electron microscope (ThermoFisher) at the electron Bio-Imaging Centre at Diamond Light Source was used to perform the FIB milling. Gallium ion beams were used to mill the cellular samples to a lamella thickness of 200nm. The ion beam parameters were set to 30kV beam energy, and 50pA-0.3nA beam current (for coarse milling) and 30pA (for fine milling). The grids with FIB-milled lamellae were stored again until cryoET was undertaken using a Titan Krios (ThermoFisher Scientific) at 300kV at eBIC. Each tilt series had 41 projections taken at −53° to +27° degrees in increments of 2° using a dose symmetric imaging scheme^(18)^. Data were collected using a Falcon 4i SelectrisX at 64,000× magnification (1.97 Å/pixel).

Motion correction of the micrographs was first performed with MotionCor2, followed by the estimation of the true defocus values using CTFFind4 on a *per-micrograph* basis. The CTFFind4 outputs were then processed using the CTFSim plugin, an Ot2Rec prototype feature, which also simulated and internally reconstructed the 3D PSF of the tilt series. The PSF volumes were truncated to the central 30 × 30 × 30 pixels as the PSF is fast-declining by nature.

Fiducial-less tilt series alignment was performed on the motion-corrected micrographs using IMOD. The patch-tracking algorithm in IMOD was configured such that there were 24 × 24 patches, each patch with the dimensions of 224 pixels squared, allowing a 15% overlap between patches. The aligned stacks were binned by a factor of 8 before tomogram reconstruction. Reconstruction was also performed using the batchruntomo tool in the IMOD suite, and the default weighted back-projection (WBP) was used. The final tomograms have a thickness of 1496Å, which translates to an overall dimensions of 512 × 512 × 1000 pixels, given an overall binning factor of 8.

Lastly, the reconstructed tomograms were deconvolved using the RLDeconv tool in Ot2Rec, with the corresponding simulated PSF volumes as deconvolution kernels. Specific parameters used in Ot2Rec for this case study are in Table S2.

The contrast-to-noise ratio (CNR), as defined in Eq. (1)^(36)^, was used to compare the tomograms before and after deconvolution with CTFSim and RLDeconv.

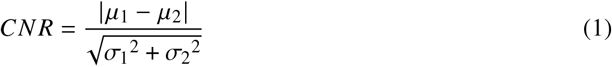

where *μ*_1_, *μ*_2_, *σ*_1_, *σ*_2_ are the average grey-values of the signal, that of the background, the standard deviation of signal grey-values, and that of the background grey-values respectively.

The CNR was calculated from the central 90% of the original and deconvolved volumes in the z-direction. Then for each *z*-slice in a volume, the Ōtsu threshold^(37)^ was calculated and used for separating^1^ the signal from the background (cf. masks shown in red in the second row of Fig 4b). The CNRs of the *i*-th *z*-slices of the two volumes were calculated using Eq. (1). The ratio between the CNR of the deconvolved volume and that of the original tomogram was calculated to gauge the extent of enhancement in image contrast. A value over 1 signifies a positive boost in image contrast after deconvolution.

#### 5.1.3. Case Study 3

Primary cortical neurons of embryonic C57BL/6J mice were dissociated and seeded on Quantifoil^®^ R 2/2 SiO2 Au200 grids that were glow discharged using a GloCube^®^ Plus (Quorum) at 20mA for 30s and treated with 0.1mg mL-1 Poly-D-Lysine (Gibco) at 37°C overnight and rinsed with PBS and left to dry. Cells were seeded onto grids and maintained in Neurobasal^®^ Medium (Gibco) supplemented with B-27TM (Gibco), 1% penicillin/streptomycin (Gibco), 2mM Glutamate (Gibco). 20% of the volume of old media was replaced with fresh media every 3-4 days. Cells were plunge frozen on DIV 14.

Data were collected with a Titan Krios G4 (ThermoFisher Scientific) transmission cryo-electron microscope at 300kV equipped with a cold field emission source gun, a Selectris (ThermoFisher Scientific) electron imaging filter, and a post-imaging filter mounted Falcon 4 (ThermoFisher Scientific) direct electron detector. Tilt series were acquired at 81,000× magnification (pixel size 1.47Å) in a dose symmetric tilt scheme^(18)^ with a tilt range of −60° to 60° imaging at 3° increments, with 2e^-^/Å dose per tilt image. Tilt series were acquired using Tomography software (ThermoFisher Scientific) and collected as EER files^(42)^.

Motion correction was first performed with MotionCor2^(26)^. Next, fiducial-less tilt series alignment was performed with IMOD^(29,30)^ and AreTomo 1.1.0^(31)^. In both cases, a binning factor of 8 was used. The IMOD aligned data was reconstructed with three methods: IMOD simultaneous iterative reconstruction technique (SIRT), AreTomo simultaneous algebraic reconstruction technique (SART), and Savu conjugate gradient least squares (CGLS) reconstruction via the ASTRA reconstruction toolbox^(43–45)^. The AreTomo aligned tilt series was reconstructed with AreTomo SART and Savu CGLS, though IMOD reconstruction will be supported in later versions of Ot2Rec. Specific parameters used in Ot2Rec for this case study are in Table S3.

### 5.2. Prototype features

#### CTFSim

As mentioned previously, the program CTFFind4 is a tool used for estimating the *true* defocus value for each micrograph collected. However, whilst it produces parameters for determining the shape of the contrast transfer function (CTF) associated with the microscope settings, it does not directly have the CTF as a standard output. In view of this, we developed an in-house plugin, named `CTFSim`, to simulate the CTF of the micrograph by reverse-engineering the equations described in the Rohou *et al.* paper^(28,46)^

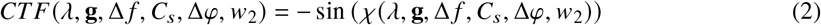

with

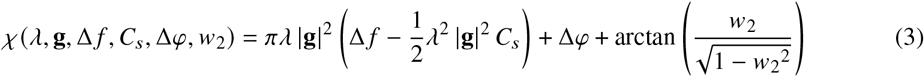

where *λ* is the associated electron wavelength, **g** is the spatial frequency vector, Δ*f* is the defocus value evaluated from CTFFind4, *C_s_* is the spherical aberration, Δ*φ* is the phase shift, and the value *w*_2_ associated with the relative phase contrast *w*_1_ via the relation 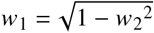.

By definition, the CTF is a function defined in the complex reciprocal space, and is hence rather difficult to visualise and use “as is”. Therefore, an additional feature of the `CTFSim` is to convert the simulated two-dimensional CTF to the point spread function (PSF) in the real space via a direct inverse Fourier transform. Lastly, since the CTF (and PSF) are calculated using the size of the motion-corrected micrographs, these images would have the same size of those micrographs. However, due to the quick-decaying feature of the PSF, the user can choose to truncate the resultant PSF image to a certain size (e.g. 30 pixels squared, by default) in order to facilitate further usage of the obtained results.

Once the per-micrograph 2D PSF profiles for a tilt-series are simulated, `CTFSim` reconstructs the profiles internally using the Weighted Backprojection (WBP) algorithm into a tomogram of the PSF.

As suggested in Croxford *et al.*^(47)^, the *cropped* 3D PSF is post-processed in a three-step process. The PSF tomogram is firstly converted into the 3D CTF with a Fourier transform. Then the pixels are “normalised”^2^ through a division by the value at zero-frequency (i.e. |**g**| =0). Lastly, the normalised CTF stack is converted back to the PSF in real-space via an inverse Fourier transform.

Finally, the processed PSF is then normalised by dividing the array elements by the global maximum of the array. Although it can be derived trivially from eq. 5 that the analytical output of the iterations should be independent of a linear scaling of the PSF, numerically speaking it is still a multi-step process and an unnormalised PSF could potentially cause numerical instabilities especially in the first fraction in eq. 5. Therefore we assert that if the PSF is to be used to deconvolve the raw tomogram via the Richardson-Lucy scheme, this extra operation is necessary.

#### RLFDeconv

The blurring of images due to the optical or electronic properties of the PSF has long been a concern for microscopists. In transmission electron microscopy, the obtained image can be modelled, in the simplest form, as^(48)^

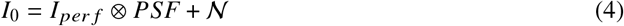

where *I*_0_ is the final output image, *I_per f_* is the perfect image, 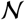 is a stochastic noise term obeying Poisson distribution, and ⊗ denotes the convolution operation. In order to reverse-engineer the perfect image *I_per f_*, an inverse operation must be carried out. We note that Eq. 4 cannot be solved exactly with any analytical or numerical method, since the noise term is unknown and cannot be isolated from the output image. Hence to obtain a reasonable solution, an approximation is necessary. Here a common approximation is that the image signal-to-noise ratio is high enough such that the noise contribution can be ignored.

However, even with such approximation, the inverse of a convolution is still unsolvable by any analytical means, as the solution is non-unique. Therefore one needs to resort to an iterative approach numerically. In 1972 and 1974, Richardson and Lucy separately discovered an iterative scheme^(49,50)^ that gives an approximation approaching an ideal solution as the number of iteration increases. Mathematically, the scheme reads

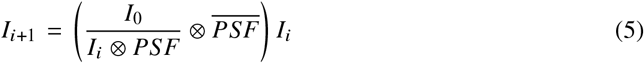

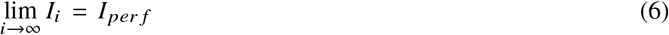

where *I_i_* is the image in the *i*-th step, and the over-bar on *PSF* denotes a flip operation of the *PSF*, hence 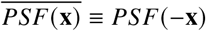.

In Ot2Rec, rather than developing an in-house solver, we have implemented a wrapper plugin for the Python library RedLionfish^(51)^ for performing the 3-dimensional deconvolution between a reconstructed tomogram and a simulated PSF stack (similarly reconstructed as the experimental tomogram). To facilitate the deconvolution of larger stacks, which might cause GPU/CPU memory issues, we also implemented the 3D version of the so-called “block-iterative” algorithm, inspired by Lee^(52)^, which breaks the volume (tomogram) into chunks, on each of which a normal Richardson-Lucy deconvolution is performed independently. The separately deconvolved chunks are then stitched back together. A merit of this method is that due to the massive reduction in the size of deconvolution operands, a much lower memory cost for the GPU/CPU can be achieved. However, the authors would like to emphasise that this block-iterative method must be used with care, as the padding setting in the convolution operations could introduce artefacts around the borders of the chunks as they are stitched back.

### 5.3. Basic usage guide

This short section is dedicated to explain the basics of user interaction with Ot2Rec.

#### 5.3.1. Installation

Ot2Rec is an open-source Linux-based program. Its code repository can be accessed at https://github.com/rosalindfranklininstitute/Ot2Rec.

The easiest way to install Ot2Rec is to download the shell script of the latest release from the Release tab on GitHub and execute the script on the command line.

An alternative method of installation, should the user wish to build the software from source, is to clone the repository to the local environment. As Ot2Rec uses miniconda, it is recommended that the user first create a new conda virtual environment (“venv”) with Python 3.8 (or above) pre-installed. Once the venv is loaded, the user can then use the command `pip install`. in the cloned root folder to automatically install Ot2Rec and its dependencies locally.

### 5.4. Basic usage

Once the virtual environment has been activated, all Ot2Rec command-line functions are available in the terminal, which generates the corresponding GUI for each plugin. All Ot2Rec commands follow a specific format which has been summarised in Table 2.

**Table 2.** Ot2Rec commands and their descriptions.

| Ot2Rec Command | Description |
| --- | --- |
| o2r.new | Creates a new Ot2Rec project. |
| o2r.<plugin>.<function>.new | Creates a new instance of each plugin. For example, o2r.imod.align.new starts a new alignment in IMOD and captures user inputs in the GUI for this plugin. The user parameters are stored in a yaml file. |
| o2r.<plugin>.<function>.run <proj_name> | Runs the commands that perform the plugin's functions on the project where <proj_name>is the project name. For example, o2r.imod.align.run TS runs IMOD alignment on all tilt series in project TS, which has user parameters stored in TS_align.yaml created from o2r.imod.align.new. |
| o2r.report.run <proj_name> --to_html --to_slides | Generate the report for the project <proj_name>as a html document and as slides. Note that this requires the o2r_report environment to be activated (detailed instructions available on Github). |

To start an image processing pipeline with Ot2Rec, the first command to be used is `o2r.new`. With this, a GUI will be displayed, allowing the user to enter some essential project-dependent parameters, with which Ot2Rec will aggregate the master metadata for downstream operations.

For subsequent steps, all the plugins contain a `new` and a `run` function. The `new` commands, like the previous `o2r.new`, prompt the user to input essential parameters for the specified plugin, then collect and pre-propulate the configuration files with metadata from previous steps. The `run` commands look for the relevant configuration files and previous metadata records, and execute the specified plugin. For all `run` operations, the project name (as defined by the user at the beginning with the `o2r.new` command) is required as a command-line argument, as it is used for seeking the correct configuration files.

## Acknowledgements

We acknowledge Diamond Light Source for access and support of the cryo-EM facilities at the UK’s national Electron Bio-imaging Centre (eBIC), funded by the Wellcome Trust, MRC and BBRSC. We also thank eBIC for ongoing collaboration and feedback on Ot2Rec and providing the initial scripts on which Ot2Rec was based.

We also thank the Core Team at the Artificial Intelligence & Informatics Theme at the Rosalind Franklin Institute, especially Drs. Joss Whittle and Laura Shemilt, for their help in configuring the computing infrastructure for running and testing of Ot2Rec.

## Data Availability Statement

The GitHub repository of Ot2Rec is open-source and can be accessed at https://github.com/rosalindfranklininstitute/Ot2Rec. Ot2Rec Report is also open-source and is available at https://github.com/rosalindfranklininstitute/ot2rec_report. All data used in this manuscript can be found on EMPIAR (case study 1: EMPIAR-10364; case study 2: EMPIAR-XXXXX; case study 3: EMPIAR-XXXXX).

## Ethical Standards

The research meets all ethical guidelines, including adherence to the legal requirements of the study country.

## Author Contributions

Software development: NBY; EMLH. Data acquisition: WT; JLRS; MG. Data processing: NBY; EMLH. Writing original draft: NBY; EMLH; WT; MD; MCD; MB. All authors approved the final submitted draft.

## Supplementary Material

- Ot2Rec Report for Case Study 1
- Ot2Rec Report for Case Study 2
- Ot2Rec Report for Case Study 3 - IMOD
- Ot2Rec Report for Case Study 3 - AreTomo
- Table S1: Configuration parameters for Case Study 1
- Table S2: Configuration parameters for Case Study 2
- Table S3: Configuration parameters for Case Study 3
- Guide on writing plugins for Ot2Rec
- Tutorial to reproduce Case Study 1 in Ot2Rec
- Movie S1: Video of 3D reconstructions from the various workflows in Case Study 3

## Footnotes

1 NB. This is only a rudimentary measures to pick out potential signals, and is by no means an attempt to automatically segment useful features from the slice.

2 We acknowledge that this use of “normalisation”, directly quoted from the source, may be an abuse of terminology, as the normalisation of a complex matrix is usually defined as the division by the norm of the matrix of associated moduli, rather than by the value at | **g**| = 0.

