## Supplementary Material for "Ot2Rec: A Semi-Automatic, Extensible, Multi-Software Tomographic Reconstruction Workflow"

---

Contents:

Ot2Rec Report for Case Study 1 — 1

Ot2Rec Report for Case Study 2 — 11

Ot2Rec Report for Case Study 3 – IMOD — 21

Ot2Rec Report for Case Study 3 – AreTomo — 50

Table S1: Configuration parameters for Case Study 1 — 70

Table S2: Configuration parameters for Case Study 2 — 71

Table S3: Configuration parameters for Case Study 3 — 72

Guide on writing plugins for Ot2Rec — 73

Tutorial to reproduce Case Study 1 in Ot2Rec — 81

Movie S1: Video of 3D reconstructions from the various workflows in Case Study 3 — 88

### Workflow diagram

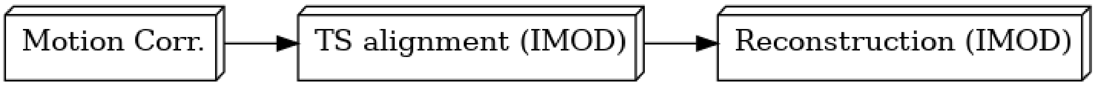


### Motion correction

Motion correction has been performed using MotionCor2.

Please cite:

1. Zheng, S., Palovcak, E., Armache, JP. et al. MotionCor2: anisotropic correction of beam-induced motion for improved cryo-electron microscopy. *Nat Methods* **14**, 331–332 (2017).

#### Summary of shifts for each tilt series

| | **mean** | **std** | **25%** | **50%** | **75%** |
|---|---|---|---|---|---|
| **TiltSeries** | | | | | |
| **1** | 0.597280 | 0.645662 | 0.158114 | 0.460000 | 0.806598 |
| **2** | 0.935666 | 1.117344 | 0.228473 | 0.664831 | 1.245311 |
| **3** | 0.506196 | 0.512616 | 0.152315 | 0.402244 | 0.698140 |
| **4** | 0.529967 | 0.593529 | 0.140357 | 0.410122 | 0.696348 |
| **5** | 0.671132 | 0.738262 | 0.177200 | 0.500100 | 0.882553 |
| **6** | 0.501673 | 0.482156 | 0.155242 | 0.433820 | 0.705762 |
| **7** | 0.423647 | 0.391669 | 0.138924 | 0.368782 | 0.604070 |
| **8** | 0.568803 | 0.542346 | 0.170880 | 0.481041 | 0.798812 |
| **9** | 0.466622 | 0.468324 | 0.136015 | 0.390512 | 0.638905 |
| **10** | 0.468918 | 0.466595 | 0.142127 | 0.399625 | 0.654905 |
| **11** | 0.552081 | 0.562856 | 0.161245 | 0.456946 | 0.763217 |

| | | | | | |
|---|---|---|---|---|---|
| **12** | 0.529845 | 0.521374 | 0.166433 | 0.452769 | 0.754321 |
| **13** | 0.669184 | 0.717781 | 0.182483 | 0.512640 | 0.906091 |
| **14** | 0.634099 | 0.676261 | 0.176918 | 0.494975 | 0.857030 |
| **15** | 0.695593 | 0.703848 | 0.205913 | 0.576281 | 0.968762 |
| **16** | 0.489171 | 0.458651 | 0.152971 | 0.417732 | 0.683520 |
| **17** | 0.417093 | 0.367335 | 0.130384 | 0.377359 | 0.610000 |

```
Out[12]: Text(0.5, 1.0, 'Euclidean Shift Means +/- Standard Deviation')
```

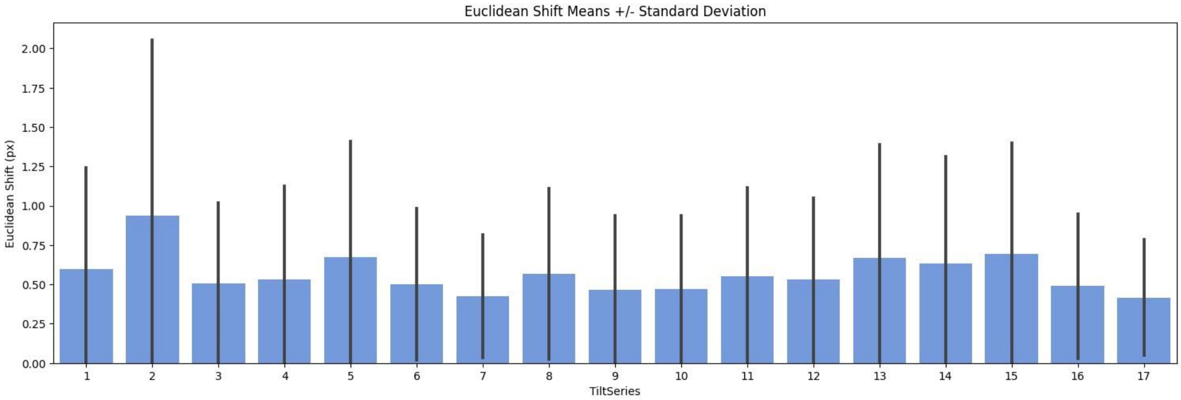


### IMOD (general)

IMOD has been used in this image processing operation.

Please cite:

1. James R. Kremer, David N. Mastronarde, J.Richard McIntosh.

Computer Visualization of Three-Dimensional Image Data Using IMOD, *Journal of Structural Biology*
**116(1)**:71-76 (1996)

2. Mastronarde DN, Held SR.

Automated tilt series alignment and tomographic reconstruction in IMOD, *Journal of Structural Biology* **197(2)**:102-113 (2017)

#### Tilt-series Alignment (IMOD)

##### Shifts between patches in A

Taken from .xf file

Out[13]:

| | Tilt series | Mean shift (A) | Shift s.d. (A) |
|---|---|---|---|
| **0** | 1 | 12.740374 | 15.802750 |
| **1** | 2 | 19.183049 | 11.365527 |
| **2** | 3 | 38.332763 | 13.413515 |
| **3** | 4 | 26.860351 | 10.567297 |
| **4** | 5 | 17.211399 | 4.665976 |
| **5** | 6 | 23.029534 | 6.364917 |
| **6** | 7 | 24.225329 | 10.100222 |
| **7** | 8 | 29.900118 | 11.788393 |
| **8** | 9 | 22.523913 | 7.419159 |
| **9** | 10 | 11.398053 | 4.347359 |
| **10** | 11 | 7.497318 | 4.166424 |
| **11** | 12 | 13.138833 | 9.176081 |
| **12** | 13 | 7.771078 | 4.478610 |
| **13** | 14 | 28.762903 | 10.508627 |
| **14** | 15 | 16.075103 | 7.398575 |
| **15** | 16 | 11.060385 | 4.149286 |
| **16** | 17 | 14.037355 | 4.974487 |

Out[14]: (0.0, 54.48671074487035)

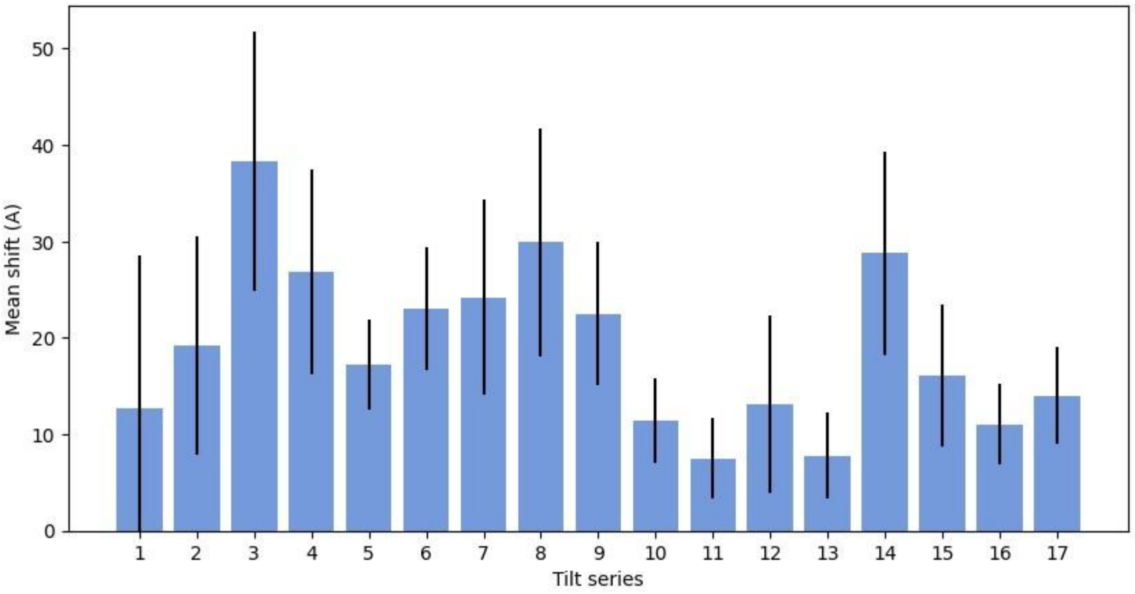


#### Errors in alignment

Taken from taLocals

| | Tilt series | Error mean (nm) | Error SD (nm) |
|---|---|---|---|
| **0** | 1 | 0.416 | 0.289 |
| **1** | 2 | 0.475 | 0.287 |
| **2** | 3 | 0.370 | 0.286 |
| **3** | 4 | 0.366 | 0.270 |
| **4** | 5 | 0.426 | 0.277 |
| **5** | 6 | 0.391 | 0.262 |
| **6** | 7 | 0.388 | 0.272 |
| **7** | 8 | 0.434 | 0.296 |
| **8** | 9 | 0.361 | 0.264 |
| **9** | 10 | 0.398 | 0.279 |
| **10** | 11 | 0.406 | 0.279 |
| **11** | 12 | 0.400 | 0.264 |

| | | | |
|---|---|---|---|
| **12** | 13 | 0.426 | 0.286 |
| **13** | 14 | 0.468 | 0.298 |
| **14** | 15 | 0.483 | 0.296 |
| **15** | 16 | 0.365 | 0.259 |
| **16** | 17 | 0.345 | 0.269 |

```
Out[16]: (0.0, 0.81795)
```

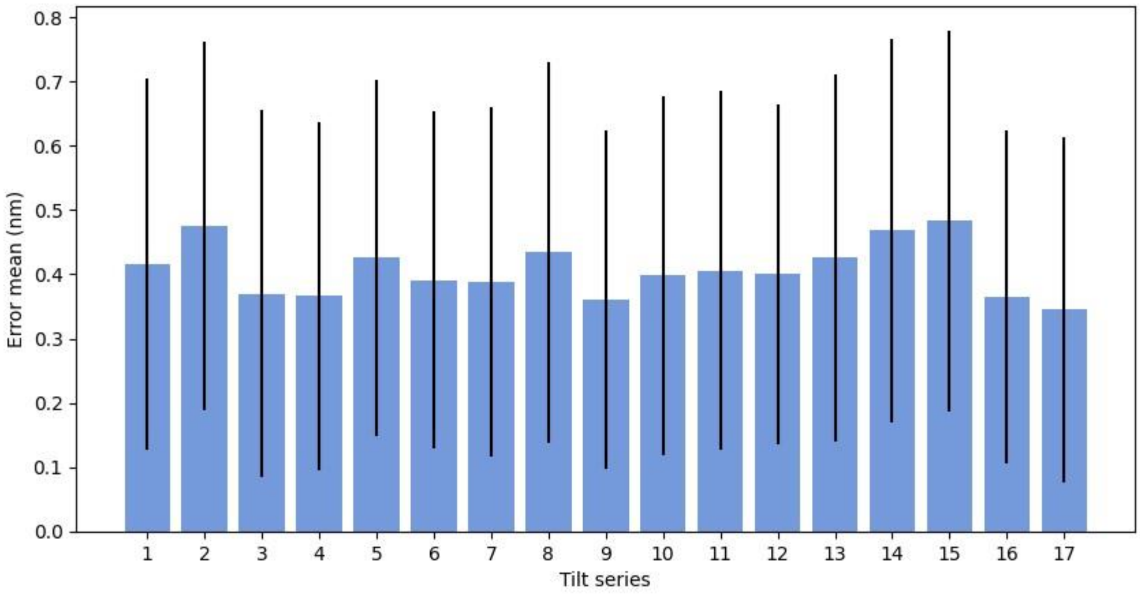


### Tomographic Reconstruction (IMOD)

Reconstruction was performed with IMOD BatchRunTomo.

Reconstruction algorithm: WBP

For more information, please see https://bio3d.colorado.edu/imod/doc/directives.html

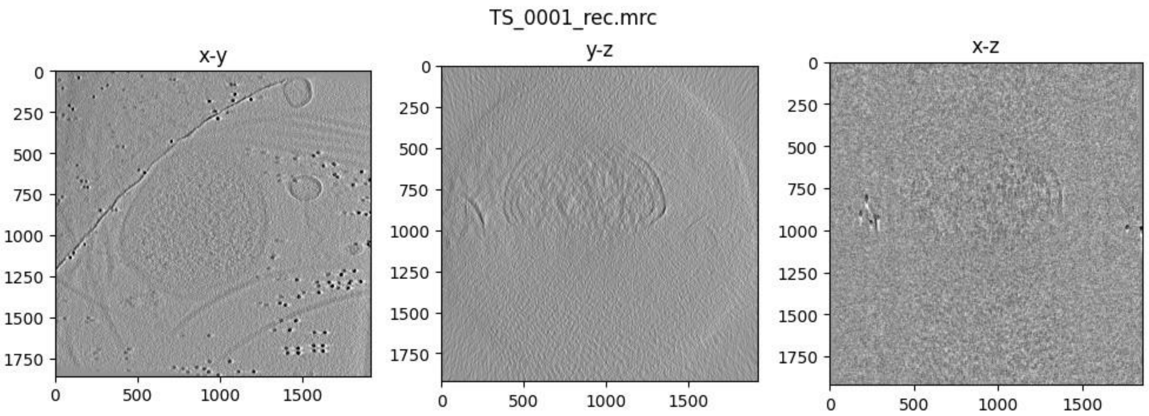
TS_0001_rec.mrc
x-y
y-z
x-z

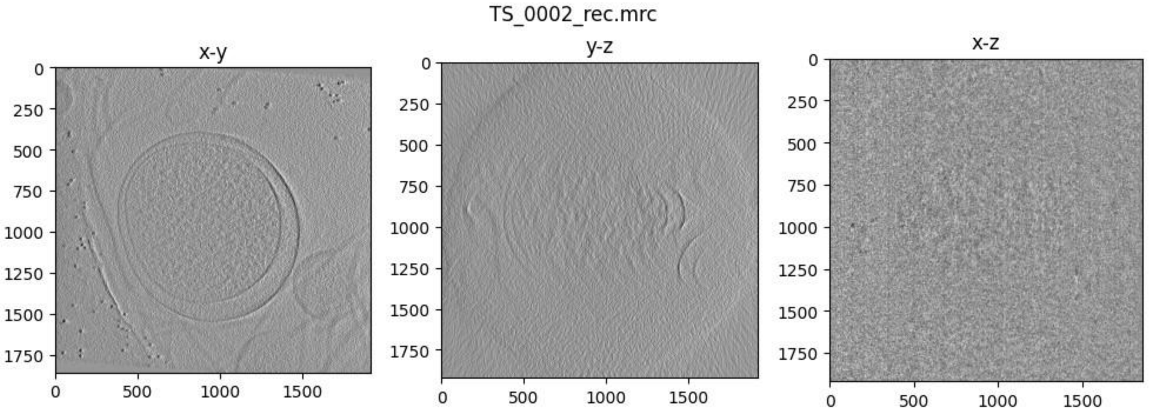
TS_0002_rec.mrc
x-y
y-z
x-z

TS_0003_rec.mrc

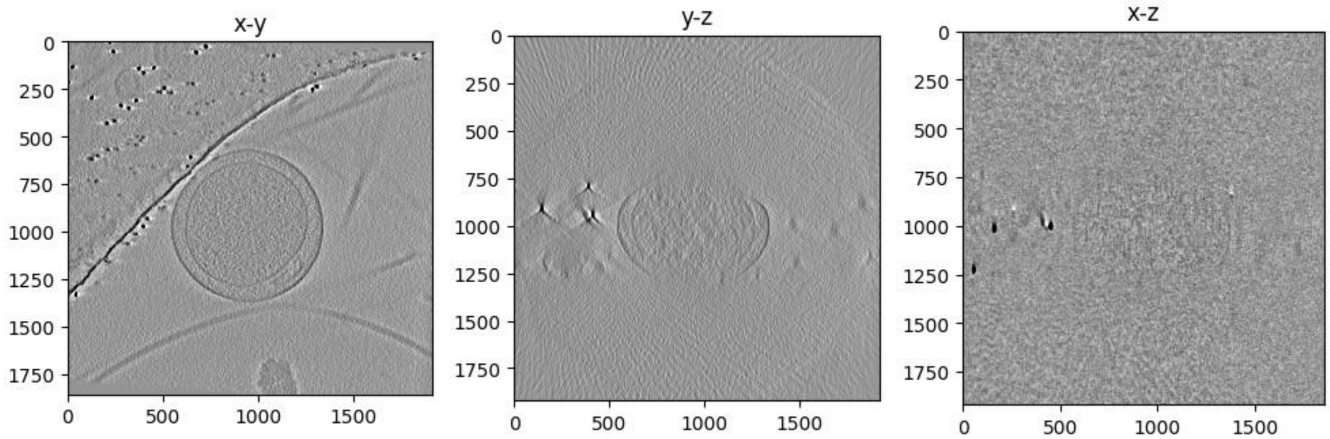


TS_0004_rec.mrc

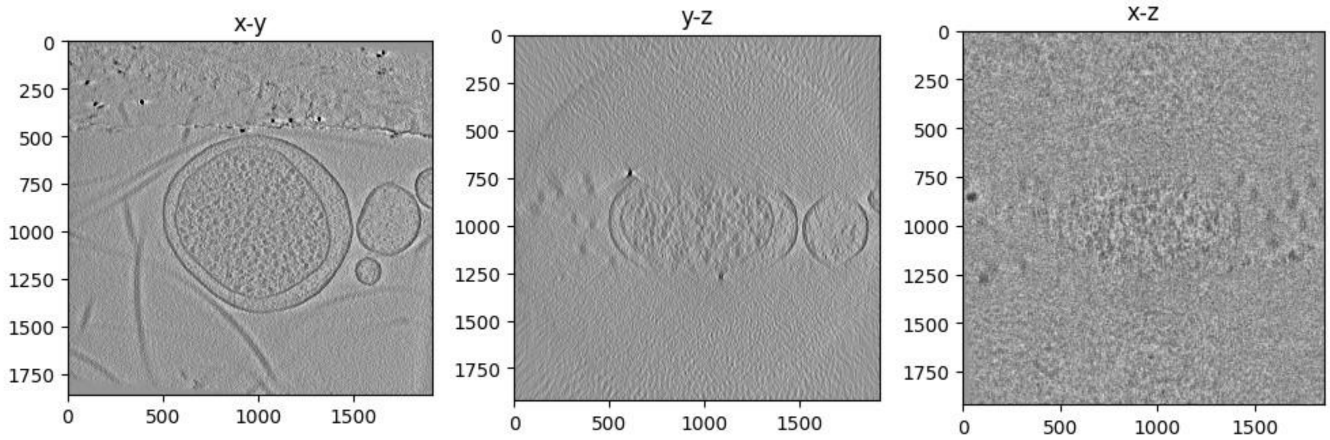


TS_0005_rec.mrc

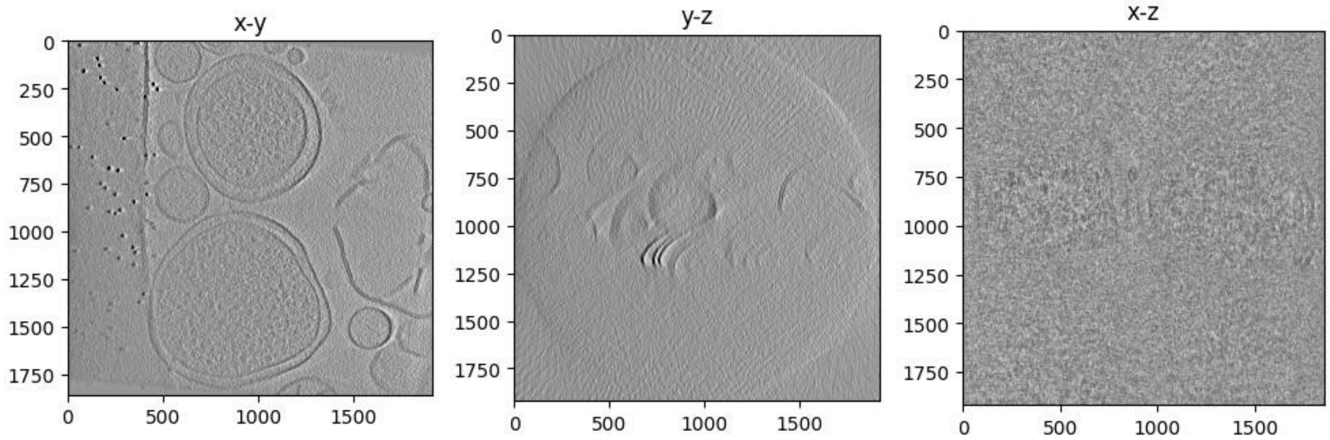


TS_0006_rec.mrc

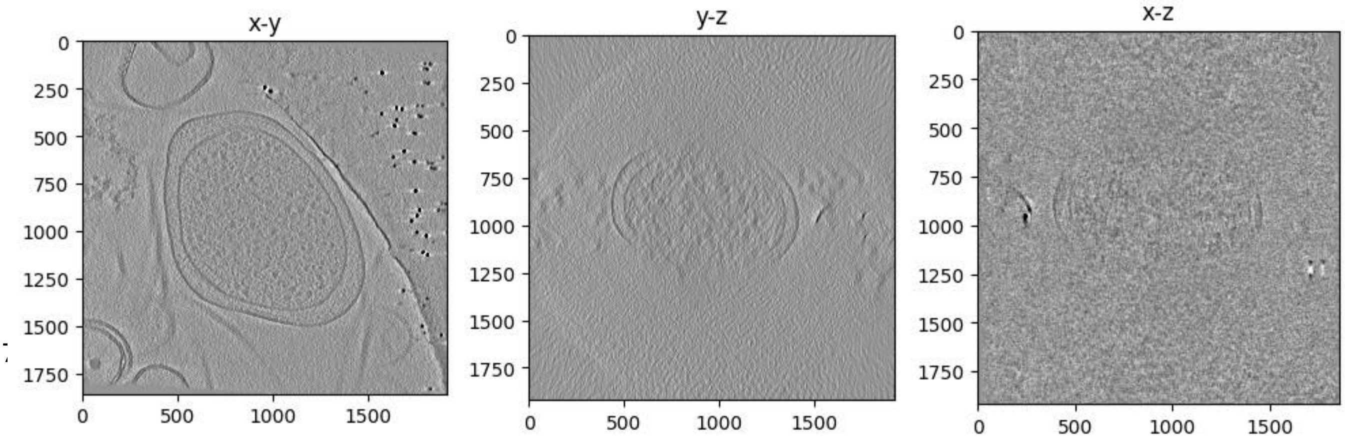

TS_0007_rec.mrc

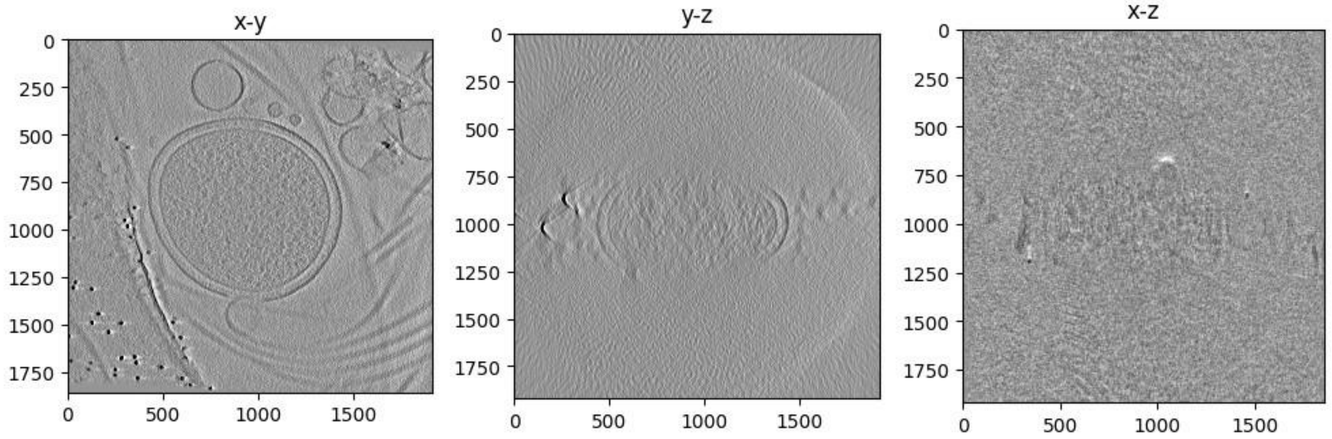


TS_0008_rec.mrc

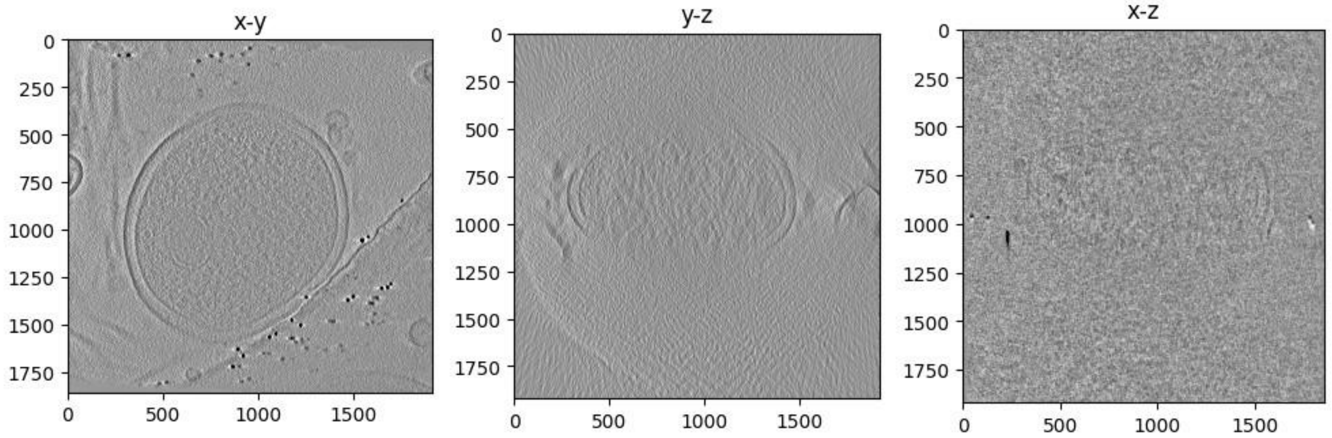


TS_0009_rec.mrc

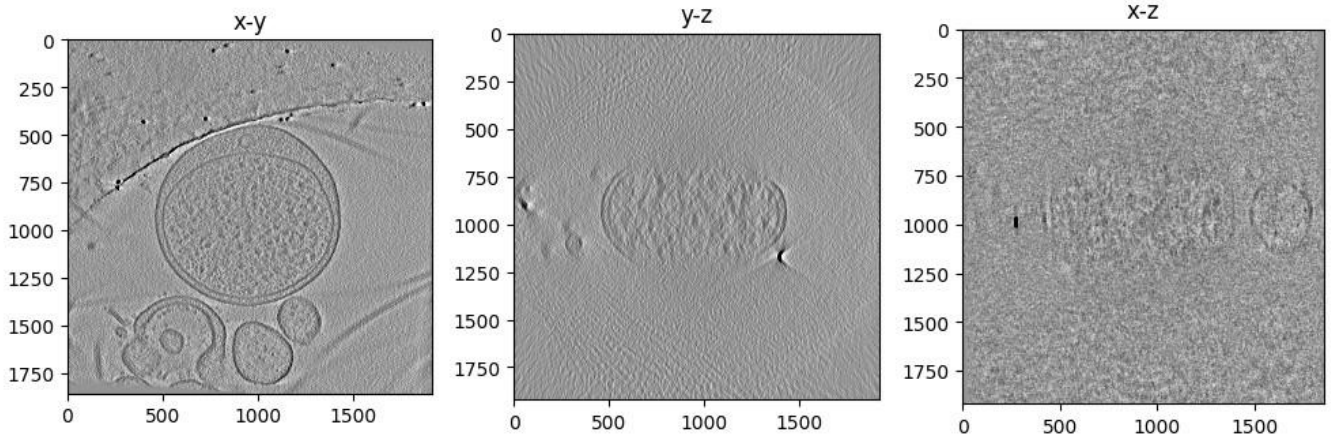


TS_0010_rec.mrc

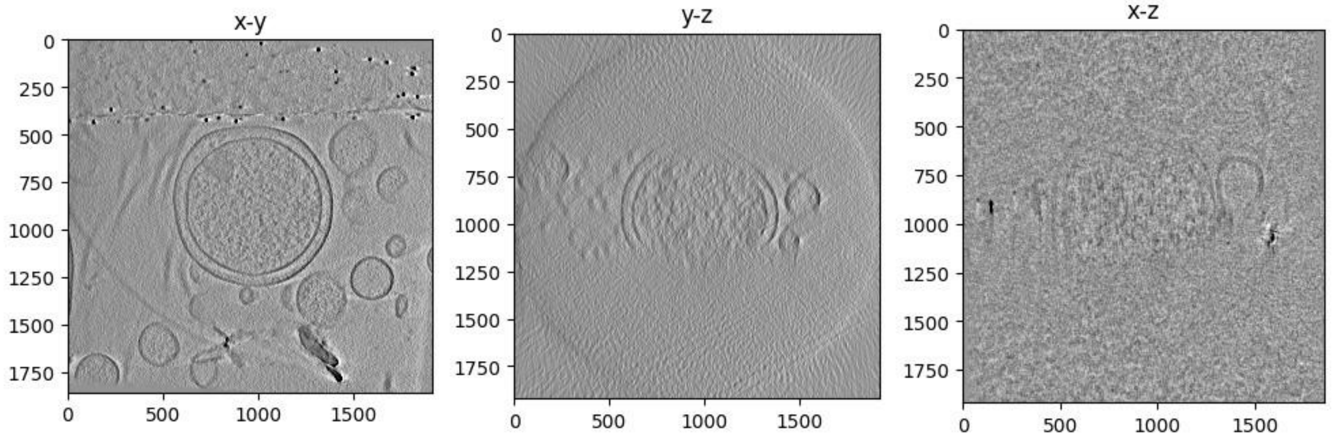

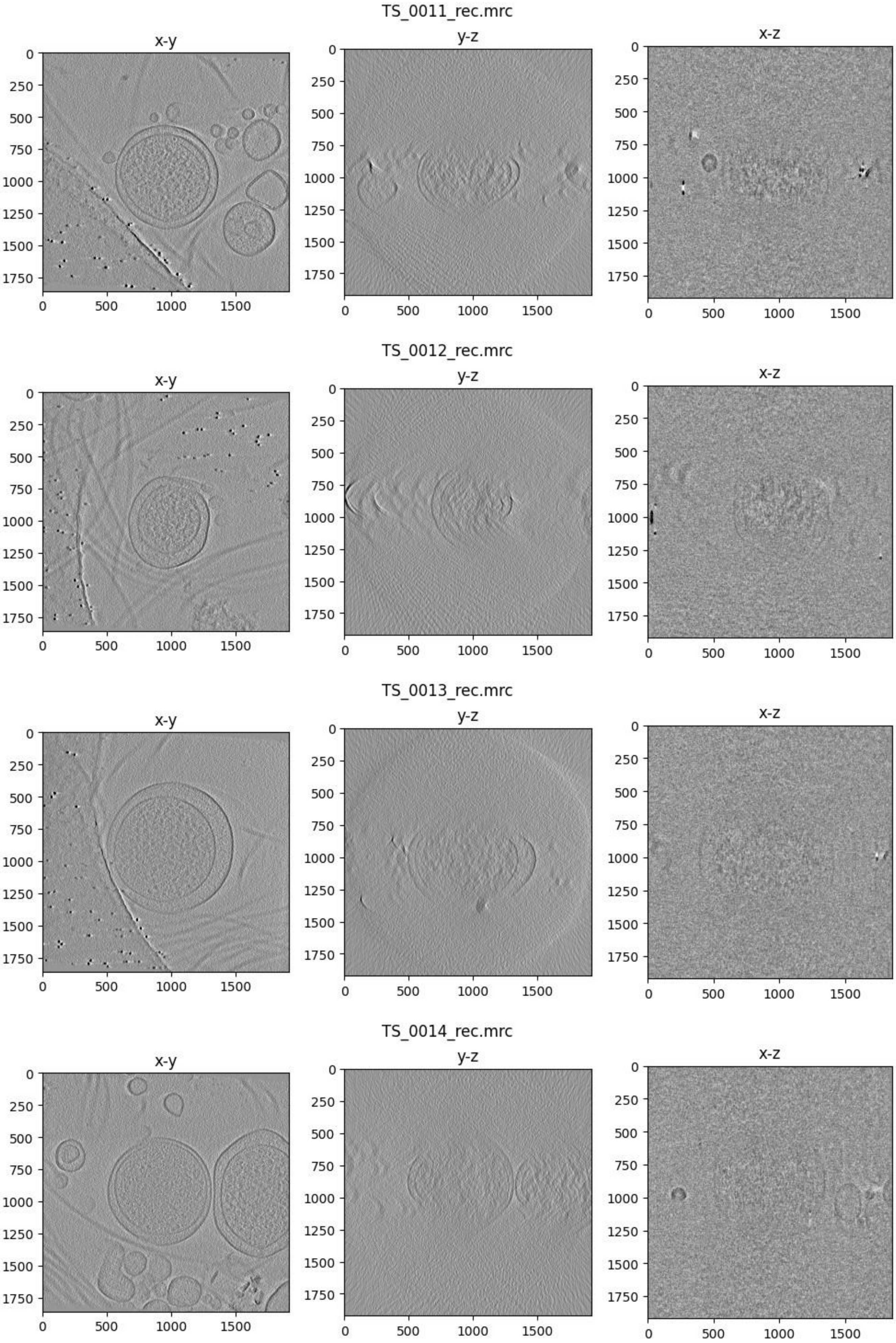
TS_0011_rec.mrc
x-y
y-z
x-z
TS_0012_rec.mrc
x-y
y-z
x-z
TS_0013_rec.mrc
x-y
y-z
x-z
TS_0014_rec.mrc
x-y
y-z
x-z

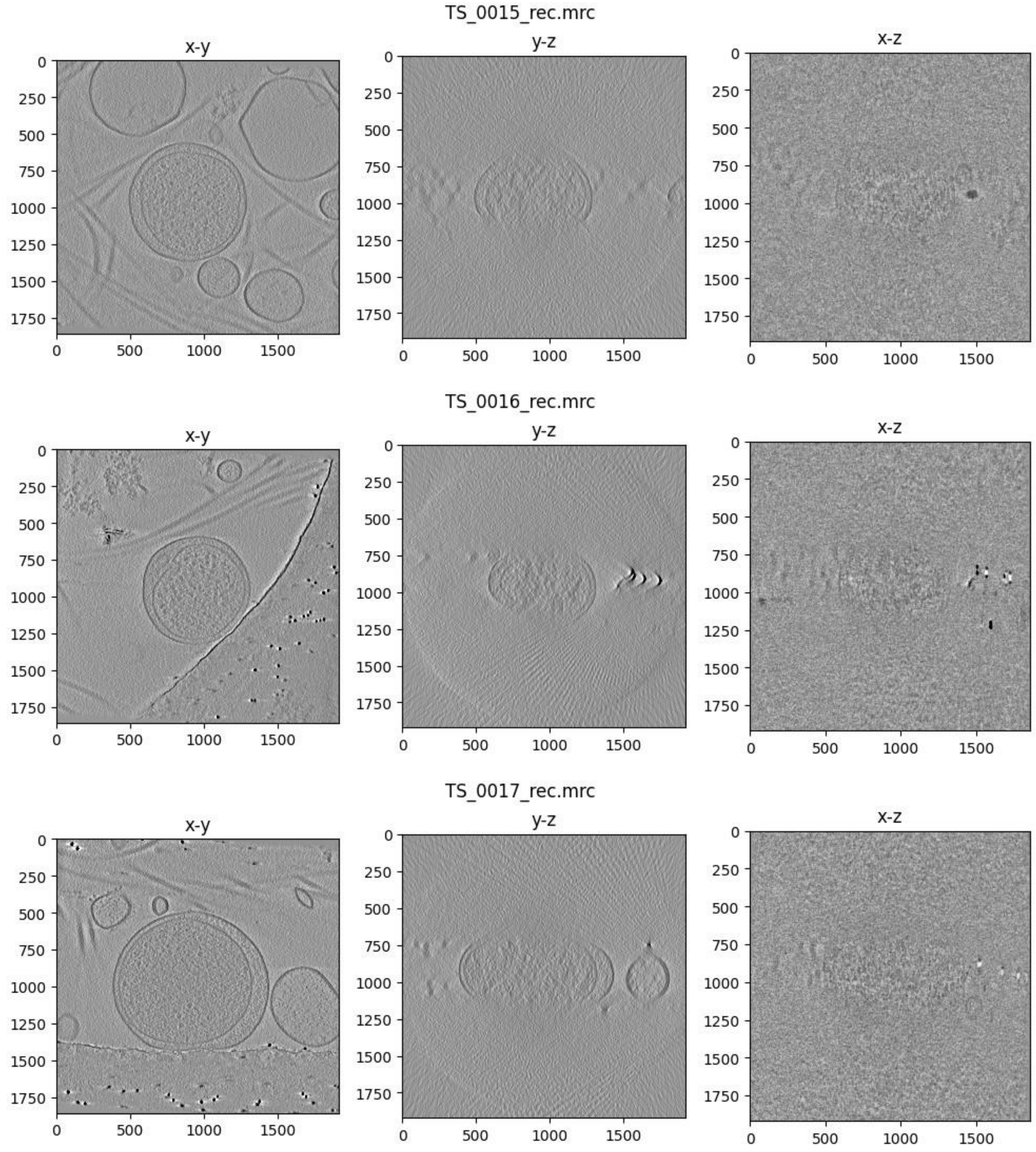
TS_0015_rec.mrc
x-y
y-z
x-z
TS_0016_rec.mrc
x-y
y-z
x-z
TS_0017_rec.mrc
x-y
y-z
x-z
0
250
500
750
1000
1250
1500
1750
0
500
1000
1500

### Workflow diagram

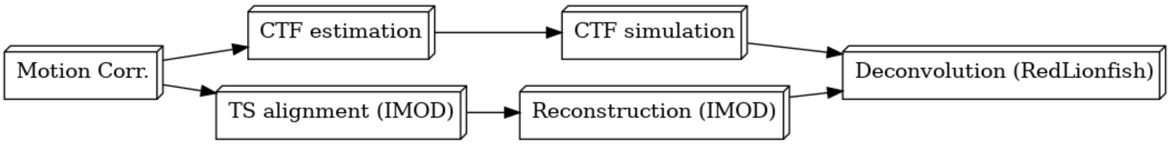


### Motion correction

Motion correction has been performed using MotionCor2.

Please cite:

1. Zheng, S., Palovcak, E., Armache, JP. et al.

MotionCor2: anisotropic correction of beam-induced motion for improved cryo-electron microscopy.
*Nat Methods* **14**, 331–332 (2017).

#### Summary of shifts for each tilt series

Out[13]:

| | count | mean | std | min | 25% | 50% | 75% | max |
|---|---|---|---|---|---|---|---|---|
| **TiltSeries** | | | | | | | | |
| **6** | 10250.0 | 0.532437 | 0.592415 | 0.0 | 0.186011 | 0.358469 | 0.629682 | 5.513765 |
| **7** | 10250.0 | 0.394030 | 0.288621 | 0.0 | 0.189737 | 0.351497 | 0.552268 | 2.871167 |
| **9** | 10250.0 | 0.631271 | 0.667577 | 0.0 | 0.226716 | 0.439318 | 0.770114 | 4.961371 |
| **10** | 10250.0 | 0.547538 | 0.402059 | 0.0 | 0.267207 | 0.489591 | 0.758172 | 3.580461 |
| **11** | 10250.0 | 0.289698 | 0.198691 | 0.0 | 0.148661 | 0.269165 | 0.406079 | 1.351481 |
| **12** | 10250.0 | 0.277824 | 0.186826 | 0.0 | 0.141421 | 0.261725 | 0.394462 | 1.213960 |
| **13** | 10250.0 | 0.449222 | 0.545662 | 0.0 | 0.187883 | 0.350143 | 0.566127 | 20.940306 |
| **14** | 10250.0 | 0.344168 | 0.241599 | 0.0 | 0.170880 | 0.314006 | 0.482701 | 1.739224 |
| **15** | 10250.0 | 0.299349 | 0.234410 | 0.0 | 0.141421 | 0.260192 | 0.410000 | 2.072342 |

```
Out[14]: Text(0.5, 1.0, 'Euclidean Shift Means +/- Standard
Deviation')
```

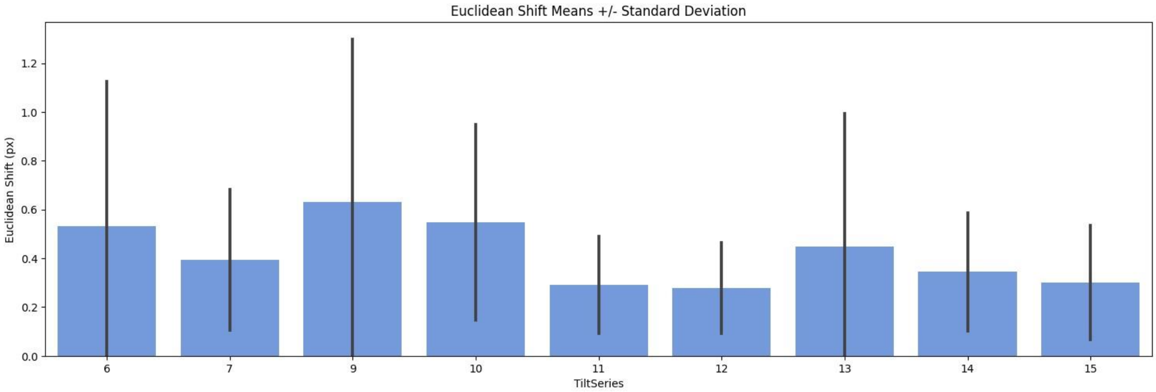


#### Summary of shifts for experiment

Euclidean shifts for all tilt series in the experiment.

```
count    92250.000000
mean         0.418393
std          0.429397
min          0.000000
25%          0.174642
50%          0.330151
75%          0.533104
max         20.940306
Name: EuclideanShift, dtype: float64
```

### CTF (Defocus) estimation

```
CTF estimation has been performed

using CTFFind4. Please cite:
```


1. Joseph A. Mindell, Nikolaus Grigorieff. Accurate determination of local defocus and specimen tilt in electron microscopy, *Journal of Structural Biology* **142(3)**:334-347 (2003)
2. Alexis Rohou, Nikolaus Grigorieff. CTFFIND4: Fast and accurate defocus estimation from electron micrographs, *Journal of Structural Biology* **192(2)**:216-221 (2015)


### CTF simulation

```
CTF simulation has been performed using

O2R-ctfsim. Please cite:
```

1. Alexis Rohou, Nikolaus Grigorieff. CTFFIND4: Fast and accurate defocus estimation from electron micrographs, *Journal of Structural Biology* **192(2)**:216-221 (2015)

### IMOD (general)

```
IMOD has been used in this image processing

operation. Please cite:
```

1. James R. Kremer, David N. Mastronarde, J.Richard McIntosh. Computer Visualization of ThreeDimensional Image Data Using IMOD, *Journal of Structural Biology* **116(1)**:71-76 (1996)
2. Mastronarde DN, Held SR. Automated tilt series alignment and tomographic reconstruction in IMOD, *Journal of Structural Biology* **197(2)**:102-113 (2017)

#### Tilt-series Alignment (IMOD)

| | Tilt series | Error mean (nm) | Error SD (nm) | Error weighted mean (nm) |
|---|---|---|---|---|
| 0 | 6 | 0.142 | 0.098 | 0.137 |
| 1 | 7 | 0.654 | 0.459 | 0.641 |
| 2 | 9 | 0.456 | 0.380 | 0.446 |
| 3 | 10 | 1.306 | 0.886 | 1.279 |
| 4 | 11 | 0.314 | 0.214 | 0.303 |
| 5 | 12 | 0.458 | 0.327 | 0.445 |
| 6 | 13 | 0.403 | 0.349 | 0.392 |
| 7 | 14 | 0.626 | 0.483 | 0.613 |
| 8 | 15 | 0.605 | 0.466 | 0.583 |

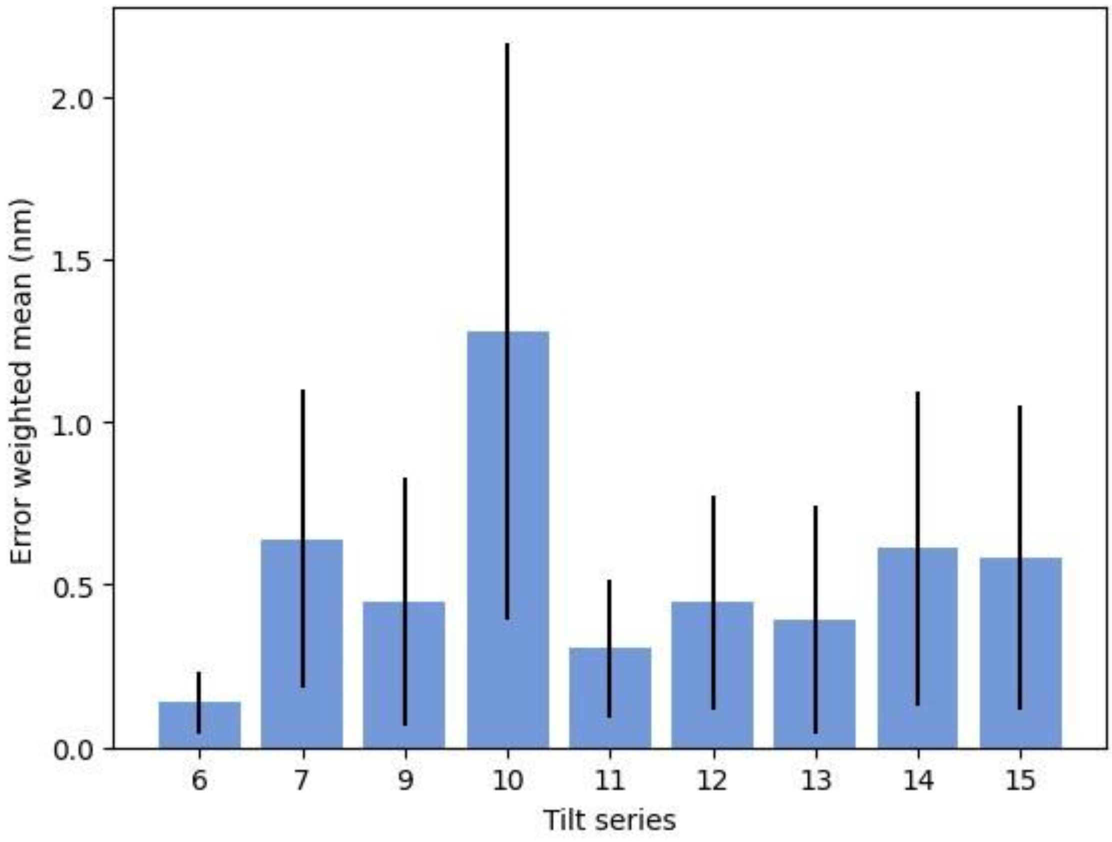


### Tomographic Reconstruction (IMOD)

Reconstruction was performed with IMOD BatchRunTomo.

Reconstruction algorithm: WBP
For more information, please see https://bio3d.colorado.edu/imod/doc/directives.html

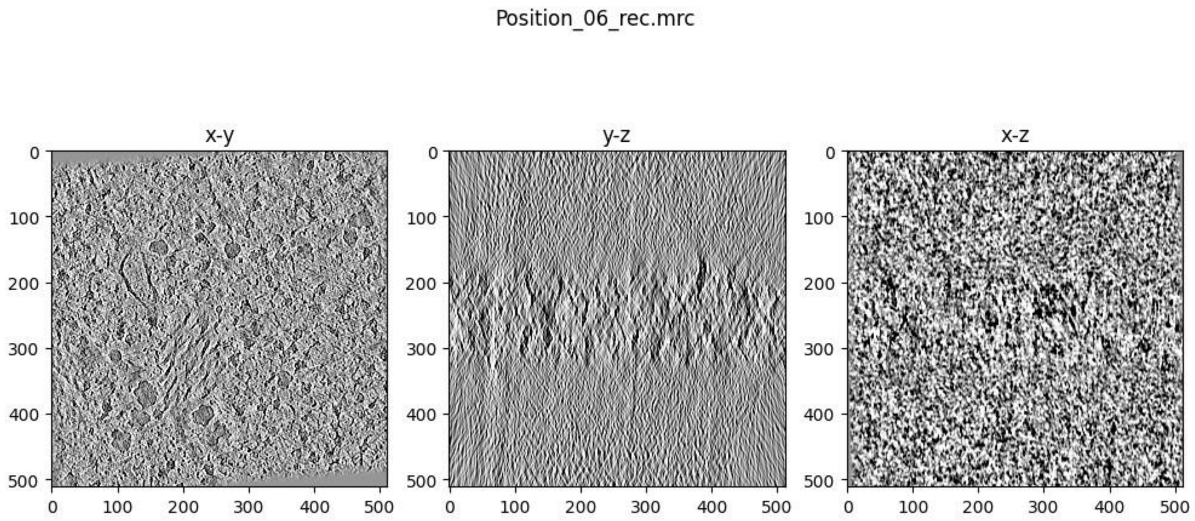
Position_06_rec.mrc
x-y
y-z
x-z
0
100
200
300
400
500

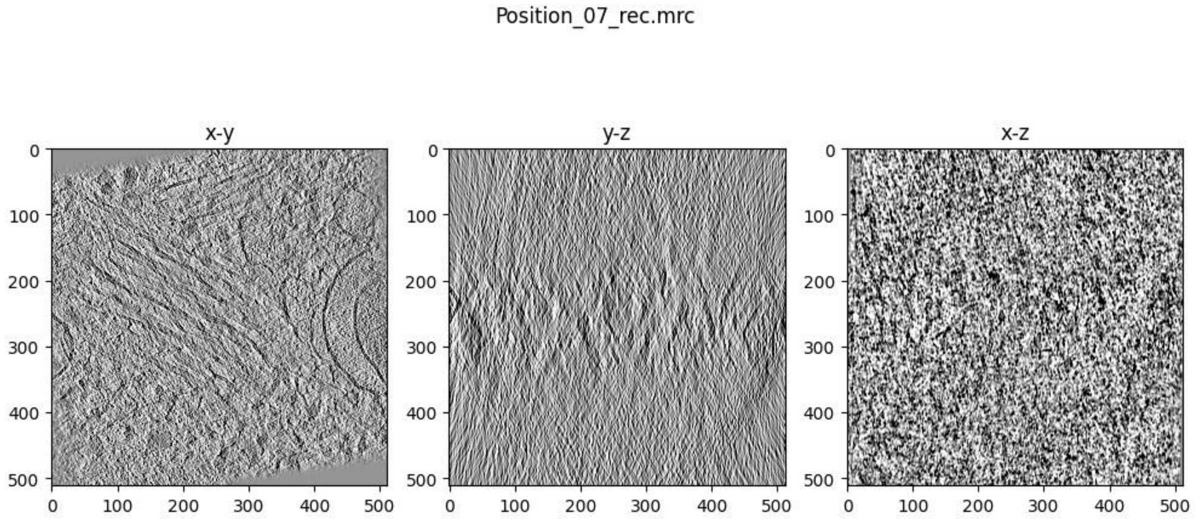
Position_07_rec.mrc
x-y
y-z
x-z
0
100
200
300
400
500

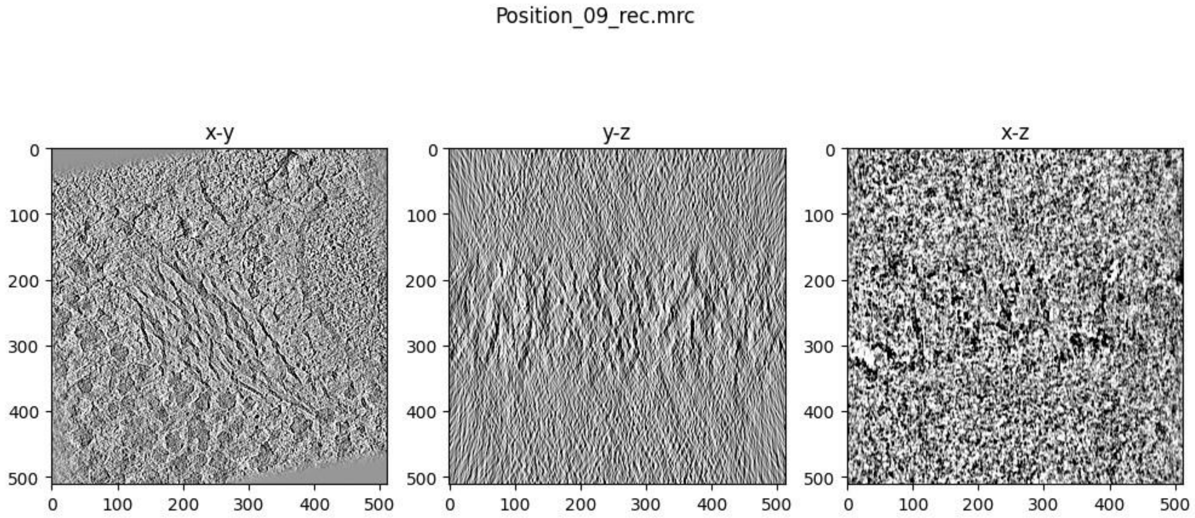
Position_09_rec.mrc
x-y
y-z
x-z
0
100
200
300
400
500

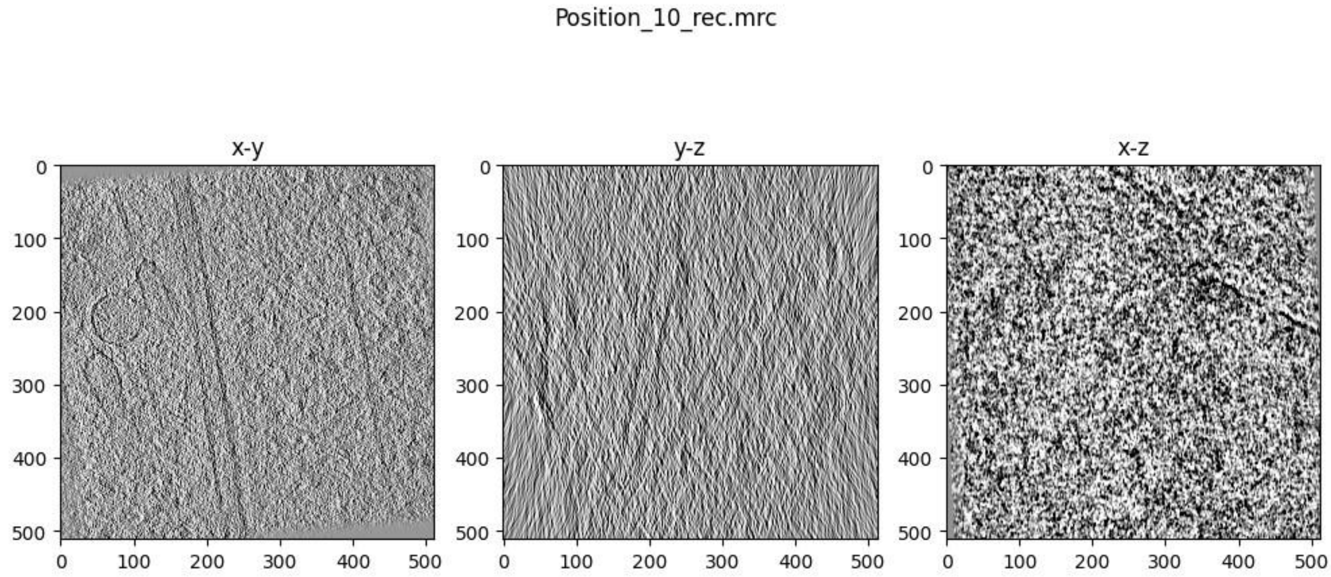
Position_10_rec.mrc
x-y
y-z
x-z

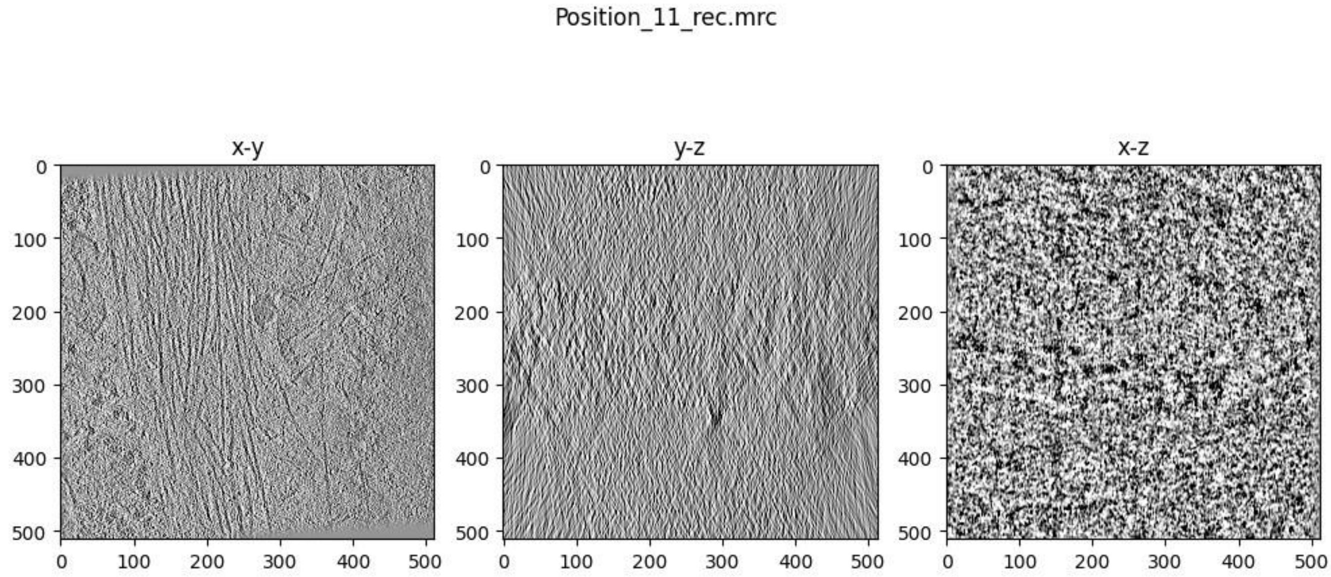
Position_11_rec.mrc
x-y
y-z
x-z

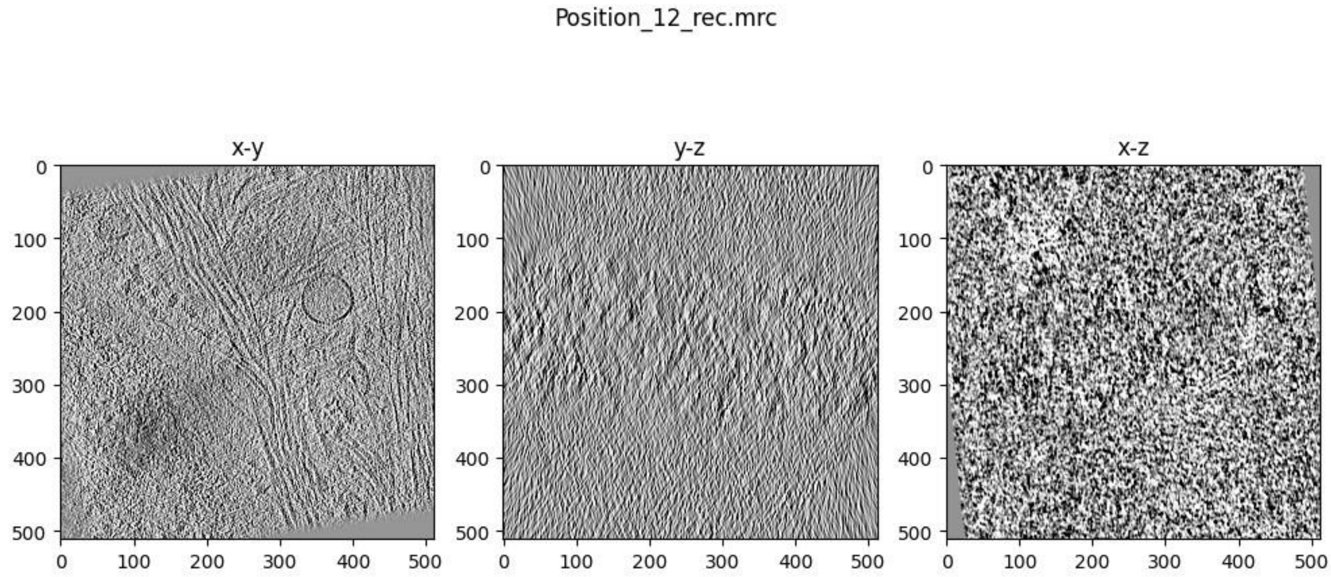
Position_12_rec.mrc
x-y
y-z
x-z

Position_13_rec.mrc

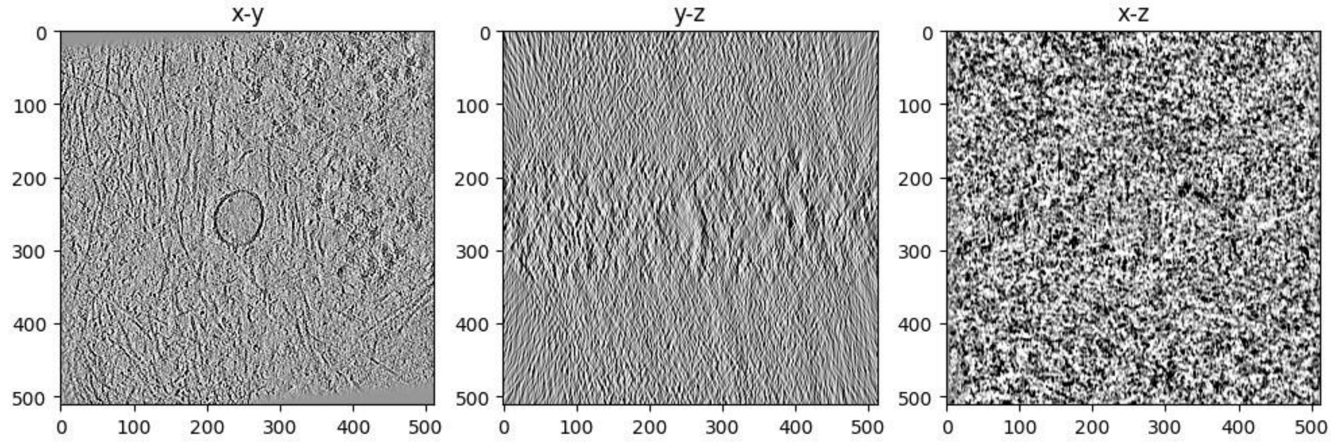


Position_14_rec.mrc

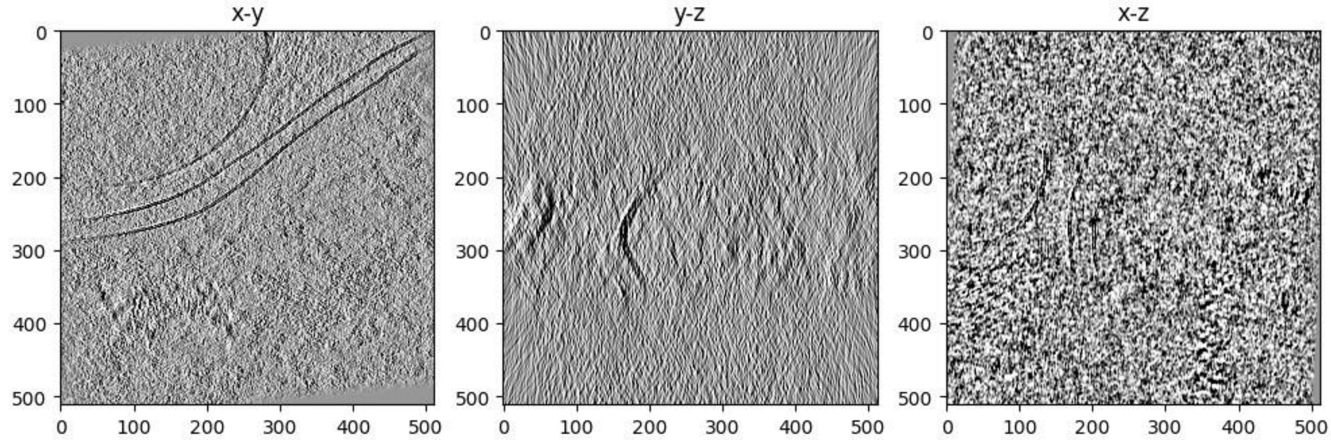


Position_15_rec.mrc

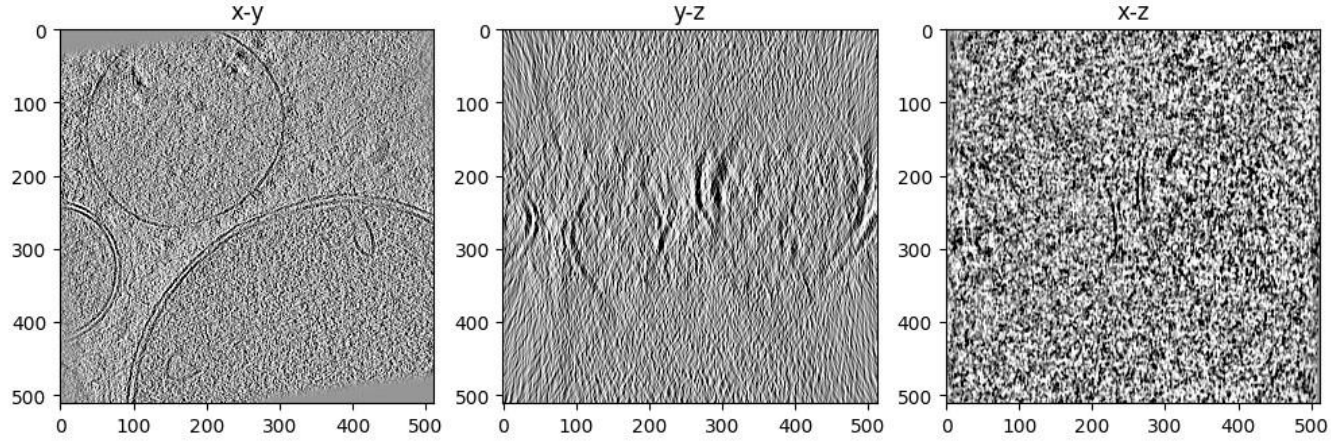

### Richardson-Lucy Deconvolution with RedLionFish

For more information, please see https://github.com/rosalindfranklininstitute/RedLionfish

#### Deconvolved thumbnails

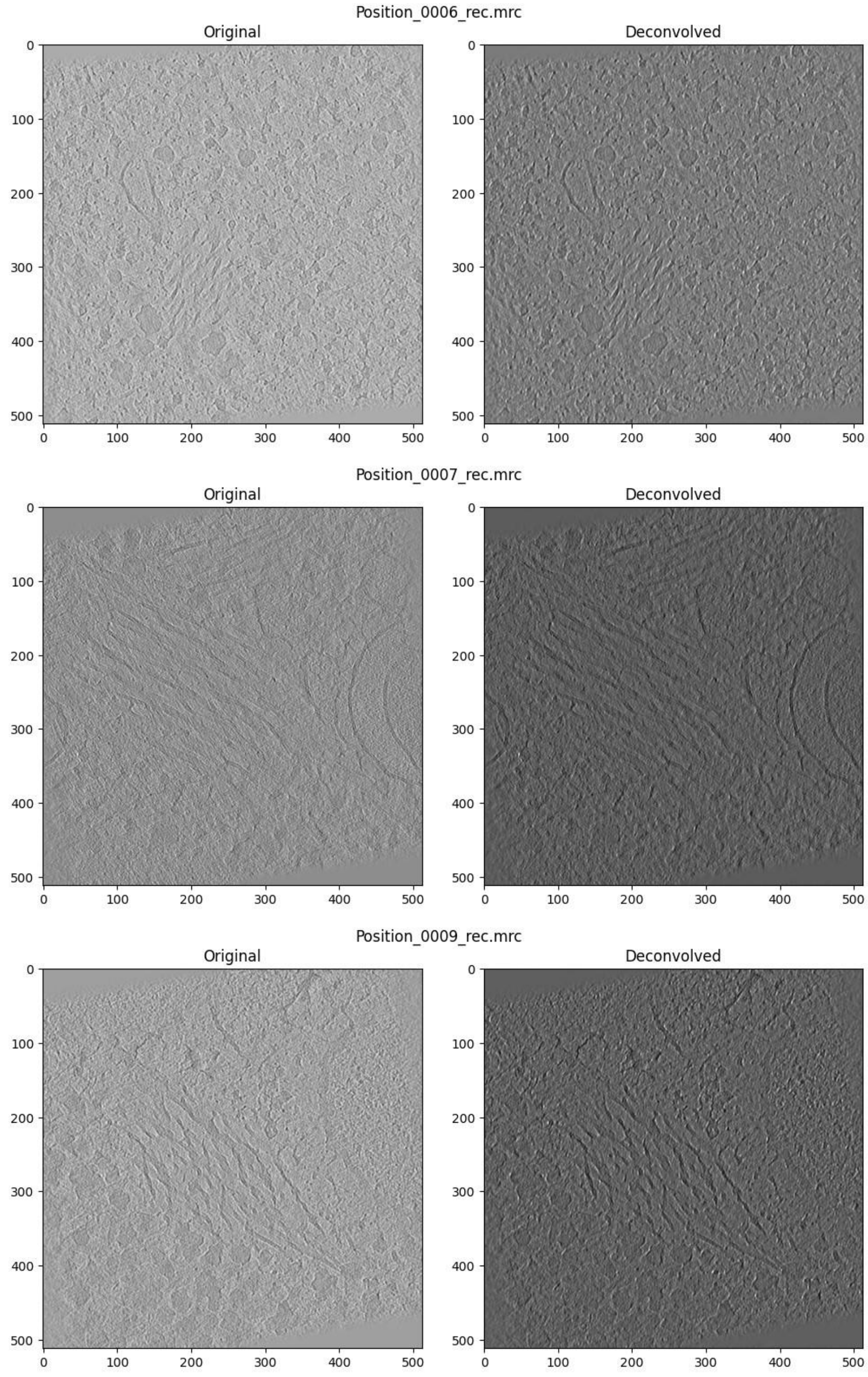

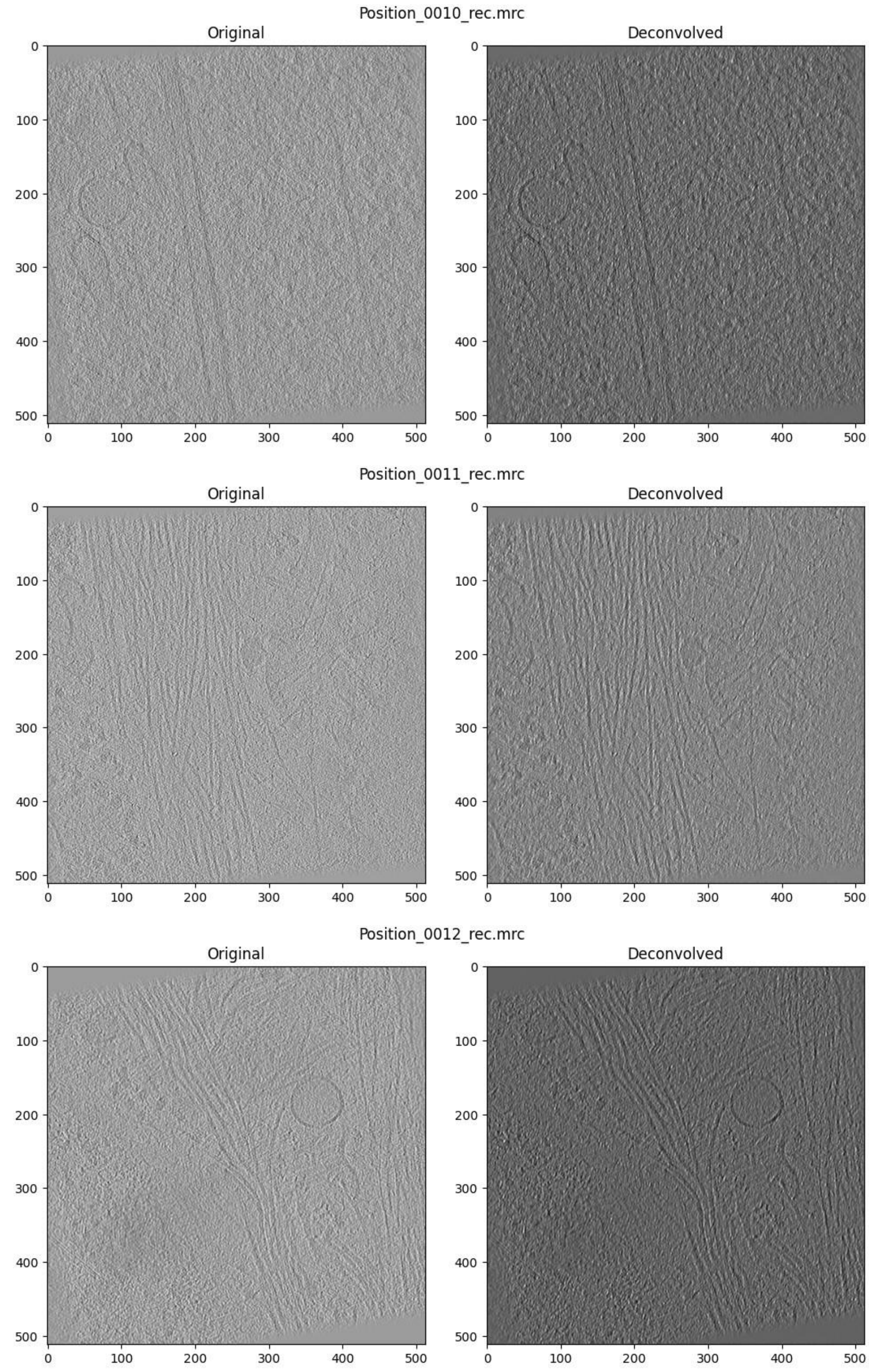
Position_0010_rec.mrc
Original
Deconvolved
0 100 200 300 400 500
Position_0011_rec.mrc
Original
Deconvolved
0 100 200 300 400 500
Position_0012_rec.mrc
Original
Deconvolved
0 100 200 300 400 500

Position_0013_rec.mrc

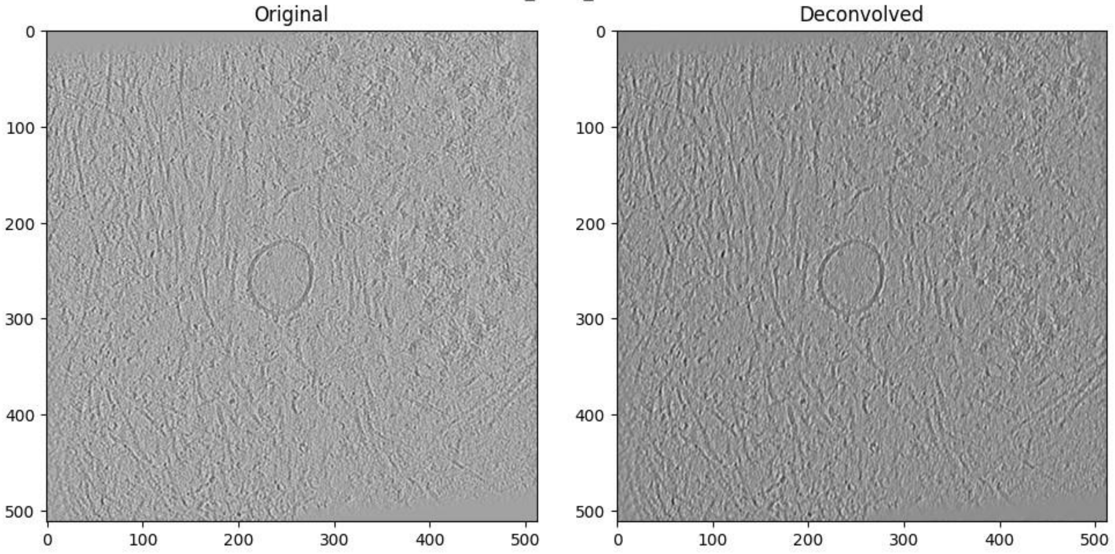


Position_0014_rec.mrc

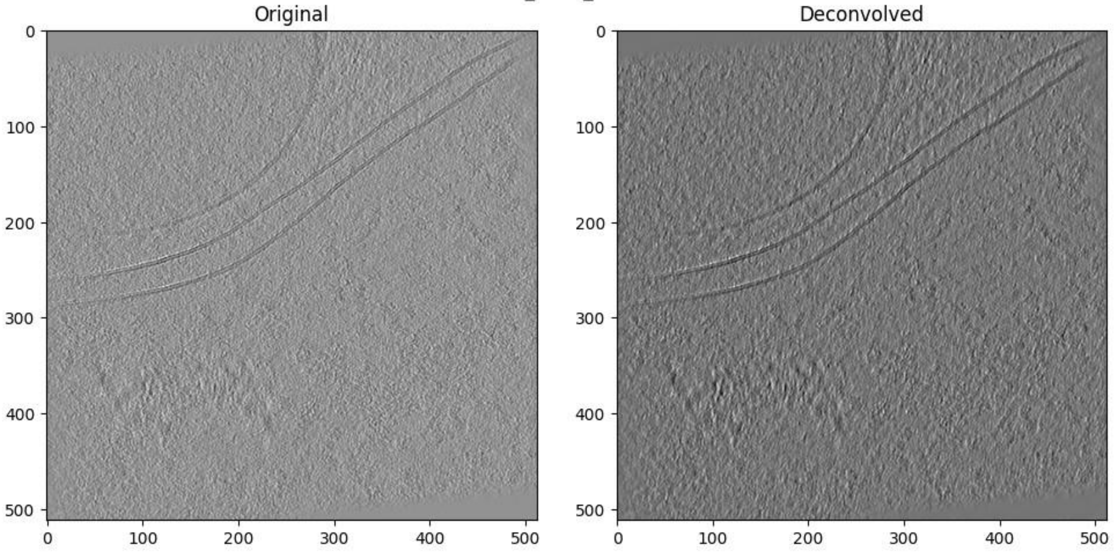


Position_0015_rec.mrc

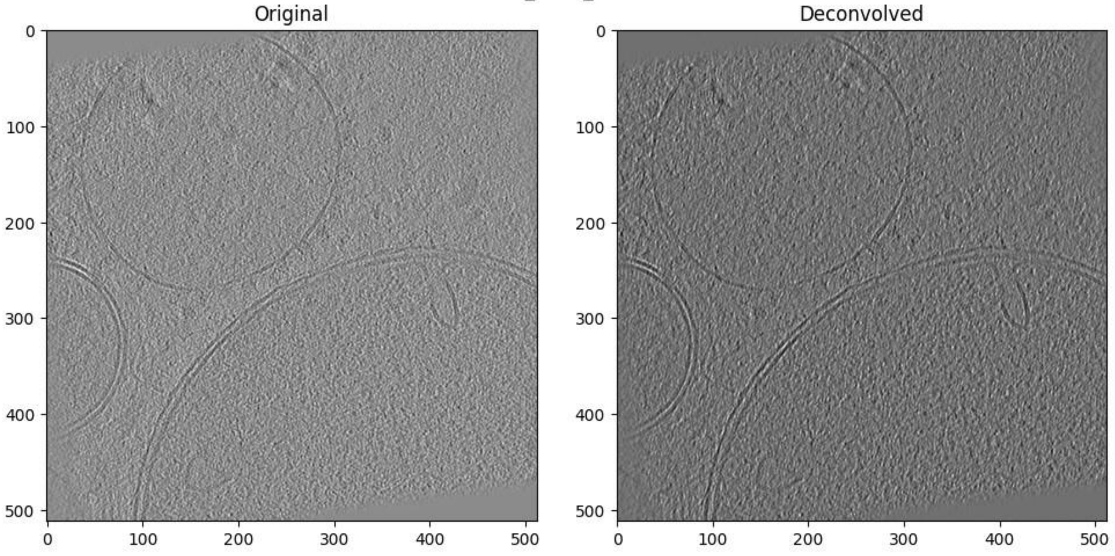

### Workflow diagram

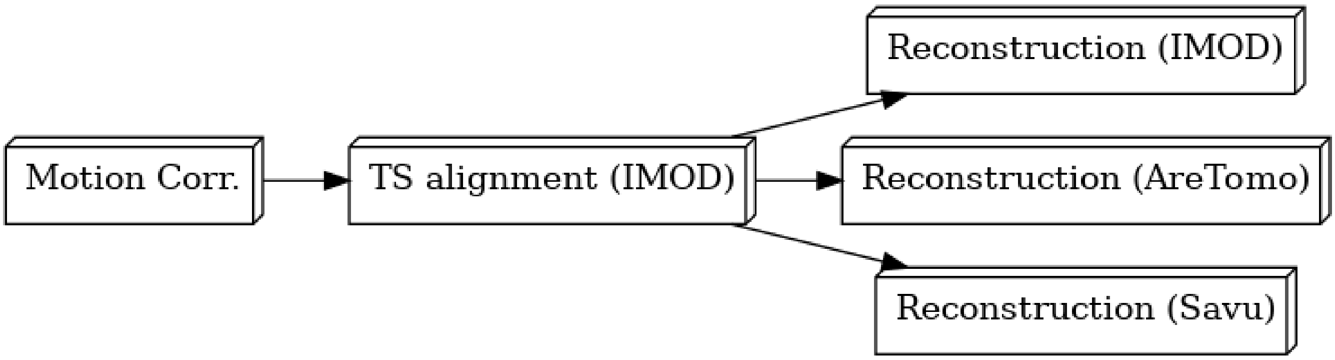


### Motion correction

Motion correction has been performed using MotionCor2.

Please cite:

1. Zheng, S., Palovcak, E., Armache, JP. et al. MotionCor2: anisotropic correction of beam-induced motion for improved cryo-electron microscopy. *Nat Methods* **14**, 331–332 (2017).

#### Summary of shifts for each tilt series

| TiltSeries | mean | std | 25% | 50% | 75% |
|---|---|---|---|---|---|
| **1** | 2.056202 | 2.656386 | 0.635059 | 1.234585 | 2.386598 |
| **2** | 1.415733 | 1.614418 | 0.521536 | 1.001249 | 1.791368 |
| **3** | 1.295799 | 1.117591 | 0.512445 | 0.965660 | 1.759602 |
| **4** | 1.661103 | 2.091689 | 0.577062 | 1.106029 | 1.981691 |
| **5** | 1.298596 | 1.033565 | 0.554437 | 1.041009 | 1.762385 |
| **6** | 1.379971 | 1.221563 | 0.605310 | 1.103087 | 1.829891 |
| **7** | 2.242506 | 2.905982 | 0.682642 | 1.346291 | 2.616104 |
| **8** | 1.402211 | 1.245235 | 0.562228 | 1.070000 | 1.881303 |

| | | | | | |
|---|---|---|---|---|---|
| **10** | 1.209009 | 0.884546 | 0.586941 | 1.011929 | 1.598405 |
| **11** | 1.871856 | 2.428340 | 0.586941 | 1.202477 | 2.268127 |
| **12** | 1.623454 | 2.318478 | 0.503289 | 0.973293 | 1.905197 |
| **13** | 6.831695 | 6.379735 | 2.314584 | 5.125159 | 9.487324 |
| **14** | 16.070258 | 49.565171 | 0.590339 | 1.156849 | 2.527969 |
| **15** | 19.491723 | 55.706889 | 0.605392 | 1.180042 | 2.618416 |
| **16** | 24.059852 | 65.467685 | 0.665733 | 1.379021 | 3.073308 |
| **17** | 20.347179 | 59.296513 | 0.617414 | 1.267793 | 2.946994 |
| **18** | 19.982279 | 56.926885 | 0.621772 | 1.191512 | 2.670431 |
| **19** | 23.301804 | 59.528379 | 1.168428 | 2.810970 | 9.441590 |
| **20** | 22.763273 | 73.947772 | 0.609590 | 1.279453 | 3.041772 |
| **21** | 15.023953 | 48.436937 | 0.653911 | 1.341827 | 2.771299 |
| **22** | 12.322676 | 45.376714 | 0.684251 | 1.385135 | 2.668122 |
| **23** | 10.351220 | 35.034275 | 0.687677 | 1.371860 | 2.622532 |
| **24** | 10.273665 | 39.577862 | 0.594643 | 1.135319 | 2.231569 |

```
Out[12]: Text(0.5, 1.0, 'Euclidean Shift Means +/- Standard Deviation')
```

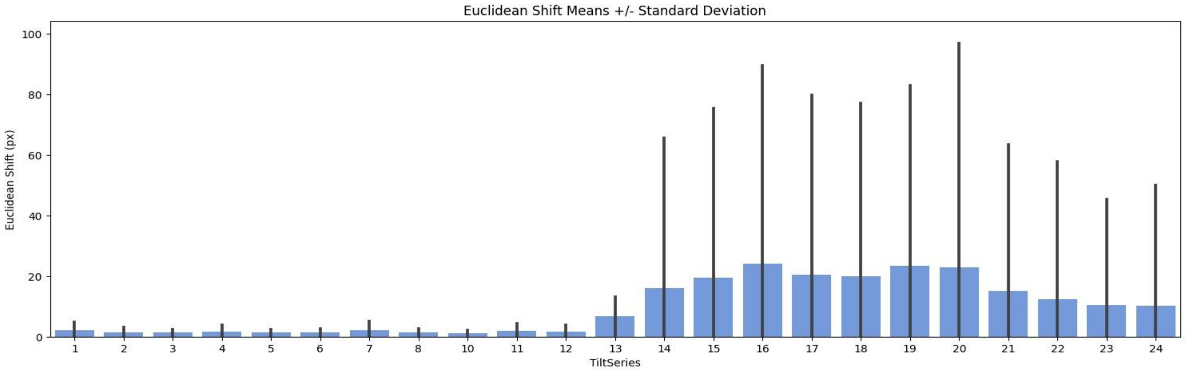

### IMOD (general)

IMOD has been used in this image processing operation.

Please cite:

1. James R. Kremer, David N. Mastronarde, J.Richard McIntosh.

Computer Visualization of Three-Dimensional Image Data Using IMOD, *Journal of Structural Biology* **116(1)**:71-76 (1996)

2. Mastronarde DN, Held SR.

Automated tilt series alignment and tomographic reconstruction in IMOD, *Journal of Structural Biology* **197(2)**:102-113 (2017)

### Tilt-series Alignment (IMOD)

#### Shifts between patches in A

Taken from .xf file

| | Tilt series | Mean shift (A) | Shift s.d. (A) |
|---|---|---|---|
| **0** | 1 | 166.236990 | 80.176748 |
| **1** | 2 | 177.115113 | 101.861063 |
| **2** | 3 | 160.303164 | 82.103914 |
| **3** | 4 | 256.280416 | 171.837799 |
| **4** | 5 | 168.619277 | 103.502143 |
| **5** | 6 | 185.401710 | 132.234260 |
| **6** | 7 | 304.877732 | 177.761258 |
| **7** | 8 | 155.059100 | 113.871254 |
| **8** | 10 | 176.344347 | 139.545098 |
| **9** | 11 | 178.296870 | 92.727205 |
| **10** | 12 | 158.912904 | 114.478130 |
| **11** | 13 | 1240.891903 | 854.958521 |
| **12** | 14 | 454.783686 | 624.699055 |
| **13** | 15 | 546.541572 | 489.618832 |
| **14** | 16 | 345.828423 | 238.146679 |
| **15** | 17 | 491.043815 | 606.243412 |
| **16** | 18 | 366.138999 | 367.136905 |
| **17** | 19 | 510.903068 | 596.798515 |
| **18** | 20 | 409.616139 | 512.951914 |
| **19** | 21 | 348.930781 | 480.343859 |
| **20** | 22 | 410.733088 | 421.374270 |
| **21** | 23 | 426.684826 | 352.738900 |
| **22** | 24 | 508.941401 | 229.217457 |

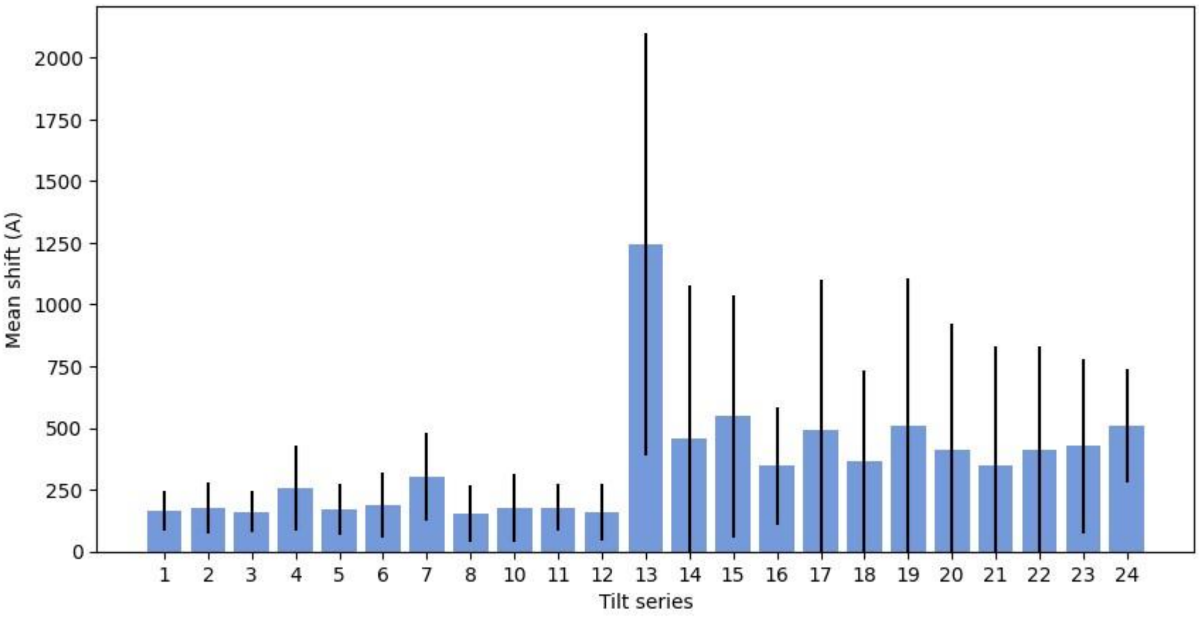
2000
1750
1500
1250
1000
750
500
250
0
Mean shift (A)
1 2 3 4 5 6 7 8 10 11 12 13 14 15 16 17 18 19 20 21 22 23 24
Tilt series

#### Errors in alignment

Taken from taLocals

| | Tilt series | Error mean (nm) | Error SD (nm) | Error weighted mean (nm) |
|---|---|---|---|---|
| **0** | 1 | 0.771 | 0.580 | 0.736 |
| **1** | 2 | 0.758 | 0.636 | 0.704 |
| **2** | 3 | 1.037 | 0.784 | 0.990 |
| **3** | 4 | 1.392 | 0.934 | 1.314 |
| **4** | 5 | 1.080 | 0.778 | 1.052 |
| **5** | 6 | 1.121 | 0.816 | 1.084 |
| **6** | 7 | 1.036 | 0.646 | 1.002 |
| **7** | 8 | 1.171 | 0.769 | 1.148 |
| **8** | 10 | 0.817 | 0.716 | 0.766 |
| **9** | 11 | 0.719 | 0.635 | 0.669 |
| **10** | 12 | 0.859 | 0.702 | 0.822 |
| **11** | 13 | 1.061 | 0.666 | 1.038 |
| **12** | 14 | 0.981 | 0.634 | 0.942 |
| **13** | 15 | 1.093 | 0.691 | 1.056 |
| **14** | 16 | 1.055 | 0.700 | 1.021 |
| **15** | 17 | 0.909 | 0.601 | 0.888 |
| **16** | 18 | 1.043 | 0.715 | 1.006 |
| **17** | 19 | 1.345 | 0.779 | 1.313 |
| **18** | 20 | 0.988 | 0.643 | 0.965 |
| **19** | 21 | 1.100 | 0.739 | 1.047 |
| **20** | 22 | 0.930 | 0.641 | 0.907 |
| **21** | 23 | 1.028 | 0.679 | 0.990 |
| **22** | 24 | 1.048 | 0.714 | 0.986 |

```
Out[16]: (0.0, 2.4423)
```

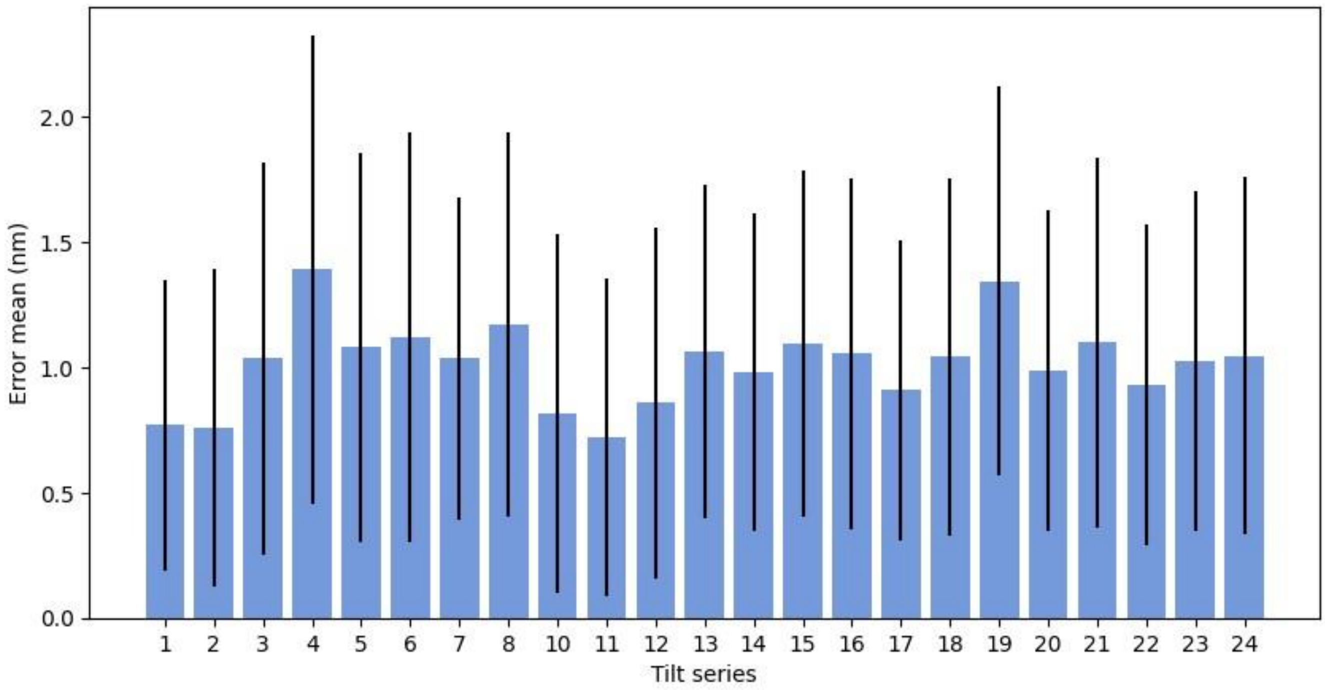

### Tomographic Reconstruction (IMOD)

Reconstruction was performed with IMOD BatchRunTomo.

Reconstruction algorithm: SIRT

For more information, please see https://bio3d.colorado.edu/imod/doc/directives.html

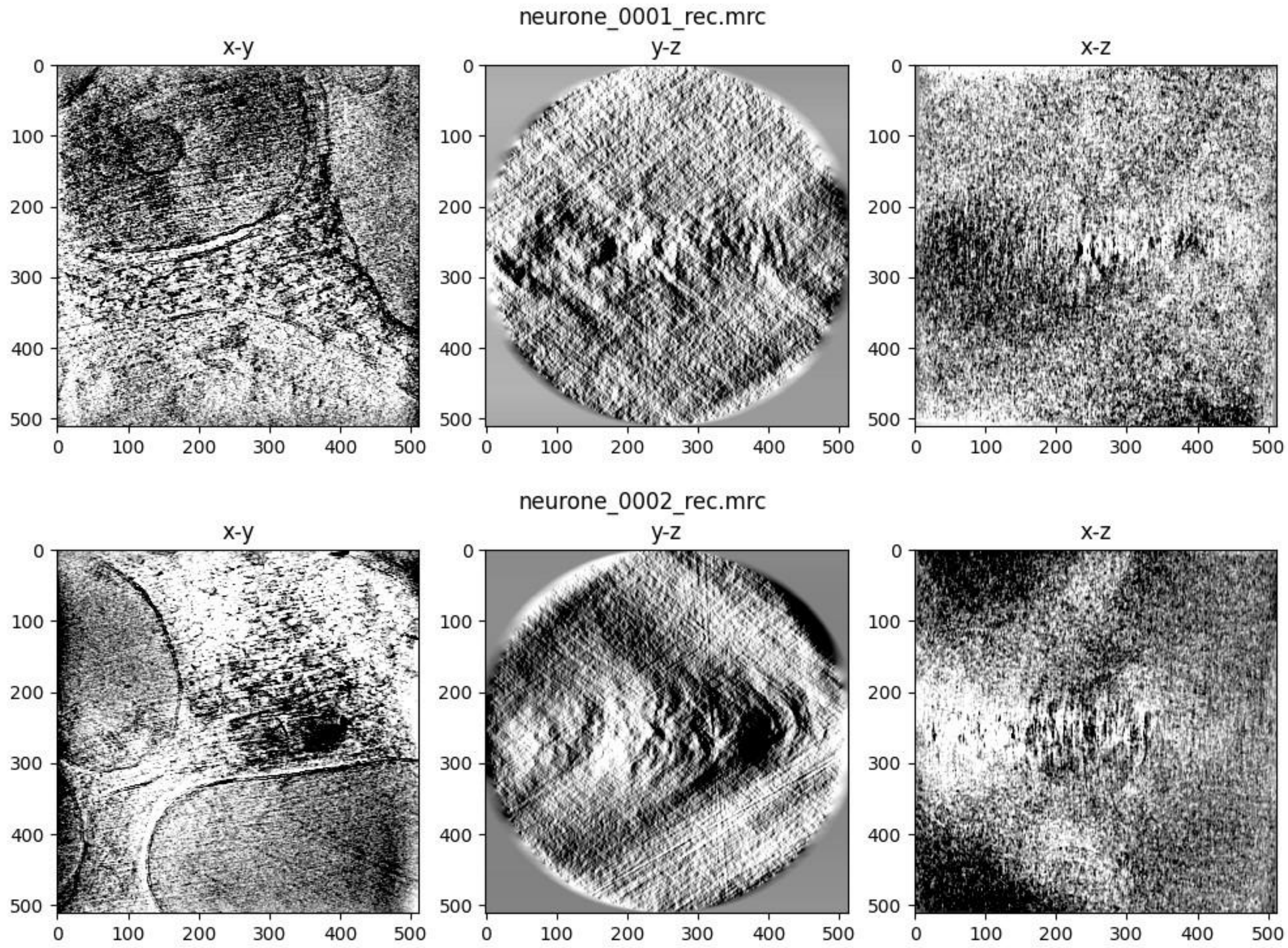

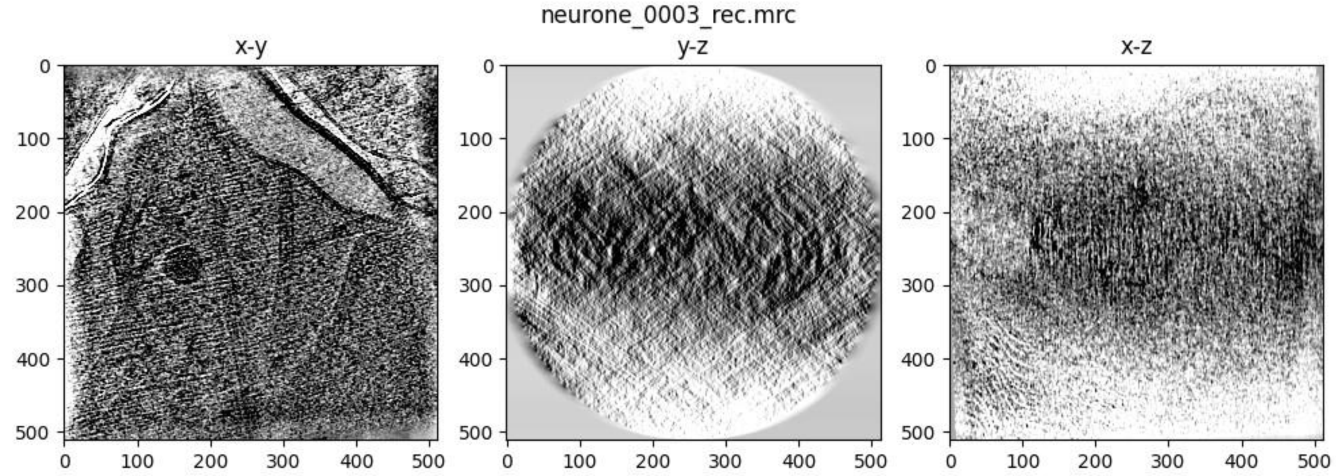
neurone_0003_rec.mrc
x-y
y-z
x-z

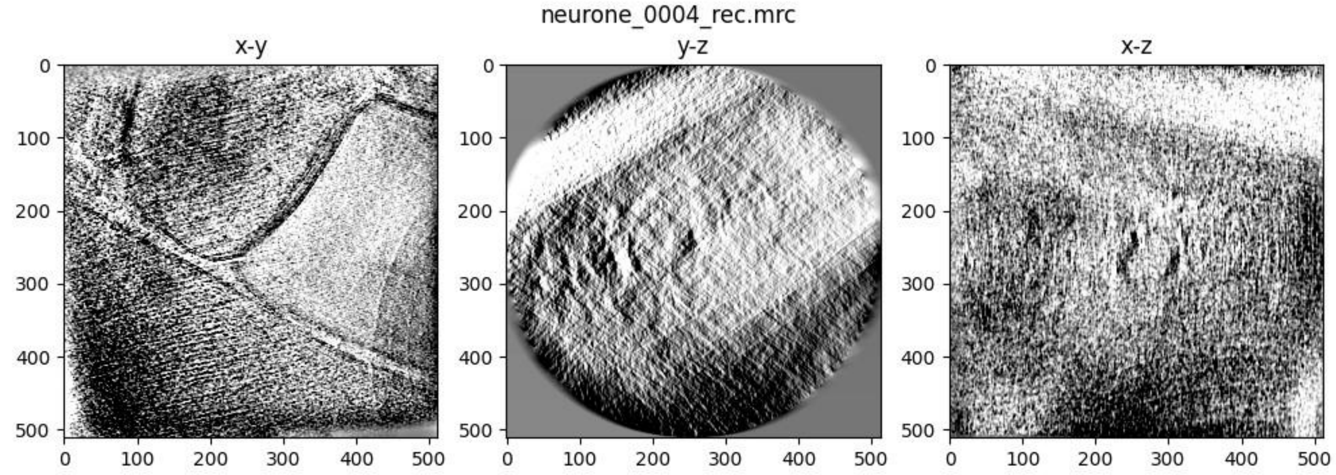
neurone_0004_rec.mrc
x-y
y-z
x-z

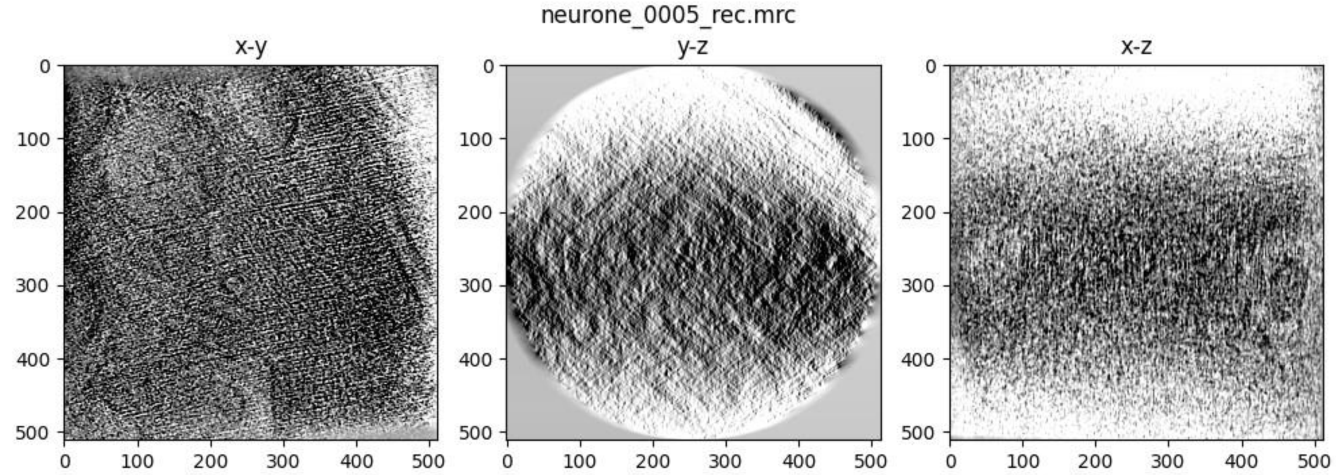
neurone_0005_rec.mrc
x-y
y-z
x-z

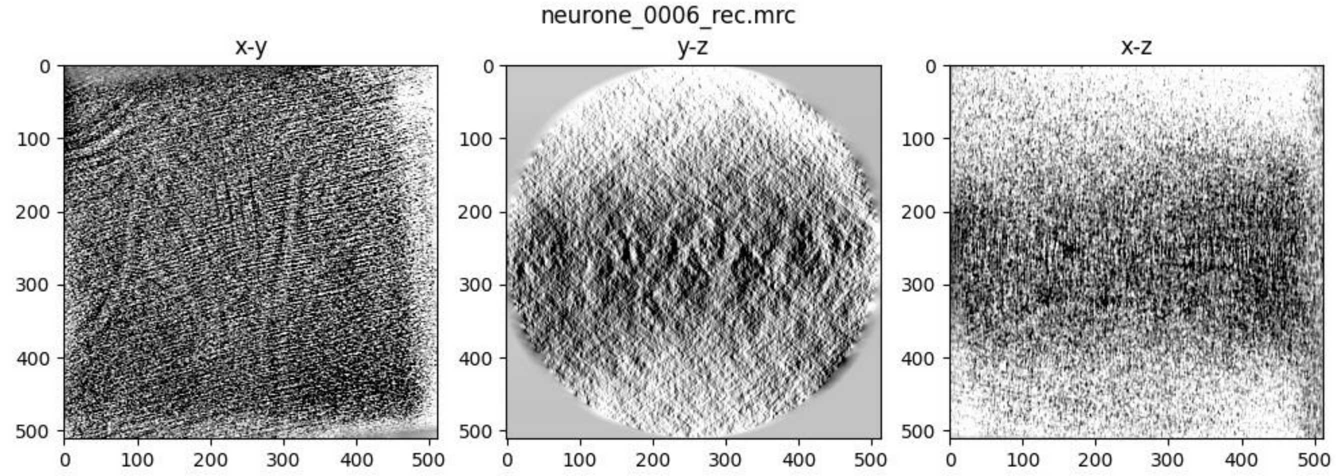
neurone_0006_rec.mrc
x-y
y-z
x-z

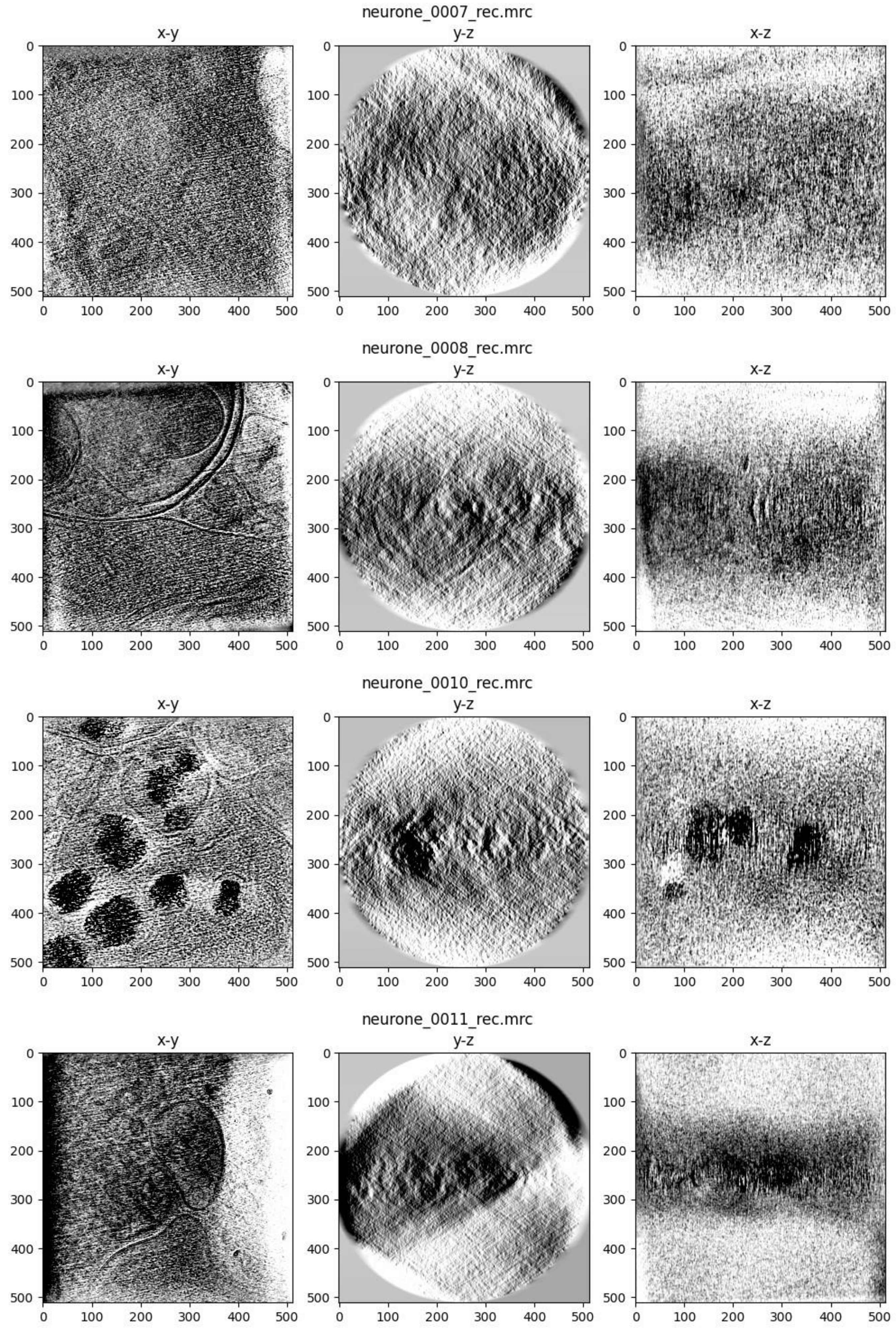
neurone_0007_rec.mrc
x-y
y-z
x-z
neurone_0008_rec.mrc
x-y
y-z
x-z
neurone_0010_rec.mrc
x-y
y-z
x-z
neurone_0011_rec.mrc
x-y
y-z
x-z

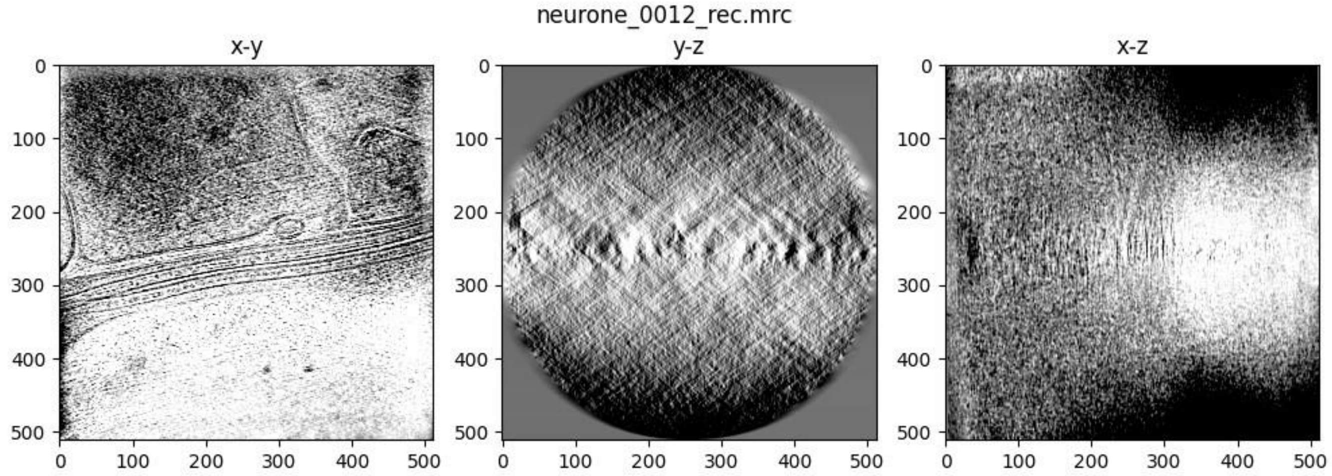
neurone_0012_rec.mrc
x-y
y-z
x-z

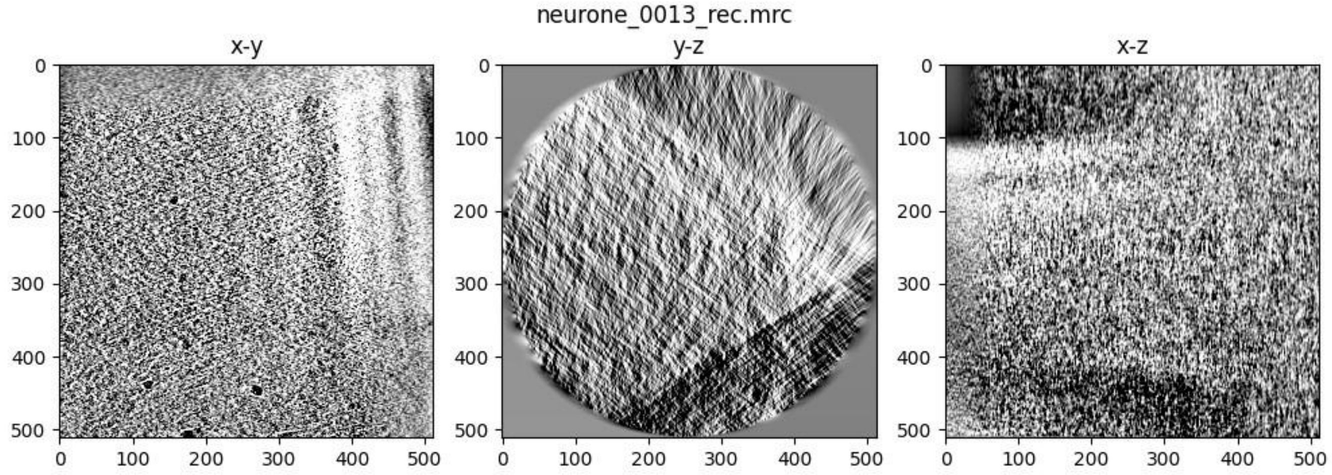
neurone_0013_rec.mrc
x-y
y-z
x-z

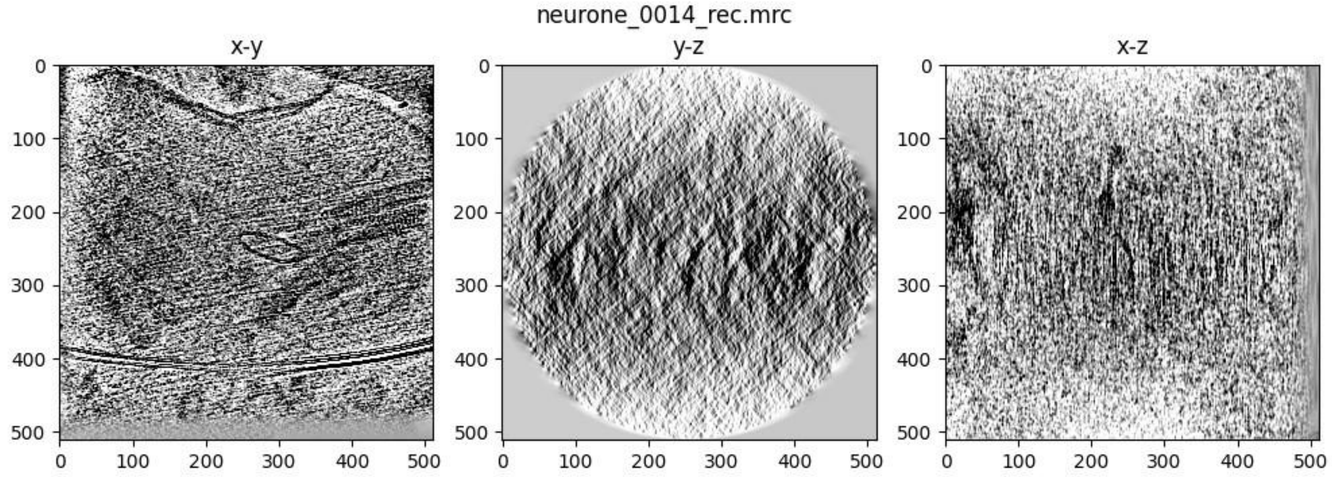
neurone_0014_rec.mrc
x-y
y-z
x-z

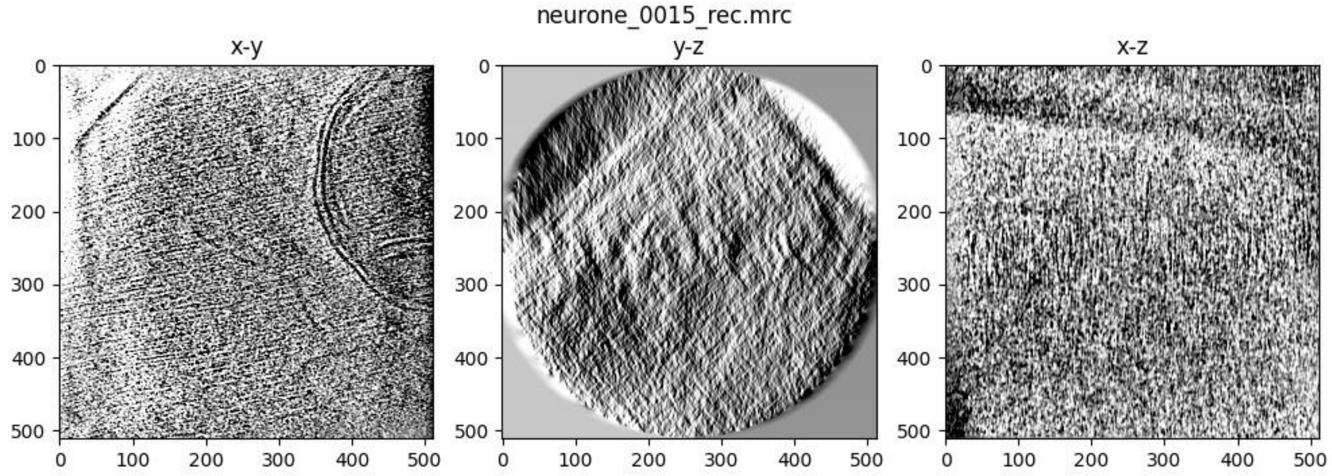
neurone_0015_rec.mrc
x-y
y-z
x-z

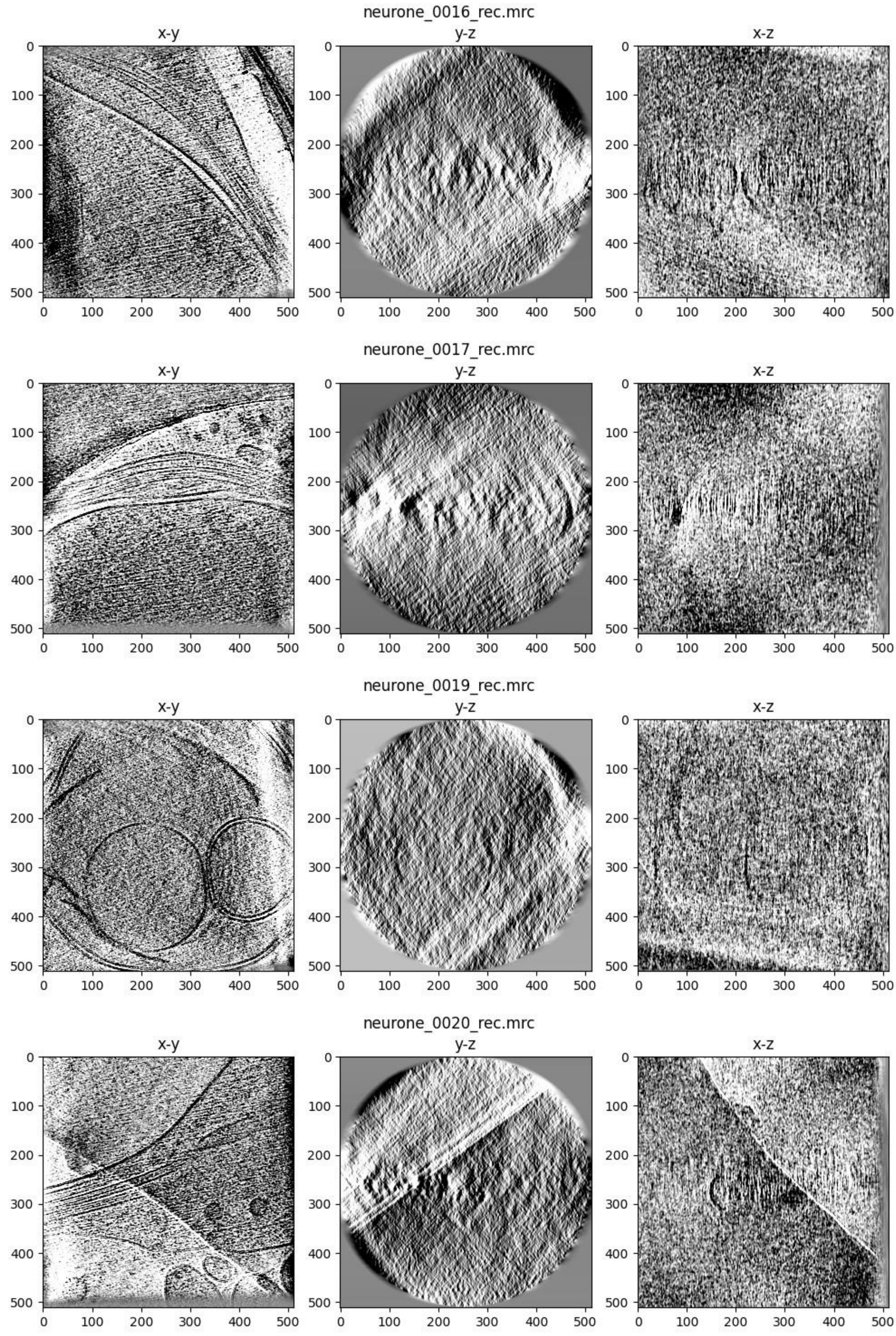
neurone_0016_rec.mrc
x-y
y-z
x-z
neurone_0017_rec.mrc
x-y
y-z
x-z
neurone_0019_rec.mrc
x-y
y-z
x-z
neurone_0020_rec.mrc
x-y
y-z
x-z

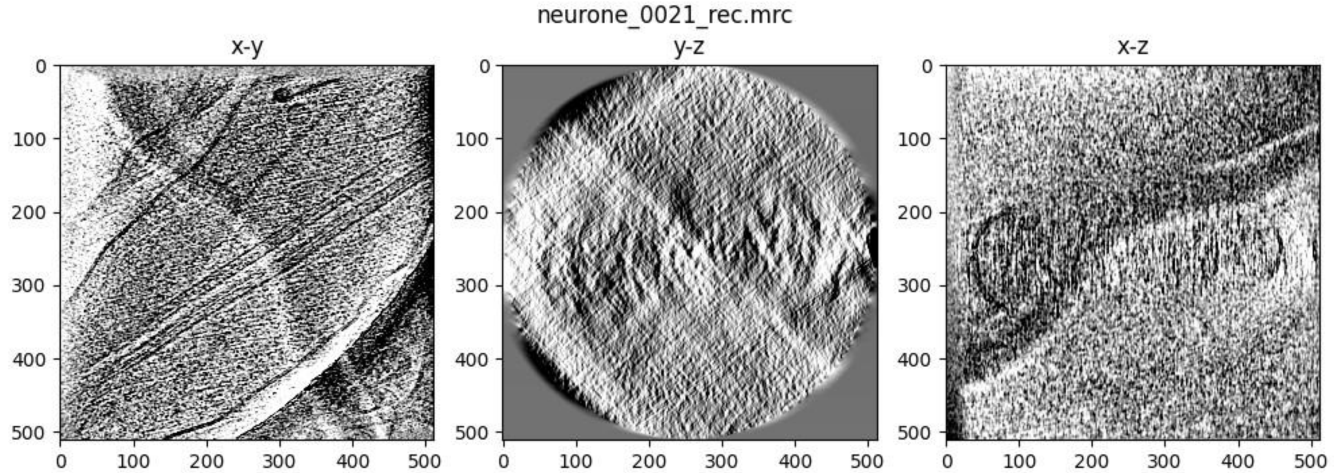
neurone_0021_rec.mrc
x-y
y-z
x-z
0
100
200
300
400
500

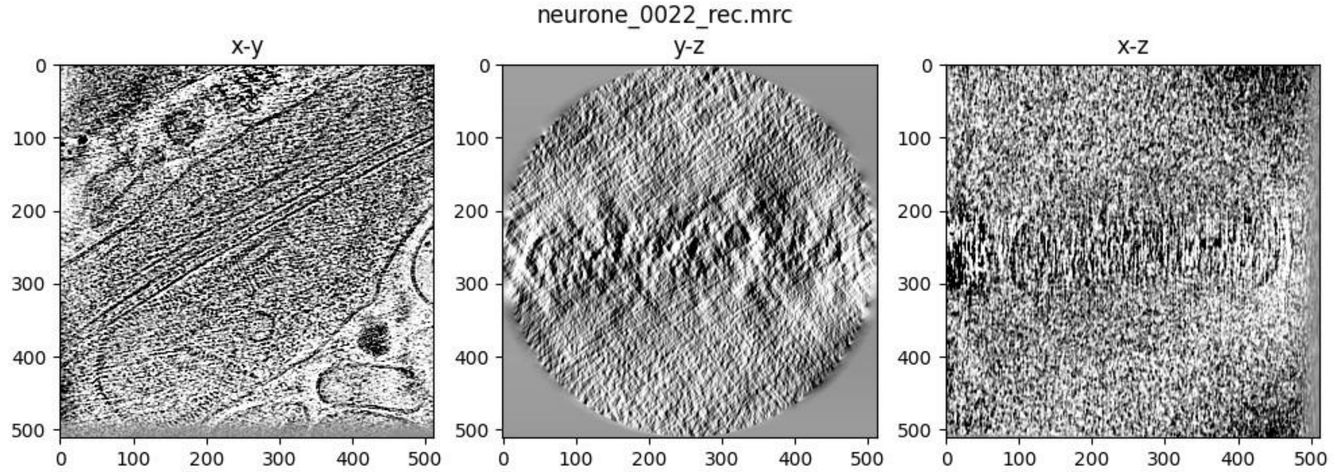
neurone_0022_rec.mrc
x-y
y-z
x-z
0
100
200
300
400
500

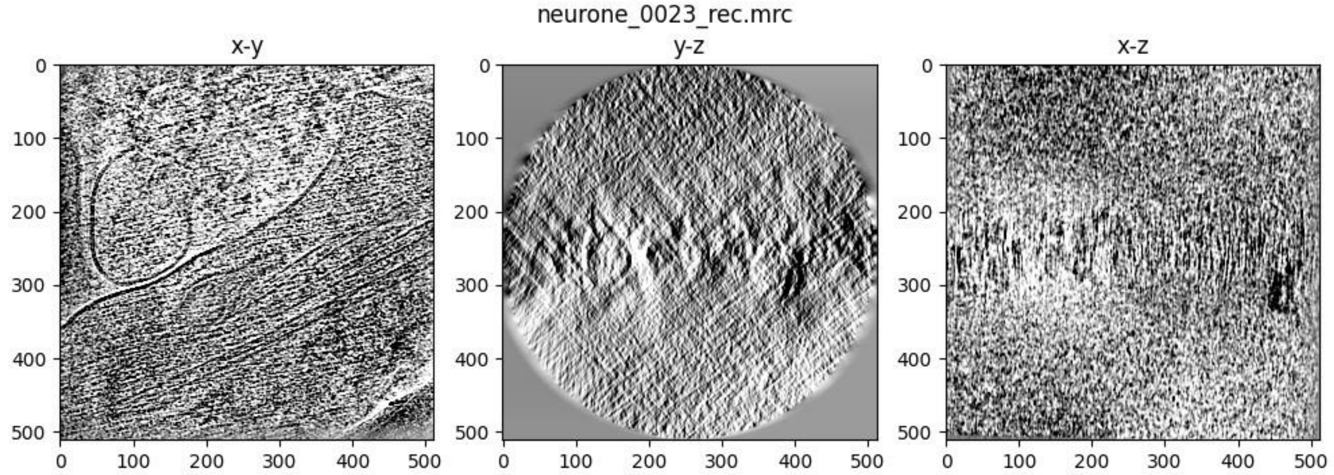
neurone_0023_rec.mrc
x-y
y-z
x-z
0
100
200
300
400
500

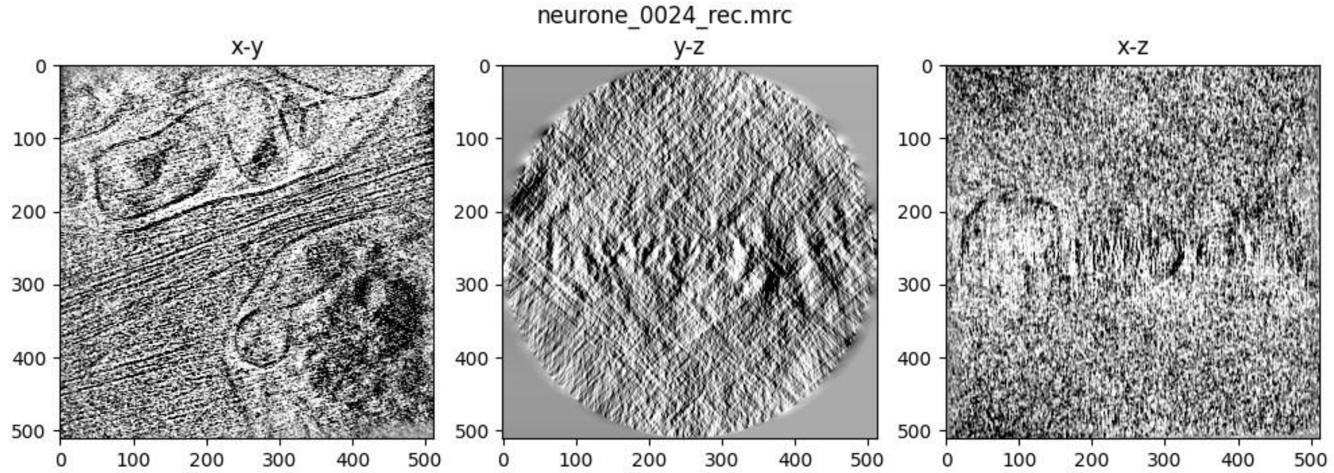
neurone_0024_rec.mrc
x-y
y-z
x-z
0
100
200
300
400
500

### AreTomo

AreTomo was used in this workflow. Please cite:

1. Shawn Zheng, Georg Wolff, Garrett Greenan, et al. AreTomo: An integrated software package for automated marker-free, motion-corrected cryo-electron tomographic alignment and reconstruction, *JSB: X* **6(100068)** , ISSN 2590-1524, (2022) DOI | ScienceDirect

For more information, please see https://msg.ucsf.edu/software.

### Tomographic Reconstruction (AreTomo)

Vol Z = 1000

Sample thickness = -1

Pixel size = 1.47

Reconstruction algorithm = SART

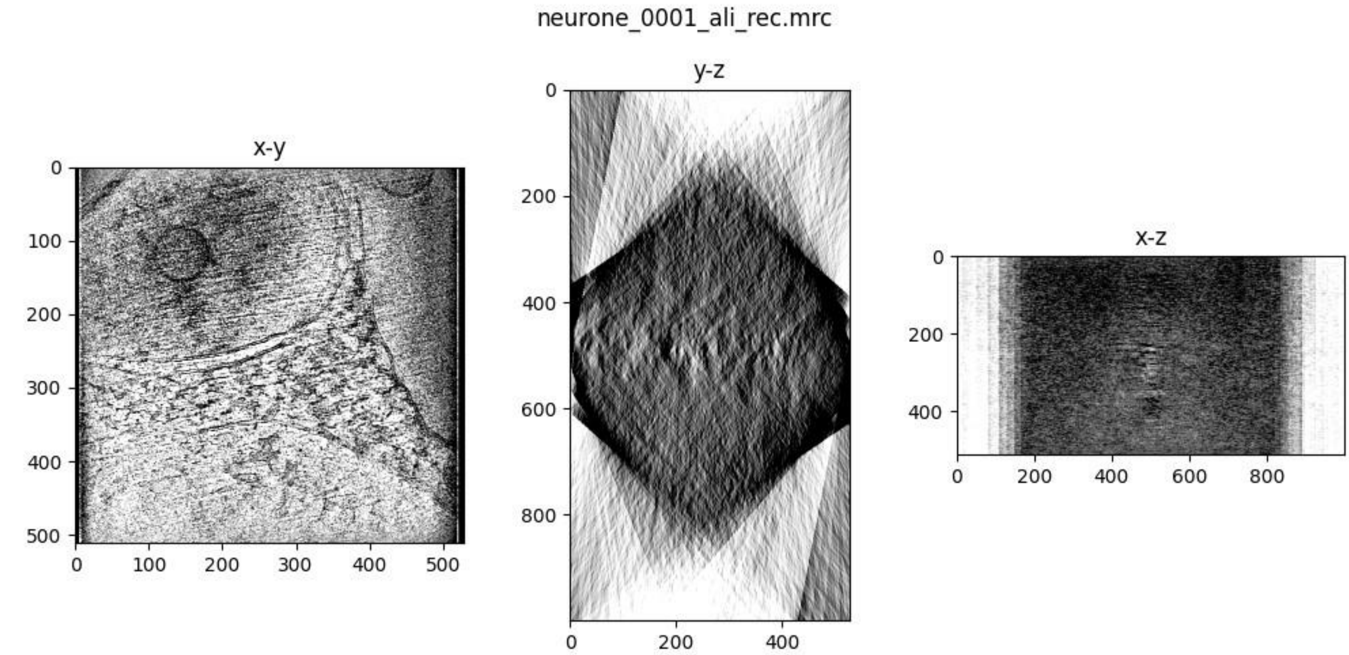
neurone_0001_ali_rec.mrc
x-y
y-z
x-z

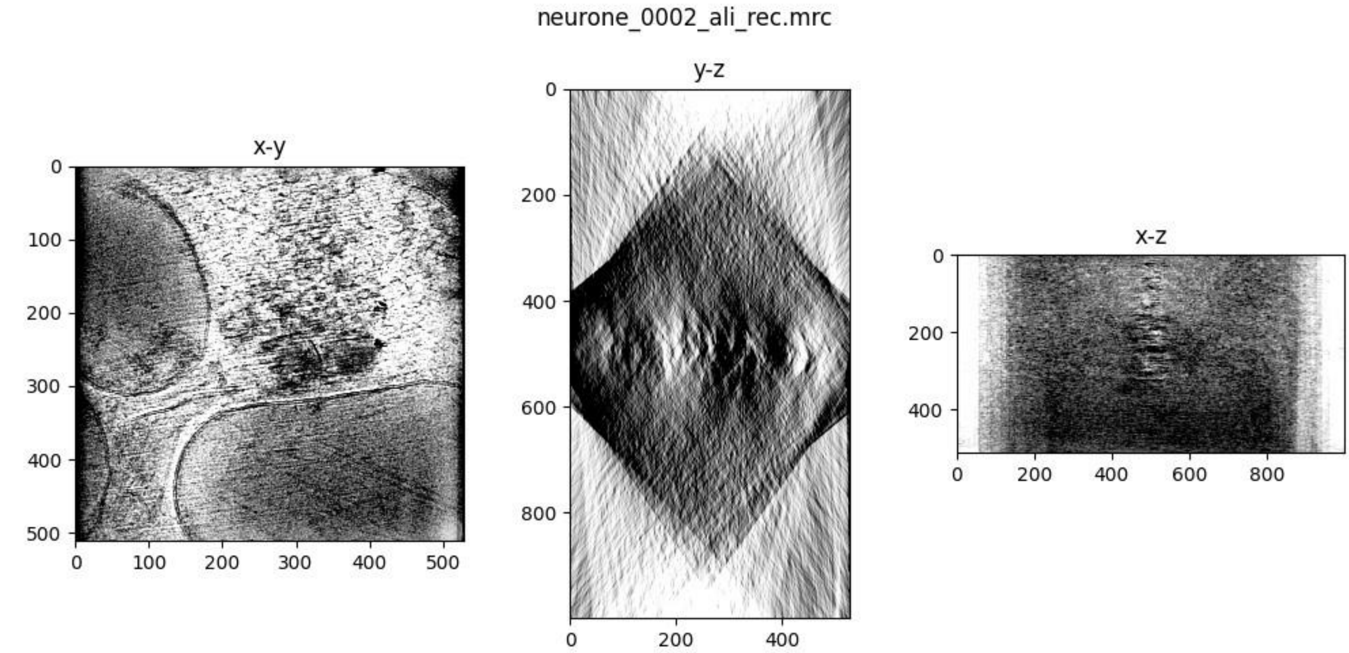
neurone_0002_ali_rec.mrc
x-y
y-z
x-z

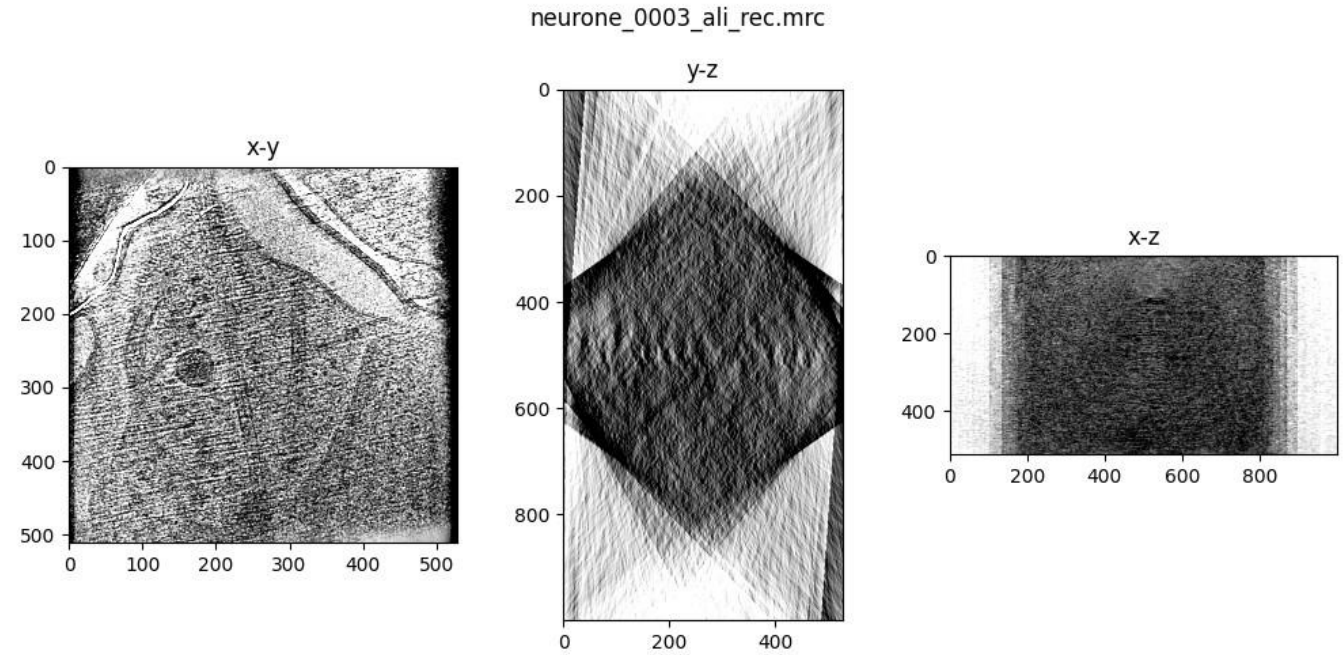
neurone_0003_ali_rec.mrc
x-y
y-z
x-z

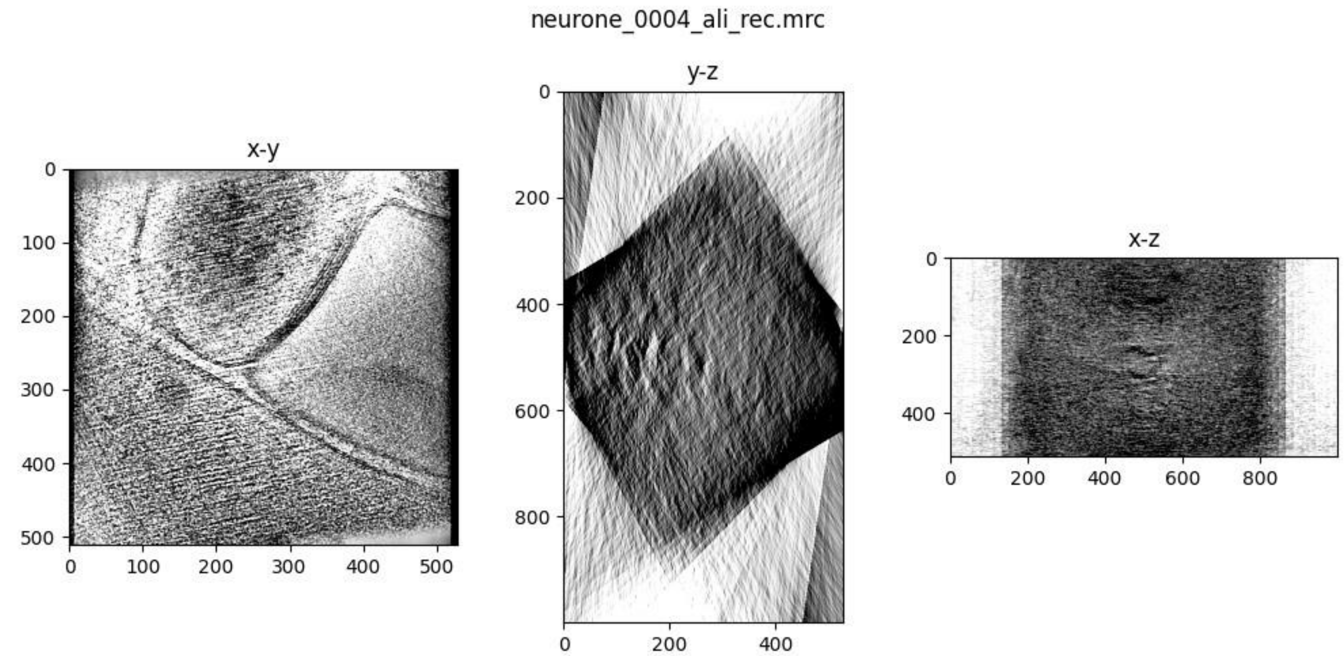
neurone_0004_ali_rec.mrc
x-y
y-z
x-z

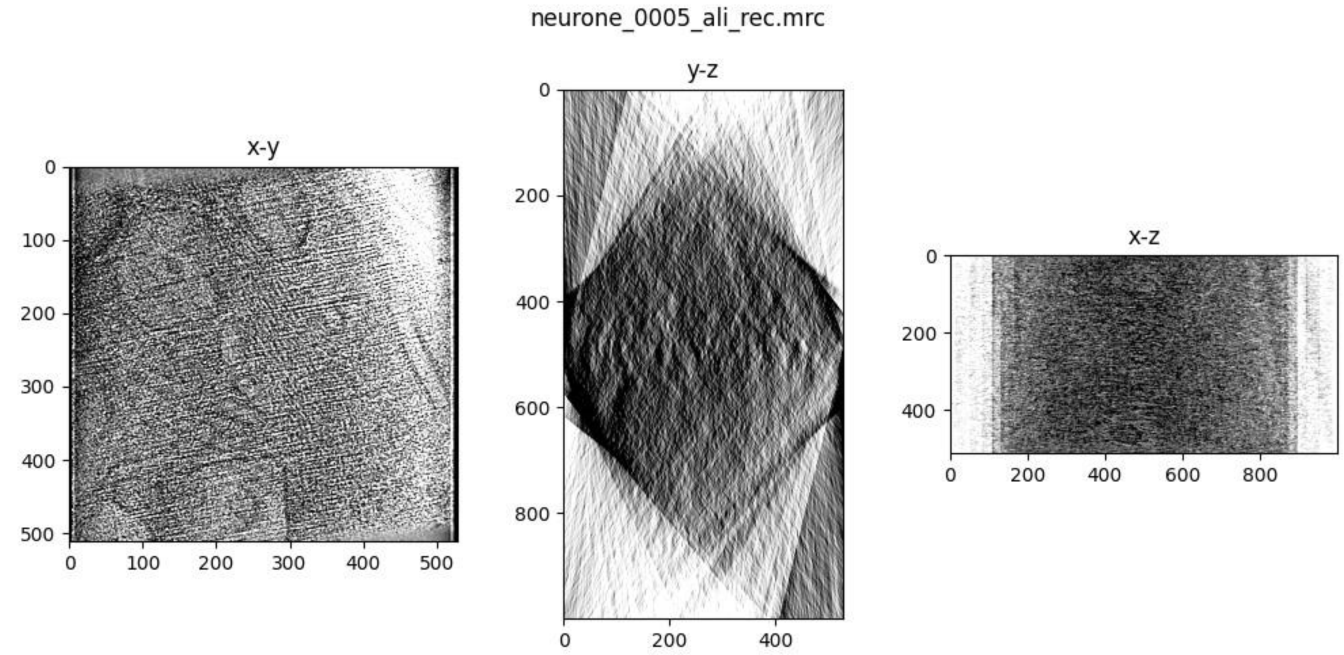
neurone_0005_ali_rec.mrc
x-y
y-z
x-z

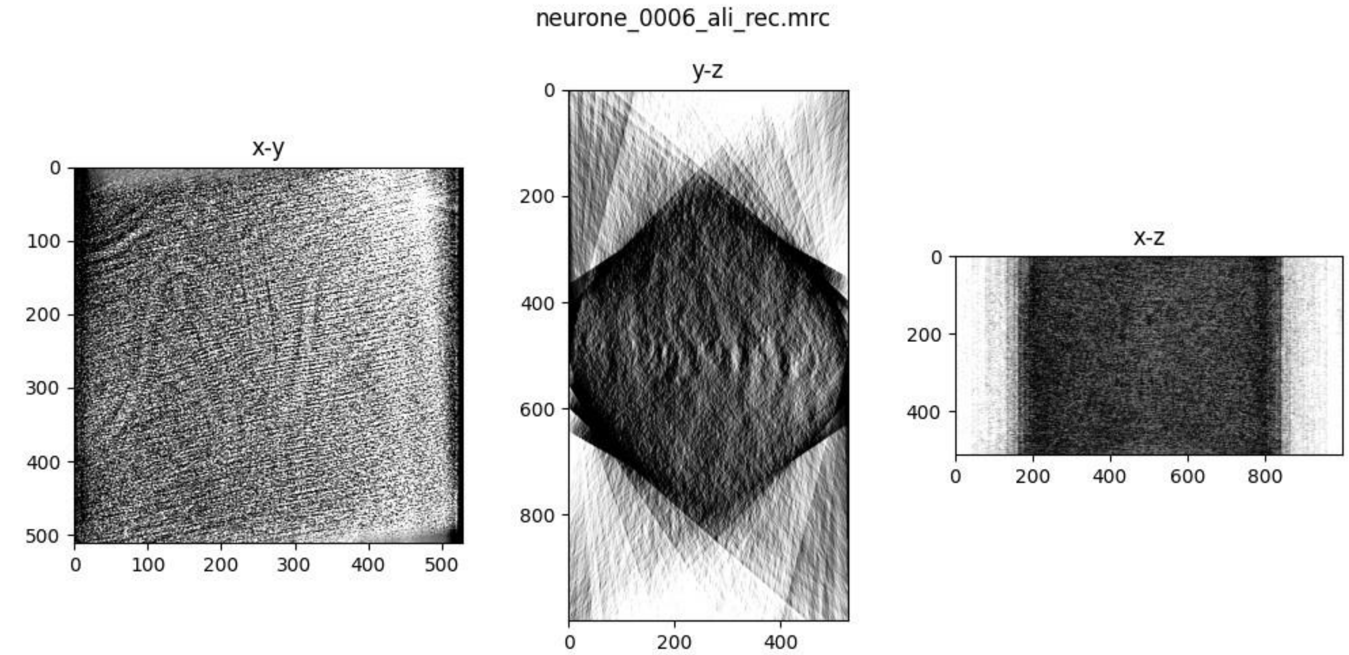
neurone_0006_ali_rec.mrc
x-y
y-z
x-z

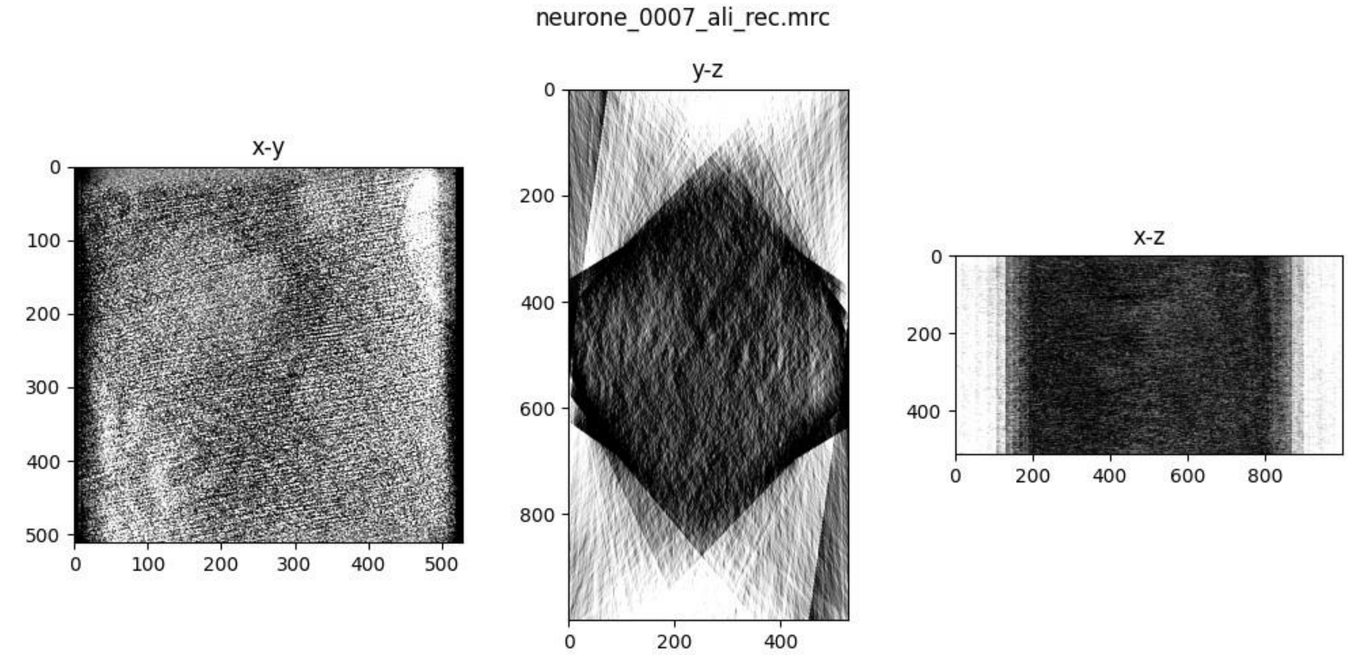
neurone_0007_ali_rec.mrc
x-y
y-z
x-z

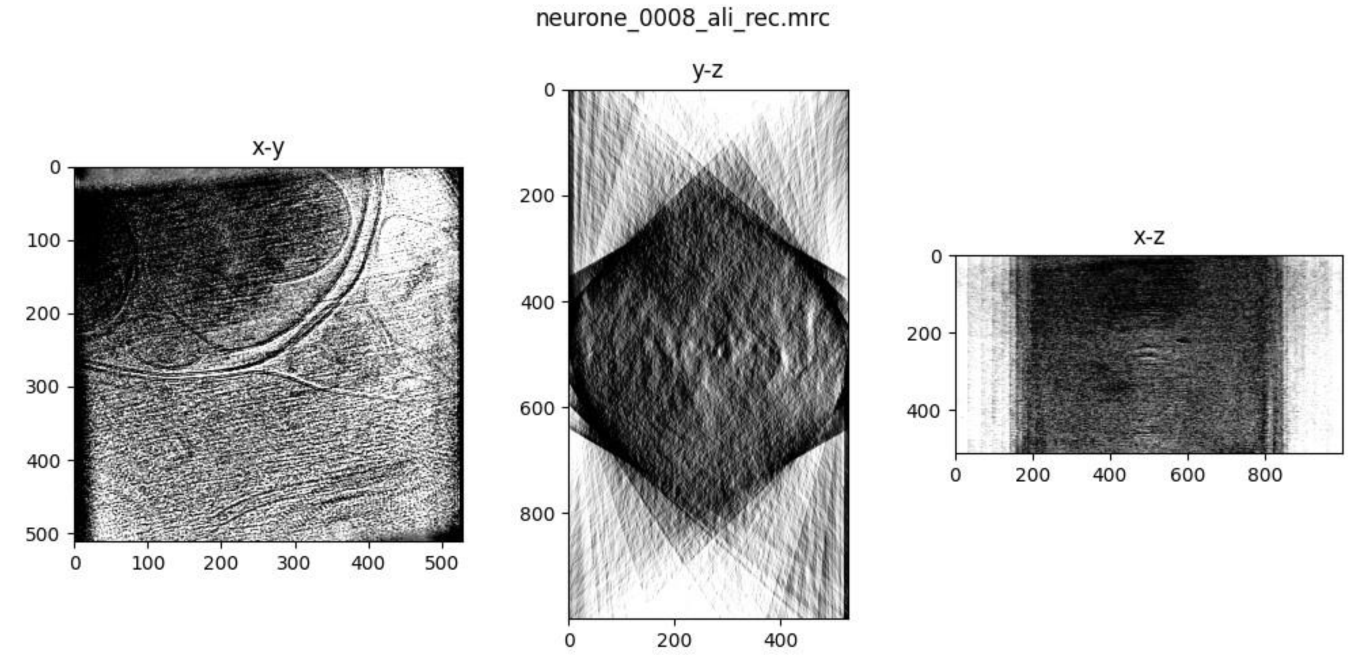
neurone_0008_ali_rec.mrc
x-y
y-z
x-z

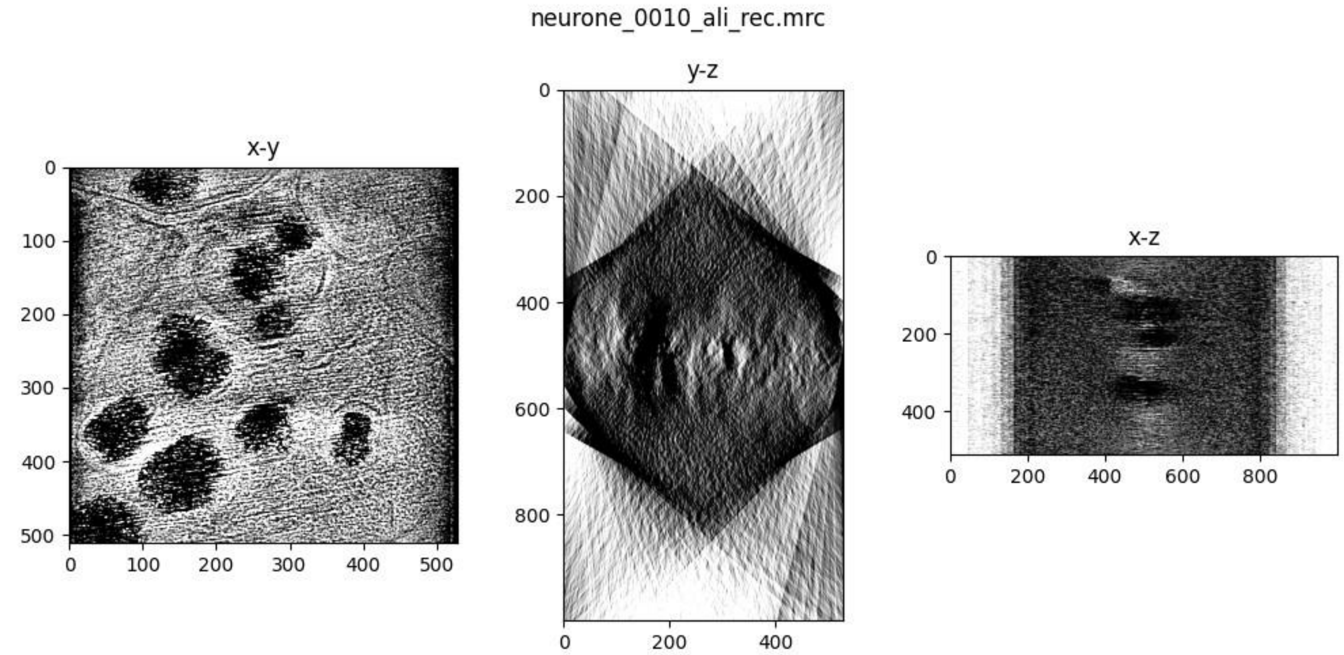
neurone_0010_ali_rec.mrc
x-y
y-z
x-z

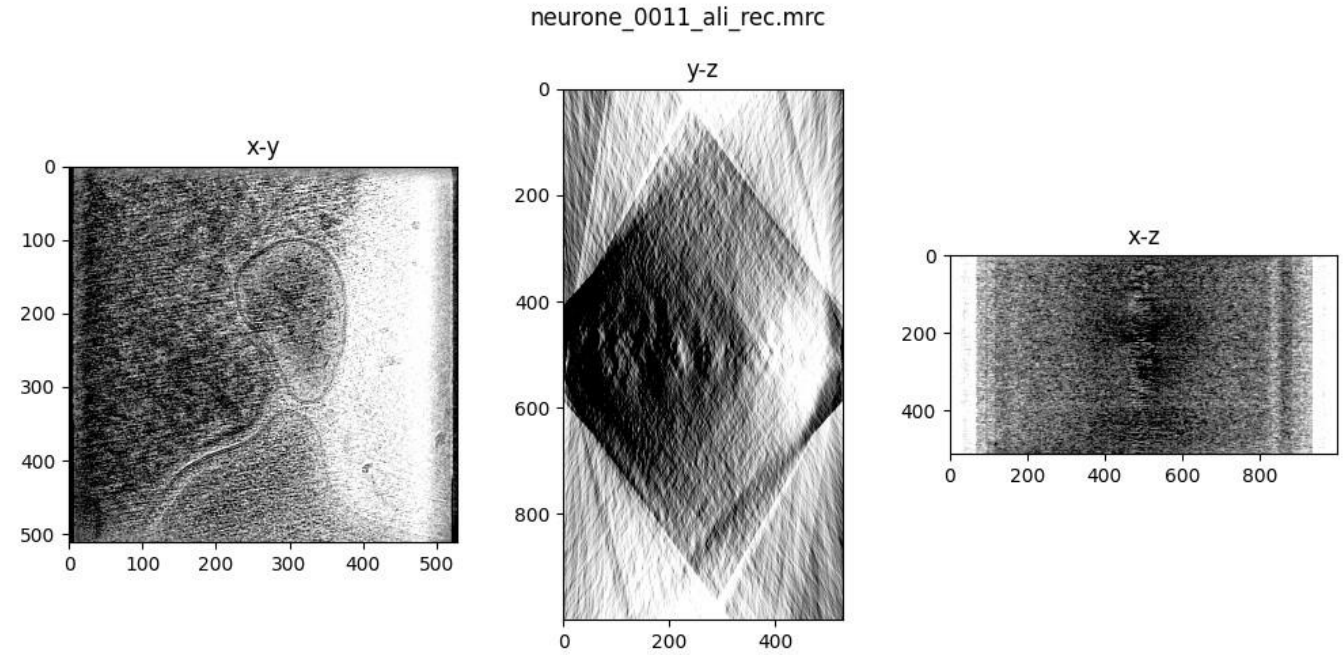
neurone_0011_ali_rec.mrc
x-y
y-z
x-z

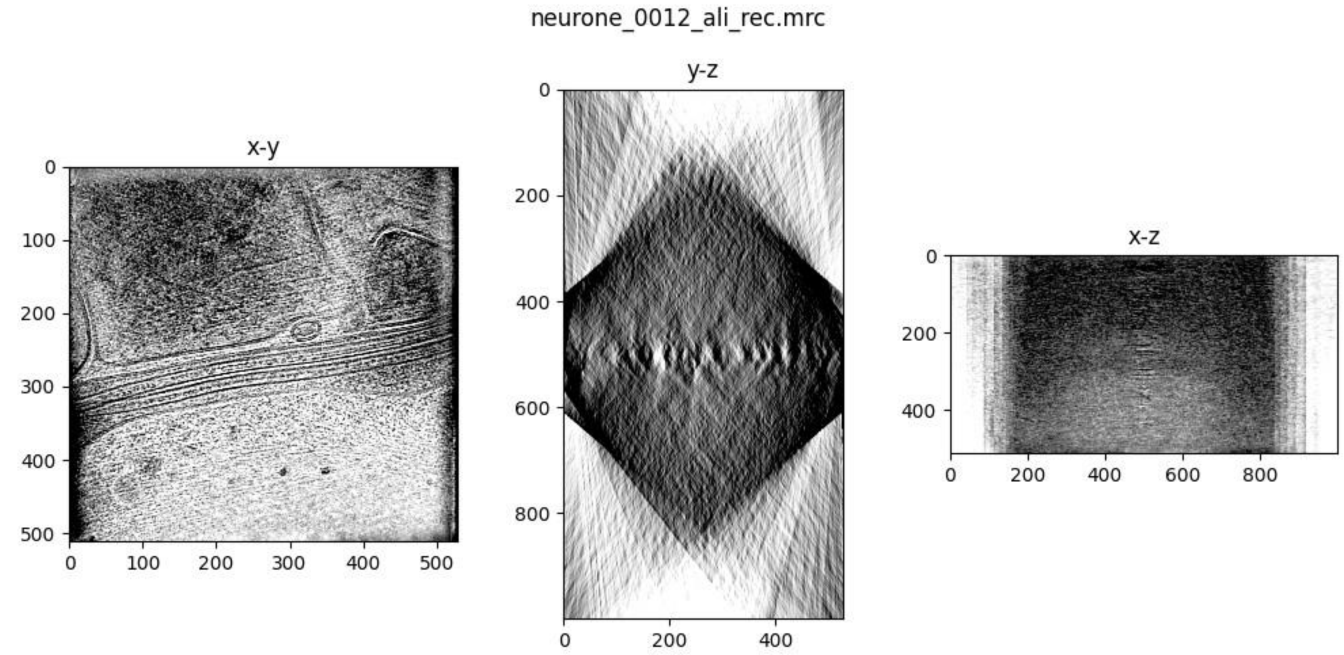
neurone_0012_ali_rec.mrc
x-y
y-z
x-z

neurone_0013_ali_rec.mrc

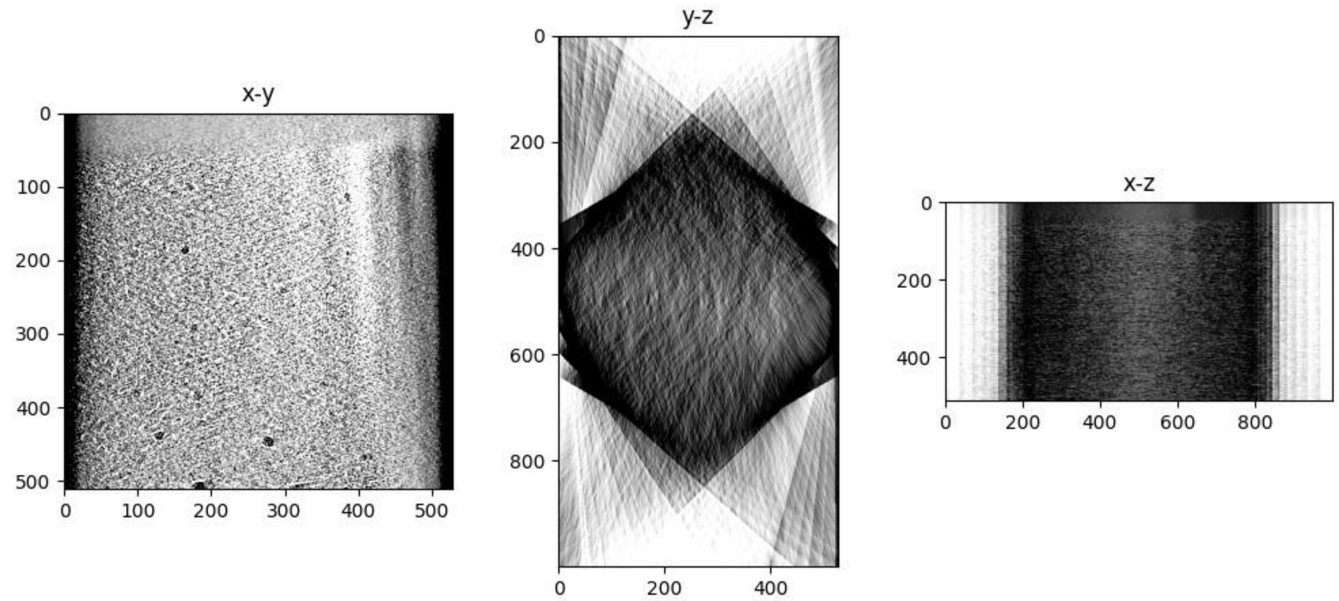


neurone_0014_ali_rec.mrc

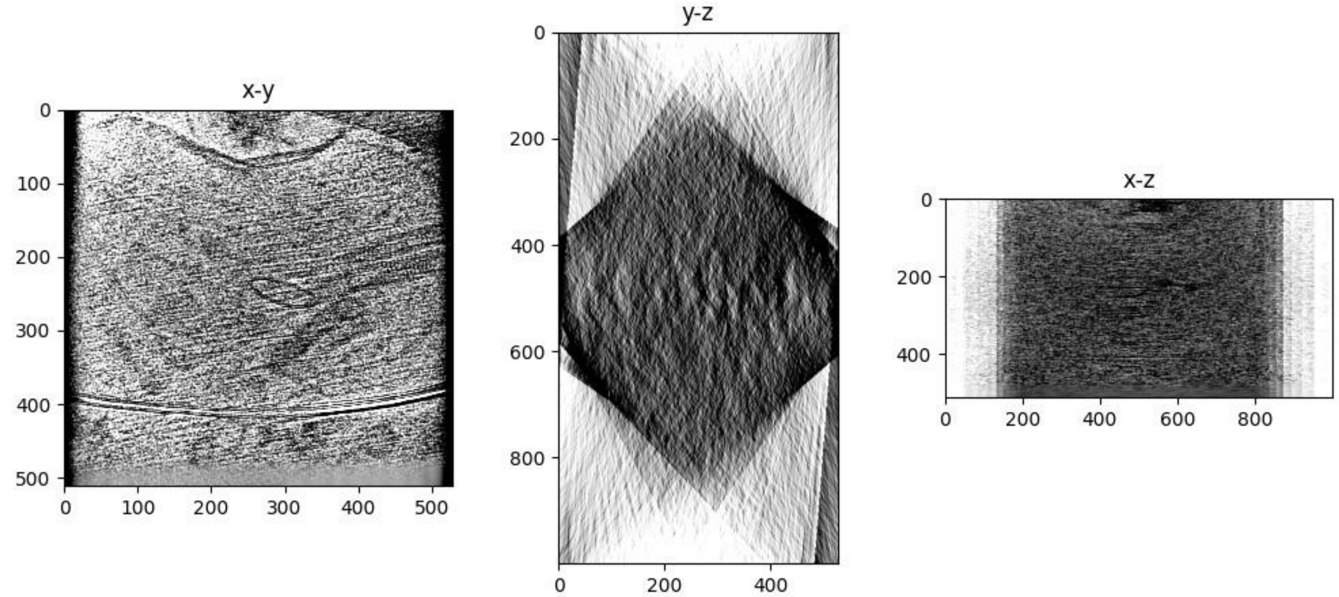


neurone_0015_ali_rec.mrc

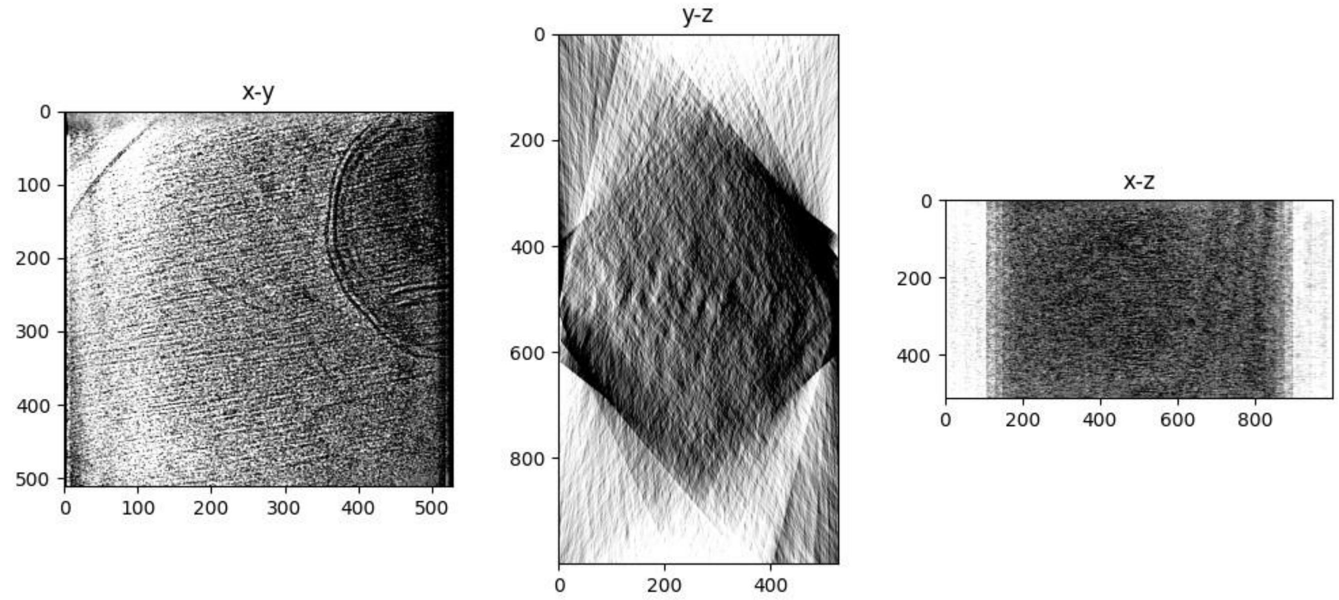

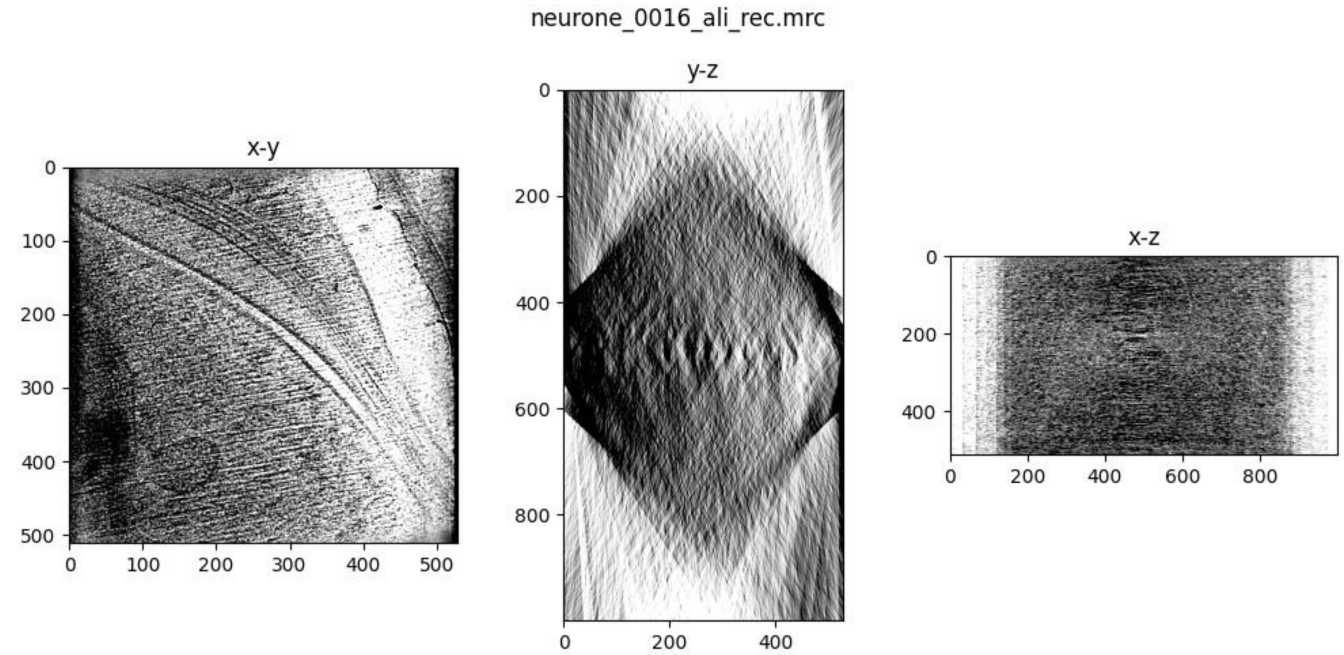
neurone_0016_ali_rec.mrc
x-y
y-z
x-z

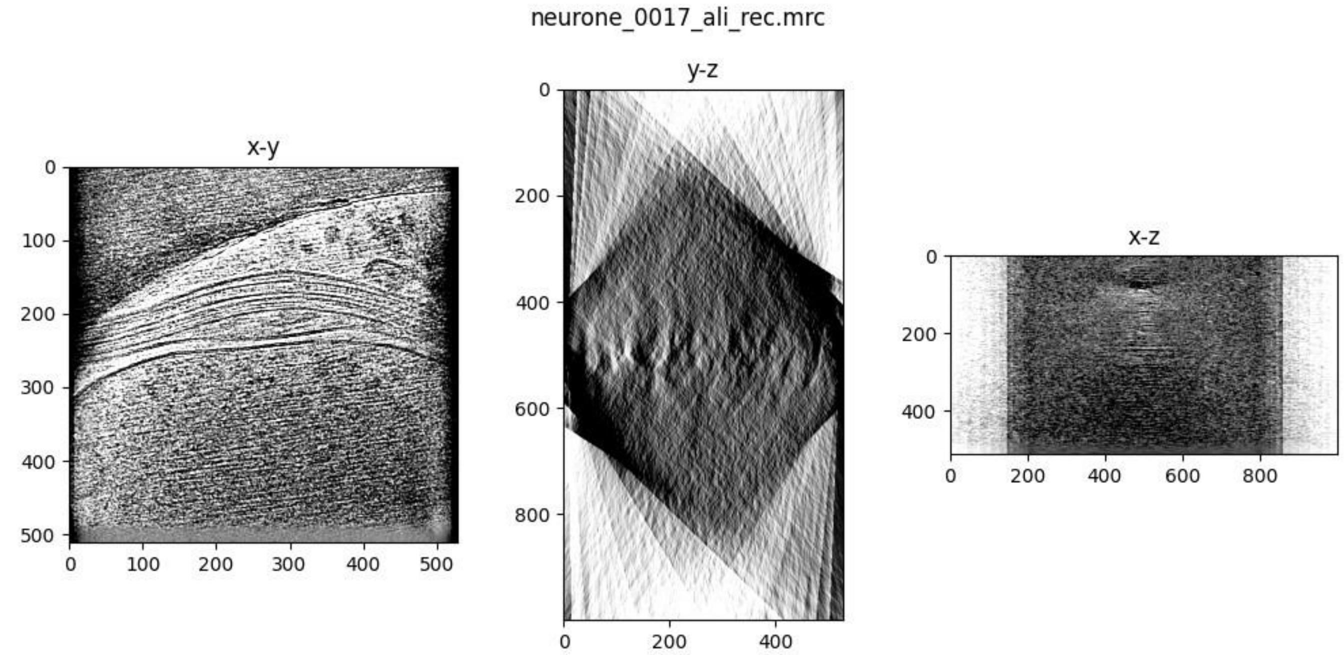
neurone_0017_ali_rec.mrc
x-y
y-z
x-z

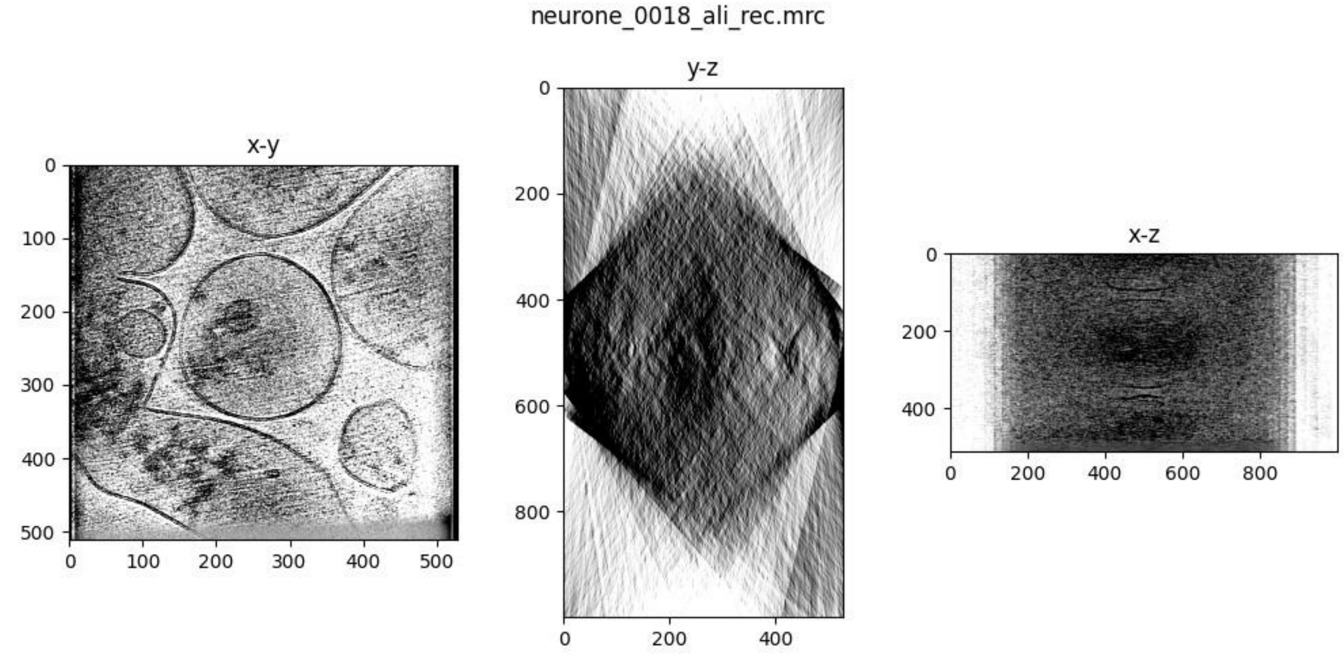
neurone_0018_ali_rec.mrc
x-y
y-z
x-z

neurone_0019_ali_rec.mrc

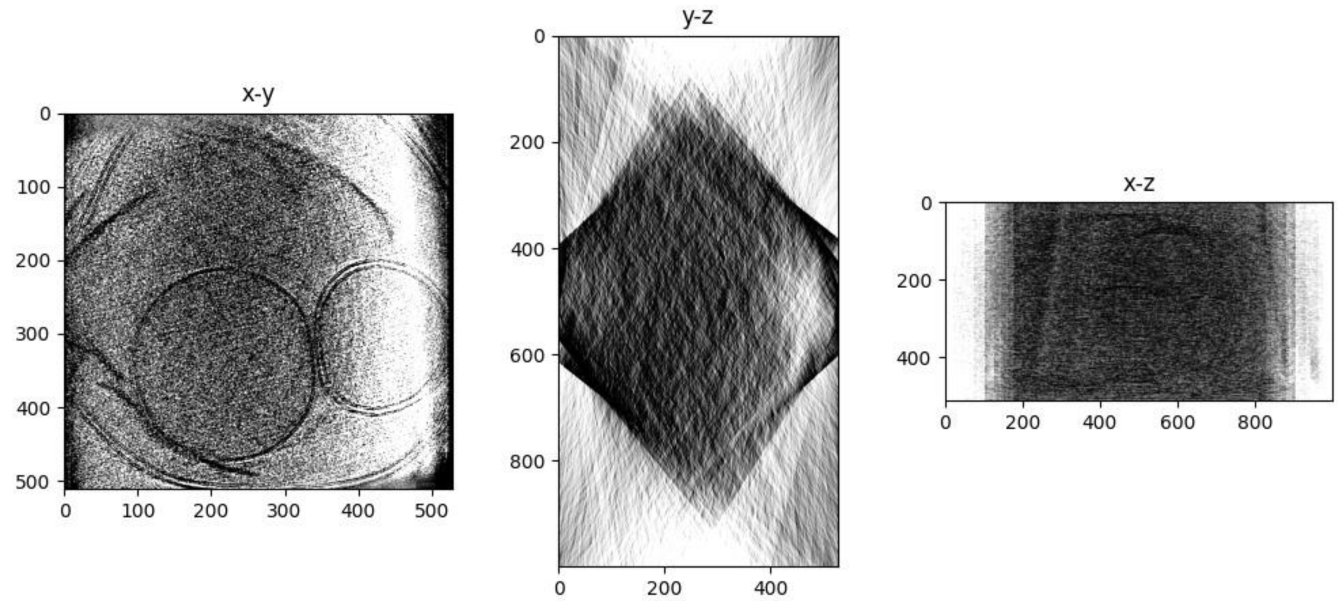


neurone_0020_ali_rec.mrc

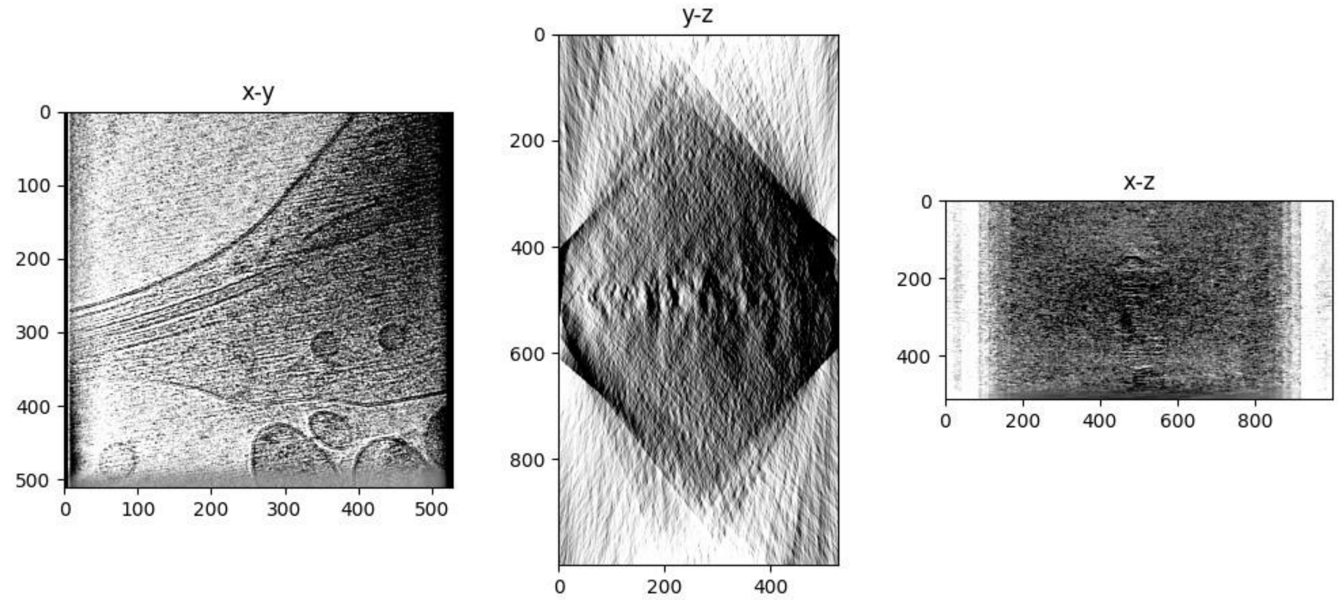


neurone_0021_ali_rec.mrc

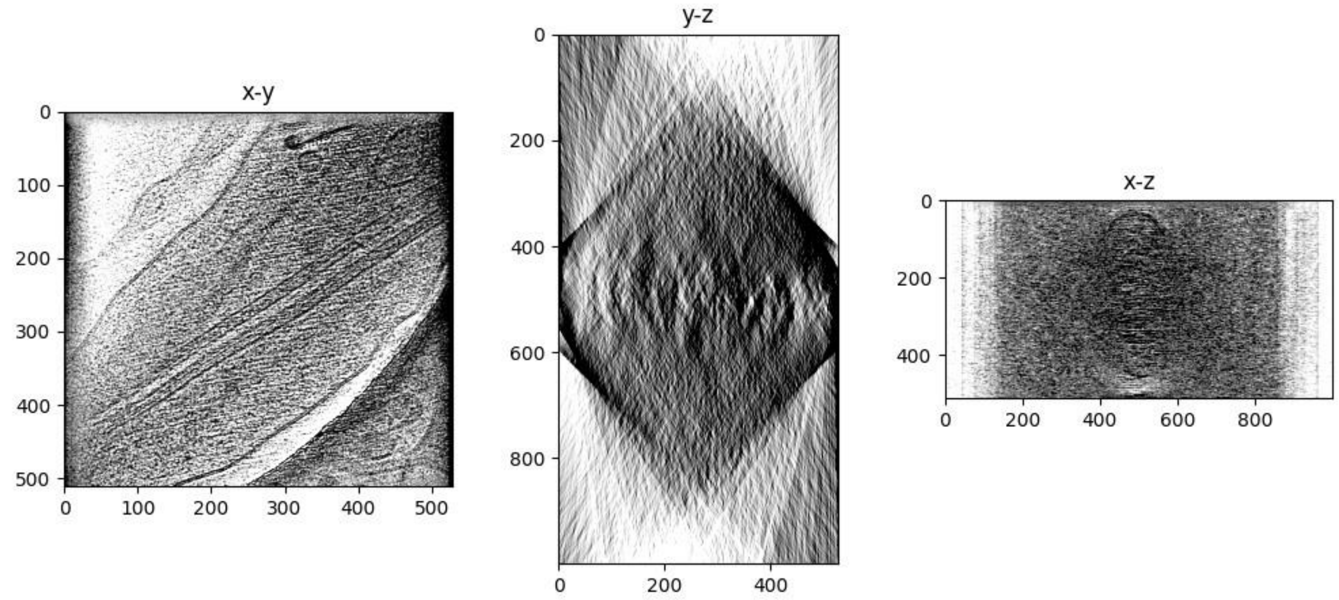

neurone_0022_ali_rec.mrc

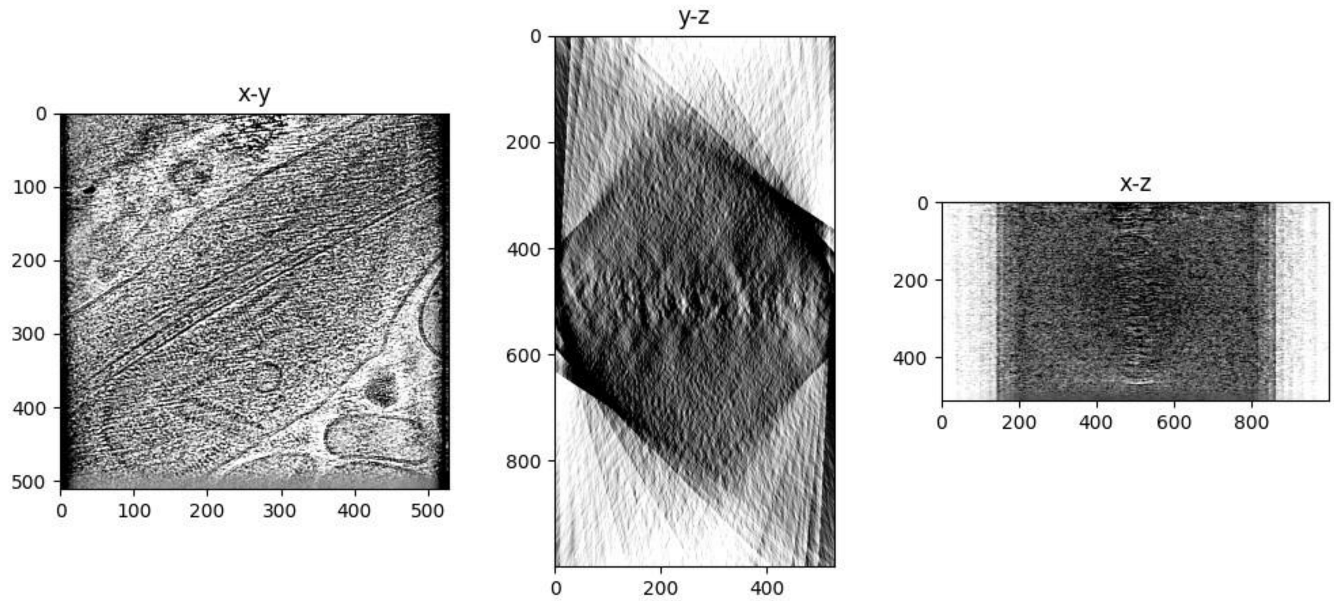


neurone_0023_ali_rec.mrc

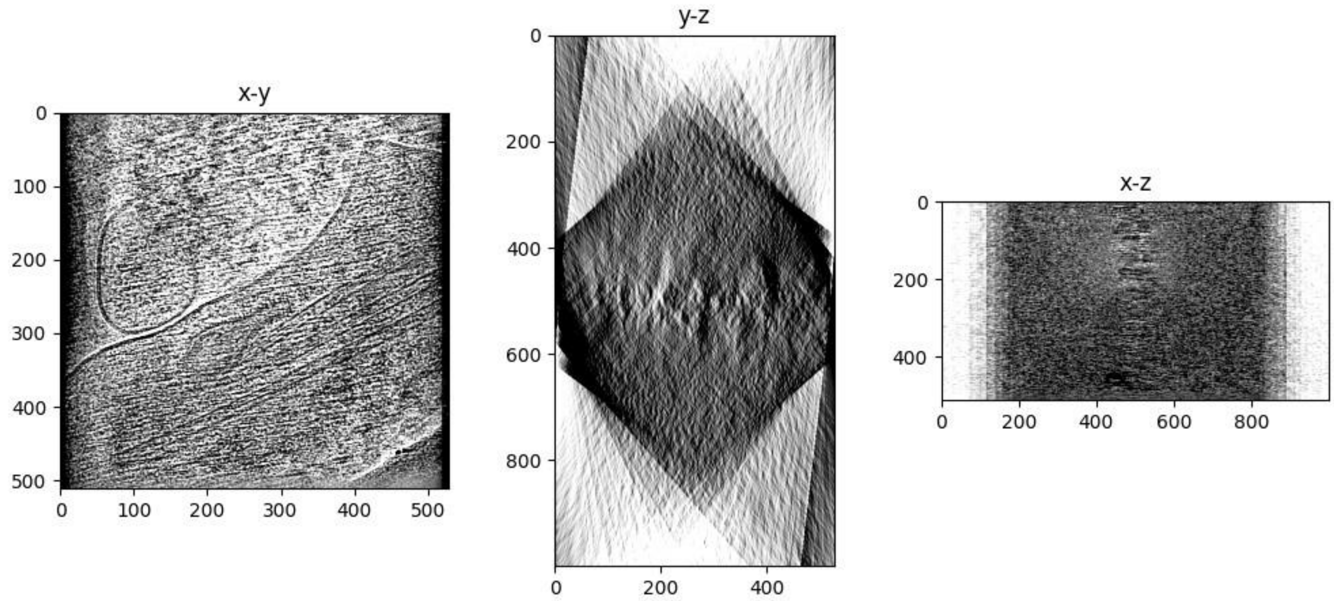


neurone_0024_ali_rec.mrc

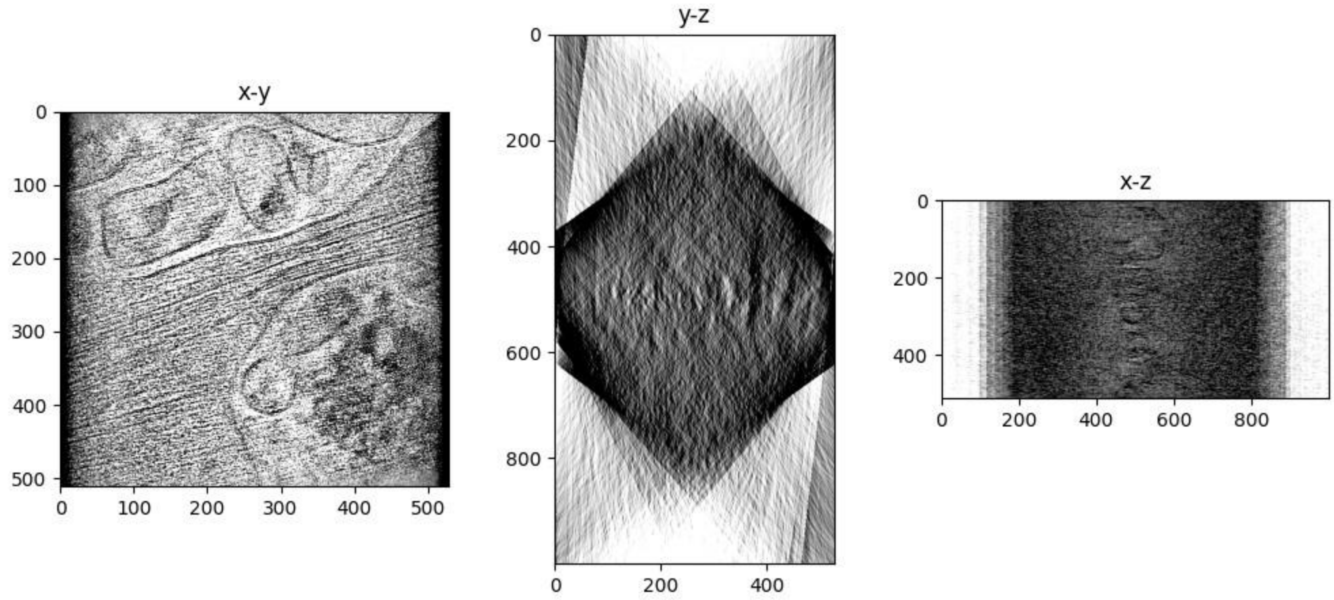

### Savu Reconstruction

Reconstruction was performed with Savu 4.0.

Please cite:


1. Wadeson, N and Basham, M. Savu: A Python-based, MPI Framework for Simultaneous Processing of Multiple, N-dimensional, Large Tomography Datasets. *arXiv* **1610.08015**. (2016)
2. Wadeson, N, Verschoyle, J, Kazantsev, D., et al. DiamondLightSource/Savu: Version 4.0. *Zenodo*. (2021) DOI 10.5281/zenodo.5095360.
3. The HDF Group. Hierarchical Data Format, version 5. (1997-2016)
4. Walker, D and Dongarra, J. MPI: a standard message passing interface. *Supercomputer* **12**: 56-68. (1996)
5. Van Aarle, W, Palenstijn, W, Cant, J, et al. Fast and flexible X-ray tomography using the ASTRA toolbox. *Optics express* **24(22)**: 25129-25147. (2016)
6. Van Aarle, W, Palenstijn, W, De Beenhouwer, J, et al. The ASTRA Toolbox: A platform for advanced algorithm development in electron tomography. *Ultramicroscopy* **157**: 35-47. (2015)
7. Palenstijn, WJ and Batenburg, KJ and Sijbers, J. Performance improvements for iterative electron tomography reconstruction using graphics processing units (GPUs). *JSB* **176(2)** 250-253. (2011) For more information, please see https://savu.readthedocs.io/en/latest/


**Note**

Savu was developed by the X-ray tomography community, so the conventional relationship between grey values and density is reversed compared to EM, i.e., the colours are reversed. The visualisations in this report have had their colours inverted to match the EM convention.

Reconstruction algorithm used: CGLS_CUDA

#### Tomogram thumbnails

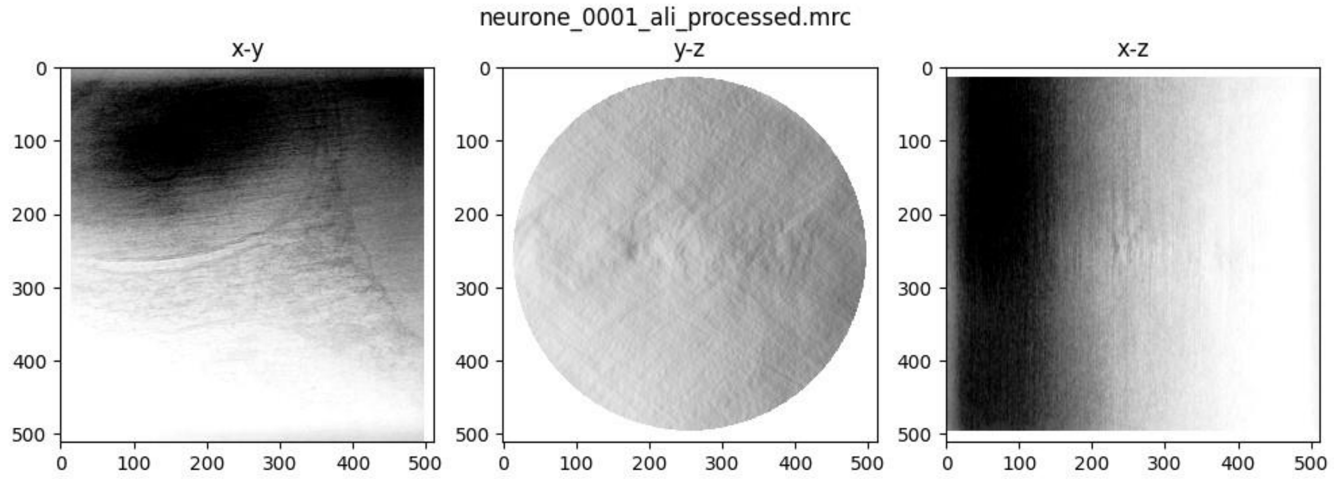

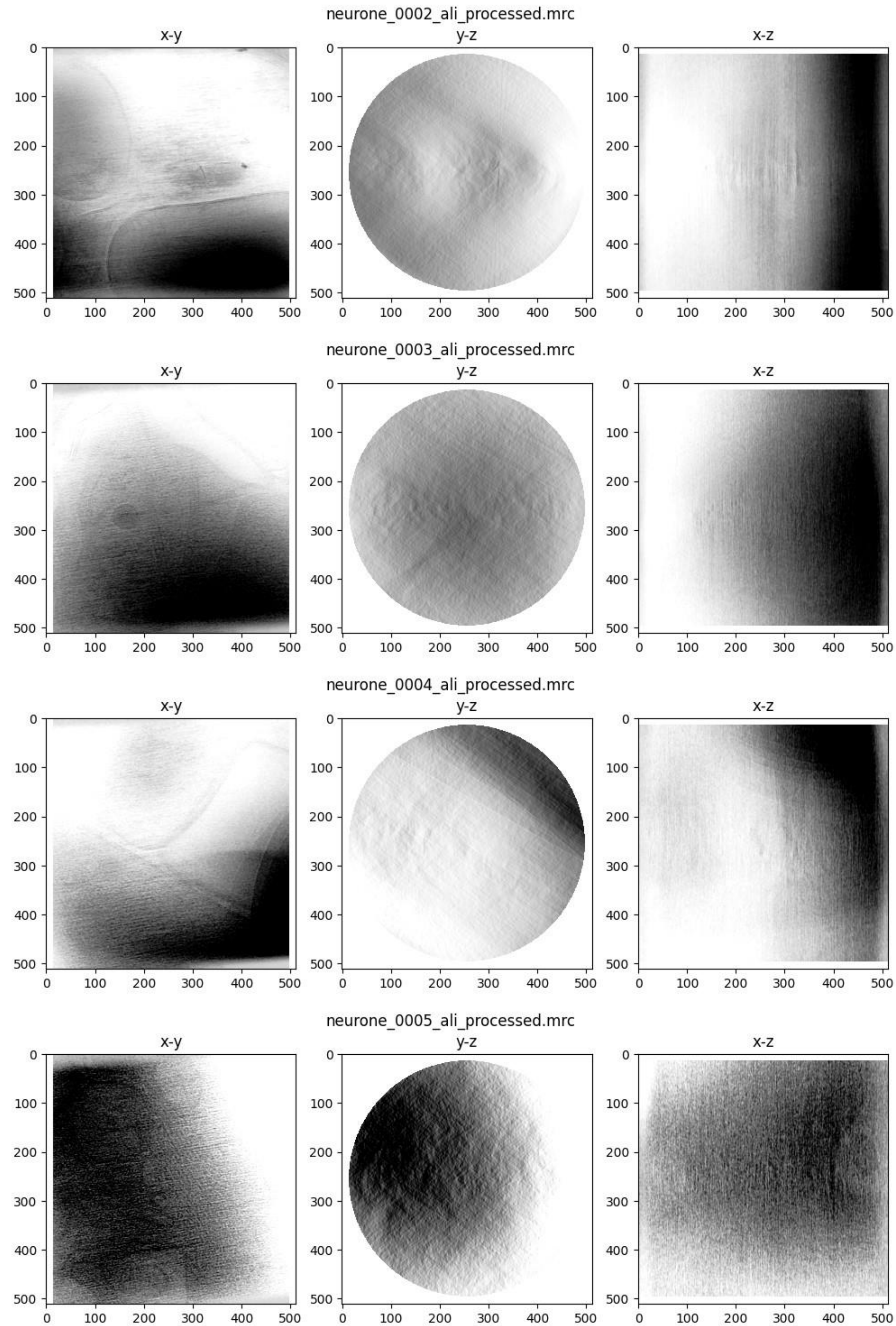

neurone_0002_ali_processed.mrc
x-y
y-z
x-z
neurone_0003_ali_processed.mrc
x-y
y-z
x-z
neurone_0004_ali_processed.mrc
x-y
y-z
x-z
neurone_0005_ali_processed.mrc
x-y
y-z
x-z
0
100
200
300
400
500

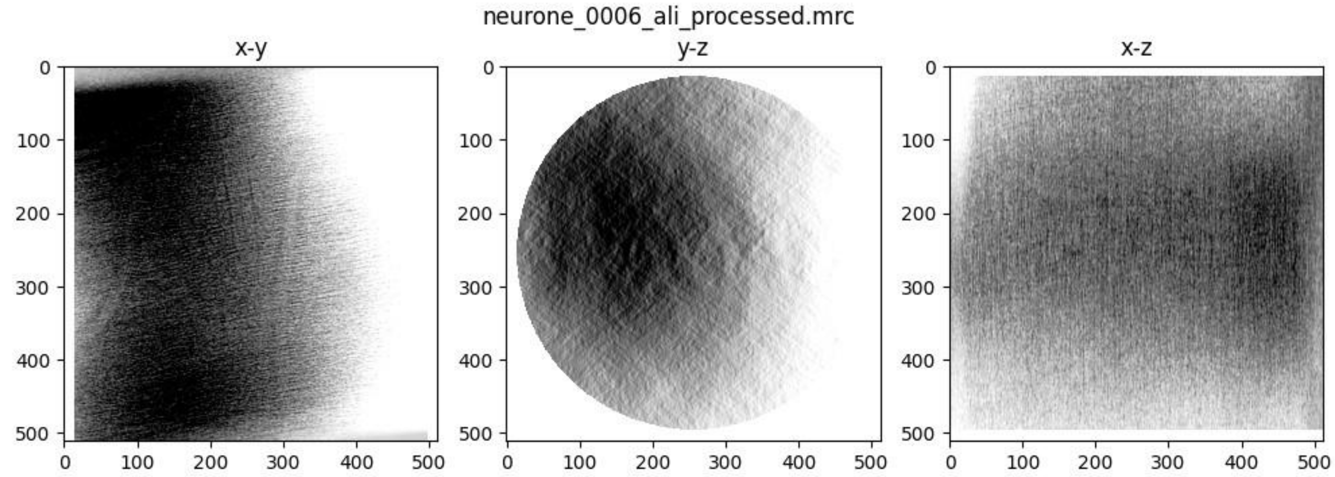
neurone_0006_ali_processed.mrc
x-y
y-z
x-z
0
100
200
300
400
500

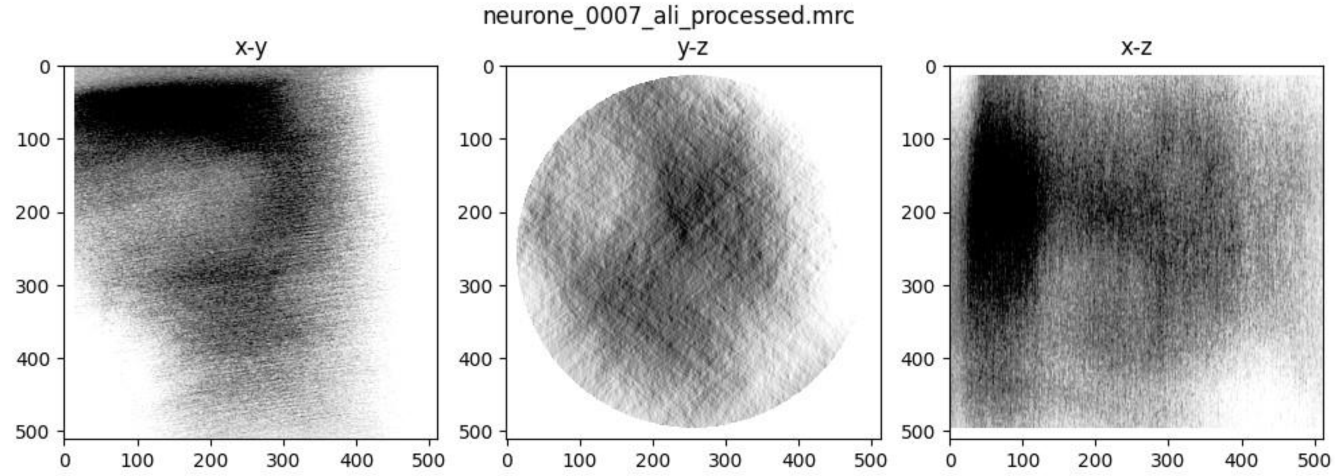
neurone_0007_ali_processed.mrc
x-y
y-z
x-z
0
100
200
300
400
500

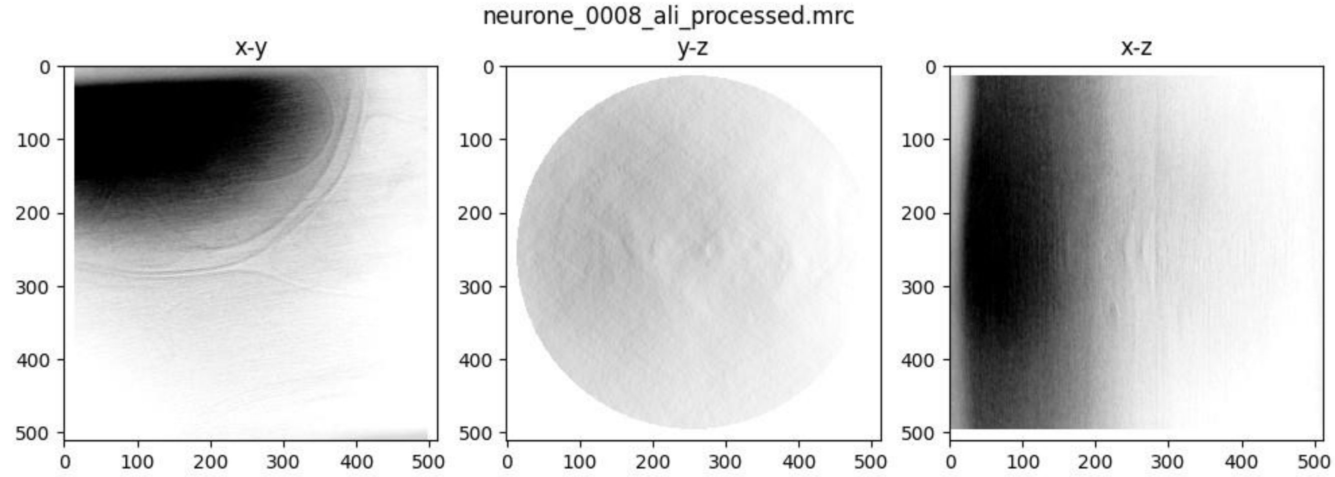
neurone_0008_ali_processed.mrc
x-y
y-z
x-z
0
100
200
300
400
500

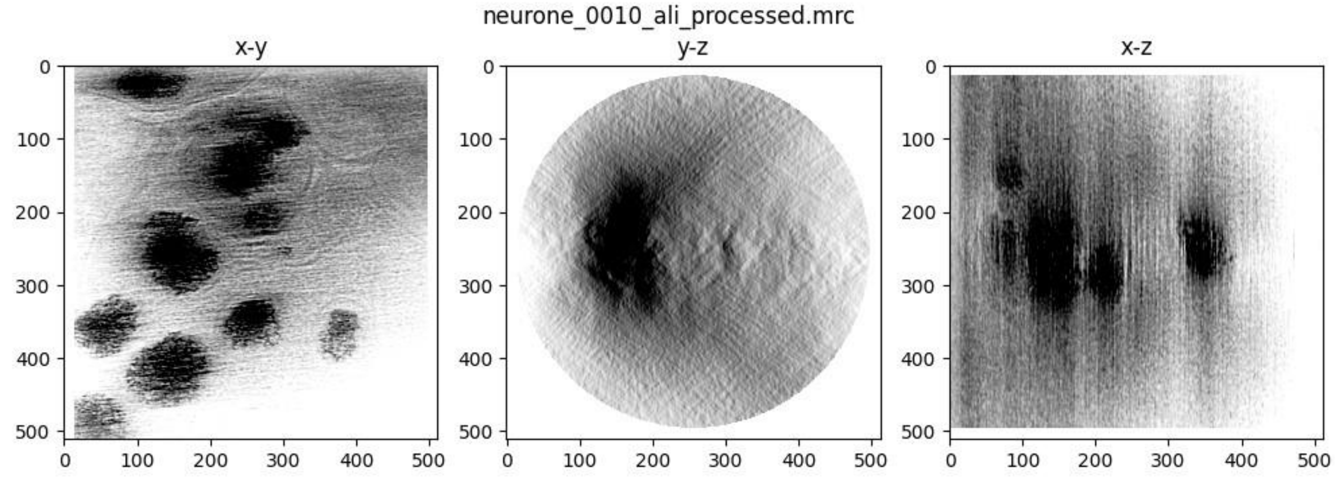
neurone_0010_ali_processed.mrc
x-y
y-z
x-z
0
100
200
300
400
500

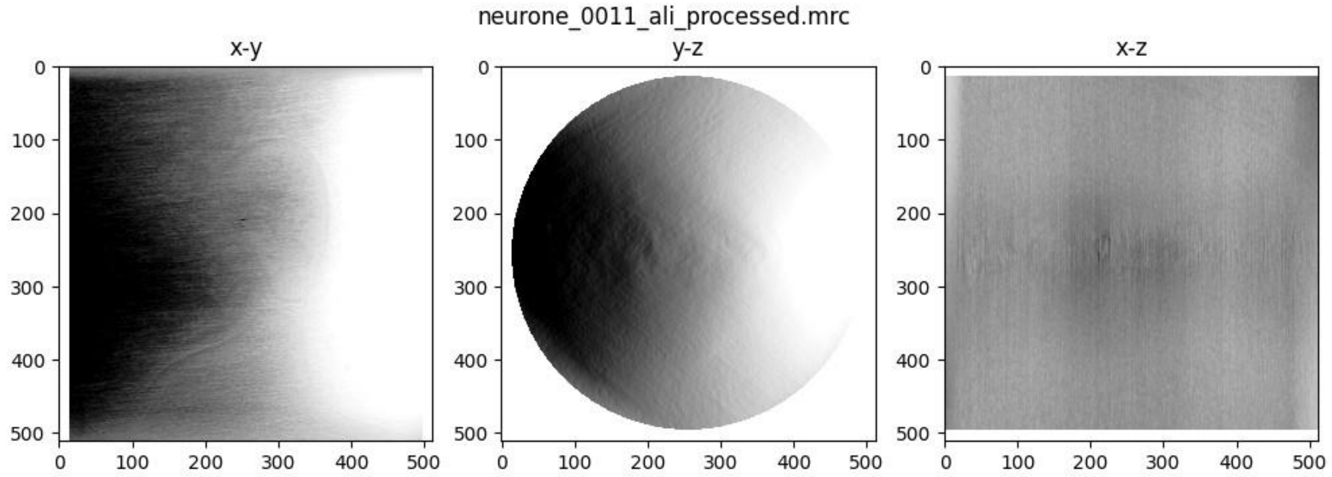
neurone_0011_ali_processed.mrc
x-y
y-z
x-z

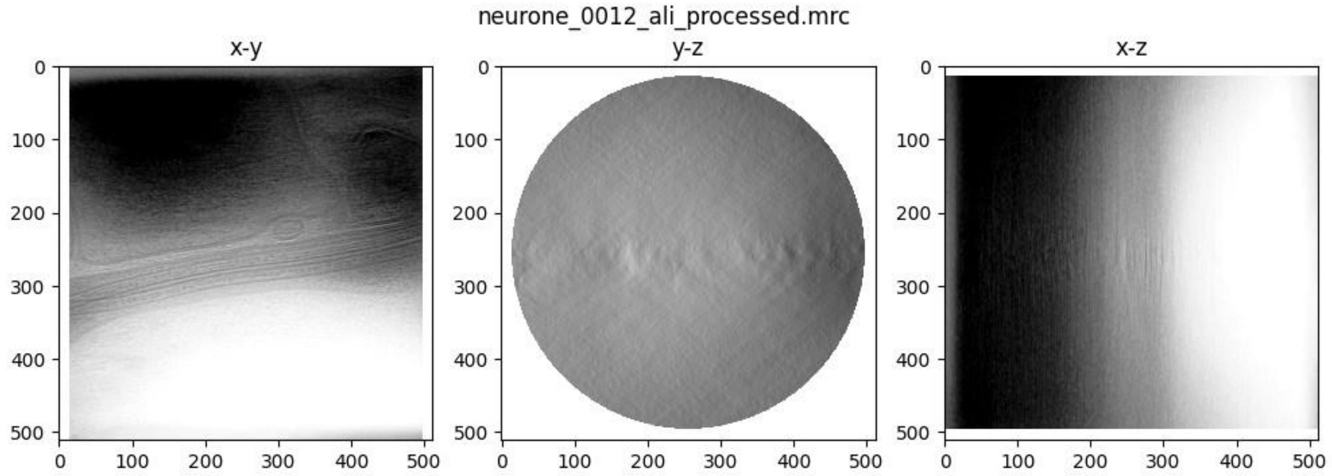
neurone_0012_ali_processed.mrc
x-y
y-z
x-z

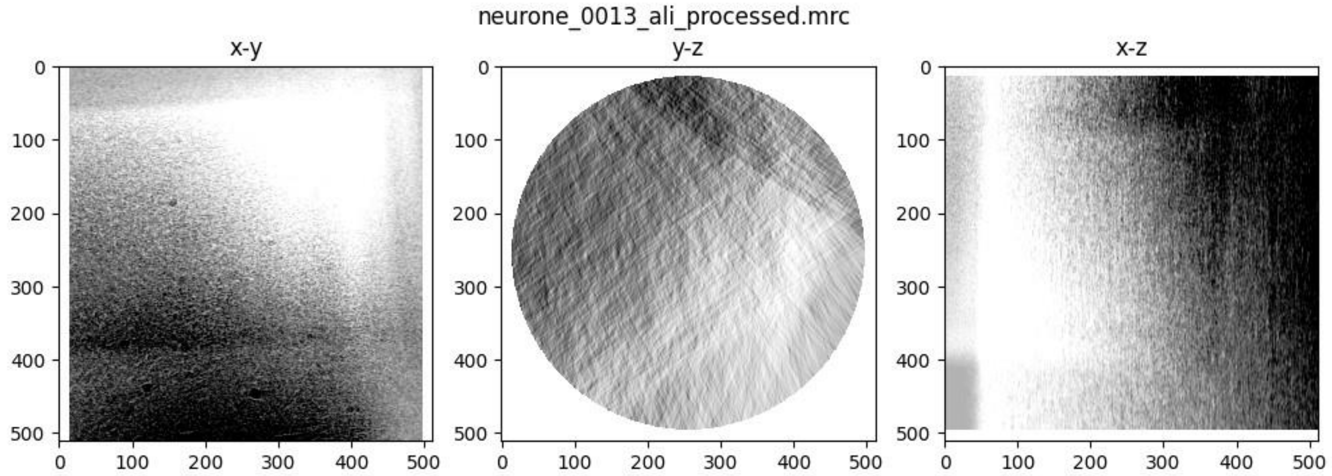
neurone_0013_ali_processed.mrc
x-y
y-z
x-z

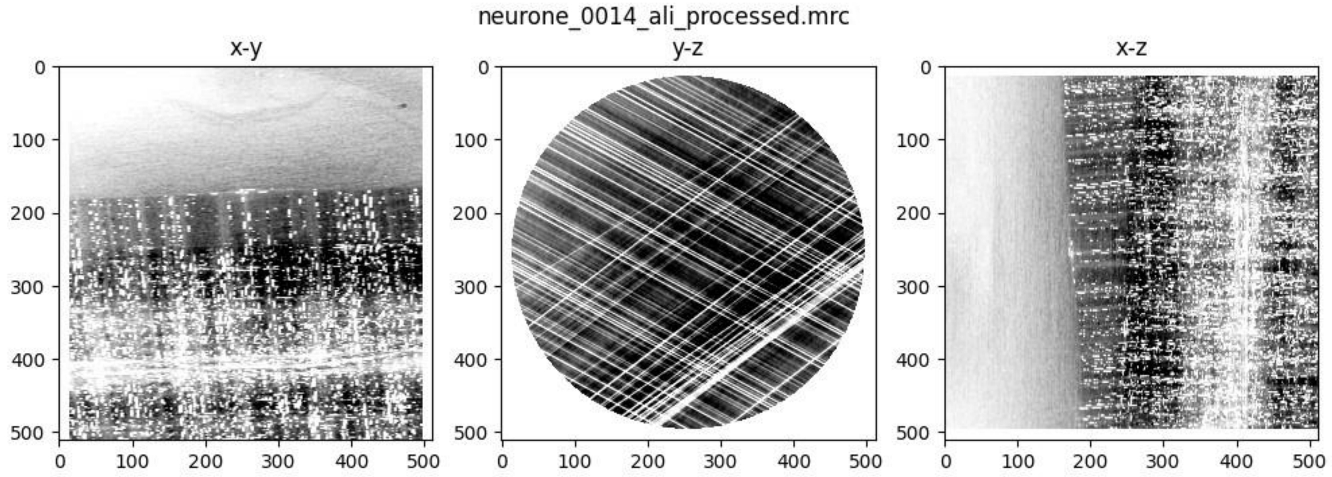
neurone_0014_ali_processed.mrc
x-y
y-z
x-z

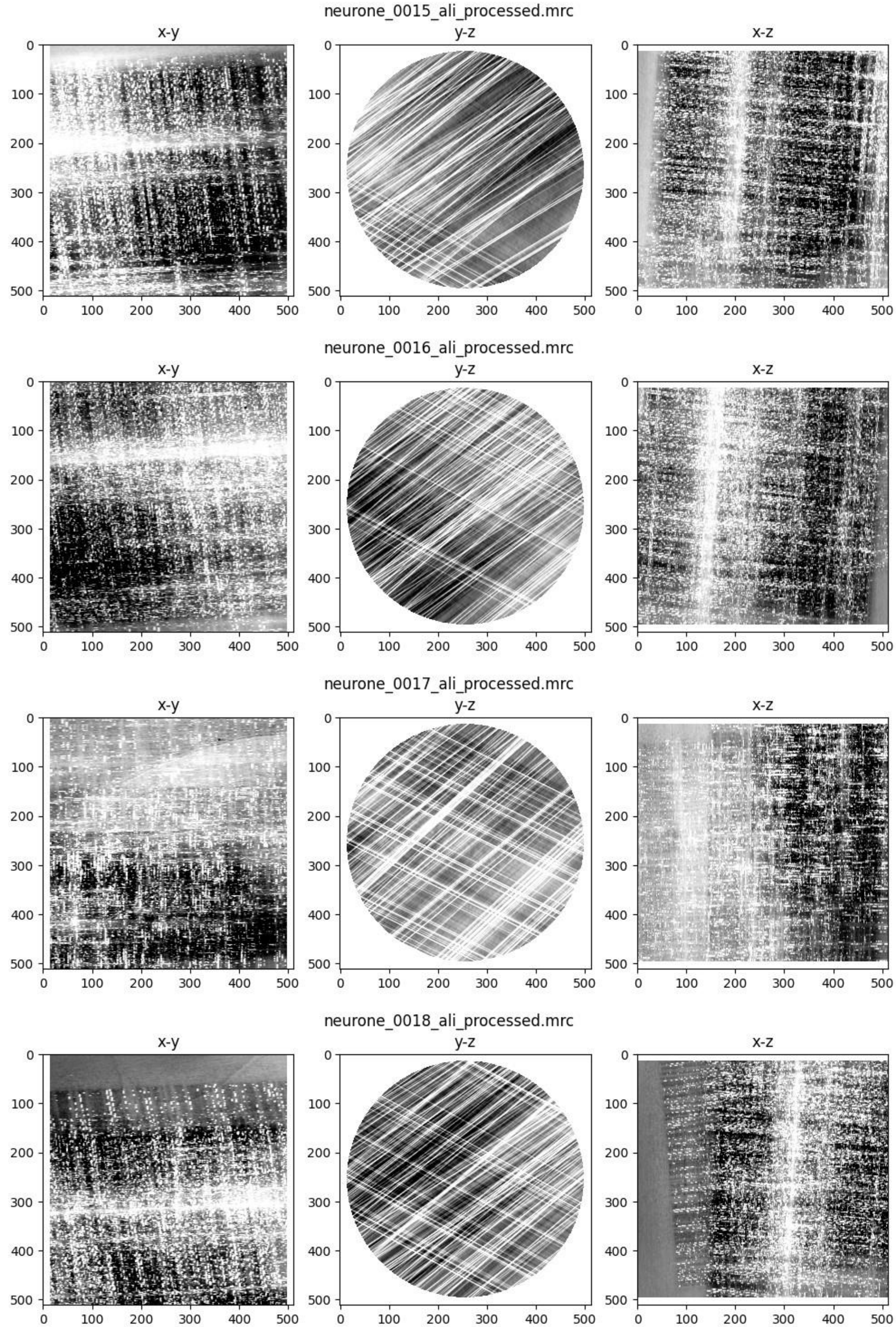
neurone_0015_ali_processed.mrc
x-y
y-z
x-z
neurone_0016_ali_processed.mrc
x-y
y-z
x-z
neurone_0017_ali_processed.mrc
x-y
y-z
x-z
neurone_0018_ali_processed.mrc
x-y
y-z
x-z

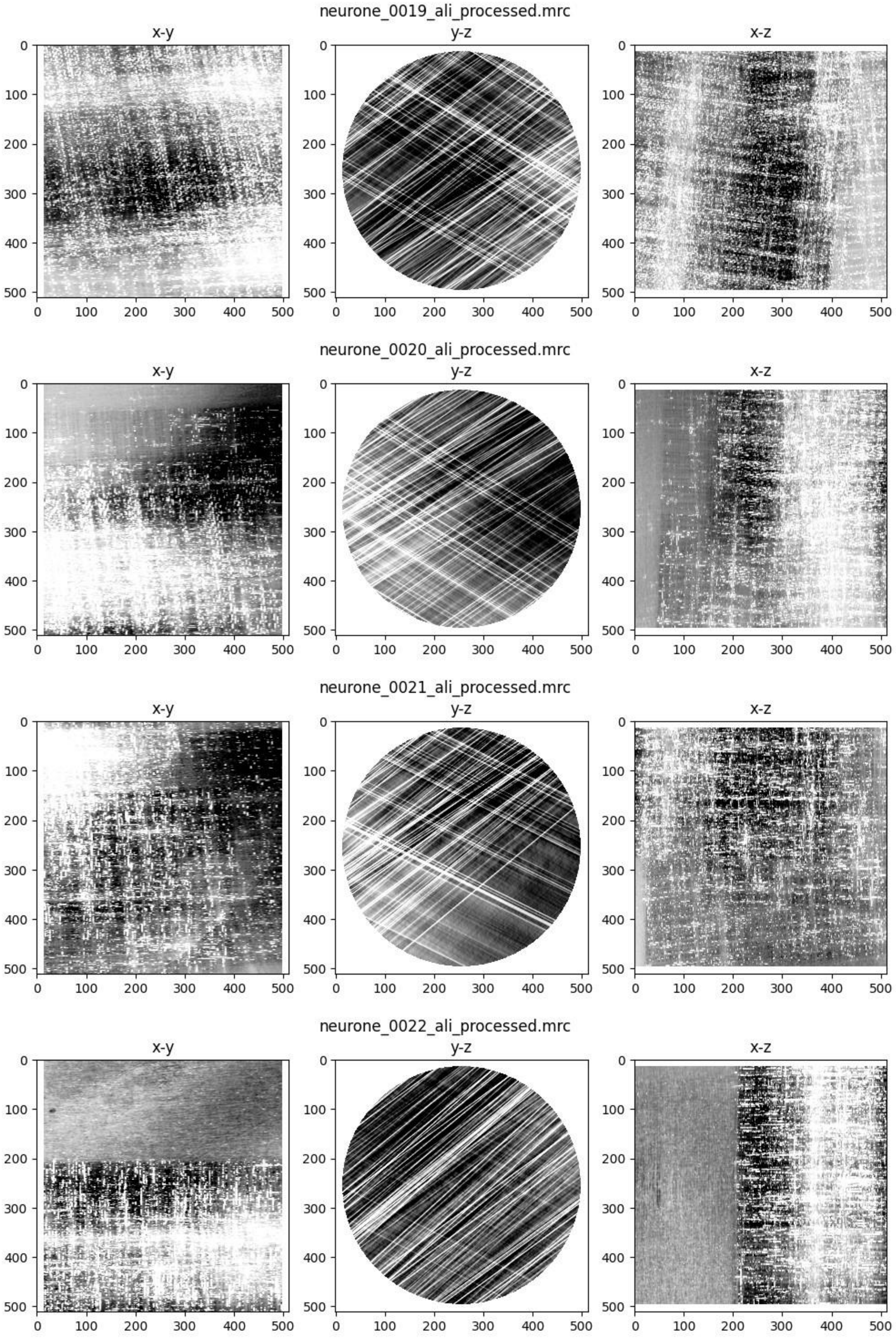
neurone_0019_ali_processed.mrc
x-y
y-z
x-z
neurone_0020_ali_processed.mrc
x-y
y-z
x-z
neurone_0021_ali_processed.mrc
x-y
y-z
x-z
neurone_0022_ali_processed.mrc
x-y
y-z
x-z

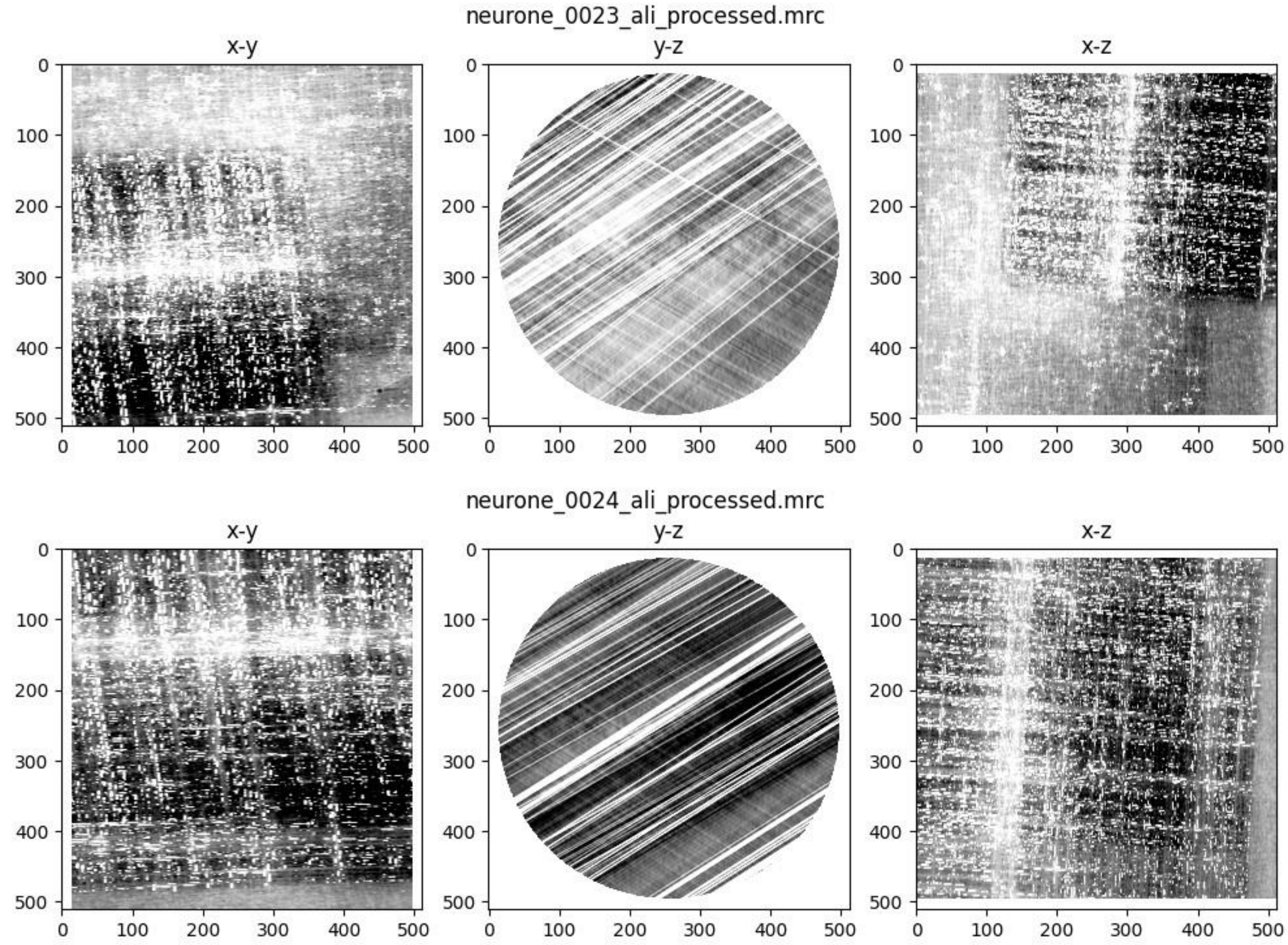
neurone_0023_ali_processed.mrc
x-y
y-z
x-z
neurone_0024_ali_processed.mrc
x-y
y-z
x-z

### Workflow diagram

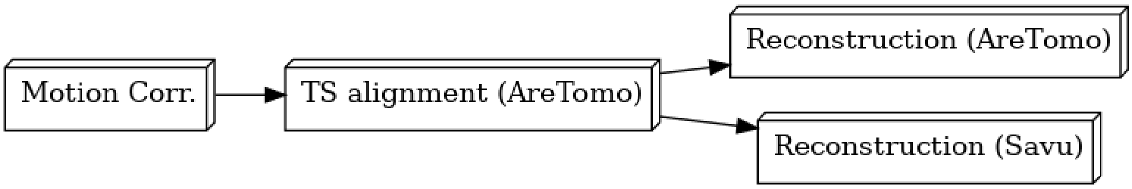


### Motion correction

Motion correction has been performed using MotionCor2.

Please cite:

1. Zheng, S., Palovcak, E., Armache, JP. et al. MotionCor2: anisotropic correction of beam-induced motion for improved cryo-electron microscopy. *Nat Methods* **14**, 331–332 (2017).

#### Summary of shifts for each tilt series

| | mean | std | 25% | 50% | 75% |
|---|---|---|---|---|---|
| **TiltSeries** | | | | | |
| **1** | 2.056202 | 2.656386 | 0.635059 | 1.234585 | 2.386598 |
| **2** | 1.415733 | 1.614418 | 0.521536 | 1.001249 | 1.791368 |
| **3** | 1.295799 | 1.117591 | 0.512445 | 0.965660 | 1.759602 |
| **4** | 1.661103 | 2.091689 | 0.577062 | 1.106029 | 1.981691 |
| **5** | 1.298596 | 1.033565 | 0.554437 | 1.041009 | 1.762385 |
| **6** | 1.379971 | 1.221563 | 0.605310 | 1.103087 | 1.829891 |
| **7** | 2.242506 | 2.905982 | 0.682642 | 1.346291 | 2.616104 |
| **8** | 1.402211 | 1.245235 | 0.562228 | 1.070000 | 1.881303 |
| **10** | 1.209009 | 0.884546 | 0.586941 | 1.011929 | 1.598405 |
| **11** | 1.871856 | 2.428340 | 0.586941 | 1.202477 | 2.268127 |

| | | | | | |
|---|---|---|---|---|---|
| **12** | 1.623454 | 2.318478 | 0.503289 | 0.973293 | 1.905197 |
| **13** | 6.831695 | 6.379735 | 2.314584 | 5.125159 | 9.487324 |
| **14** | 16.070258 | 49.565171 | 0.590339 | 1.156849 | 2.527969 |
| **15** | 19.491723 | 55.706889 | 0.605392 | 1.180042 | 2.618416 |
| **16** | 24.059852 | 65.467685 | 0.665733 | 1.379021 | 3.073308 |
| **17** | 20.347179 | 59.296513 | 0.617414 | 1.267793 | 2.946994 |
| **18** | 19.982279 | 56.926885 | 0.621772 | 1.191512 | 2.670431 |
| **19** | 23.301804 | 59.528379 | 1.168428 | 2.810970 | 9.441590 |
| **20** | 22.763273 | 73.947772 | 0.609590 | 1.279453 | 3.041772 |
| **21** | 15.023953 | 48.436937 | 0.653911 | 1.341827 | 2.771299 |
| **22** | 12.322676 | 45.376714 | 0.684251 | 1.385135 | 2.668122 |
| **23** | 10.351220 | 35.034275 | 0.687677 | 1.371860 | 2.622532 |
| **24** | 10.273665 | 39.577862 | 0.594643 | 1.135319 | 2.231569 |

```
Out[12]: Text(0.5, 1.0, 'Euclidean Shift Means +/- Standard Deviation')
```

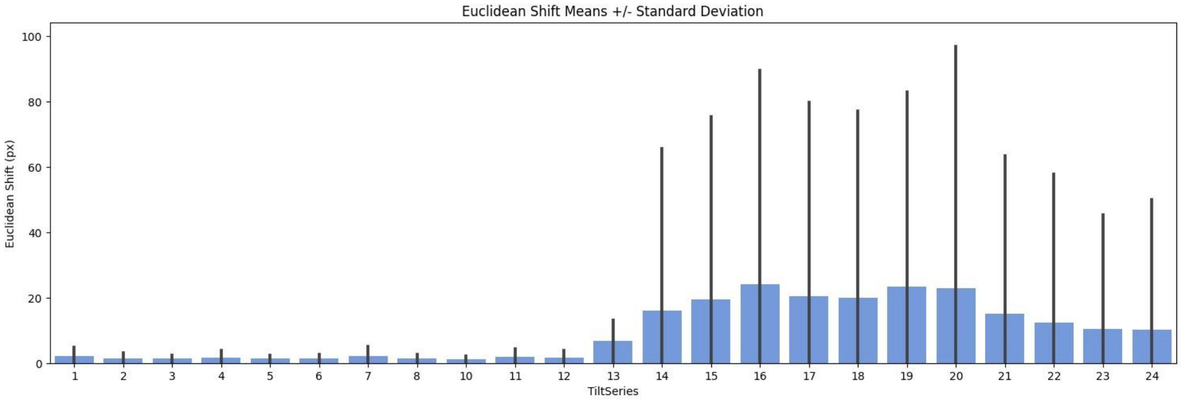

### AreTomo

AreTomo was used in this workflow. Please cite:

1. Shawn Zheng, Georg Wolff, Garrett Greenan, et al. AreTomo: An integrated software package for automated marker-free, motion-corrected cryo-electron tomographic alignment and reconstruction, *JSB: X* **6(100068)** , ISSN 2590-1524, (2022) DOI | ScienceDirect

For more information, please see https://msg.ucsf.edu/software.

#### AreTomo Alignment

Out[15]:

| | Tilt series | Mean shift (px) | Shift s.d. (px) | Mean shift (A) | Shift s.d. (A) |
|---|---|---|---|---|---|
| **1** | 1 | 249.846455 | 134.187956 | 293.819432 | 157.805036 |
| **2** | 2 | 167.342715 | 105.078591 | 196.795033 | 123.572423 |
| **3** | 3 | 156.732914 | 85.524085 | 184.317907 | 100.576324 |
| **4** | 4 | 257.709137 | 154.558138 | 303.065945 | 181.760370 |
| **5** | 5 | 145.165522 | 106.232851 | 170.714654 | 124.929833 |
| **6** | 6 | 171.819790 | 121.486764 | 202.060072 | 142.868434 |
| **7** | 7 | 316.277157 | 234.309473 | 371.941936 | 275.547940 |
| **8** | 8 | 152.049511 | 100.580919 | 178.810225 | 118.283161 |
| **9** | 10 | 209.886436 | 145.100036 | 246.826449 | 170.637642 |
| **10** | 11 | 191.660644 | 141.893540 | 225.392917 | 166.866803 |
| **11** | 12 | 203.012721 | 102.486559 | 238.742960 | 120.524194 |
| **12** | 13 | 409.359774 | 357.633795 | 481.407095 | 420.577343 |
| **13** | 14 | 169.928989 | 126.169199 | 199.836491 | 148.374978 |
| **14** | 15 | 365.881024 | 171.289044 | 430.276084 | 201.435915 |
| **15** | 16 | 438.967867 | 237.565736 | 516.226212 | 279.377306 |

| | | | | | |
|---|---|---|---|---|---|
| **16** | 17 | 300.202655 | 219.939474 | 353.038322 | 258.648822 |
| **17** | 18 | 162.764817 | 127.174397 | 191.411424 | 149.557091 |
| **18** | 19 | 248.243619 | 341.288754 | 291.934496 | 401.355575 |
| **19** | 20 | 267.172073 | 131.402679 | 314.194358 | 154.529551 |
| **20** | 21 | 172.336549 | 138.699108 | 202.667781 | 163.110151 |
| **21** | 22 | 242.213529 | 178.690878 | 284.843110 | 210.140472 |
| **22** | 23 | 330.917261 | 281.295930 | 389.158699 | 330.804013 |
| **23** | 24 | 823.669692 | 476.013402 | 968.635558 | 559.791760 |

```
Out[16]: (0.0, 2000.0)
```

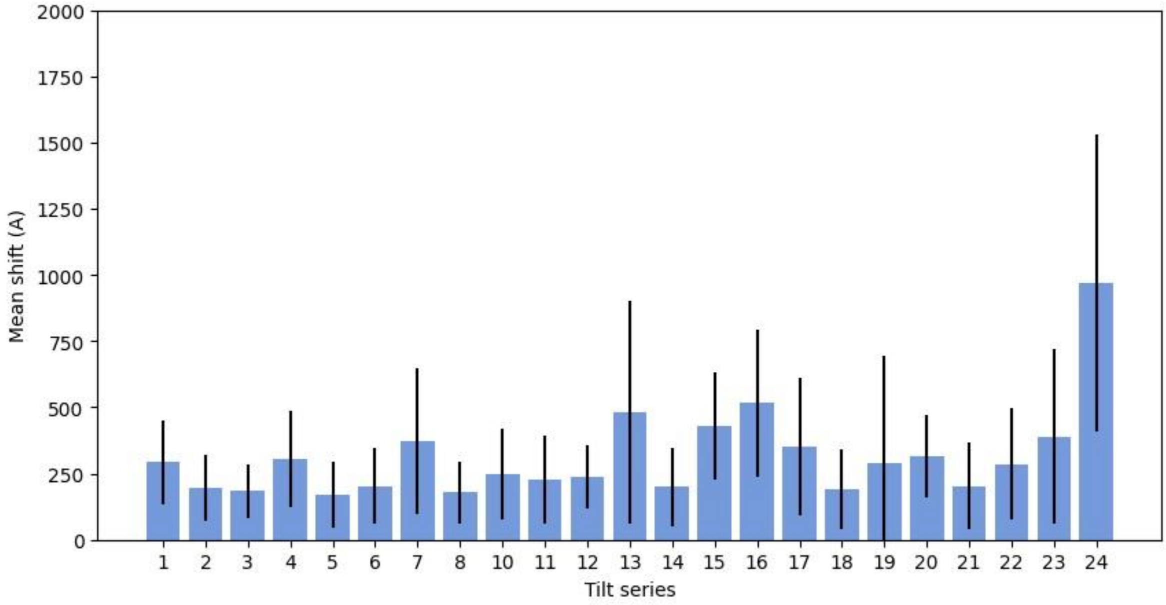

### Tomographic Reconstruction (AreTomo)

Vol Z = 1000

Sample thickness = -1

Pixel size = 1.47

Reconstruction algorithm = SART

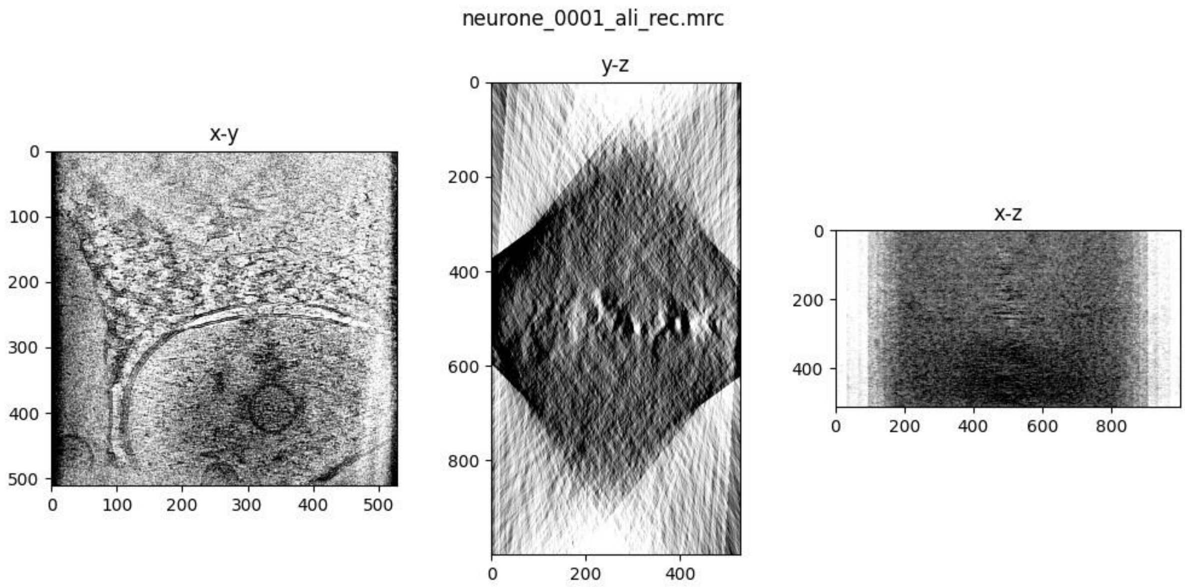

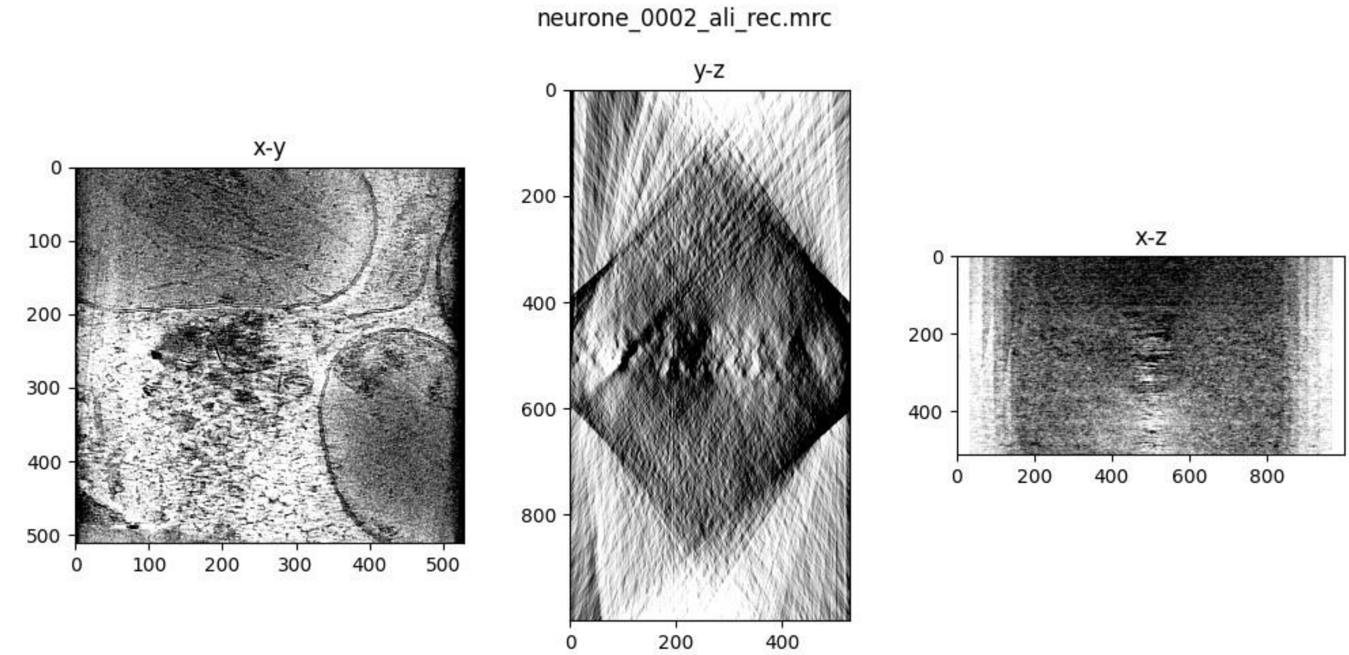
neurone_0002_ali_rec.mrc
x-y
y-z
x-z

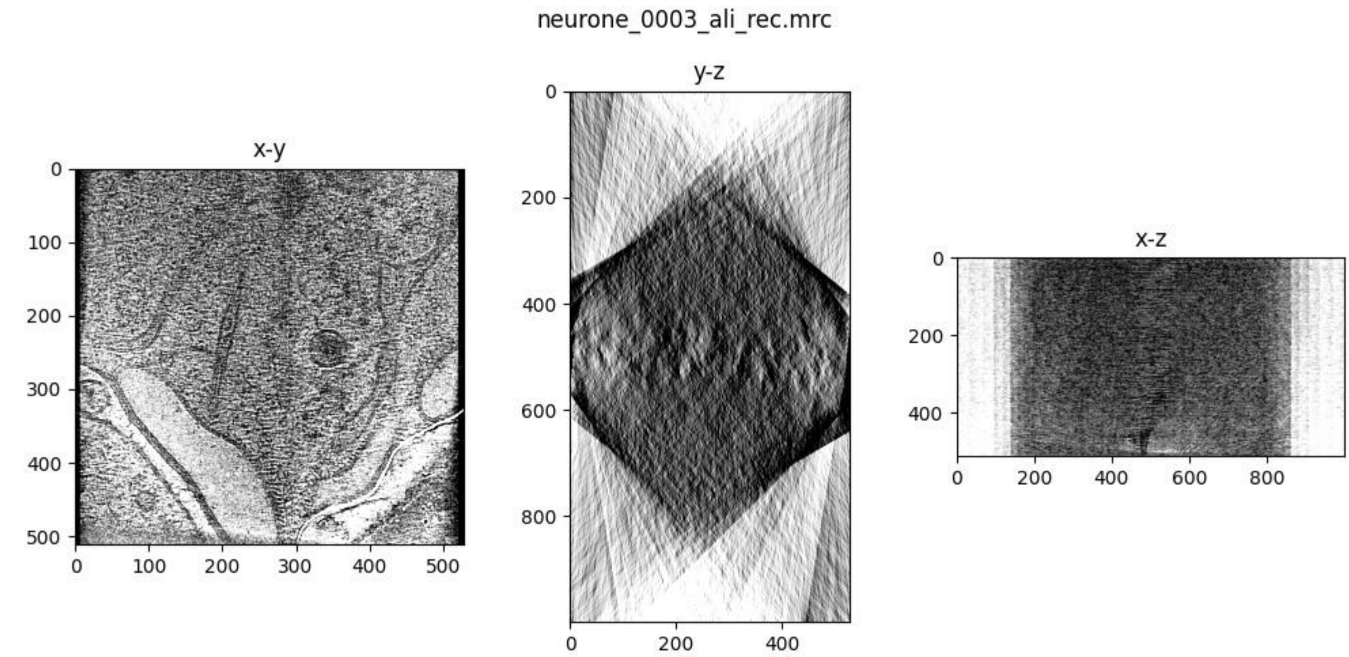
neurone_0003_ali_rec.mrc
x-y
y-z
x-z

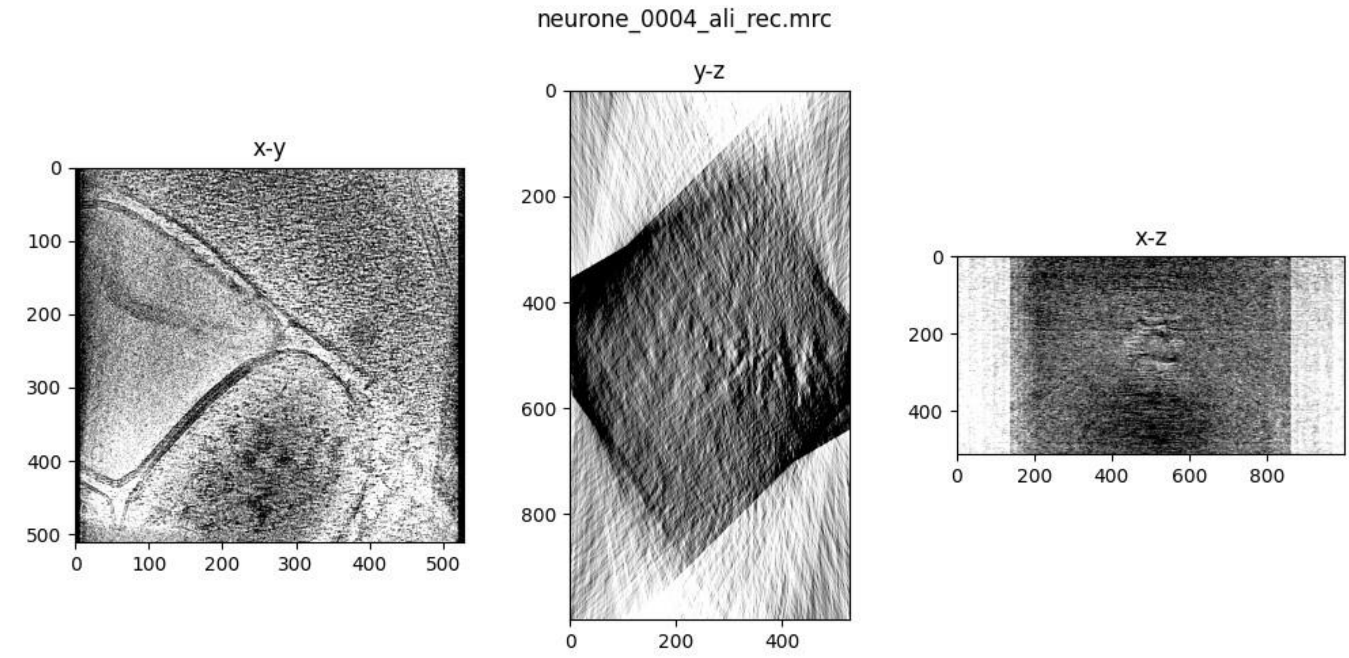
neurone_0004_ali_rec.mrc
x-y
y-z
x-z

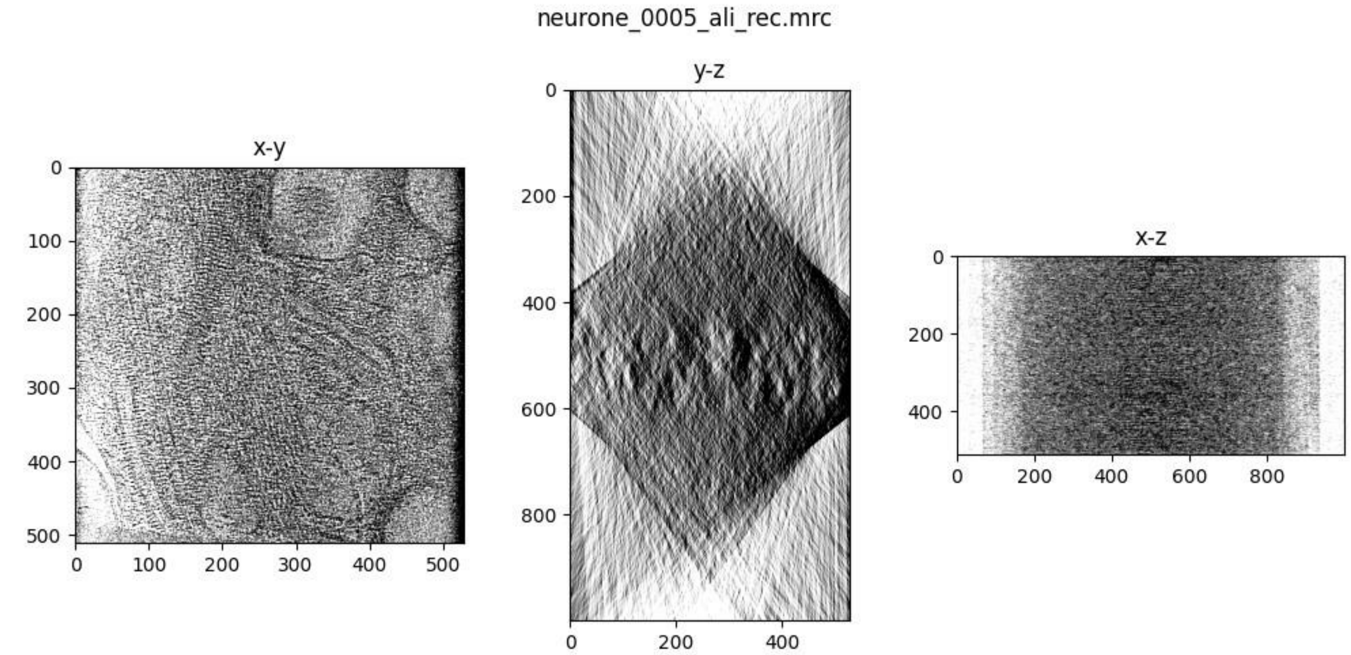
neurone_0005_ali_rec.mrc
x-y
y-z
x-z

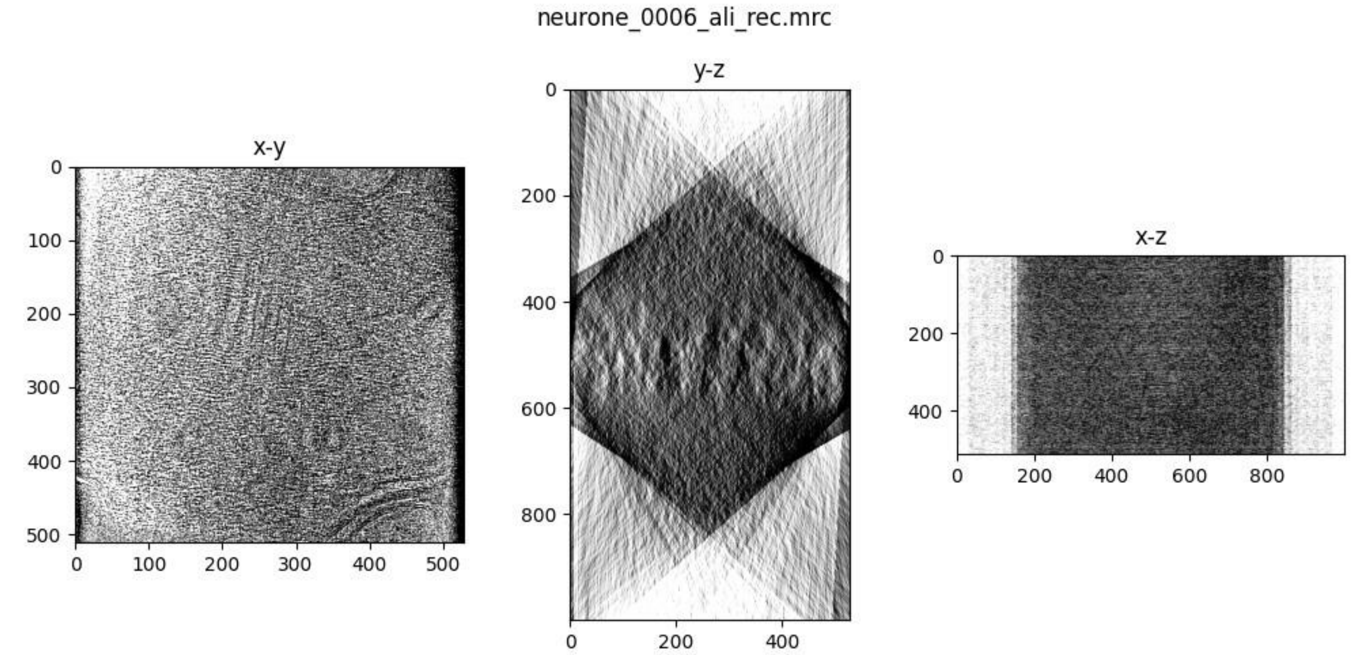
neurone_0006_ali_rec.mrc
x-y
y-z
x-z

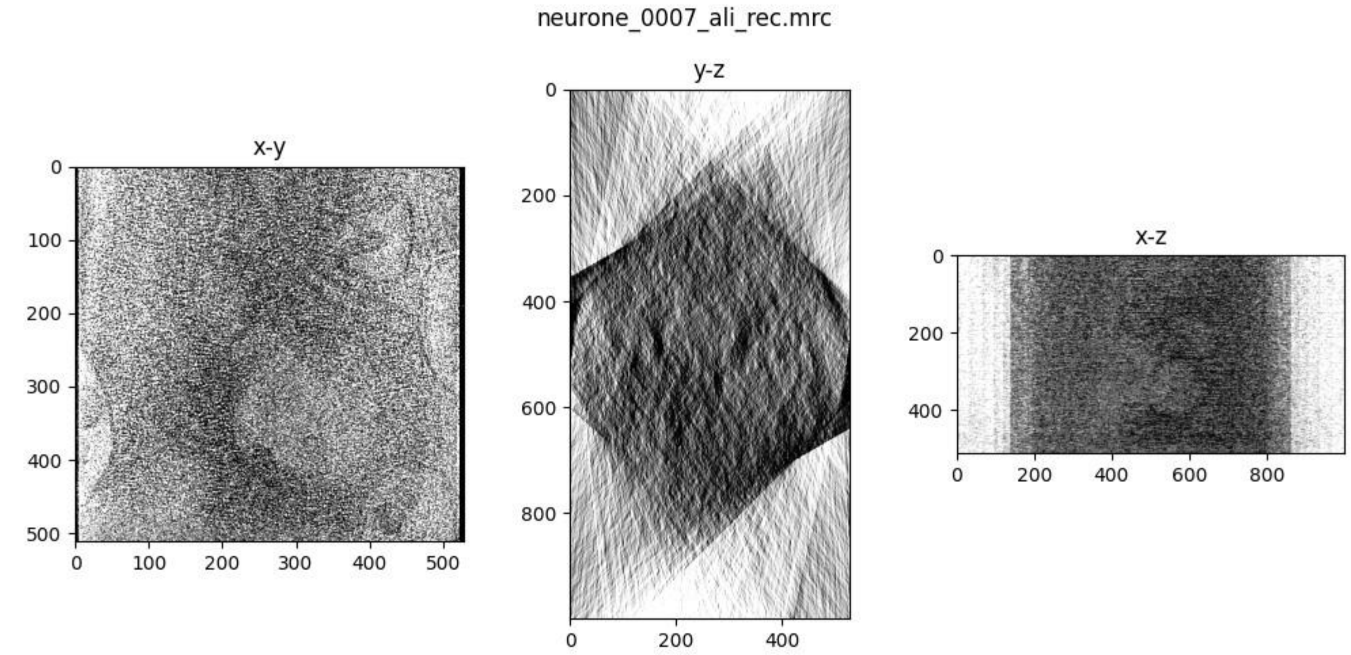
neurone_0007_ali_rec.mrc
x-y
y-z
x-z

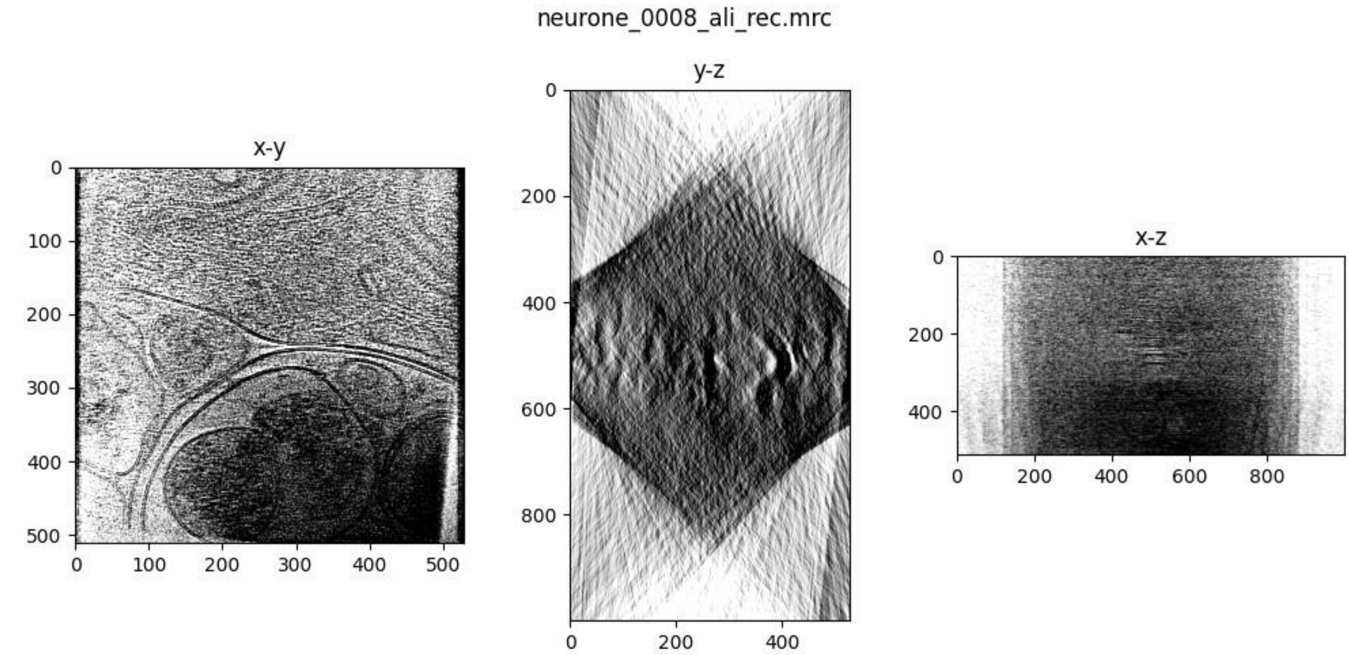
neurone_0008_ali_rec.mrc
x-y
y-z
x-z

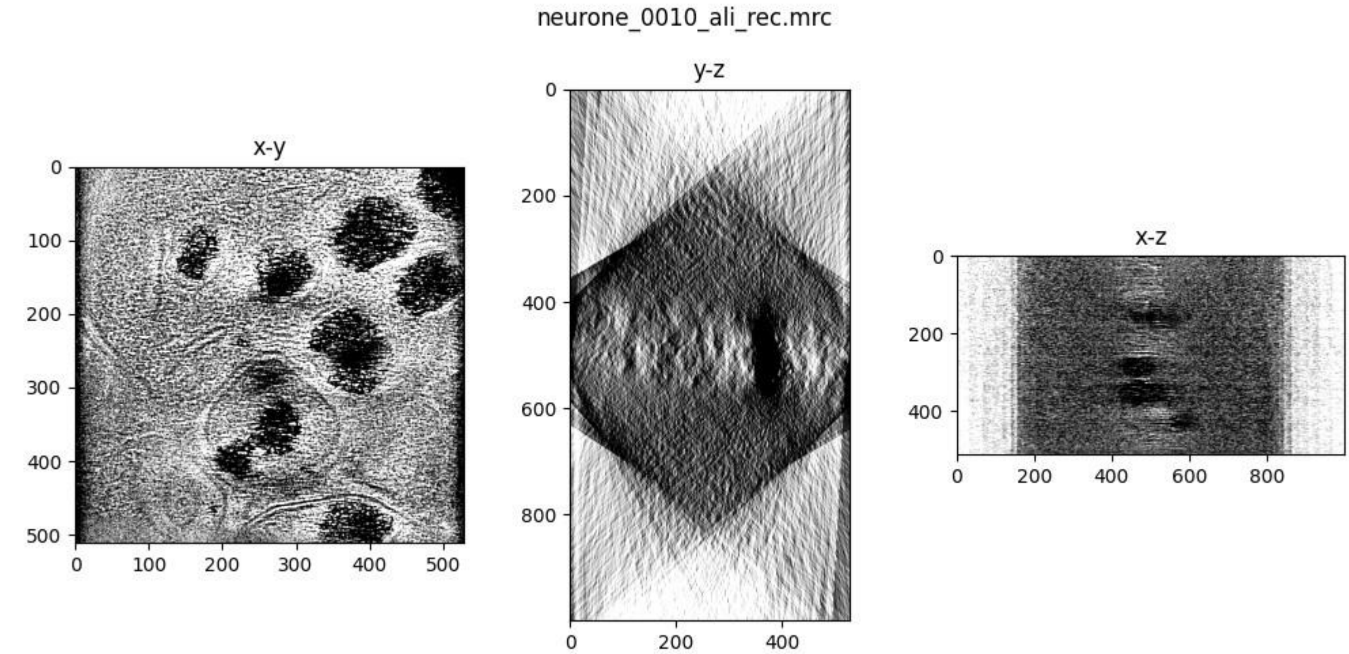
neurone_0010_ali_rec.mrc
x-y
y-z
x-z

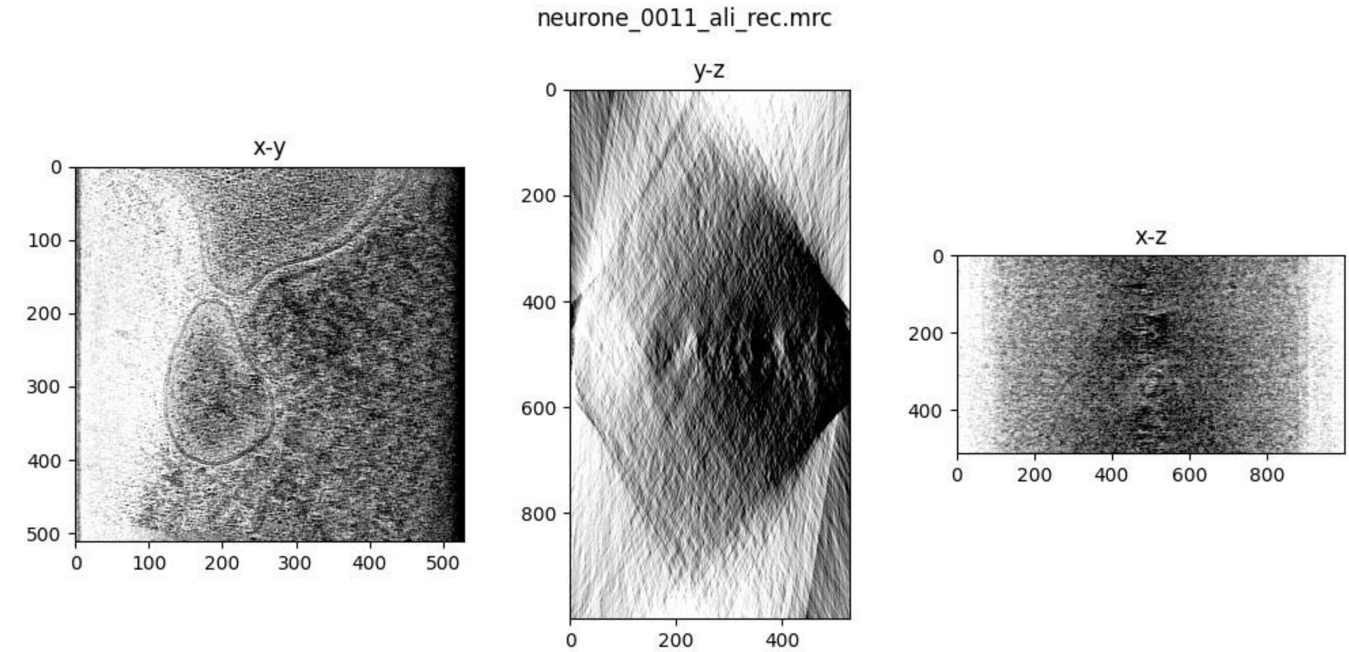
neurone_0011_ali_rec.mrc
x-y
y-z
x-z

neurone_0012_ali_rec.mrc

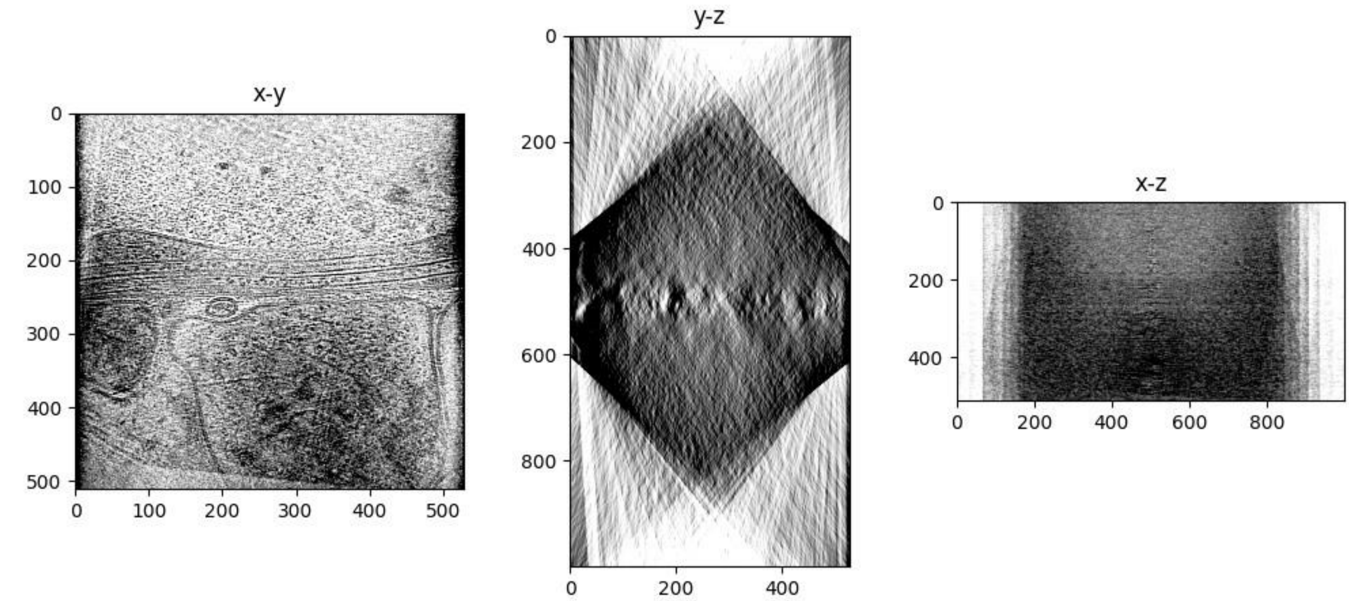


neurone_0013_ali_rec.mrc

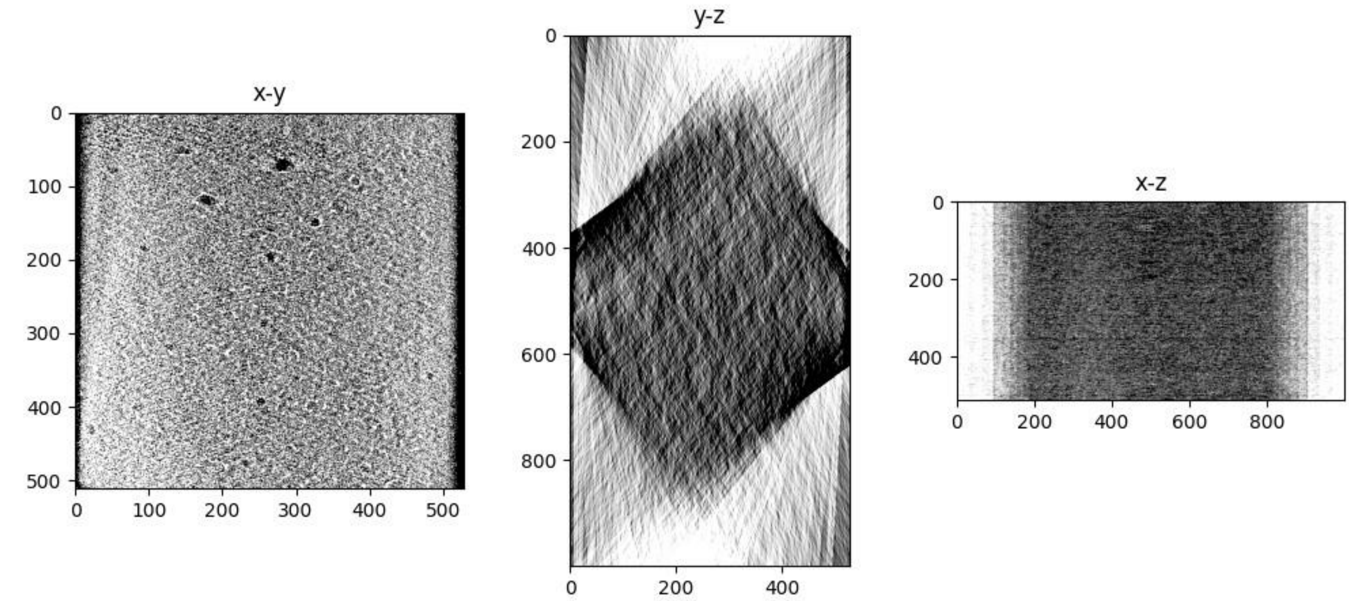


neurone_0014_ali_rec.mrc

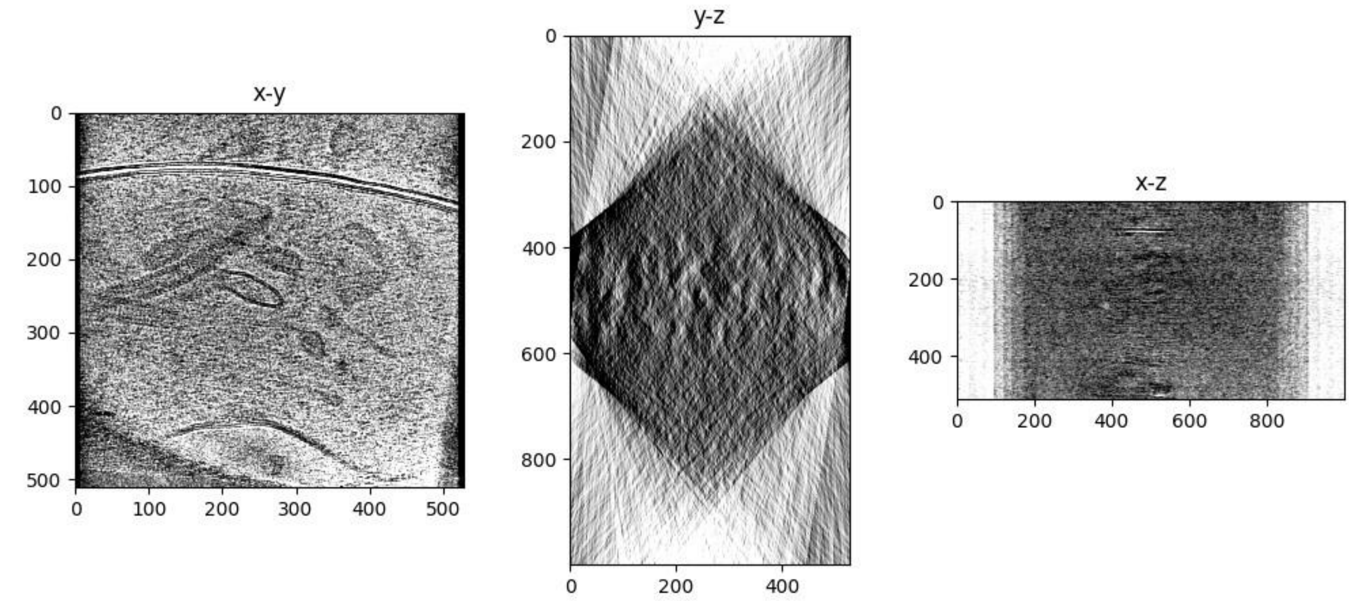

neurone_0015_ali_rec.mrc

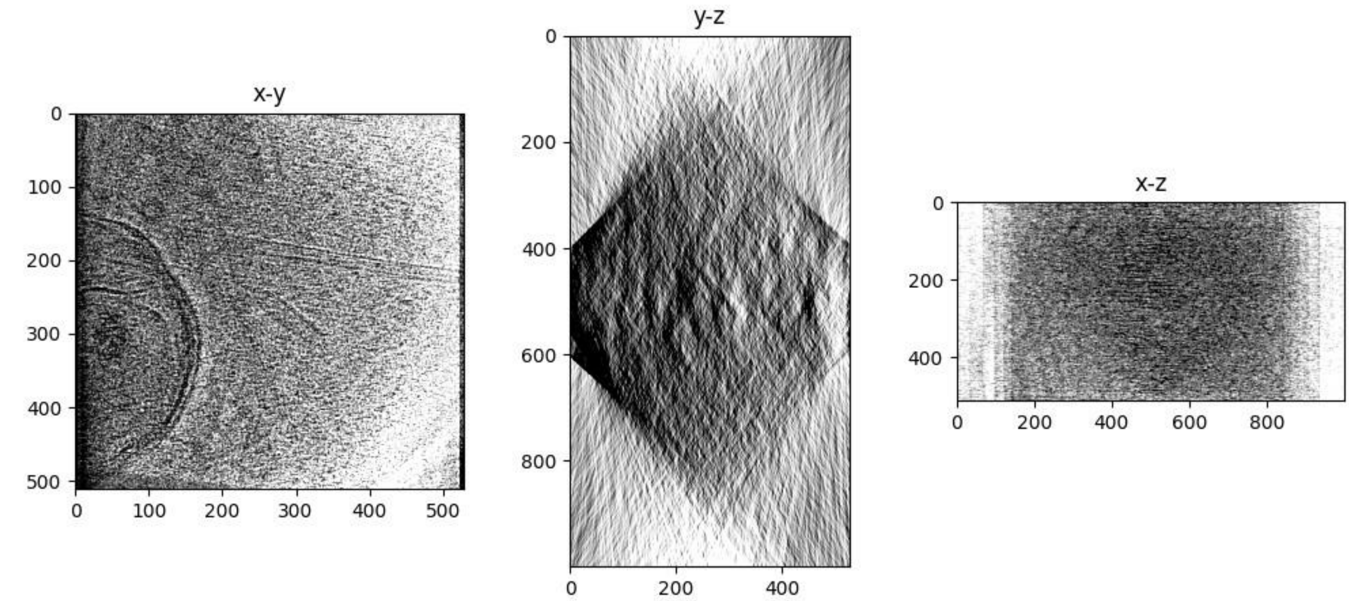


neurone_0016_ali_rec.mrc

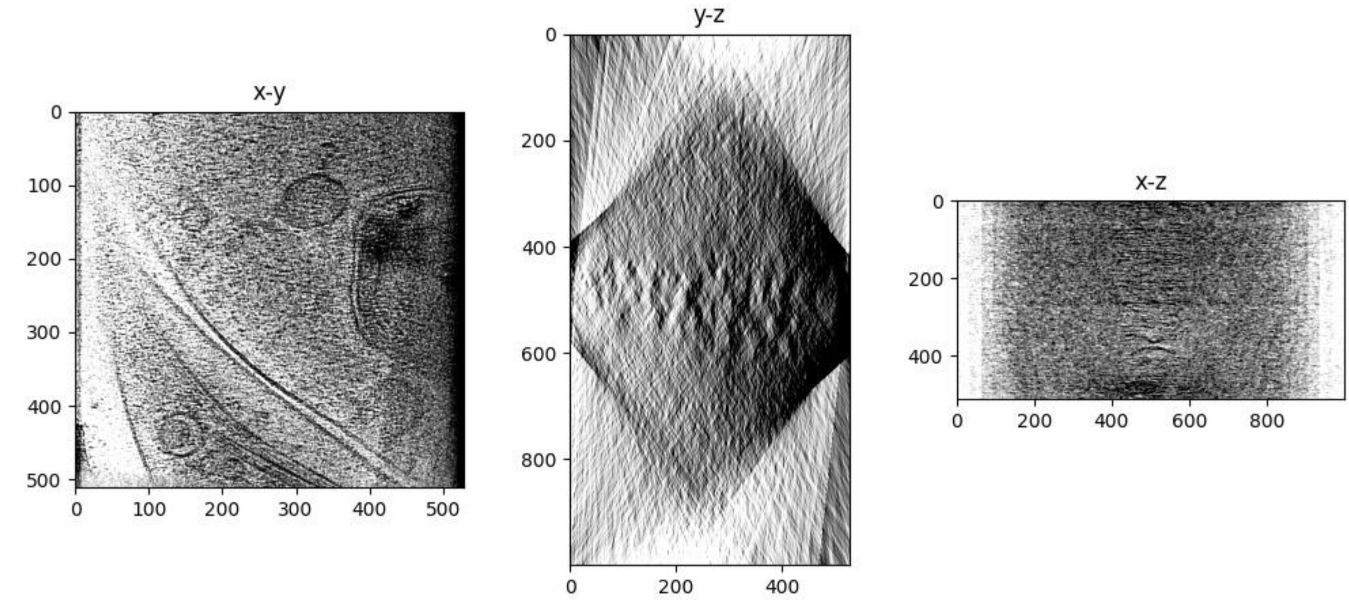


neurone_0017_ali_rec.mrc

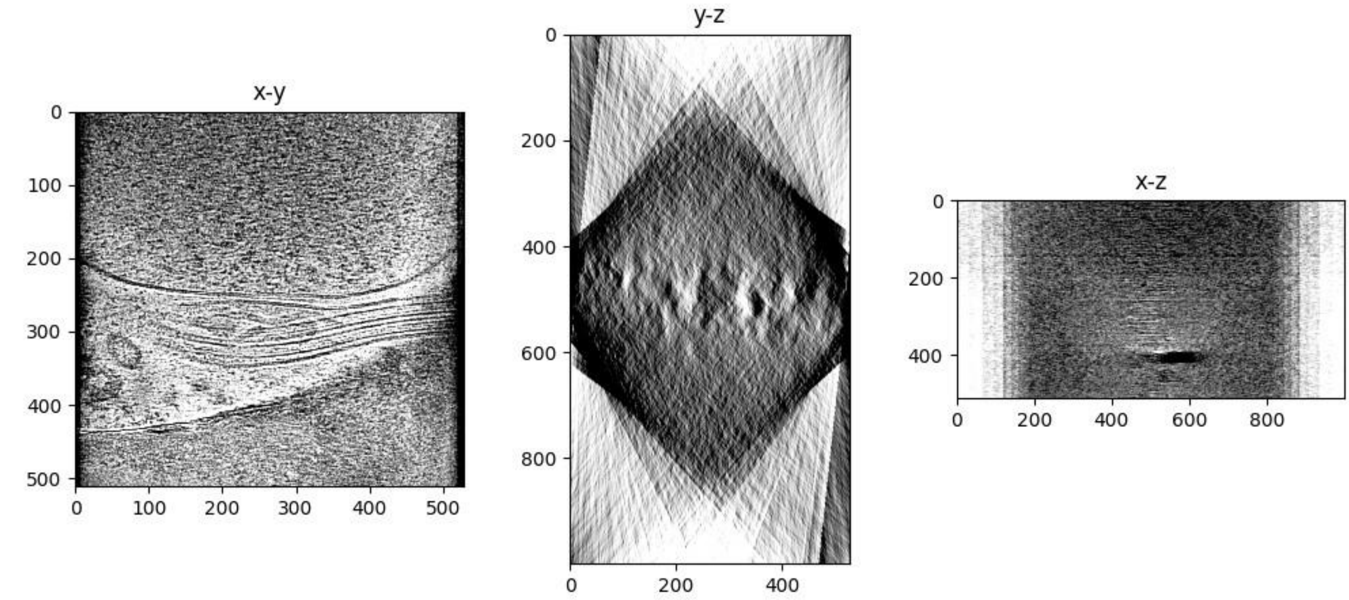

neurone_0018_ali_rec.mrc

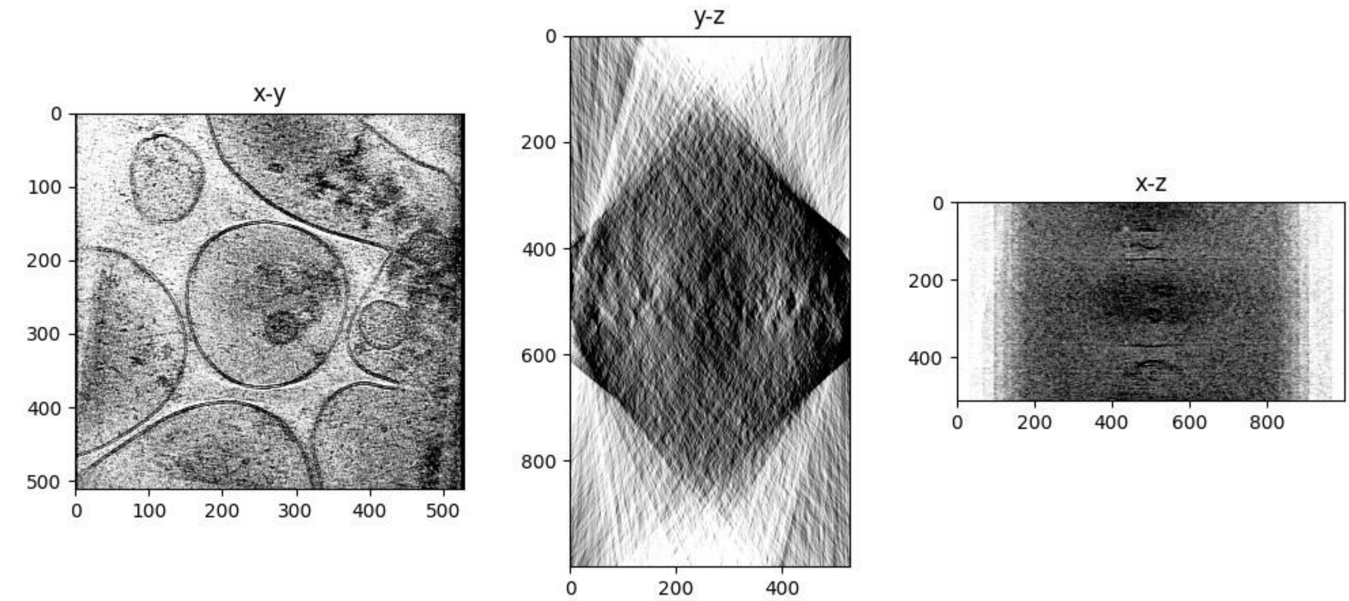


neurone_0019_ali_rec.mrc

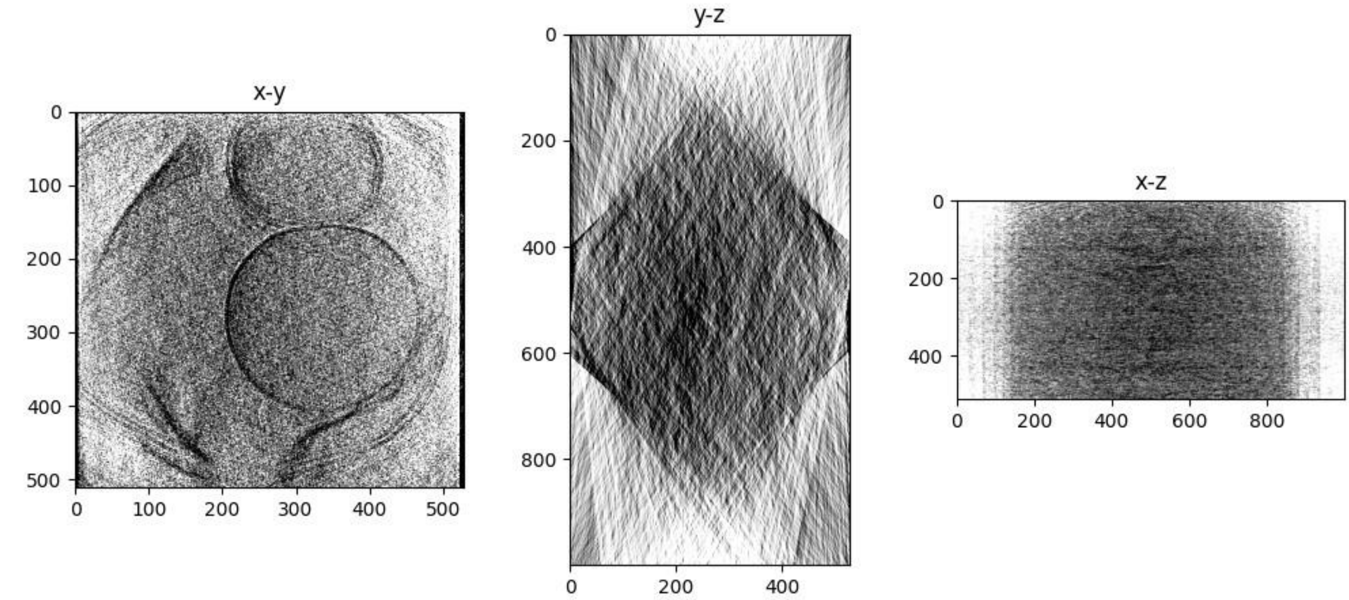


neurone_0020_ali_rec.mrc

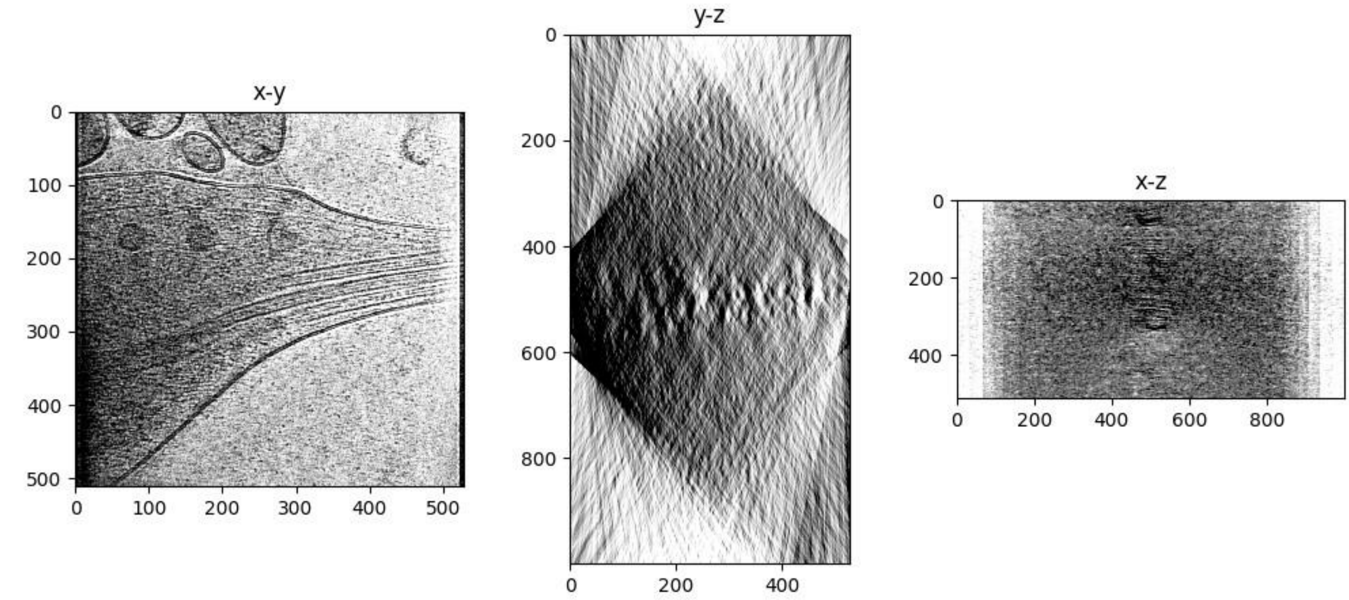

neurone_0021_ali_rec.mrc

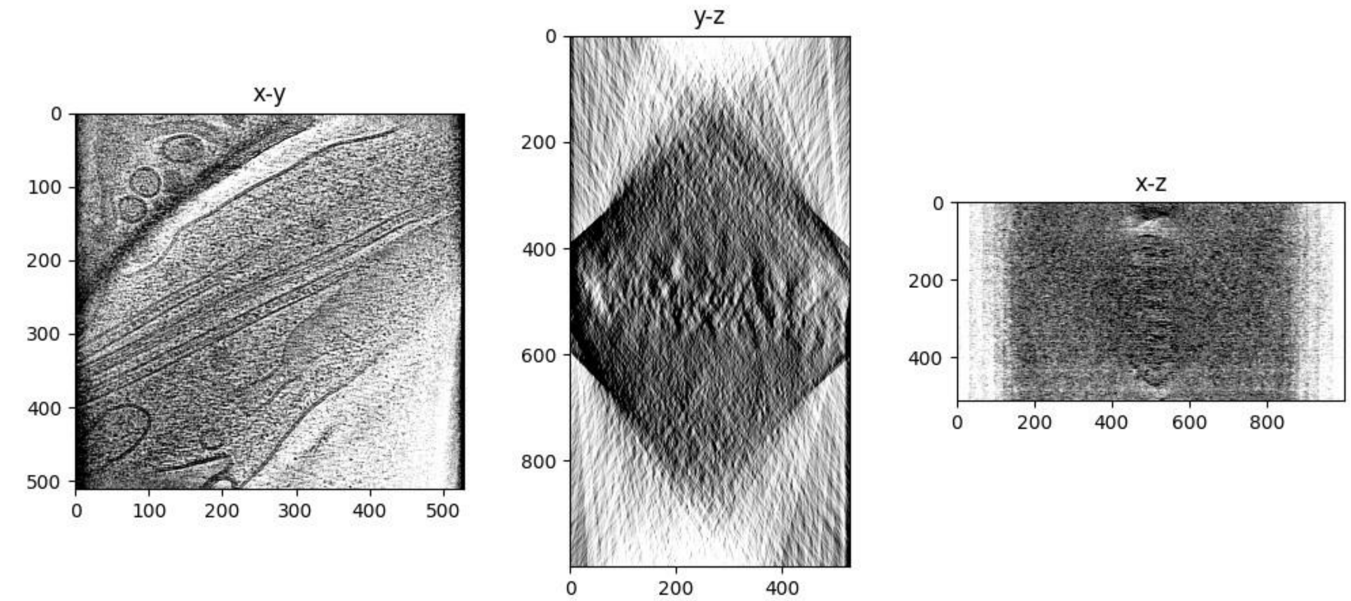


neurone_0022_ali_rec.mrc

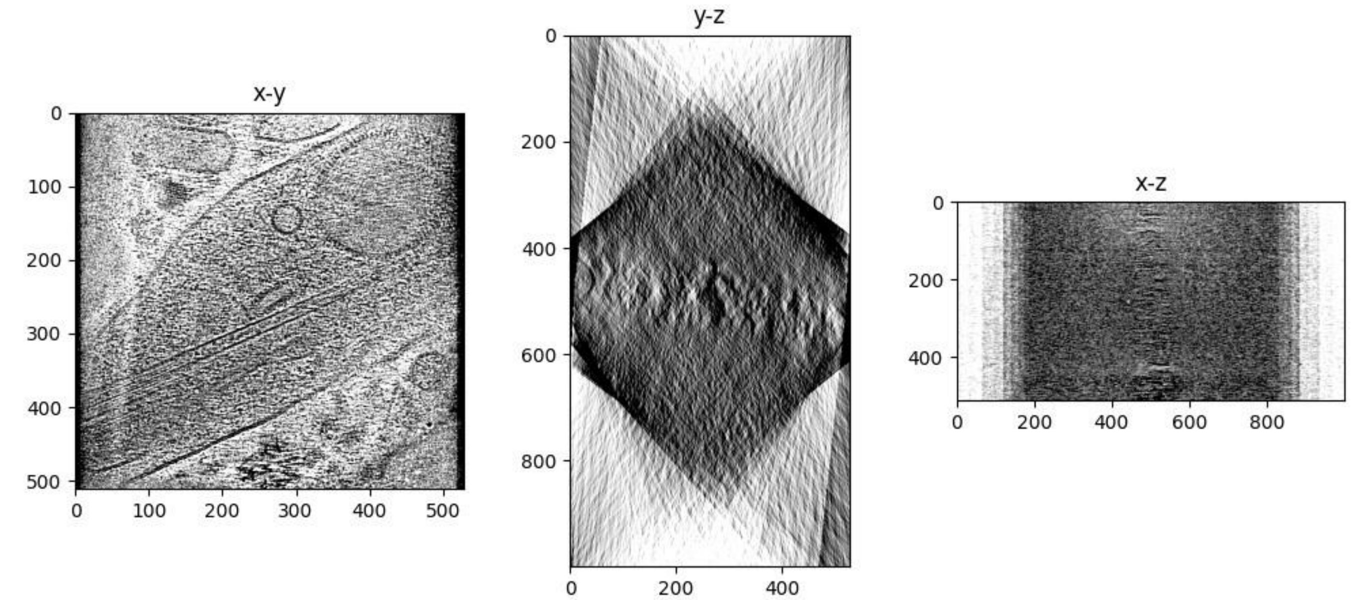


neurone_0023_ali_rec.mrc

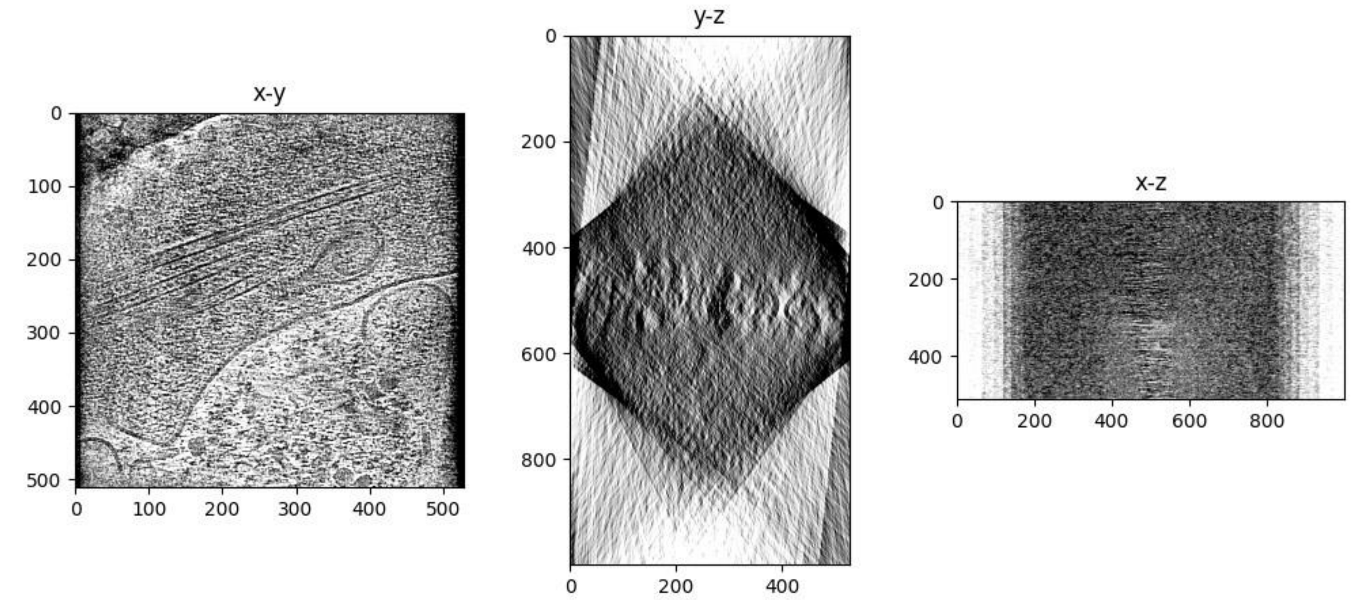

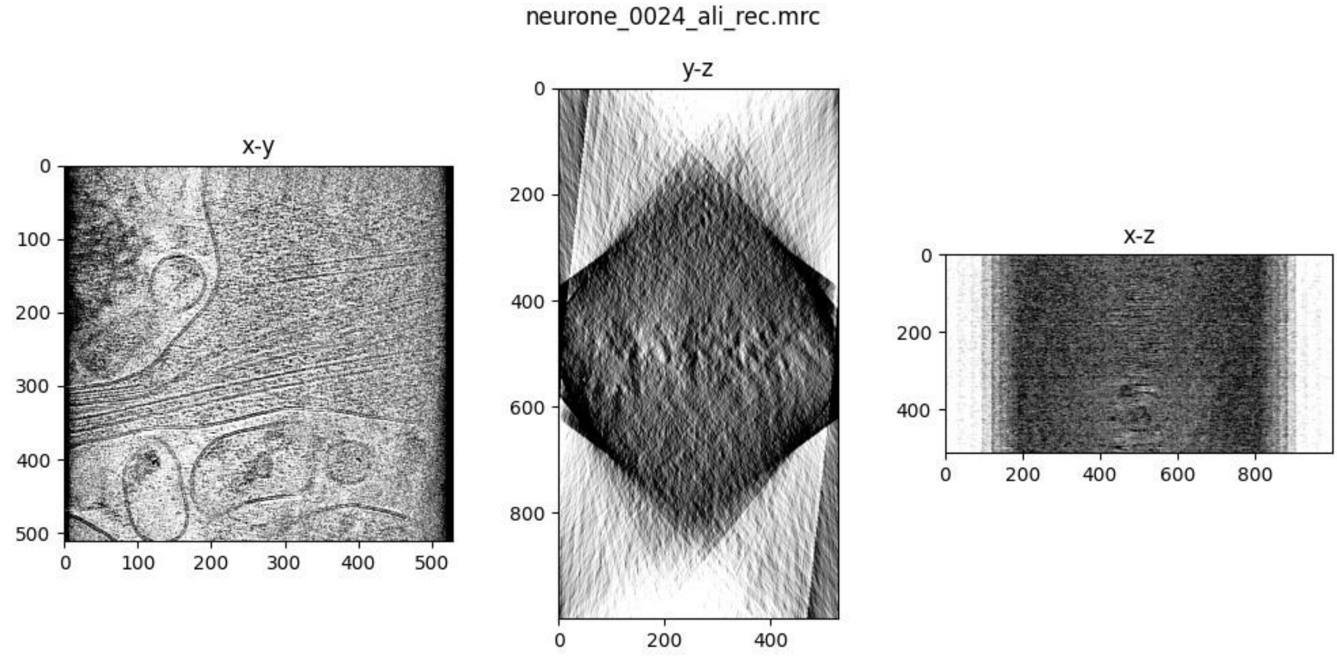
neurone_0024_ali_rec.mrc
x-y
y-z
x-z
0
100
200
300
400
500
600
800

### Savu Reconstruction

Reconstruction was performed with Savu 4.0.

Please cite:


1. Wadeson, N and Basham, M. Savu: A Python-based, MPI Framework for Simultaneous Processing of Multiple, N-dimensional, Large Tomography Datasets. *arXiv* **1610.08015**. (2016)
2. Wadeson, N, Verschoyle, J, Kazantsev, D., et al. DiamondLightSource/Savu: Version 4.0. *Zenodo*.
 (2021) DOI 10.5281/zenodo.5095360.
3. The HDF Group. Hierarchical Data Format, version 5. (1997-2016)
4. Walker, D and Dongarra, J. MPI: a standard message passing interface. *Supercomputer* **12**: 56-68. (1996)
5. Van Aarle, W, Palenstijn, W, Cant, J, et al. Fast and flexible X-ray tomography using the ASTRA toolbox. *Optics express* **24(22)**: 25129-25147. (2016)
6. Van Aarle, W, Palenstijn, W, De Beenhouwer, J, et al. The ASTRA Toolbox: A platform for advanced algorithm development in electron tomography. *Ultramicroscopy* **157**: 35-47. (2015)
7. Palenstijn, WJ and Batenburg, KJ and Sijbers, J. Performance improvements for iterative electron tomography reconstruction using graphics processing units (GPUs). *JSB* **176(2)** 250-253. (2011)

For more information, please see https://savu.readthedocs.io/en/latest/

**Note**

Savu was developed by the X-ray tomography community, so the conventional relationship between grey values and density is reversed compared to EM, i.e., the colours are reversed. The visualisations in this report have had their colours inverted to match the EM convention.

Reconstruction algorithm used: CGLS_CUDA

#### Tomogram thumbnails

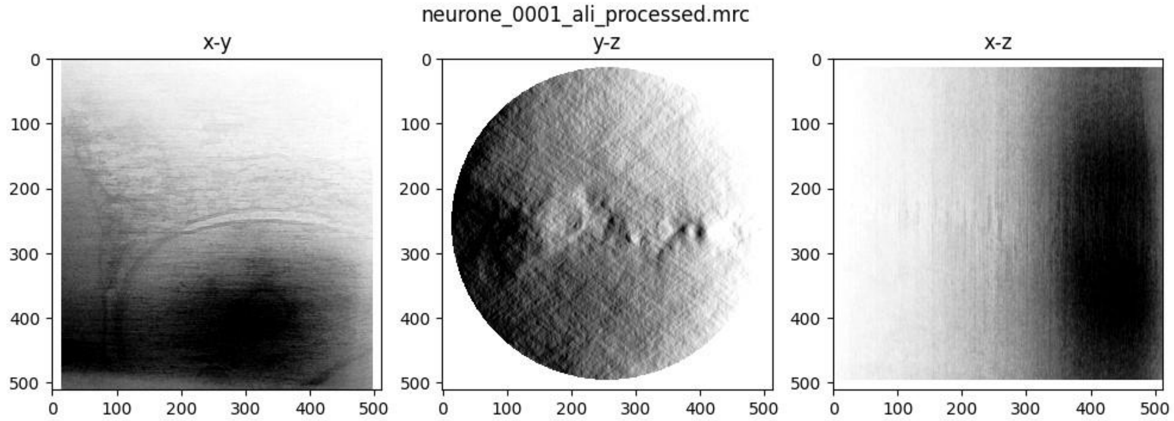


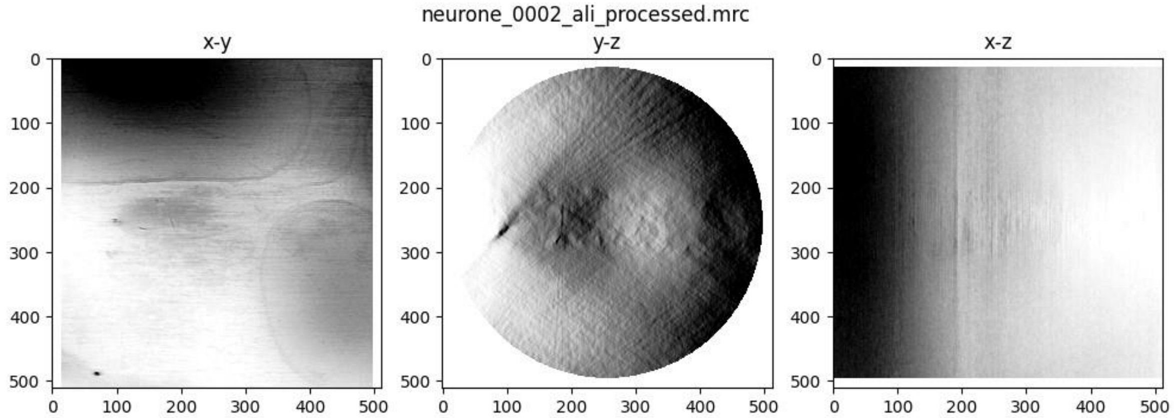


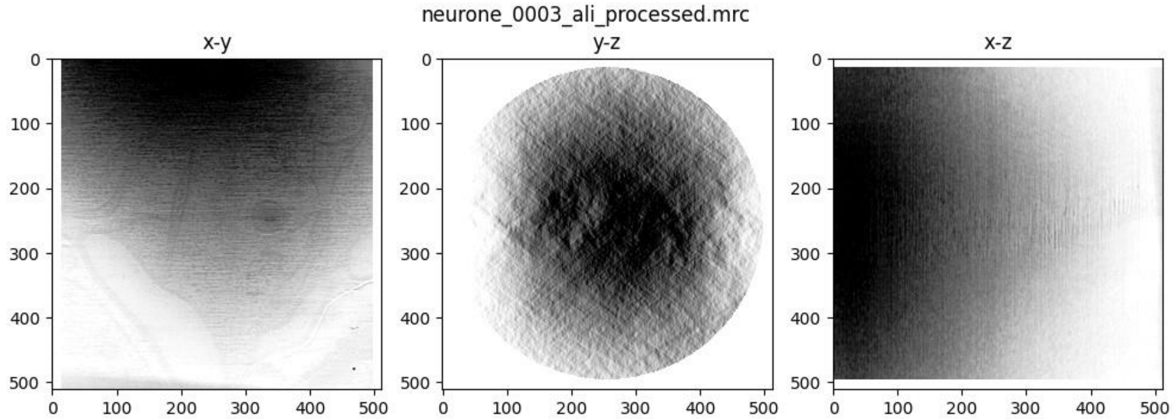


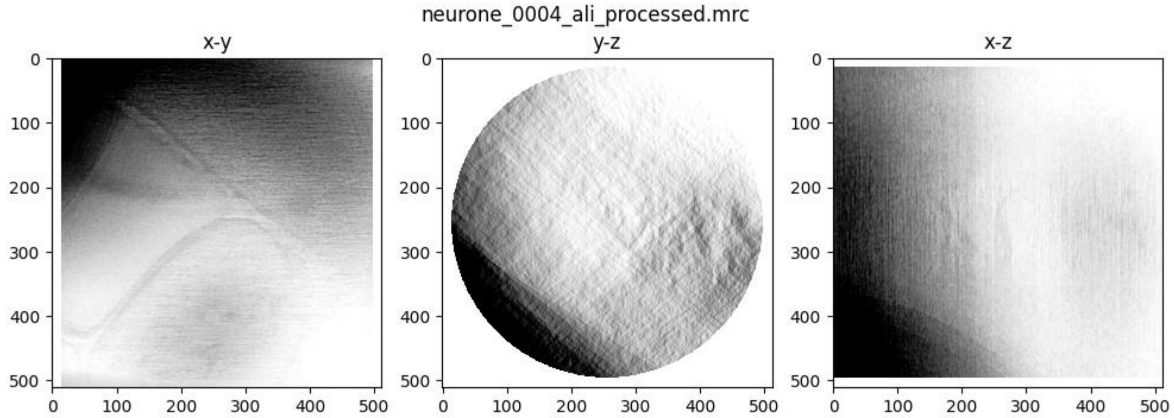

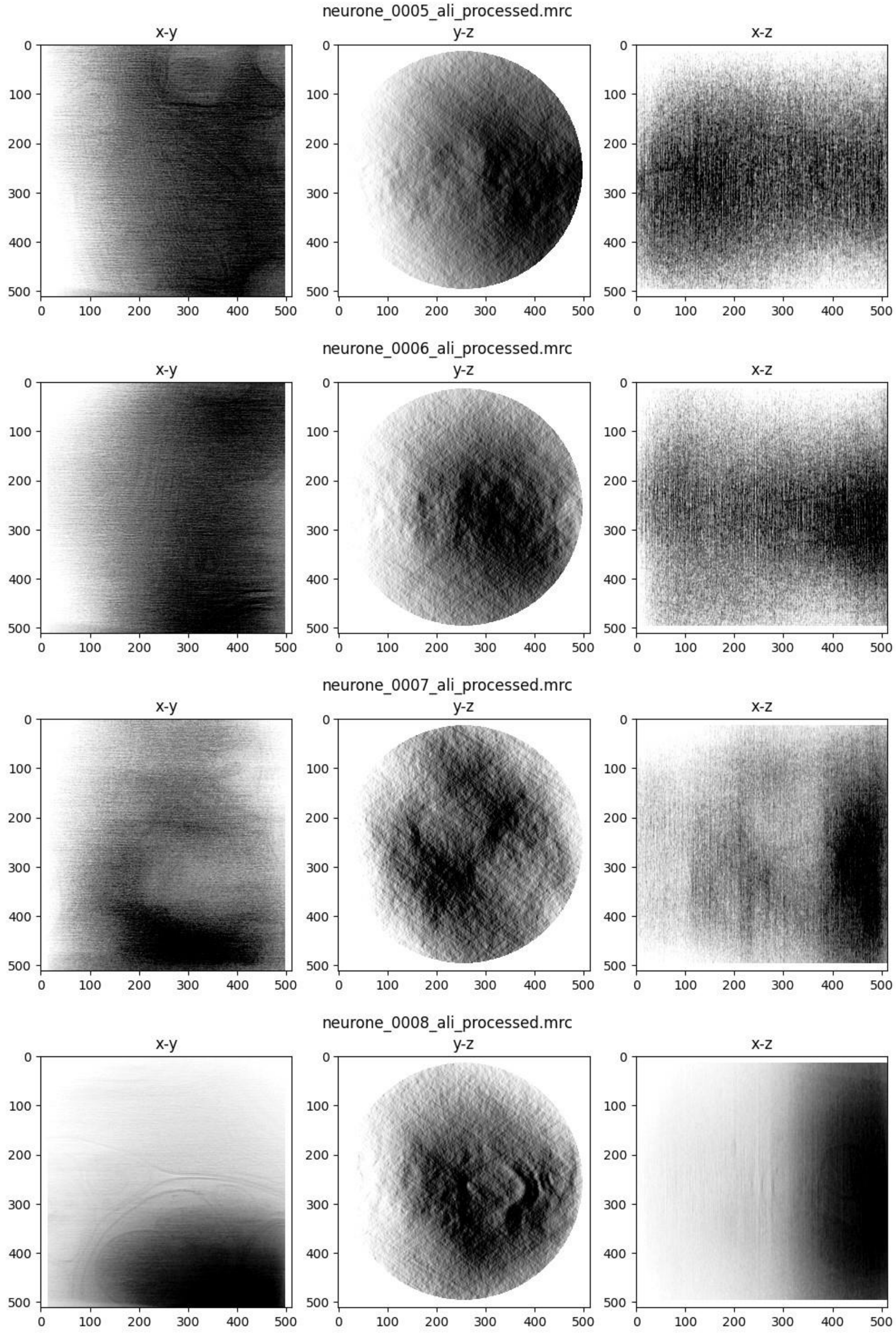
neurone_0005_ali_processed.mrc
x-y
y-z
x-z
neurone_0006_ali_processed.mrc
x-y
y-z
x-z
neurone_0007_ali_processed.mrc
x-y
y-z
x-z
neurone_0008_ali_processed.mrc
x-y
y-z
x-z

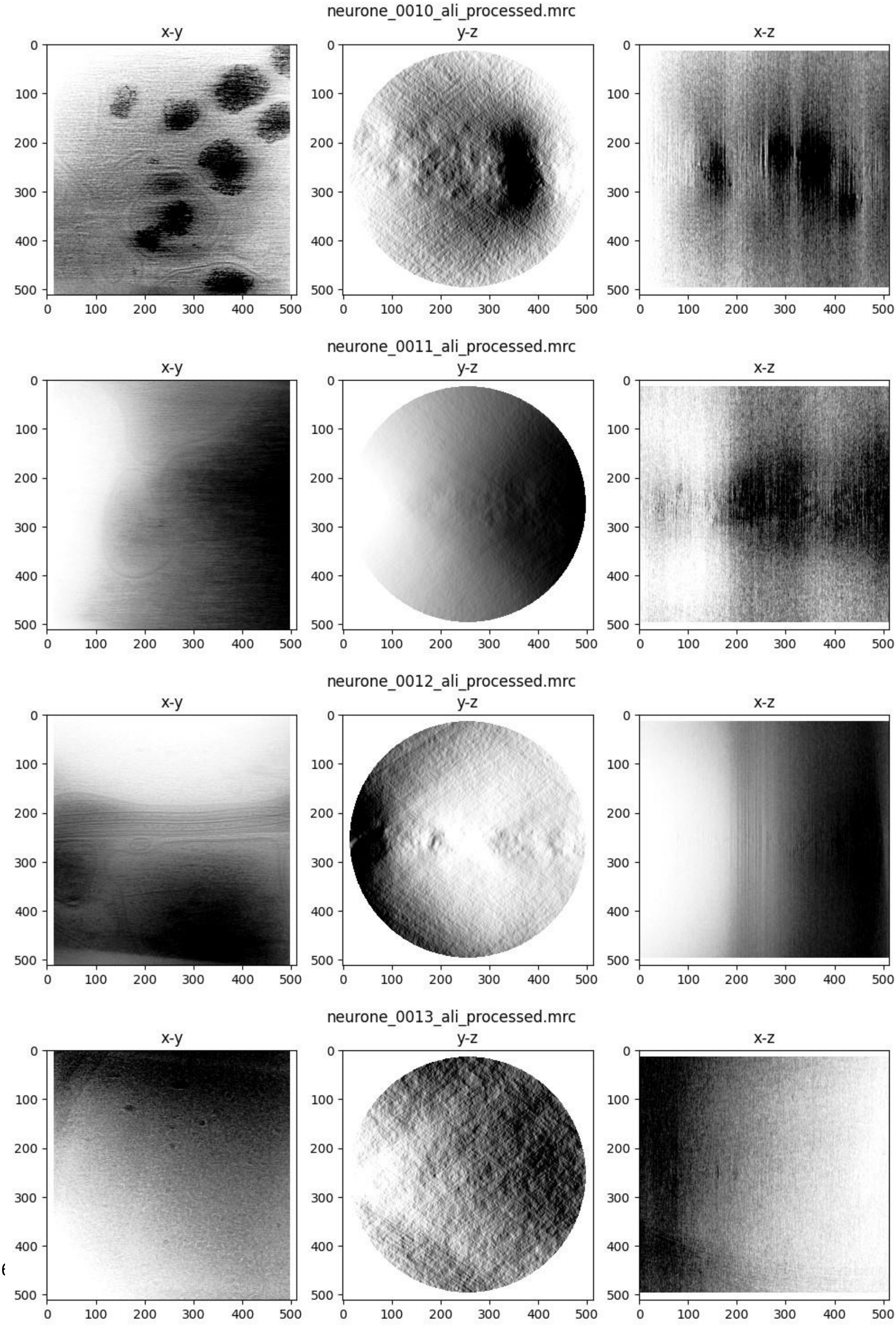
neurone_0010_ali_processed.mrc
x-y
y-z
x-z
neurone_0011_ali_processed.mrc
x-y
y-z
x-z
neurone_0012_ali_processed.mrc
x-y
y-z
x-z
neurone_0013_ali_processed.mrc
x-y
y-z
x-z

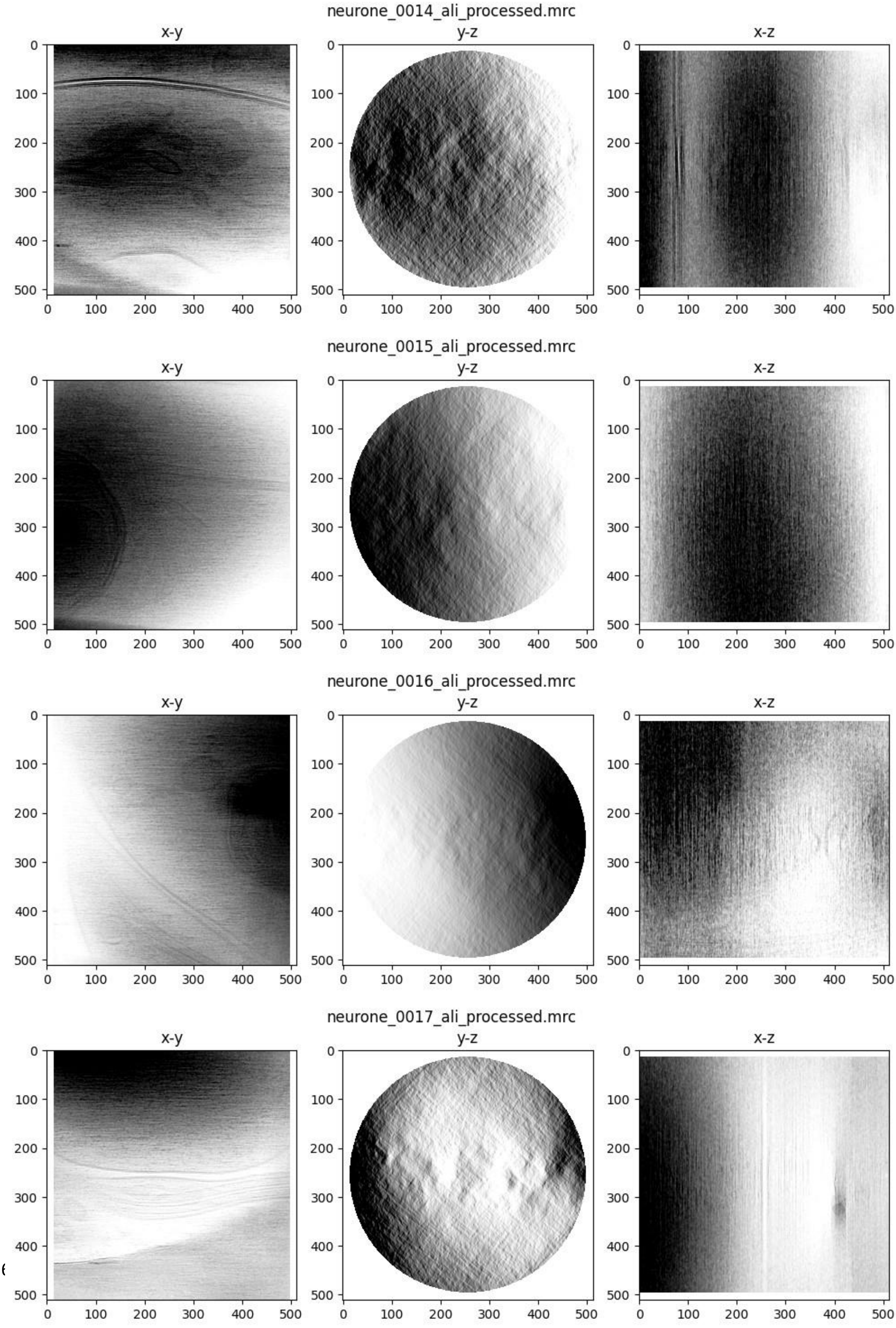
neurone_0014_ali_processed.mrc
x-y
y-z
x-z
neurone_0015_ali_processed.mrc
x-y
y-z
x-z
neurone_0016_ali_processed.mrc
x-y
y-z
x-z
neurone_0017_ali_processed.mrc
x-y
y-z
x-z

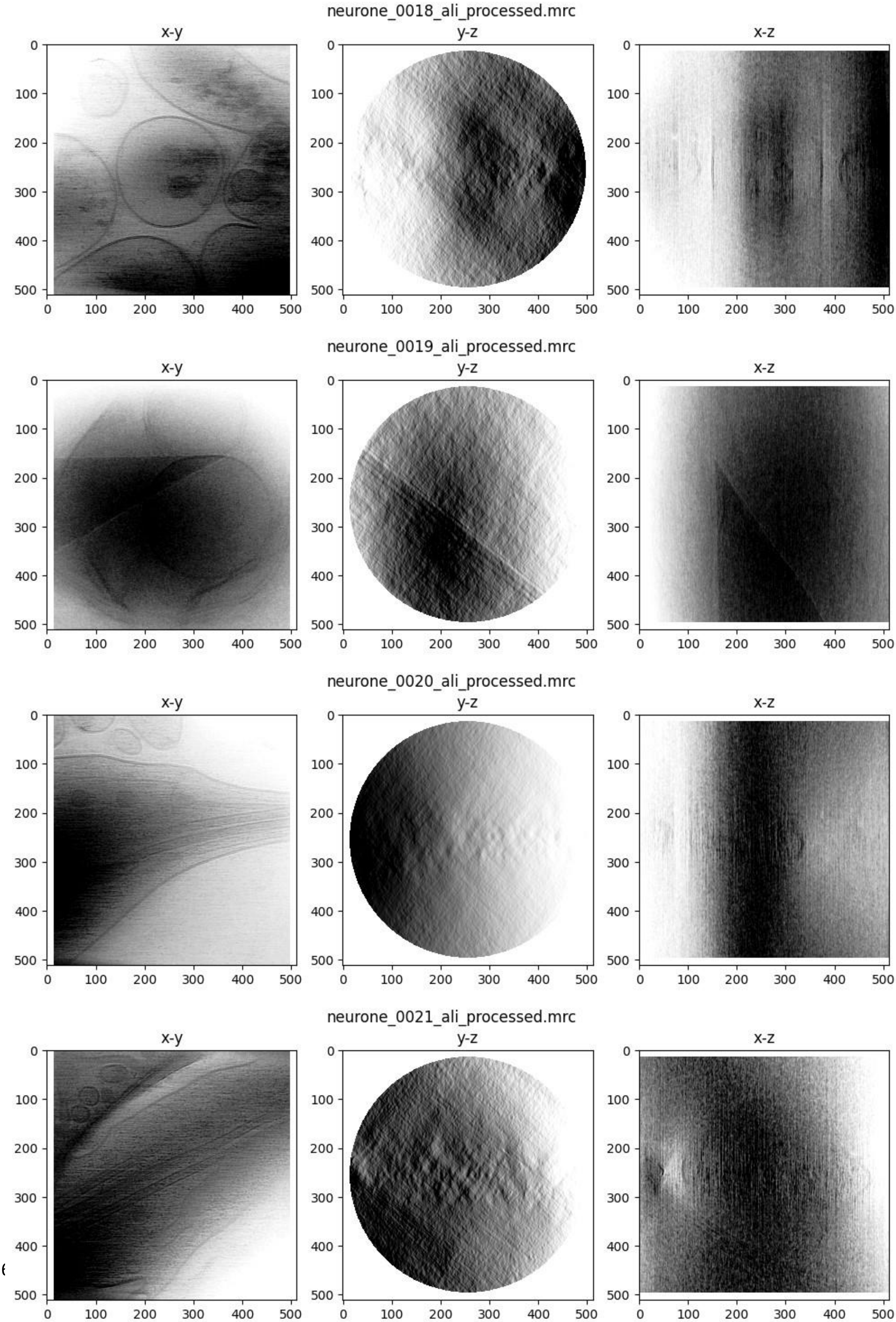

neurone_0018_ali_processed.mrc
x-y
y-z
x-z
neurone_0019_ali_processed.mrc
x-y
y-z
x-z
neurone_0020_ali_processed.mrc
x-y
y-z
x-z
neurone_0021_ali_processed.mrc
x-y
y-z
x-z
0
100
200
300
400
500

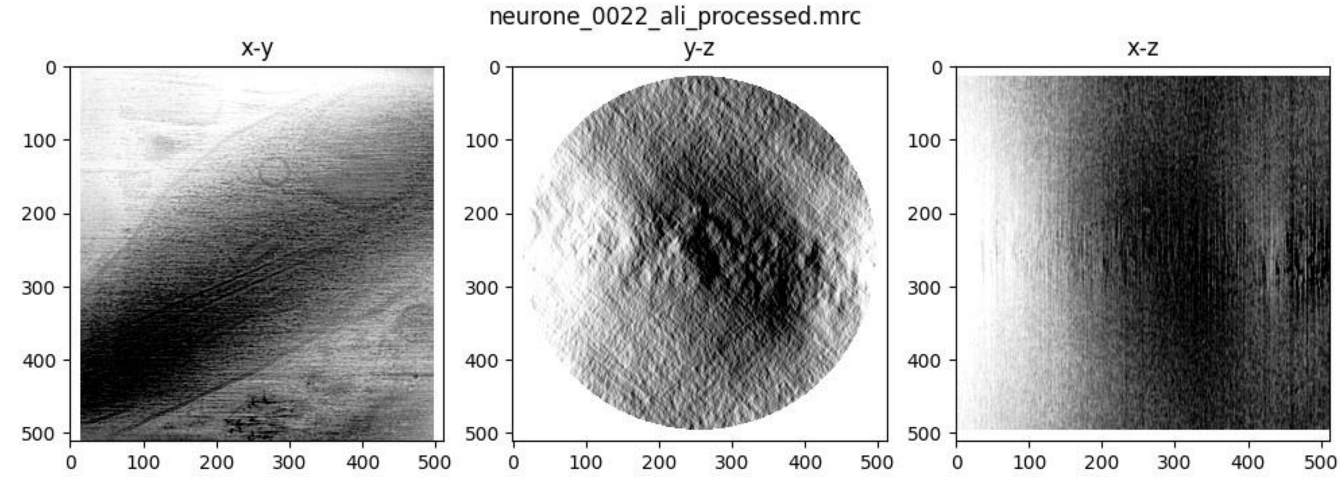
neurone_0022_ali_processed.mrc
x-y
y-z
x-z

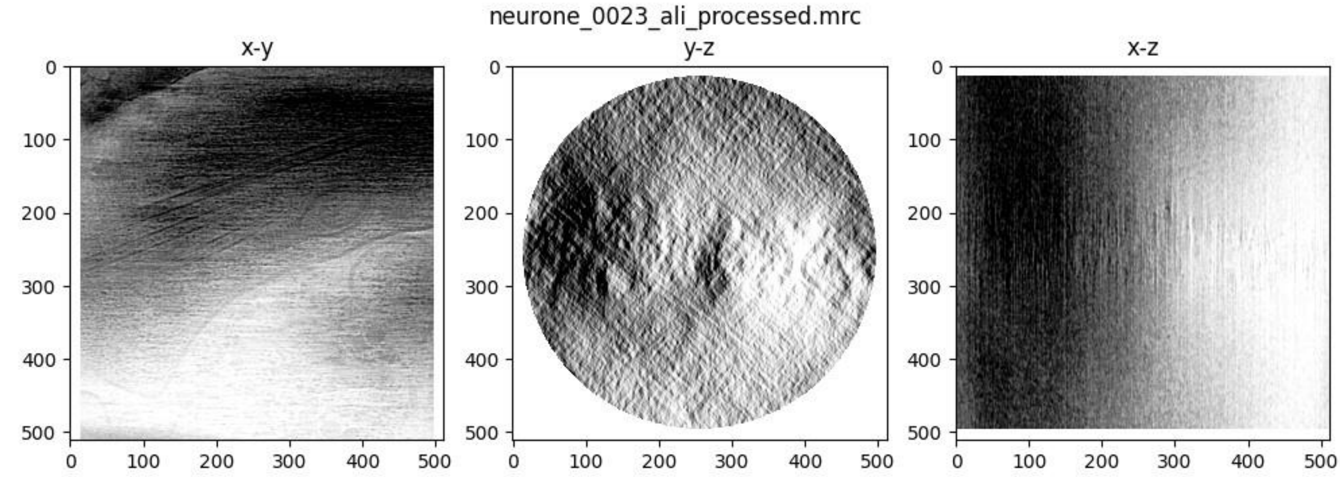
neurone_0023_ali_processed.mrc
x-y
y-z
x-z

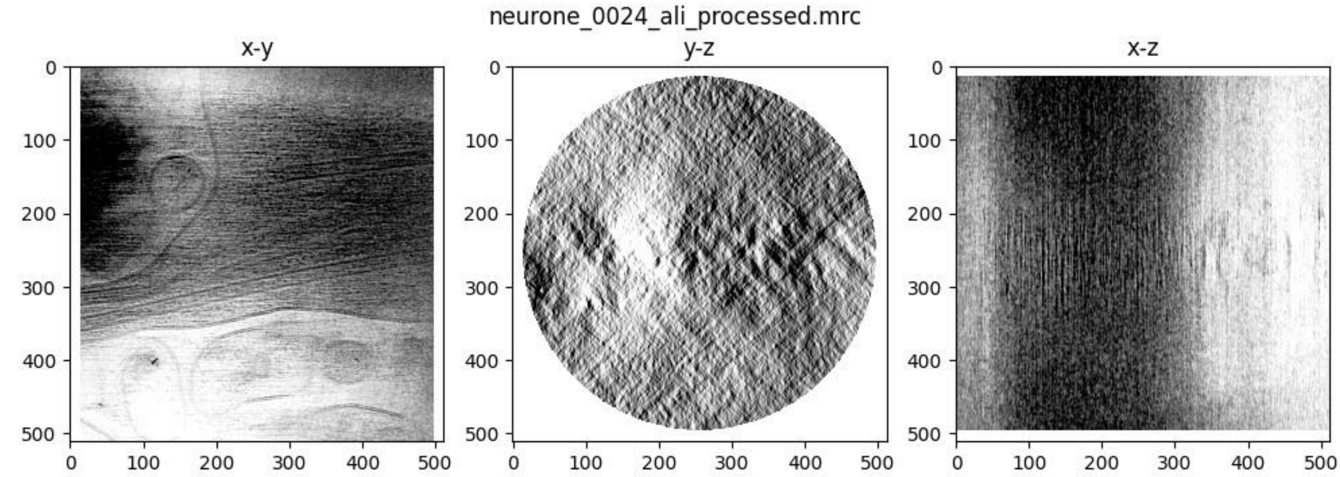
neurone_0024_ali_processed.mrc
x-y
y-z
x-z

Table S1: Configuration parameters for Case Study 1.

| | |
|---|---|
| **Motion correction** | |
| **Input filetype** | MRC |
| **Gain reference** | No gain |
| **Pixel size (A)** | 2.243 |
| **Desired pixel size (A)** | 2.243 |
| **Discard frames** | None |
| **Tolerance** | 0.5 |
| **Max number of iterations** | 10 |
| **Patch size** | 5 x 5 x 20 |
| **Use subgroups** | Yes |
| **IMOD Alignment** | |
| **Excluded views** | None |
| **Use rawtlt** | Yes |
| **Pixel size (nm)** | 0.2243 |
| **Rotation angle (deg)** | 175.51 |
| **Stack bin factor** | 2 |
| **Remove x-rays** | Yes |
| **Coarse align** | |
| **Patch size** | 210 x 203 |
| **Number of patches** | 24 x 24 |
| **Number of iterations** | 4 |
| **Limits on shifts** | 2, 2 |
| **Adjust tilt angles** | Yes |
| **Fine align** | |
| **Number of surfaces** | 1 |
| **Magnification option** | fixed |
| **Tilt option** | fixed |
| **Rotation option** | group |
| **Beam tilt option** | fixed |
| **Use robust fitting** | Yes |
| **Weight all contours** | Yes |
| **IMOD Reconstruction** | |
| **Use rawtlt** | Yes |
| **Pixel size (nm)** | 0.4486 |
| **Rotation angle (deg)** | 175.51 |
| **Do positioning** | No |
| **Unbinned thickness** | 1920 |
| **Correct CTF** | No |
| **Erase gold** | No |
| **2D filtering** | No |
| **Bin factor** | 1 |
| **Reconstruction thickness** | 1920 |
| **Algorithm** | WBP |
| **Trim volume** | Yes |
| **Trim volume reorientation** | Rotate |

Table S2: Configuration parameters for Case Study 2

| **Motion correction** | |
|---|---|
| **Input filetype** | MRC |
| **Gain reference** | No gain |
| **Pixel size (A)** | 1.87 |
| **Desired pixel size (A)** | 1.87 |
| **Discard frames** | None |
| **Tolerance** | 0.5 |
| **Max number of iterations** | 10 |
| **Patch size** | 5 x 5 x 20 |
| **Use subgroups** | Yes |
| **IMOD Alignment** | |
| **Excluded views** | None |
| **Use rawtlt** | Yes |
| **Pixel size (nm)** | 0.187 |
| **Rotation angle (deg)** | -83.96 |
| **Stack bin factor** | 8 |
| **Remove x-rays** | Yes |
| **Coarse align** | |
| **Patch size** | 224 x 224 |
| **Number of patches** | 24 x 24 |
| **Number of iterations** | 4 |
| **Limits on shifts** | 2, 2 |
| **Adjust tilt angles** | Yes |
| **Fine align** | |
| **Number of surfaces** | 1 |
| **Magnification option** | Fixed |
| **Tilt option** | Fixed |
| **Rotation option** | Group |
| **Beam tilt option** | Fixed |
| **Use robust fitting** | Yes |
| **Weight all contours** | Yes |

| **IMOD Reconstruction** | |
|---|---|
| **Use rawtlt** | Yes |
| **Pixel size (nm)** | 1.496 |
| **Rotation angle (deg)** | -83.96 |
| **Do positioning** | No |
| **Unbinned thickness** | 4096 |
| **Correct CTF** | No |
| **Erase gold** | No |
| **2D filtering** | No |
| **Bin factor** | 8 |
| **Reconstruction thickness** | 4096 |
| **Algorithm** | WBP |
| **Trim volume** | Yes |
| **Trim volume reorientation** | Rotate |
| **CTFFind4** | |
| **Pixel size (A)** | 1.87 |
| **Voltage (kV)** | 300 |
| **Spherical aberration** | 2.7 |
| **Amp contrast** | 0.8 |
| **Amp spec size** | 512 |
| **Min resolution** | 30 |
| **Max resolution** | 5 |
| **Defocus min** | 5000 |
| **Defocus max** | 50000 |
| **Defocus step** | 500 |
| **Astigmatism type** | None |
| **Exhaustive search** | No |
| **Astigmatism restraint** | No |
| **Phase shift** | No |

Table S3: Configuration parameters for Case Study 3

| | |
|---|---|
| **Motion correction** | |
| **Input filetype** | EER |
| **Gain reference** | No gain |
| **Pixel size (A)** | 1.47 |
| **Desired pixel size (A)** | 1.47 |
| **Discard frames** | None |
| **Tolerance** | 0.5 |
| **Max number of iterations** | 10 |
| **Patch size** | 5 x 5 x 20 |
| **Use subgroups** | Yes |
| **IMOD Alignment** | |
| **Excluded views** | None |
| **Use rawtlt** | Yes |
| **Pixel size (nm)** | 0.147 |
| **Rotation angle (deg)** | -84.22 |
| **Stack bin factor** | 8 |
| **Remove x-rays** | Yes |
| **Coarse align** | |
| **Patch size** | 224 x 224 |
| **Number of patches** | 24 x 24 |
| **Number of iterations** | 4 |
| **Limits on shifts** | 2, 2 |
| **Adjust tilt angles** | Yes |
| **Fine align** | |
| **Number of surfaces** | 1 |
| **Magnification option** | Fixed |
| **Tilt option** | Fixed |
| **Rotation option** | Group |
| **Beam tilt option** | Fixed |

| | |
|---|---|
| **Use robust fitting** | Yes |
| **Weight all contours** | Yes |
| **IMOD Reconstruction** | |
| **Use rawtlt** | Yes |
| **Pixel size (nm)** | 1.176 |
| **Rotation angle (deg)** | -84.22 |
| **Do positioning** | No |
| **Unbinned thickness** | 512 |
| **Correct CTF** | No |
| **Erase gold** | No |
| **2D filtering** | No |
| **Bin factor** | 8 |
| **Reconstruction thickness** | 4096 |
| **Algorithm** | SIRT |
| **SIRT iterations** | 10 |
| **Trim volume** | Yes |
| **Trim volume reorientation** | Rotate |
| **AreTomo alignment** | |
| **Bin factor** | 8 |
| **AreTomo reconstruction** | |
| **Rotation angle** | -84.22 |
| **Bin factor** | 1 |
| **VolZ** | 1000 |
| **Pixel size (A)** | 1.47 |
| **Algorithm** | SART |
| **Savu reconstruction** | |
| **Algorithm** | CGLS |
| **Number of iterations** | 100 |
| **Centre of rotation** | 256 |

Guide to writing plugins for Ot2Rec, available at
https://github.com/rosalindfranklininstitute/Ot2Rec/wiki/Tutorial:-How-to-write-an-Ot2Rec-plugin

### Guide to writing plugins for Ot2Rec v0.2.x

#### Introduction

Plugins in Ot2Rec achieve four main objectives:

1. Capture user arguments for the task. (These are saved in a `yaml` file)
2. Generate the correct commands to call other programs to complete the task, e.g. IMOD
3. Save the metadata along the way
4. (Optionally) add a section to the report showing how the task performed.

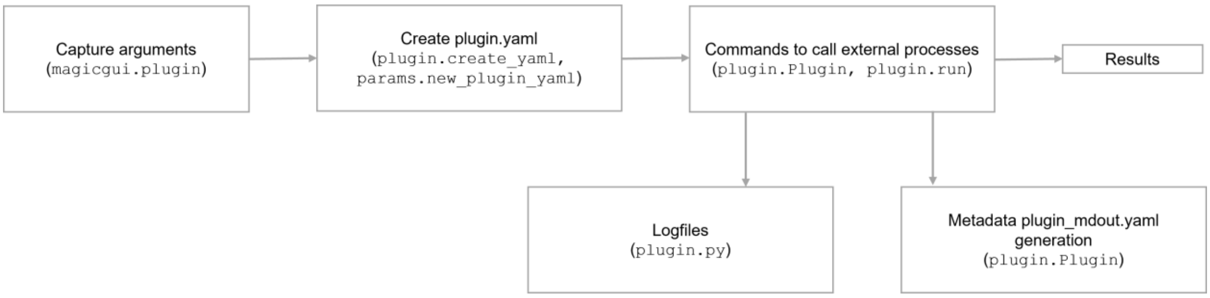


This guide will show you how to write an Ot2Rec plugin called `plugin`.

#### To-Do list

- Add the user arguments to `magicgui.py`
- Add the desired layout of `plugin.yaml` to `params.py`
- Create `plugin.py`, which holds most of the logic behind your new plugin.
- Add your command-line binding to `setup.py`. This binding allows users to call your plugin straight from the terminal.

#### Step 0: Setup your conda environment

Clone Ot2Rec from Github:

```
git clone https://github.com/rosalindfranklininstitute/Ot2Rec.git
```

Create a new conda environment and install Ot2Rec in editable mode:

```
conda create -n ot2rec
```

```
conda activate ot2rec
pip install -e .
```

This should update your installation of Ot2Rec when changes are made so you are always running the latest version.

#### Step 1: Capture user arguments

We use `magicgui` to generate Ot2Rec's GUI to capture user arguments. These argument capture functions are added to the `magicgui.py` file. One GUI is created for each plugin, an example is shown below:
**add image here**

Here is a minimal example of the magicgui functions we need for our plugin.

```
@mg(
    call_button="Create config file",
    layout="vertical",
    result_widget=False,

    project_name={"label": "Project name *"},
    pixel_size={"label": "Pixel size in A",
                "min": 0.001,
    },
    rootname={"label": "Rootname of current project (required if different from
project name"},
    suffix={"label": "Suffix of project files"},
    input_mrc_folder={
        "label": "Folder containing input mrc's",
        "mode": "d",
    },
    output_path={
        "label": "Path to output folder",
        "mode": "d",
    },
    choice_abc={
        "label": "Choice between a, b, c",
        "choices": ["a", "b", "c]
    },
)
def get_args_plugin(
        project_name="",
        pixel_size=0.00,
        rootname="",
        suffix="",
        input_mrc_folder=Path("./plugin_input"),
        output_path=Path("./plugin_output"),
```

```
        recon_algo="WBP",
):
    return locals()
```

First, we use the `magicgui` decorator `@mg` to create our widgets. The widgets we have on all our plugins are:

- `project_name` | name of the dataset we are processing
- `rootname` | prefix of all the files we want to work with, defaults to the `project_name` if not specified.
- `suffix` | suffix added to the end of the filenames, can be left as `None`.
- some sort of input, e.g., `input_mrc_folder`
- an output path, e.g., `output_path`

### Step 2: Create template of yaml to hold arguments

The `plugin.yaml` file holds the information needed to set up the task to be performed, e.g., input filepaths, parameters, output filepaths. Some of these are captured from the `magicgui` in the previous section, but others can be calculated or populated with default values.
The `yaml` file is first generated from a template in `params.py`, its values are then populated by code in `plugin.py` with user arguments captured from `magicgui.py`.
An example `yaml` file for our plugin looks like:

```
System:
    process_list:
    - 1
    - 2
    - 3
    output_path: ./plugin_processed
    output_rootname: neurone
    output_suffix: ''
Plugin_setup:
    pixel_size: 1.0
    input_mrc:
    - ./aligned/neurone_0001/neurone_0001.st
    - ./aligned/neurone_0002/neurone_0002.st
    - ./aligned/neurone_0003/neurone_0003.st
    output_mrc:
    - ./aligned/neurone_0001/neurone_0001_ali.mrc
    - ./aligned/neurone_0002/neurone_0002_ali.mrc
    - ./aligned/neurone_0003/neurone_0003_ali.mrc
    tilt_angles:
    - ./aligned/neurone_0001/neurone_0001.rawtlt
    - ./aligned/neurone_0002/neurone_0002.rawtlt
    - ./aligned/neurone_0003/neurone_0003.rawtlt
    choice: a
```

Here, the `process_list`, `input_mrc`, `output_mrc`, and `tilt_angles` fields are populated automatically in the `plugin.update_yaml` function in `plugin.py`.
To create the template `plugin.yaml` file, we add the following to `params.py`:

```
def new_plugin_yaml(args):
    """
    Subroutine to create yaml file for plugin

    ARGS:
    args (Namespace) :: Namespace containing user parameter inputs
    """

    plugin_yaml_name = args.project_name.value + '_plugin.yaml'

    plugin_yaml_dict = {
        'System': {
            'process_list': None,
            'output_path': str(args.output_path.value),
            'output_rootname': args.project_name.value if args.rootname.value is
None else args.rootname.value,
            'output_suffix': args.suffix.value,
        },

        'Plugin_setup': {
            'pixel_size': args.pixel_size.value,
            'input_mrc': None,
            'output_mrc': None,
            'tilt_angles': None,
            'choice': args.choice.value,
        }
    }

    with open(plugin_yaml_name, 'w') as f:
        yaml.dump(plugin_yaml_dict, f, indent=4, sort_keys=False)
```

Don't worry about populating all the fields just now, just fill in those which can be obtained from the `args`, i.e., the user arguments captured from `magicgui`.
Next, we will need to create the `plugin.py` file, which will hold most of the logic needed for the plugin. We will go through this in two parts, here we will write the sections which deal with creating the `yaml` to setup the plugin.
Add the `create_yaml` and `update_yaml` methods to `plugin.py`.

```
def update_yaml(args):
    """Method to update yaml file

    Here we set the process list, specific filepaths, and check inputs

    ARGS:
    args (magicgui.FunctionGUI) :: magicgui object containing user input
    """
```

```python
    # Read in template yaml file
    plugin_yaml_name = f"{args["project_name"]}_plugin.yaml"
    plugin_params = prmMod.read_yaml(
        project_name=args["project_name"],
        filename=plugin_yaml_name
    )

    # Set input filepaths, can be by searching for specific filenames in the input
mrc folder, then update the yaml file
    input_files = glob(f"{args['input_mrc']}/*.mrc")
    plugin_params.params["Plugin_setup"]["input_mrc"] = input_files

    # Can do similar for the output filepaths

    # Set process list, this is usually the tilt series index, can be parsed from
filepaths

    # Update yaml
    with open(Path(plugin_yaml_name), "w") as f:
        yaml.dump(plugin_params.params, f, indent=4, sort_keys=False)

def create_yaml(input_mgNS=None):
    """
    Subroutine to create new yaml file for Plugin
    """

    # Parse user inputs
    if input_mgNS is None:
        args = mgMod.get_args_plugin.show(run=True).asdict()
    else:
        args = input_mgNS

    # Create the yaml file, then automatically update it
    prmMod.new_plugin_yaml(args)
    update_yaml(args)
```

Now the functions to create the yaml should be in place, and we can move on to generating and running commands based on the yaml file.

#### Step 3: Functions to do `plugin`'s work

The `Plugin` class in `plugin.py` creates the results folders, generates commands to run the plugin, runs the plugin, and saves the metadata.
Let's write the `Plugin` class in stages. First, set up the `Plugin` class. All `Ot2Rec` plugin classes have the same fundamental attributes

- `project_name`
- `params_in`: parameters read from the yaml file

- `logger_in`: a logger object to create the logfile `o2r_plugin.log`
- `md_out`: dictionary of metadata to pass to `plugin_mdout.yaml`.

In the `__init__`, we also want to set up the results folder structure, which is done in the `_get_internal_metadata()` function.

```
class Plugin:
    def __init__(self, project_name, params_in, logger_in):
        self.proj_name = project_name
        self.params = params_in.params
        self.logObj = logger_in
        self.md_out = {}

        self._get_internal_metadata()

    def _get_internal_metadata(self):
        """Prep internal metadata for processing and checking

        ** See other plugins for code to reuse **

        Generally, this sets the input and output filepaths and creates results
folders.

        You can also pass information to the md_out, which is a metadata yaml file
written after processing. e.g., putting in filepaths to the results.
        """
        for curr_ts in self.params["System"]["process_list"]:
            subfolder = (f"{self.basis_folder}/"
                         f"{self.rootname}_{curr_ts:04d}{self.suffix}")
            os.makedirs(subfolder, exist_ok=True)

            self.md_out["plugin_output_dir"][curr_ts] = subfolder
            self.md_out["plugin_output_file"][curr_ts] = f"{subfolder}/example.st
```

Next, we want to generate the commands needed to run the plugin. These are the same commands we would use if we were to use the plugin program directly (e.g., IMOD, AreTomo) in the terminal.

Add the following to the `Plugin` class.

```
    def _get_plugin_command(self, i):
        """Get command to call an external process for the i-th tilt series"""
        cmd = [
            "plugin",
            "-input",
            self.params["Plugin_setup"]["input_mrc"][i],
            "-output",
            self.params["Plugin_setup"]["output_mrc"][i],
            "-tiltangle",
            self.params["Plugin_setup"]["tilt_angles"][i],
```

```
            "-choice",
            self.params["Plugin_setup"]["choice]
        ]

        return cmd
```
We also want to add the runner functions to the `Plugin` class.
```
    def _run_plugin(self, i):
        """Run the plugin for the i-th tilt series"""
        cmd = self._get_plugin_command(i)
        plugin_run = subprocess.run(
            cmd,
            stdout=subprocess.PIPE,
            stderr=subprocess.STDOUT,
            encoding="ascii",
            check=True
        )
        self.logObj(plugin_run.stdout) # save stdout to log

    def run_plugin_all(self):
        for i, ts in enumerate(self.params["System"]["process_list"]):
            self._run_plugin(i)
        self.export_metadata()

    def export_metadata(self):
        yaml_file = self.proj_name + "_plugin_mdout.yaml"
        with open(yaml_file, "w") as f:
            yaml.dump(self.md_out, f, indent=4, sort_keys=False)
```
Lastly, we will add the `run` function to `plugin.py`, which is called by `o2r.plugin.run`.
```
def run():
    """
    Method to run plugin
    """
    # argparser to collect the project name
    parser = argparse.ArgumentParser()
    parser.add_argument("project_name",
                        type=str,
                        help="Name of current project")
    args = parser.parse_args()

    # Check if prerequisite files exist
    plugin_yaml_name = f"{args["project_name"]}_plugin.yaml"
    if not os.path.isfile(plugin_yaml_name):
        raise IOError("Error in Ot2Rec.main.run_plugin: plugin yaml file not
found.")

    # Read in config and metadata
    plugin_config = prmMod.read_yaml(
```

```
        project_name=args.project_name,
        filename=plugin_yaml_name
    )

    # Create Logger object
    logger = logMod.Logger(log_path="o2r_plugin.log")

    # Create Plugin object
    plugin_obj = Plugin(
        project_name=args.project_name,
        params_in=plugin_config,
        logger_in=logger
    )

    # Run AreTomo commands
    plugin_obj.run_plugin_all()
```

#### Step 4: Add CLI binding

Lastly, add the plugin's entry points to `setup.py` to use it in the command line.
In `setup.py` under `entry_points`, add:
```
"o2r.plugin.new=Ot2Rec.plugin:create_yaml",
"o2r.plugin.run=Ot2Rec.plugin:run",
```

#### Step 5: Share your plugin

If you'd like your plugin to be part of the public Ot2Rec, raise a pull request and we'll review your code and merge it if everything works well.

Tutorial to reproduce Case Study 1, available from https://github.com/rosalindfranklininstitute/Ot2Rec/wiki/Tutorial:-Basic-reconstruction-workflow-in-Ot2Rec

This tutorial covers tomogram reconstruction of EMPIAR 10364 with Ot2Rec. Here we've assumed that Ot2Rec, Ot2Rec Report, MotionCor2 and IMOD are already installed on your system. If you need help with this, please see the Ot2Rec / Ot2Rec Report wikis, the MotionCor2 installation page, or the IMOD installation page. You'll need to run this on a GPU enabled Linux system. For reference, our system we used for this tutorial was a virtual Linux machine with 12 Intel Xeon Gold 5218 2.30GHz CPUs and 1 Tesla V100 GP.

1. Download the example data

2. Open a terminal and navigate to where the data is stored (replace /path/to/data with the actual path)

```
cd /path/to/data
```

3. Create a new folder – this will be where your processing will be stored.

```
mkdir processing
cd processing
```

4. Start a new Ot2Rec project by running

```
o2r.new
```

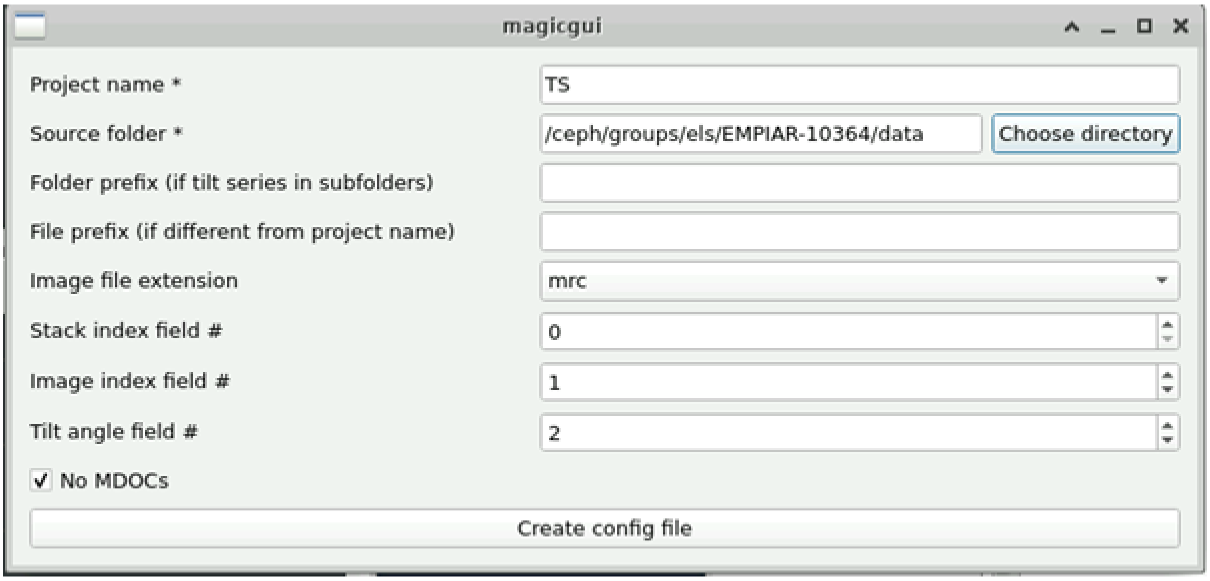


This creates a new Ot2Rec project and brings up the GUI to fill in the required parameters.

For this dataset, an example image filename is:

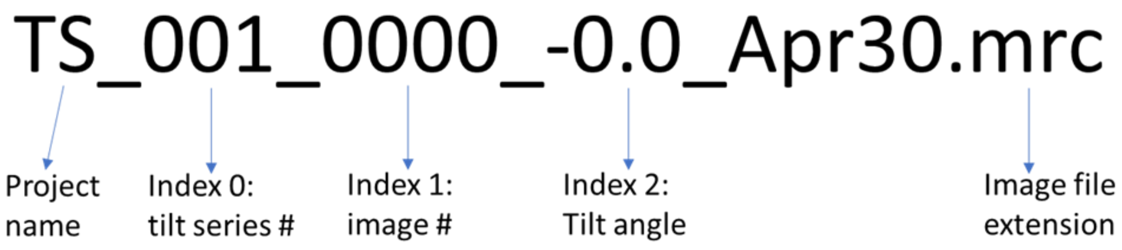


Click on create config file and close the GUI. There will now be 3 new files created:

- new_proj.yaml – a logfile capturing output from Ot2Rec
- TS_proj.yaml – a metadata file showing the parameters you've just filled into the GUI
- TS_master_md.yaml – this holds all a record of the tilt angles, filenames, tilt series number, and image number, which have been extracted from the filenames. Ot2Rec uses this information to generate the commands to run processes on all the tilt series.

5. We will now set up the Ot2Rec motioncor2 plugin.

First, ensure that motioncor2 is available on your system (RFI users: use `module load motioncor2`)

```
o2r.mc.new
```

Fill in the project name and pixel size of the data. The pixel size can usually be found in the mdoc or in the mrc header of your images. The MC2 output folder field sets where the motion corrected images will be stored, in this case, in the motioncor folder. You can change this if you'd like to. We have left the remaining parameters at their default values but you are free to change them if you would like to. Note that the path to mc2 executable field will probably be specific to your institution or system, depending on how motioncor2 is installed.

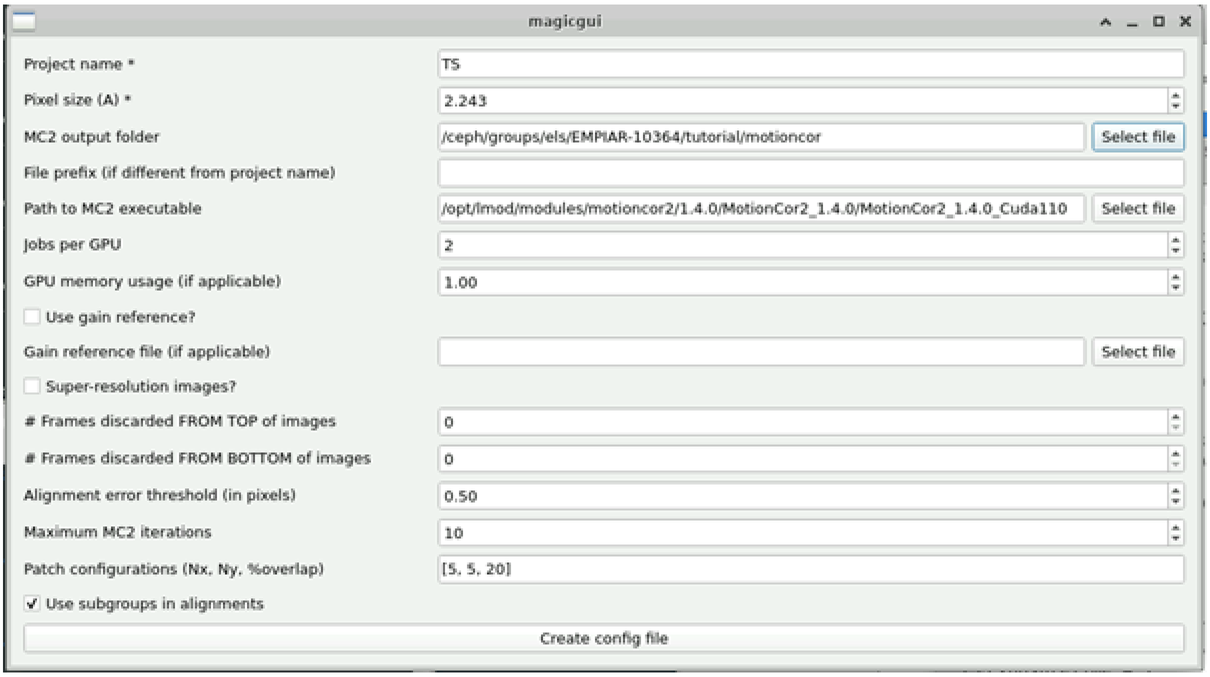


Again, click on the create config file button, close the dialog and two files will be created in your processing folder.

- o2r_motioncor2.log – this stores the output of motioncor which appears in the terminal.
- TS_mc2.yaml – this stores the user input you've added earlier, along with a process list.

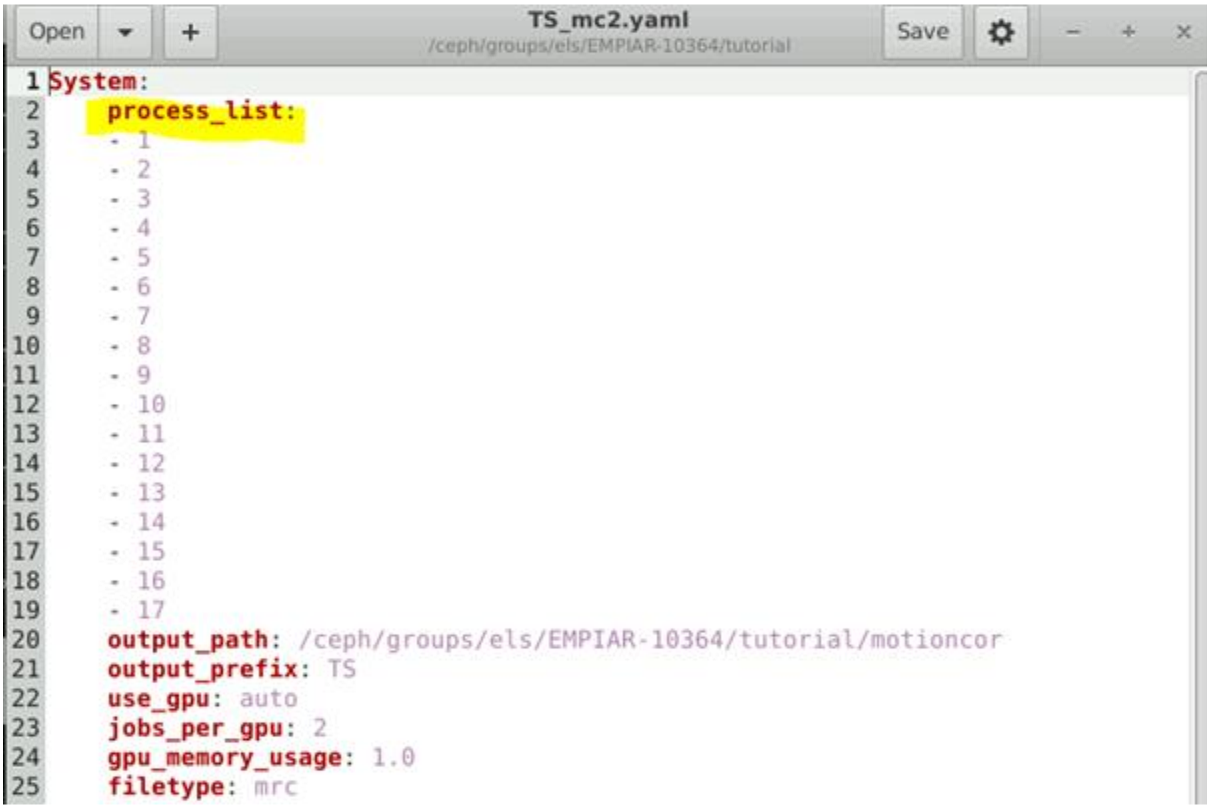

The process list is the list of tilt series that will be processed if we run this plugin. You can edit this manually to only use a selection of the tilt series if you would like to.

6. To run the motioncor2 plugin, we run

```
o2r.mc.run TS
```

Running Ot2Rec plugins requires the project name to be specified, so in this case we add "TS" after the run command to tell Ot2Rec to run the motion correction plugin on the project TS.

This will likely take a while but you can monitor the progress in the terminal. When complete, your motioncor folder should look like this:

| Name | Size | Type |
|---|---|---|
| TS_0001_2.0.mrc.log0-Patch-Frame.log | 5.4 KiB | application log |
| TS_0001_2.0.mrc.log0-Patch-Full.log | 193 bytes | application log |
| TS_0001_2.0.mrc.log0-Patch-Patch.log | 5.8 KiB | application log |
| TS_0001_2.0.mrc | 54.3 MiB | unknown |
| TS_0001_4.0.mrc.log0-Patch-Frame.log | 5.4 KiB | application log |
| TS_0001_4.0.mrc.log0-Patch-Full.log | 193 bytes | application log |
| TS_0001_4.0.mrc.log0-Patch-Patch.log | 5.8 KiB | application log |
| TS_0001_4.0.mrc | 54.3 MiB | unknown |

7. Now we will set up the tilt series alignment in IMOD.

```
o2r.imod.align.new
```

Fill in the relevant parameters, referring to the mdoc if necessary. Only parameters with *'s are compulsory to fill in.

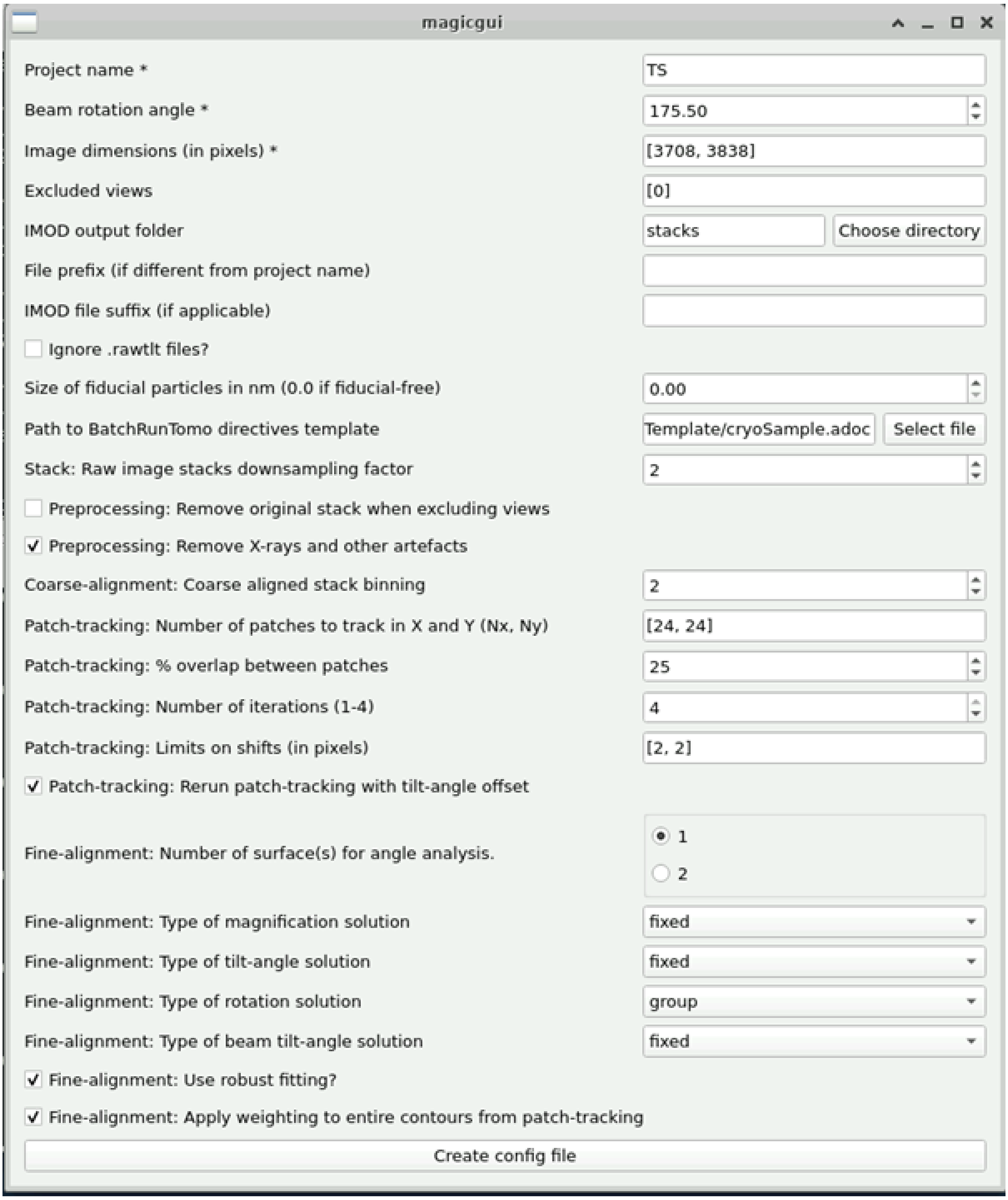


Again, create the config file and close the dialog box. Two files will be created:

- o2r_imod_align.log – holds the output from IMOD's alignment method
- TS_align.log – holds the configuration parameters from the input you just filled in.

Note that you may have to change the Path to BatchRunTomo directives template depending on where IMOD is installed on your system.

8. Run the alignment step. Ensure IMOD is loaded (RFI users: use `module load imod`)
   Run the Ot2Rec IMOD alignment plugin:

```
o2r.imod.align.run TS
```

IMOD will then create the stacks (.st files) from the motion corrected files, using the filepaths saved in the TS_mc2_mdout.yaml file from step 6. After that, alignment will start. You can monitor the process in the terminal. If you'd like to view your aligned tilt series, you can open the _ali.mrc files in IMOD's 3dmod or in Fiji.

9. Set up the IMOD reconstruction process.

```
o2r.imod.recon.new
```

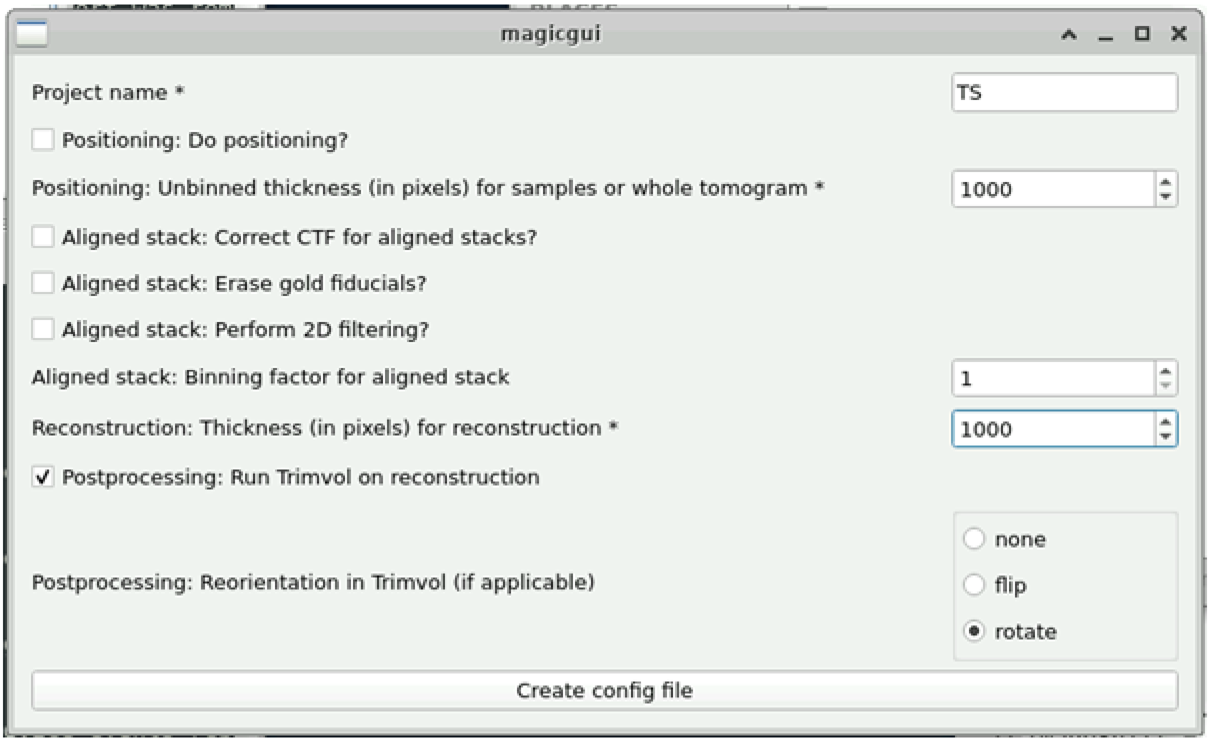


The thickness parameters determine how many z-slices will be in the reconstruction. Larger thickness values means you are less likely to inadvertently exclude regions of your specimen in the tomogram, but there might be lots of empty space in the tomogram. You can experiment with these parameters by doing a test run – generate the TS_recon.yaml file and edit the process list to only do one tilt series and see if you are happy with the result. Once you are, then you can generate a new TS_recon.yaml by rerunning this step with the parameters you have chosen.

10. Run the IMOD reconstruction process

```
o2r.imod.recon.run TS
```

11. Generate the Ot2Rec report. *(temporary fix – this step will be redundant in the future)*

Run `o2r.imod.align.stats` to generate the statistics for the alignment step to go into the report.
Activate the Ot2Rec report environment (see the Github for detailed instructions if required).

Generate the report by running:

```
o2r.report.run TS --to_html --to_slides
```

This generates the report as a html document that you can print to a pdf, or to a slide deck that you can also open in the browser. The report will be in your processing folder and it will be called "report.html" or "report.slides.html".

Movie S1: Video of 3D reconstructions from the various workflows in Case Study 3

Full animation available at DOI: 10.6084/m9.figshare.21732491

IMOD Alignment | AreTomo Alignment

IMOD SIRT

AreTomo SART

Savu CGLS

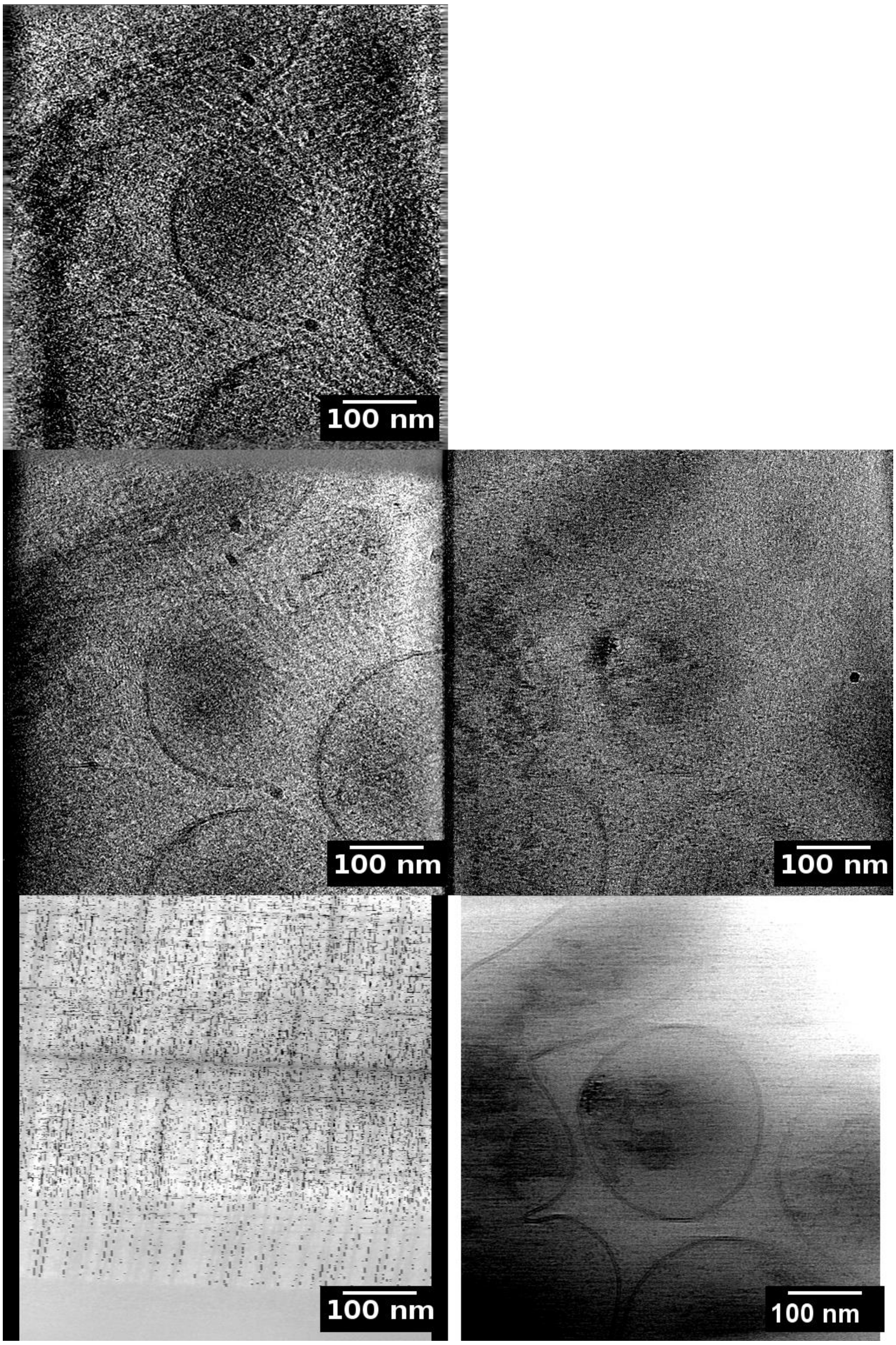
